# A near-complete human pancreatic cancer genome

**DOI:** 10.64898/2026.05.01.722316

**Authors:** Justin Wagner, Ayse G. Keskus, Keisuke K. Oshima, T. Rhyker Ranallo-Benavidez, Jennifer McDaniel, Shu-Cheng Chuang, Mile Sikic, Dehui Lin, Luis F. Paulin, Adam C. English, Fritz J. Sedlazeck, Elizabeth M. Munding, J. Zachary Sanborn, Andrew Carroll, Pi-Chuan Chang, Daniel E. Cook, Kishwar Shafin, Joep de Ligt, Rayan Hassaine, Daniel Cameron, Severine Catreux, Yeonghun Lee, Lisa Murray, Sean Truong, Christian Brueffer, Aleksey V. Zimin, Erin Cross, Matthew McGowan, Michael Vernich, Andrew S. Liss, Jean-Pierre Kocher, Zachary Stephens, Tanveer Ahmad, Asher Bryant, Nathan Dwarshuis, Hua-Jun He, Zhiyong He, Nathan D. Olson, Francoise Thibaud-Nissen, Dmitry Antipov, Sergey Koren, Adam Phillippy, Arang Rhie, Melissa Wilson, Rajeeva Lochan Musunuri, Giuseppe Narzisi, Miten Jain, Aaron M. Wenger, Stephen Eacker, Sayed Mohammad Ebrahim Sahraeian, Paul C. Boutros, Yash Patel, Takafumi N. Yamaguchi, Joseph McConnell, Matthew Borchers, Jennifer L. Gerton, Paxton Kostos, Tamara A. Potapova, Andrea Guarracino, Maryam Jehangir, Hila Benjamin, Mohammed Faizal Eeman Mootor, Yuan Xu, Mobin Asri, Karen H. Miga, Jimin Park, Benedict Paten, Ruibang Luo, Zhenxian Zheng, Jae Young Choi, Linh Nguyen, Pankaj Vats, Dan R. Robinson, Josh N. Vo, Shenghan Gao, Ghulam Murtaza, Andrew B. Stergachis, Jane Ranchalis, Christopher E. Mason, Haoyu Cheng, Yuichi Shiraishi, Yoshitaka Sakamoto, Danilo Dubocanin, Floris P. Barthel, Chunlin Xiao, Glennis A. Logsdon, Mikhail Kolmogorov, Justin M. Zook

**Affiliations:** National Institute of Standards and Technology, Material Measurement Laboratory, 100 Bureau Dr., Gaithersburg, MD 20899, USA; National Cancer Institute, Cancer Data Science Laboratory, 9000 Rockville Pike, Bethesda, MD 20892, USA; University of Pennsylvania, Department of Genetics, Epigenetics Institute, 3400 Civic Center Blvd., Smilow Center for Translational Research, Philadelphia, PA 19104, USA; Translational Genomics Research Institute, Bioinnovation and Genome Sciences, 445 N 5th St 4th Floor, Phoenix, AZ 85004, USA; A*STAR, Genome Institute of Singapore, 60 Biopolis St, Singapore 138672; Baylor College of Medicine (BCM), Human Genome Sequencing Center (HGSC), One Baylor Plaza, Houston, TX 77030, USA; Baylor College of Medicine (BCM) and Rice University, One Baylor Plaza, Houston, TX 77030, USA; Dovetail Genomics, part of Cantata Bio, Research and Development, 100 Enterprise Way, Suite A101, Scotts Valley, CA 95066, USA; Google Research, Google Inc., 1600 Amphitheatre Pkwy, Mountain View, CA, USA; Hartwig Medical Foundation, Data Services, Science Park 404, 1098 XH, Amsterdam, The Netherlands; Illumina, DRAGEN, 5200 Illumina Way, San Diego, CA 92122, USA; InSilico Consulting AB, Kulgränden 15B, SE-22649 Lund, Sweden; Johns Hopkins University, Department of Biomedical Engineering, Wyman Park Building S237, Baltimore, MD 21205, USA; KromaTiD, 6899 Winchester Cir Suite 100, Boulder, CO 80301, USA; Lund University, Division of Oncology, Department of Clinical Sciences Lund, Medicon Village, Scheeletorget 1, SE-22381 Lund, Sweden; Massachusetts General Hospital and Harvard Medical School, Department of Surgery, Boston, MA 02114, USA; Mayo Clinic, Department of Quantitative Health Sciences, 200 First Street SW, Rochester, MN 55905, USA; National Center for Biotechnology Information, National Library of Medicine, National Institutes of Health, 45 Center Drive, Bethesda, MD 20894, USA; Genome Informatics Section, Center for Genomics and Data Science Research, National Human Genome Research Institute, National Institutes of Health, Bethesda, MD, USA; New York Genome Center (NYGC), New York, NY 10013, USA; Northeastern University, Genome Technology Laboratory, 360 Huntington Ave, Boston, MA 02115, USA; PacBio, 1305 O’Brien Drive, Menlo Park, CA 94025, USA; Phase Genomics Inc., Seattle, WA, USA; Roche Sequencing Solutions, 2881 Scott Blvd, Santa Clara, CA 95050, USA; Sanford Burnham Prebys Medical Discovery Institute, Cancer Center, 10901 North Torrey Pines Road, La Jolla, CA 92037, USA; SA Pathology, Data and Bioinformatics Innovation, Department of Genetics and Molecular Pathology, Frome Rd., Adelaide, South Australia 5000, Australia; Stowers Institute, 1000 E. 50th Street, Kansas City, MO 64110, USA; Ultima Genomics Inc., 4425 Technology Dr, Fremont, CA 94538, USA; University of California Los Angeles (UCLA), Department of Human Genetics, Los Angeles, CA, USA; University of California Santa Cruz (UCSC), Department of Biomolecular Engineering, Santa Cruz, CA, USA; University of California Santa Cruz (UCSC), UC Santa Cruz Genomics Institute, Santa Cruz, CA, USA; University of Hong Kong, School of Computing and Data Science, 301 Chow Yei Ching Bldg., HKU, Pokfulam Rd., Hong Kong; University of Kansas, Department of Ecology and Evolutionary Biology, Haworth Hall, 1200 Sunnyside Ave, Lawrence, KS 66045, USA; NVIDIA Corporation, 2788 San Tomas Expressway, Santa Clara, CA 95051; University of Michigan-Ann Arbor, Department of Pathology, 1500 E Medical Center Dr, Ann Arbor, MI 48109, USA; University of Washington, Department of Genome Sciences, William H. Foege Hall, 3720 15th Ave NE, Seattle, WA 98195, USA; Weill Cornell Medicine, Department of Systems and Computational Biomedicine, 1300 York Ave, New York, NY 10065, USA; Yale School of Medicine, Department of Biomedical Informatics and Data Science, New Haven, CT, USA; National Cancer Center Research Institute, Division of Genome Analysis Platform Development, 5-1-1 Tsukiji, Chuo-ku, Tokyo 104-0045, Japan; National Human Genome Research Institute, National Institutes of Health, Comparative Genomics and Reproductive Health Section, Center for Genomics and Data Science Research, Bethesda, MD, USA; Stanford University, Department of Genetics, Stanford, CA 94305, USA; University of Washington, Division of Medical Genetics, Department of Medicine, Seattle, WA 98195, USA; NIH Intramural Sequencing Center, National Institutes of Health, Rockville, MD 20852, USA

**Author notes:** Corresponding author: Justin Zook. These authors contributed equally (Justin Wagner, Ayse G. Keskus, Keisuke K. Oshima, T. Rhyker Ranallo-Benavidez, Jennifer McDaniel). Co-senior authors (Floris P. Barthel, Chunlin Xiao, Glennis A. Logsdon, Mikhail Kolmogorov, Justin M. Zook).

## Abstract

Genomic changes are central to cancer, so sequencing is essential for understanding tumor evolution and advancing precision medicine, but resolving somatic variants and methylation is challenging in regions with repeats, reference gaps, and germline variants.^1–4^ To identify the full spectrum of somatic variants, including complex rearrangements, we construct and curate near-complete, haplotype-resolved assemblies of the most recent common ancestor of an early-passage, broadly-consented hypodiploid pancreatic cancer cell line and matched normal tissues.^5^ The tumor assembly recapitulates all 35 chromosomes observed by karyotyping, including translocation-induced hybrid chromosomes containing dynamic dicentric and fused centromeres, nested foldback inversions causing 14 breakpoints with a haplotype switch in a single derivative chromosome, and centromeric satellite tandem duplications up to 136 kbp. Our tumor-normal assembly-based benchmark expands previous multi-technology ensemble benchmark approaches, adding >8,000 somatic small variants altering ∼40 kbp and 149 somatic SVs altering 939 kbp, including 2,764 small variants and 126 SVs not represented on GRCh38. Additionally, 5,111 somatic small variants change representation because they alter germline variants on GRCh38, impacting mutational signatures. Most source-resolved somatic LINE insertions originate from two non-reference germline LINE insertions with accessible promoter elements. These assemblies demonstrate that centromeric, acrocentric, telomeric, and other polymorphic regions, conventionally excluded from analysis, harbor extensive somatic and epigenetic changes and provide curated paired tumor-normal benchmarks to advance algorithms for repetitive regions of cancer genomes.

## Introduction

Tumor genomes often have complex somatic changes relative to the germline genome from which they evolve.^1^ It is common for tumor cells to gain or lose chromosome arms due to somatic structural changes that occur inside highly repetitive satellite DNA sequences in the centromeres.^4^ These repetitive regions in normal genomes have only recently been tractable to resolve with the advent of complete “Telomere-to-Telomere” (T2T) assemblies of effectively haploid^6^ and diploid genomes.^7^ For somatic variants, mapping tumor and normal reads to a donor-specific assembly (DSA, or a haplotype-resolved assembly of the normal genome) has been shown to improve somatic variant calling and epigenetic analysis.^2,3,8,9^ DSAs expand prior simulated^10,11^ and tumor/normal benchmarks for legacy cell lines^12–14^ that largely excluded repetitive regions, lacked phasing, and focused on one variant type. While normal DSAs are now possible, tumor genome assemblies are fragmented with current algorithms due to subclonal variation between cells and large, nearly-identical duplication events.

Here, we use extensive genomic data from a hypodiploid tumor cell line and normal pancreatic and duodenal tissues from the same recently broadly-consented germline XX individual with pancreatic ductal adenocarcinoma (PDAC).^5,15^ In this genome from HG008, we previously reported clonal somatic variants in *KRAS*, *TP53*, *SMAD4*, and *CDKN2A*, four of the most commonly mutated genes in PDAC.^5^ With recent advances in telomere-to-telomere assembly, we investigated generating an assembly of a tumor and normal matched sample using a recipe of Hi-C, approximately 17x and 58x HiFi for normal and tumor, respectively, and 16x standard ONT for normal and 27x ultralong ONT for tumor (coverages per haplotype). Here, we generate phased benchmarks for all variant types from near-complete haplotype-resolved assemblies of the normal genome and the genome of the most recent common ancestor of the tumor cell line. With extensive curation and polishing, these assemblies precisely resolved phased small variants, structural variants, and copy number variants, including complex rearrangements involving multiple chromosomes, translocations with breakpoints in centromeres, and repeat expansions and contractions in regions missing from the reference genome. We establish a curated, near-complete tumor–normal assembly-based benchmark and use it to quantify somatic variation omitted or misrepresented by mapping-based approaches, while resolving the sequence and chromatin organization of rearranged cancer centromeres.

## Results

### Assemblies recover truncal SVs and CNVs

To assess tractability and structural accuracy of HG008 tumor genome assembly, we used the directional genomic hybridization (dGH) SCREEN assay^16^ (Fig. 1a). Almost all 80 cells from passages 21 and 53 shared common structural changes matching assembly-based copy number profiles and structural variants (Supplementary Table S1), apart from subclonal variants occurring in 1 to 4 cells (like the chr19 segment in der(18)t(18;19;X)(q;?;q) in Fig. 1a and Supplementary Table S2). While subclonal variants differ between passages, the variants characterized in this work are relatively stable and in most cells across passages. Thus, the mutations are truncal (*i.e.* occur in the trunk of the evolutionary tree, leading to the <u>m</u>ost <u>r</u>ecent <u>c</u>ommon <u>a</u>ncestor, MRCA). Most are also clonal (i.e. present in all cells), but a subset are subclonal due to loss in descendent passages, mostly due to large deletions.^11^

**Fig. 1:**
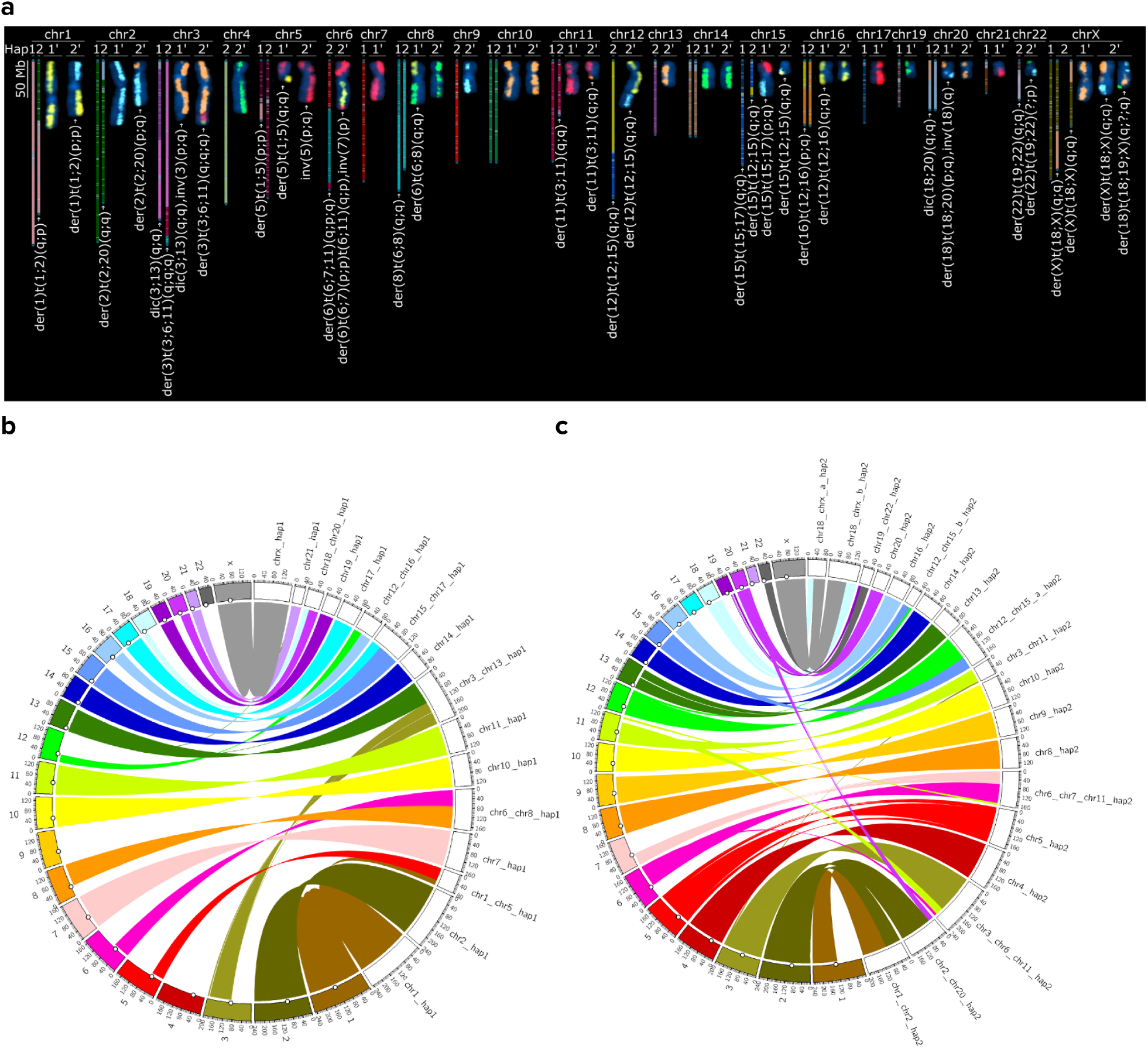
Near-complete haplotype-resolved tumor and normal assemblies. (a) KaryoScope visualization of the near-complete tumor assembly matches all chromosomes consistently seen in dGH SCREEN karyotyping (representative cell shown), which paints one strand of each chromosome one of five distinct colors. For each chromosome, KaryoScope is on the left and dGH is on the right, including International System for Human Cytogenomic Nomenclature (ISCN) notations. (b-c) Alignments of tumor and normal assembly haplotypes (right and left sides of each circos plot, respectively) reveal large chromosomal-scale deletions, duplications, and translocations. Large chromosome-scale CNVs >10Mbp were caused by loss of whole chromosomes (chr4_hap1, chr9_hap1, chr17_hap2, chr21_hap2, and chr22_hap1) or by unbalanced translocations between chromosomes (e.g. chr3p, chr5q, chr6q, chr8p, chr12q, chr16p, and chr18q in hap1, and chr1p, chr6p and chr7q in hap2).

We generated haplotype-resolved, near-complete assemblies of the MRCA of the hypodiploid HG008 PDAC tumor cell line and its matched normal genome by adapting T2T assembly workflows.^17^ By curating the assembly graph, we achieved T2T scaffolds for all 35 tumor chromosomes with 8 gaps (all due to rDNA arrays) and 44 T2T scaffolds for the normal genome. Systematic comparison of dGH to assembly-based karyotyping using KaryoScope^18^ showed complete concordance in translocation partner identification across all 16 truncal inter-chromosomal rearrangements, including unidentified ‘marker’ chromosomes from standard G-banded karyotyping (Table S2). Polishing using Element^19,20^ and Onso^21^ short reads fixed > 100,000 indel errors, particularly in homopolymers.^7^ Validation through k-mers, Hi-C mapping, haplotype-specific markers, and gene-completeness analysis confirmed high structural and phasing accuracy (Supplementary Figures 1-4). While some errors remained, particularly around gaps due to rDNA arrays, these assemblies captured all truncal variation detected by karyotyping and reference-based approaches, providing a comprehensive, high-resolution foundation for identifying clonal somatic variants and tumor chromosome structure (Fig. 1).

The assemblies completely resolve truncal somatic CNVs and the translocations causing them, including complex, centromeric, and acrocentric breakpoints typically left unresolved (Supplementary Fig 5).^4^ The prevalence of large deletions in HG008 (Extended Data Fig. 1) aligns closely with chromosomes and chromosome arms commonly lost in PDAC and in the chromosome-scale LOH signature from TCGA.^22^ Translocations also caused three large duplications or CNLOH events (Supplementary Fig 6). The assembly phases all events, including loss of each chr6 arm on opposite haplotypes, deleting a copy of the MHC region, which may affect immunogenicity (Supplementary Table S3).^23^ Phasing also elucidates a balanced chr18_hap2/chrX_hap2 translocation on the copy of chrX that is mostly transcriptomically silenced (Supplementary Fig 7). Interchromosomal translocations in HG008 are frequently complex in the sense that they cannot be described as a single pair of breakpoints, commonly due to non-reciprocal inversions at the breakpoint. In addition to three reciprocal inversions, we resolve nine non-reciprocal foldback inversions related to complex interchromosomal translocations, five inversions that do not cause duplications near translocations, and one inversion related to a tandem duplication (Supplementary Fig 8).

The tumor assembly resolved several translocations involving multiple chromosomes. For example, chromoplexy resulted in a complex series of translocations involving chromosomes 3, 6, 7, and 11 to form 3 new hybrid chromosomes (Extended Data Fig 2). The most complex somatic structural variation and copy number variation occurred on the parts of two haplotypes of chr19 attached to most of chr22 (Fig. 2). The translocation between chr22 and chr19 occurred with a foldback inversion in the pericentromere of chr22, between the rDNA and centromere on the normal assembly. On chr19, the assembly resolves a nested series of foldback inversions with 11 breakpoints causing copy numbers from 1 to 5, consistent with multiple breakage-fusion-bridge (BFB) cycles, and stabilization by telomere capture with the last 8 Mbp of chr19 coming from the opposite haplotype (Supplementary Fig 2c). The resulting CNLOH has the imprinted gene *ZNF331* remaining unmethylated on both copies and expressed but the *PEG3* paternal allele changing from unmethylated in the normal tissues to methylated in the tumor cell line. The assembly resolves another acrocentric short arm translocation between chr15 and chr17, with 210 bp of chr11 in between (Supplementary Fig 9). All of these large translocations were confirmed by Hi-C contact maps and curation, forming a comprehensive benchmark (Supplementary Fig 10).

**Fig. 2:**
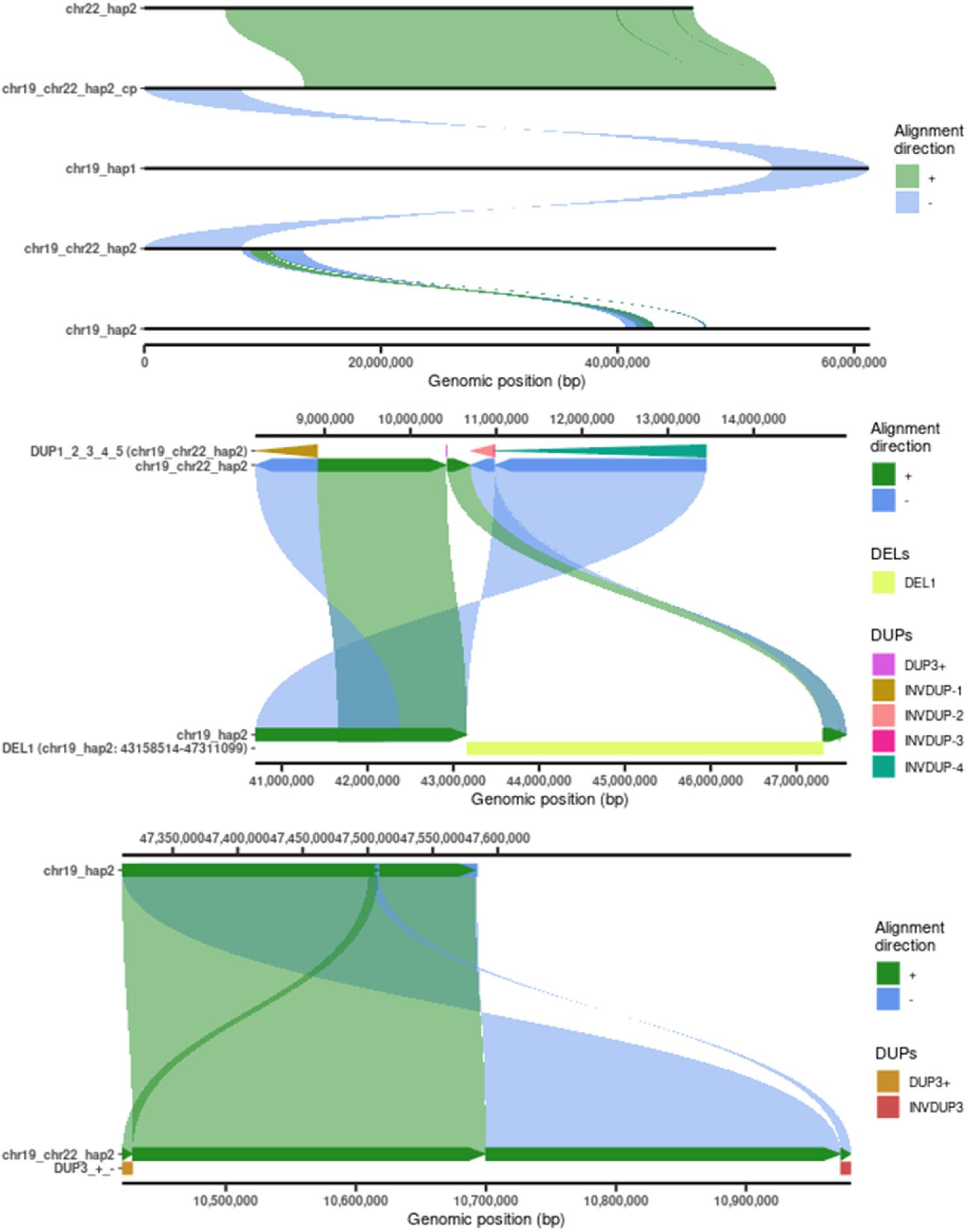
The chr19_chr22_hap2 hybrid tumor chromosome assembly reveals complex foldback inversions at multiple resolutions, plus a haplotype switch. der(22)t(19;22)(q;p) has a breakpoint in the acrocentric short arm of chr22, causing loss of 6.8 Mbp of the chr22 acrocentric short arm including the rDNA array and distal junction, as well as a 97 kbp foldback inverted duplication of chr22 adjacent to the centromere. It is connected to a series of nested foldback inversions of chr19_hap2, before being connected to the last 8 Mbp of chr19_hap1, causing CNLOH. The assembly resolves high-level structure (top) as well as complex inverted duplications of 8 kbp to 2.5 Mbp fragments at basepair resolution (middle and bottom).

### Variant types enriched in distinct repeats

Directly comparing the tumor and normal assembly haplotypes gives a more precise and comprehensive view into how different types and sizes of somatic variants are associated with repetitive genomic regions (Fig. 3). In particular, assemblies reduce biases caused by germline variation and missing reference sequences in repetitive regions. These variants are discovered by comparing tumor and normal assembly haplotypes, and independently confirmed by support from tumor reads and not normal reads aligned to the normal assembly. All variant types except tandem duplications > 100 kbp are enriched in repetitive regions (Fig. 3 and Supplementary Fig 11). Somatic insertions of all size ranges are enriched in repetitive regions, with >96% of insertions from 1 bp to 100 kbp occurring in homopolymers, tandem repeats, or centromeric satellites, and only LINE-1 insertions occurring mostly in non-repetitive regions (Fig. 3c). Similar to germline variants, while most truncal somatic variants are small (21,608 variants <50 bp, modifying 27,457 bp), the fraction of the genome modified by large somatic variants is higher (214 variants ≥50bp, modifying 27,971,033 bp, not including large CNVs caused by 86 translocation and inversion breakpoints) (Supplementary Fig 11). Insertion:deletion ratios are a complex function of repeat unit size, repeat length, and variant type (Fig 3 and Supplementary Figs 11-12).

**Fig. 3:**
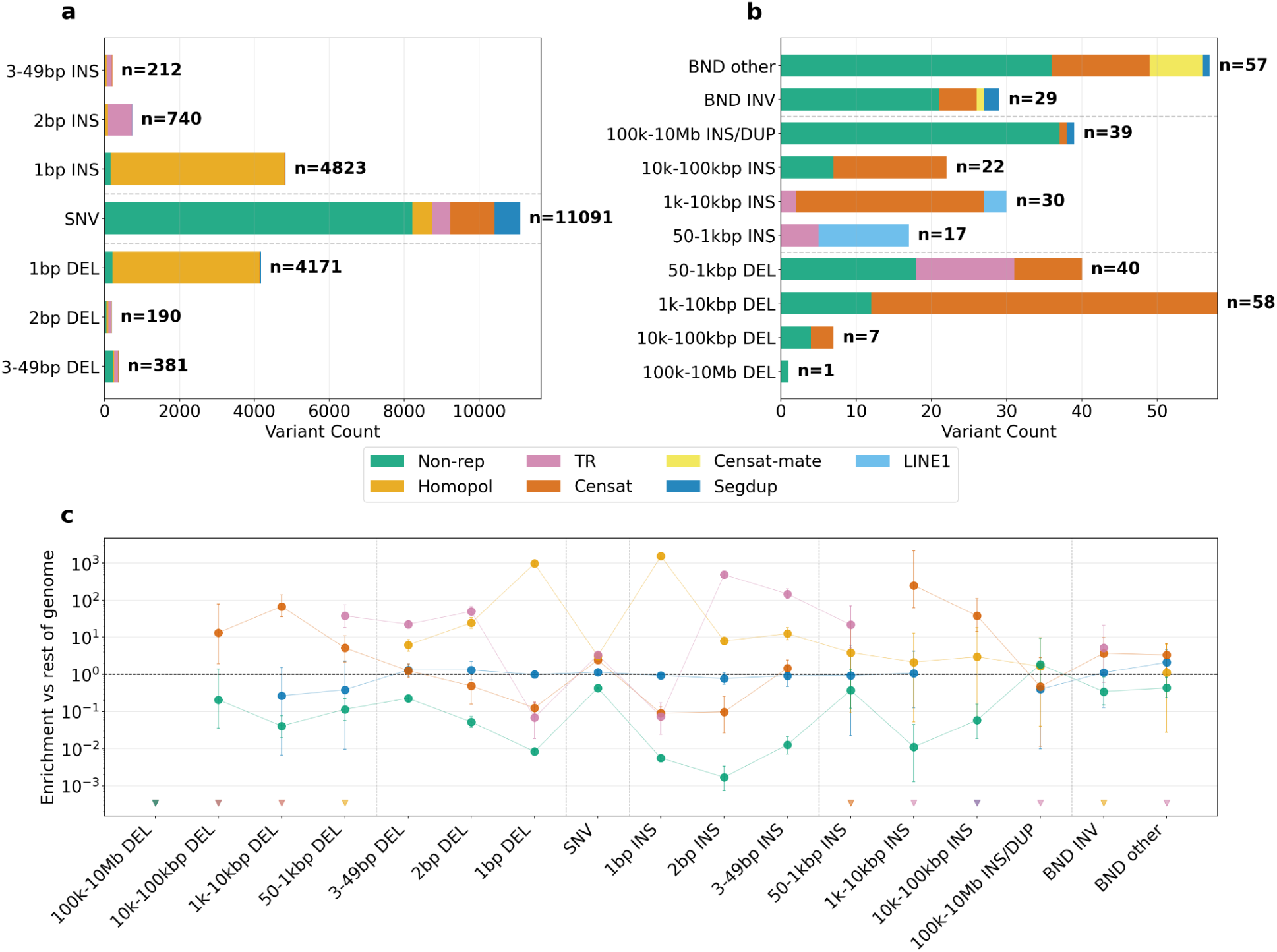
Somatic variant types are enriched in different repetitive regions, based on confirmed truncal variants derived from comparing the tumor and normal assemblies. (a) Truncal small variant counts show substantial numbers of SNVs in repetitive regions and most indels in homopolymers and TRs. (b) Truncal SV counts show many SVs in repetitive regions, particularly for 50 bp to 100 kbp insertions. (c) Variants across types and sizes are generally enriched in repetitive regions relative to the whole genome. Categories are mutually exclusive in panels a and b, but not c. Error bars are exact 95% confidence intervals for the ratio of two Poisson rates (conditional binomial, Clopper-Pearson). Triangles indicate no variants. Dashed lines separate variant types.

While most variant types are associated with particular repeat types, large somatic deletions and tandem duplications that could be represented on GRCh38 tend to have simple breakpoints in non-repetitive regions. Important genes affected by these variants include a 20 kb deletion of the remaining copy of *CDKN2A*, a 2 Mbp tandem duplication of the *KRAS* G12V mutated gene that was associated with advanced disease,^24^ and Internal tandem duplications in *NFE2L2*, *UBR3*, and *BBS9*, with RNA-seq support for the *UBR3* event (Extended Data Fig 3). For other variant types, repeat structures dictate the types of mutations that occur and the challenges in detecting them, so we next investigated the challenges and potential biological relevance within each major repeat class.

### Four centromere types in tumor chromosomes

We examine centromere structure in the rearranged HG008 tumor chromosomes because centromeres have long been implicated in aneuploidy^25^ and chromosome arm loss and were recently shown to have higher somatic mutation rates.^2,3^ Of 16 hybrid tumor chromosomes composed of multiple normal chromosomes, 9 clearly retained only one centromere with breakpoints outside the centromeres (2 reciprocal translocation pairs in Supplementary Fig 13 and 5 non-reciprocal translocations in Extended Data Fig. 2 and Supplementary Fig 9 and 14). However, 7 had a breakpoint in or near one or both centromeres (Supplementary Table S4-7), causing chromosome arm losses in HG008-T that are common in all cancers and/or the chromosome-scale LOH signature (*e.g.*, chromosomes 1p, 3p, 6p, 6q, 8p, 18q) in TCGA samples^22^ and in PDAC samples (6p, 6q, 8p, 18q).^25^

To determine if these 7 chromosomes had formed putative functional dicentrics, we identified all centromere dip regions (CDRs), which are associated with kinetochore function.^26^ We further performed CENP-A DiMeLo-seq and long-read CUT&RUN to identify CENP-A chromatin domains, which serve as the foundation for the assembly of the kinetochore. We found that one chromosome formed a putative functional dicentric in the batch of cells used for de novo assembly, with the chr6 and chr7 CDRs located ∼1 Mbp apart in the α-satellite HOR array (**Fig. 4a, Extended Data Fig. 4**). The assembly-source cell culture contained two CDRs, consistent with a putatively dicentric state. In an independent culture, the chr6 region was methylated and lacked detectable CENP-A enrichment, whereas the chr7 region retained centromeric chromatin, resulting in the formation of a monocentric similar to that observed in some Robertsonian translocations,^27^ and highlighting the dynamic nature of this centromere. We further analyzed Fiber-seq for dichromatin organization, which is associated with kinetochore function, across 3 clones of HG008-T and identified dichromatin around the chr7 CDR but not the chr6 CDR in the clone that lost the chr6 CDR (**Extended Data Fig. 5**). We also observed another chromosome that had fused within both chr12 and chr16 CDRs, resulting in a single CDR site with CENP-A chromatin enrichment composed of two different α-satellite HOR arrays (**Fig. 4b, Supplementary Figs. 14 and 15**). We identified three pseudodicentric chromosomes with one functional centromere and one inactivated centromere. The chr8 CDR is deleted by the interchromosomal translocation (**Fig. 4c**). Large inversions related to the translocation delete the CDR in chr3, but do not delete the CDR in chr18 so its CDR is inactivated by methylation and does not have CENP-A signal (**Extended Data Fig. 6 and Supplementary Figs. 8, 16-18).** Two monocentric chromosomes had breakpoints within the chr1 α-satellite HOR or immediately adjacent to the chr22 α-satellite HOR but preserved the CDRs (**Supplementary Figs. 19-22**). Together, we observe four different types of centromeres in the tumor chromosomes with translocations: one dynamic dicentric, one hybrid centromere, three pseudodicentrics with different causes of centromere inactivation, and eleven monocentrics, underscoring the extreme plasticity of human centromeres in order to preserve function by modulating either genetic or epigenetic landscapes.

**Fig. 4.**
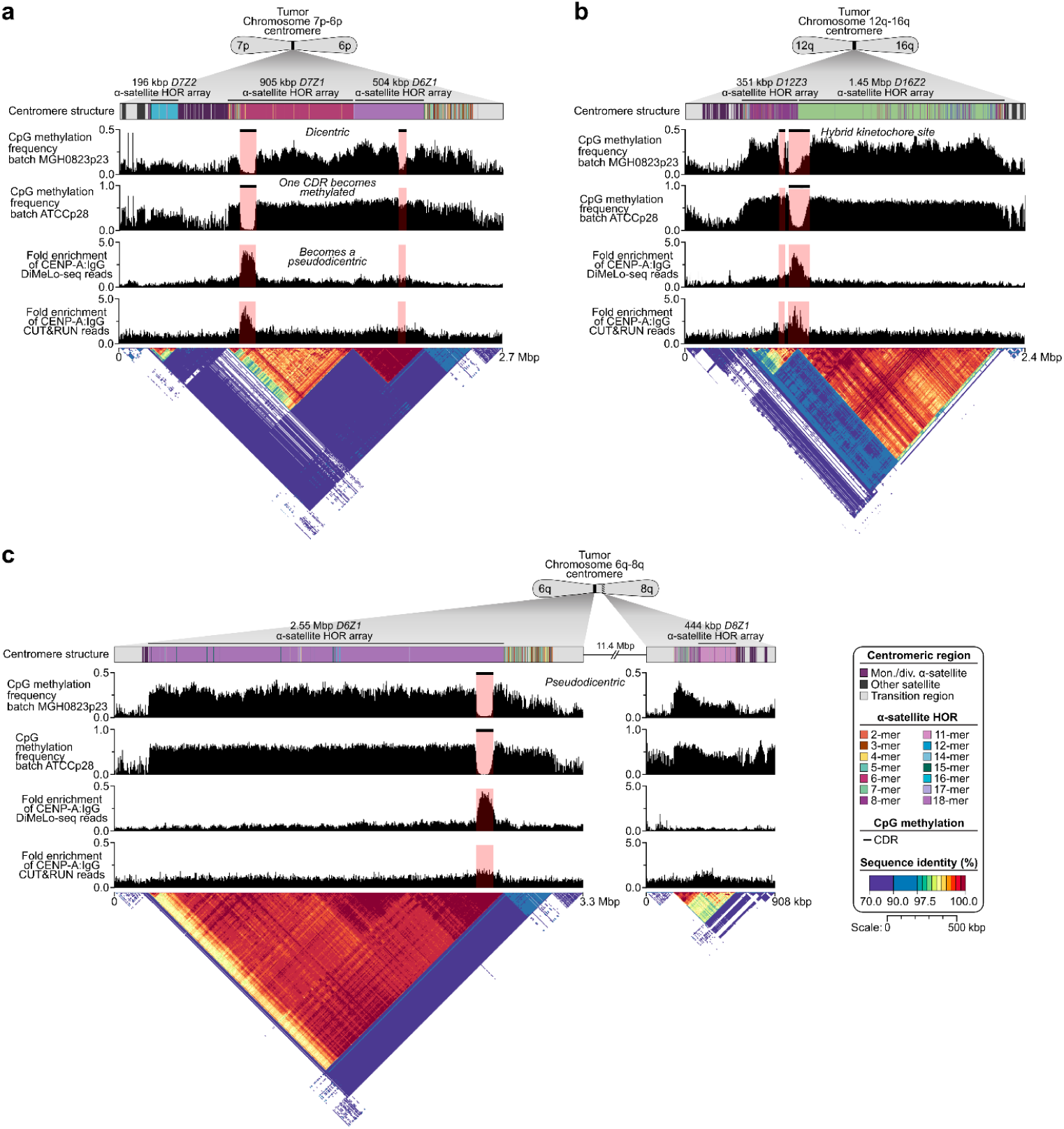
Plasticity of human centromeres during tumorigenesis. The top methylation track matches cell batch MGH0823p23 used for the assembly, whereas the second track matches the ATCC p28 cell batch used for CENP-A DiMeLo-seq and CUT&RUN measurements of kinetochore protein binding. CDRs are low methylation regions associated with the site of kinetochore assembly, highlighted by black bars and pink rectangles in the CpG methylation track. a) A dynamic dicentric chromosome is formed upon fusion of 6p and 7p, resulting in two CDRs within the *D7Z1* and *D6Z1* α-satellite HOR arrays in one batch of tumor cells, but only a single CDR with CENP-A binding in *D7Z1* in a different batch of cells. b) A monocentric chromosome is formed when chromosomes 12q and 16q fuse, creating a hybrid CDR with CENP-A that resides on two different types of α-satellite HOR arrays (*D12Z3* and *D16Z2*). c) A pseudodicentric chromosome is formed upon translocation of 6q and 8q, resulting in an active CDR site and CENP-A on the *D6Z1* α-satellite HOR array and an inactivated CDR site on the *D8Z1* α-satellite HOR array. The inactivation of the *D8Z1* α-satellite HOR is caused by the translocation that removes ∼2 Mbp of the *D8Z1* α-satellite HOR array.

### Assemblies reveal centromeric deletions and duplications

The HG008 assemblies enabled precise resolution of many translocations, insertions, and deletions in centromere satellites, where most 1-100 kbp SVs occur. 300 SVs were confirmed as likely true truncal somatic SVs by most tumor but not normal PacBio HiFi or ONT reads aligned to the combined haplotypes. Chromosome 1 is particularly rich in challenging SVs with 24 of 27 truncal SVs in satellite repeats, including insertions in both haplotypes of a small HSat3 array and a deletion of one monomer in a 5S rDNA array outside the centromere (Supplementary Fig 23).

Analysis of all 100 somatic insertions and deletions in centromeric satellite DNA regions of HG008-T revealed that somatic insertions were significantly larger than deletions (Supplementary Fig 24). The four largest confirmed somatic insertions of 62 kbp to 136 kbp are larger than germline or somatic de novo centromeric insertions seen in recent studies (Supplementary Fig 25). As expected, smaller somatic SVs < 2 kb in α-satellite HOR arrays showed strong periodicity at 171 bp intervals (Supplementary Fig 24). Of 39 CenSat insertions larger than 2 kb, 37 were simple tandem duplications, one was a complex duplication in HSat2, and one was an L1 insertion (Supplementary Table S8 and Supplementary Fig. 26).

Small somatic variants that could not be lifted to GRCh38 were enriched for SNVs vs. small indels, likely because centromeric satellites contain fewer homopolymers. Similar to what was found recently for UV-damage somatic SNVs in a melanoma cell line,^2^ somatic SNVs in HG008 (not from UV damage) were also enriched in centromeric satellites relative to non-repetitive regions, particularly in CDRs of α-satellite HORs, dHORs, HSat2, and HSat3 (Supplementary Fig 27). Mutational profiles were mostly similar between HORs and non-repetitive regions, but those in HSat2 were significantly different, primarily due to differences in sequence composition (Fig. 5a and Supplementary Fig 28). All nine putative kataegis clusters of 6 or more variants with intermutation distance ≤1 kbp outside centromeric satellites showed hallmarks of APOBEC kataegis (strand-coordinated C>N substitutions in each cluster except chr13_hap2, with 65 of 88 in TCW base context, Supplementary Fig 29). In contrast, none of the nine clusters within centromeric satellites had hallmarks of APOBEC: two were suboptimal alignments around SVs, and in the other seven the tumor alleles matched other copies in the same array due to possible gene conversion or complex repair pathways. These likely differing mutational processes highlight the importance of refining the definition of kataegis in centromeres.

**Fig. 5:**
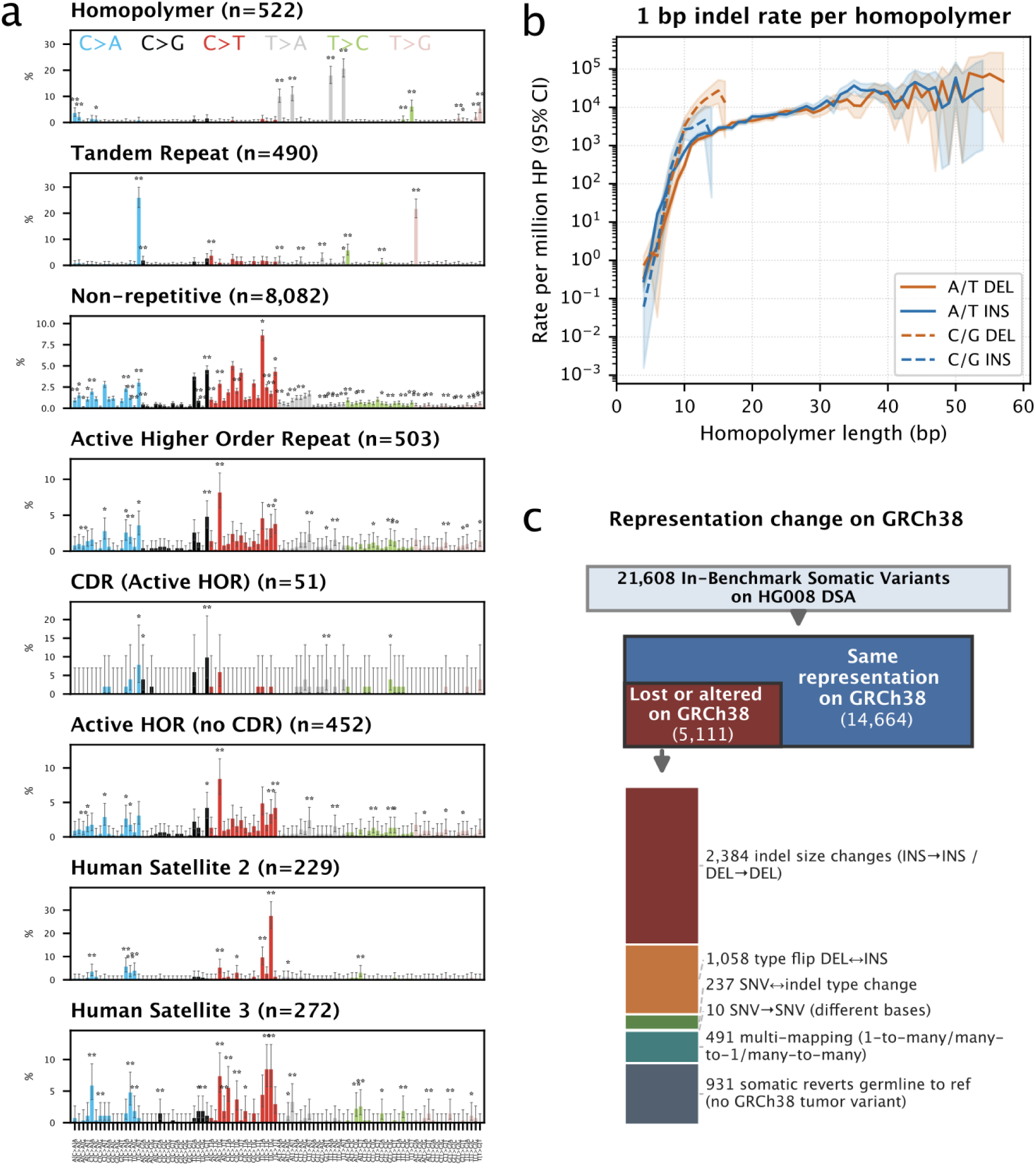
Challenging Somatic Small Variants. (a) 96-channel SBS spectrum varies across some repetitive regions using the DSA. Error bars, 95% Wilson CIs. * BH-FDR q<0.05, ** q<0.01 (two-sided exact conditional binomial test of the channel’s SNV rate per bp vs the rest of the genome, corrected across all 768 region × channel tests). (b) Truncal somatic 1 bp INDEL rate vs. A/T and C/G homopolymer length with 95% confidence interval shaded. (c) Many somatic small variants are lost or have altered representation on GRCh38 vs the true somatic change on the DSA due to germline variants.

### Germline variants hide somatic variants in repeats

Somatic variant calling in homopolymers and tandem repeats faces multiple challenges caused by high rates of germline variants, somatic mutations, systematic sequencing errors, and mapping errors. For example, all STRs and VNTRs with somatic SVs also have large germline insertions and/or deletions in HG008, so that the true somatic variant size is not apparent on GRCh38, and four also have evidence for mosaicism in the normal tissues. In addition to TR expansions and contractions, breakpoints on one end of a complex 36 Mbp somatic inversion on chr7 fall inside a 611 bp VNTR that also has germline indels. Some somatic small variants are hidden inside germline insertions in TRs, including a stop-gain somatic SNV near the end of a 63 bp germline insertion in a VNTR in *PRB4*. Multiple VNTRs like the one in *PRB4* are also located inside segmental duplications, causing additional mapping challenges (more examples in Supplementary Figs 30-33).

Accurate truncal small variants in homopolymers and TRs were identified by requiring support in accurate short reads and/or long reads from the tumor but not normal when mapped to the DSA. Deletions are enriched in very short and very long G/C homopolymers, whereas insertions are enriched in short and 35-36 bp A/T homopolymers and dinucleotide TRs (Fig. 5b, Extended Data Fig 7). In addition, most large indels inside homopolymers are deletions in long homopolymers (Supplementary Fig 33).

Variants are missed by reference-based methods because they cannot be lifted to GRCh38 (such as in centromeres and telomeres), fall in regions with germline CNVs, revert a germline variant to reference, change type or size due to germline variation, or fall in difficult regions (Supplementary Figs 34-37). 5,111 of 19,775 variants in GRCh38 benchmark regions changed type, size, or reverted to reference (Fig. 5c, Supplementary Fig 38). These changes can affect mutational signatures in repeats (Fig 5a, Supplementary Figs 28, 39-41). The reversions to reference are generally consistent with the number expected by somatic and germline 1 bp indel rates in homopolymers (Supplementary Table S9). While most chromosomes had similar somatic rates between haplotypes present in the tumor, the inactive chrX_hap2 has 1.32 to 1.72-fold as many SNVs and 1.16 to 1.50-fold as many indels (exact binomial 95% CIs), consistent with previous studies showing hypermutation of inactive X in cancer and lymphoblastoid cell lines.^28,29^

### Somatic LINEs from two germline LINEs

The 14 insertion SVs outside tandem duplications, tandem repeats, and centromeric satellites were all LINE retrotranspositions, plus one L1 in an α-satellite (Supplementary Table S10). Strikingly, 6 of 9 with an identifiable source were derived from two germline L1 insertions not in GRCh38: five from the chr14 source that has a population frequency of 14 % and one from chr15 that has 5.3 % population frequency in CoLoRSdb long reads.^30^ This suggests that germline ‘non-reference’ L1 insertions were particularly important for this individual’s somatic L1 insertion rate. The germline L1s and the other identifiable source L1’s on chr11 and chr22 had hypomethylated promoters in tumor but not normal reads (Extended Data Fig. 8 and Supplementary Fig 42). We performed Fiber-seq on 3 tumor cell line clones, confirming that these hypomethylated promoters contained significantly increased local chromatin accessibility relative to other full length L1s with both intact ORFs (Extended Data Fig. 8), consistent with these promoters being active. Six additional L1 somatic insertions were detected where the source L1 could not be determined. The somatic L1 insertions have modal poly-A tail lengths ranging from 29 bp to 128 bp, and many have highly variable lengths between cells based on HiFi reads (which might be caused by rapid shortening^31^), with the largest range from 26 bp to 152 bp for an SV with a bimodal distribution (Supplementary Fig 43). Some poly-A’s are substantially longer than a previously reported germline L1S distribution of 10 bp to 65 bp,^32^ similar to those seen recent disease-causing insertions.^33^ Of note, transduction sequences of somatic L1 insertions with interspersed poly-A tracts reveal not only the source L1 element, but also occasionally past lineages. In one example, the source L1 on chr15 was itself likely derived from either chr6 or chrX. One somatic L1 insertion lies within the centromeric α-satellite HOR (400 kbp upstream of the CDR) on chr1_hap2, so it is only detected using the DSA. While germline L1 insertions have recently been identified across most chromosomes’ α-satellite HORs, they are relatively rare on chr1.^34^

### Telomeres erode to distal TVR, with two translocations

Somatic changes are particularly prominent in the telomeres. While the allele telomere lengths in the pancreatic normal tissue are 6991 ± 2134 bp (1 SD, n=91), the telomeres of the tumor cell line are almost completely eroded, with lengths of 549 ± 216 bp (1 SD, n=68) (Extended Data Fig. 9 and Supplementary Fig 44). Interestingly, most of the telomeres in the tumor cell line eroded down to the most distal telomere variant repeat (TVR) but not significantly further, suggesting that telomeres effectively begin at the canonical tract and that the preceding regions comprised of canonical TTAGGG repeats interspersed with TVRs may not have telomeric function. However, TVRs were observed to be frequently altered, with an enrichment of somatic deletions up to 182 bp in size affecting TVRs, possibly attributable to somatic restructuring during cycles of telomere shortening and elongation.^35^ Telomeric translocations that may be related to shortened telomeres occur at the chr13q telomere with chr3, and in the chr15p acrocentric short arm telomere with chr17.

### Ribosomal DNA landscape is altered in the tumor cell line

The human genome contains hundreds of ribosomal RNA (rRNA) gene copies arranged in ribosomal DNA (rDNA) tandem arrays, which exist as two functionally distinct families: the 45S rDNA arrays, located on the short arms of the five acrocentric chromosomes and the 5S rDNA array, located on chromosome 1. Their repetitive structure has historically made rDNA arrays difficult to assemble, but new T2T genomes have now resolved the 5S array and substantially improved resolution of the 45S arrays.^6^ Here, we examine the 45S and 5S arrays in the context of cancer, because proliferation depends heavily on ribosome output.

The 45S rDNA arrays have been proposed as recombinational hotspots in cancer genomes,^36^ and loss of 45S rDNA copies in cancer has been previously reported.^37,38^ The total copy number estimate from short-read WGS indicated that the HG008 tumor cell line lost nearly half of its 45S rDNA copies relative to the two normal tissues (Fig 6A). rDNA loss was accompanied by loss of the distal junction (DJ), a large, inverted repeat element distal to 45S arrays on acrocentric chromosomes, indicating that an entire genomic region containing an rDNA array was lost (Fig 6A, B).^39^ Cytogenetic analysis with rDNA FISH probes quantified each array copy number (Fig 6C) and confirmed the assembly-based result that this copy number reduction derived from the loss of three rDNA arrays: loss of one entire copy of chromosomes 21 and 22, and loss of other chr22 array in chr19-chr22 BFB event (Figs 2 and 6B, Supplementary FIg. 45). Notably, the structural rearrangement of one copy of chromosome 15 involved terminal acrocentric p-arm fusion with a fragment of chromosome 17, preserving its rDNA array and DJ regions on this rearranged chromosome (Supplementary Fig. 45).

**Fig. 6.**
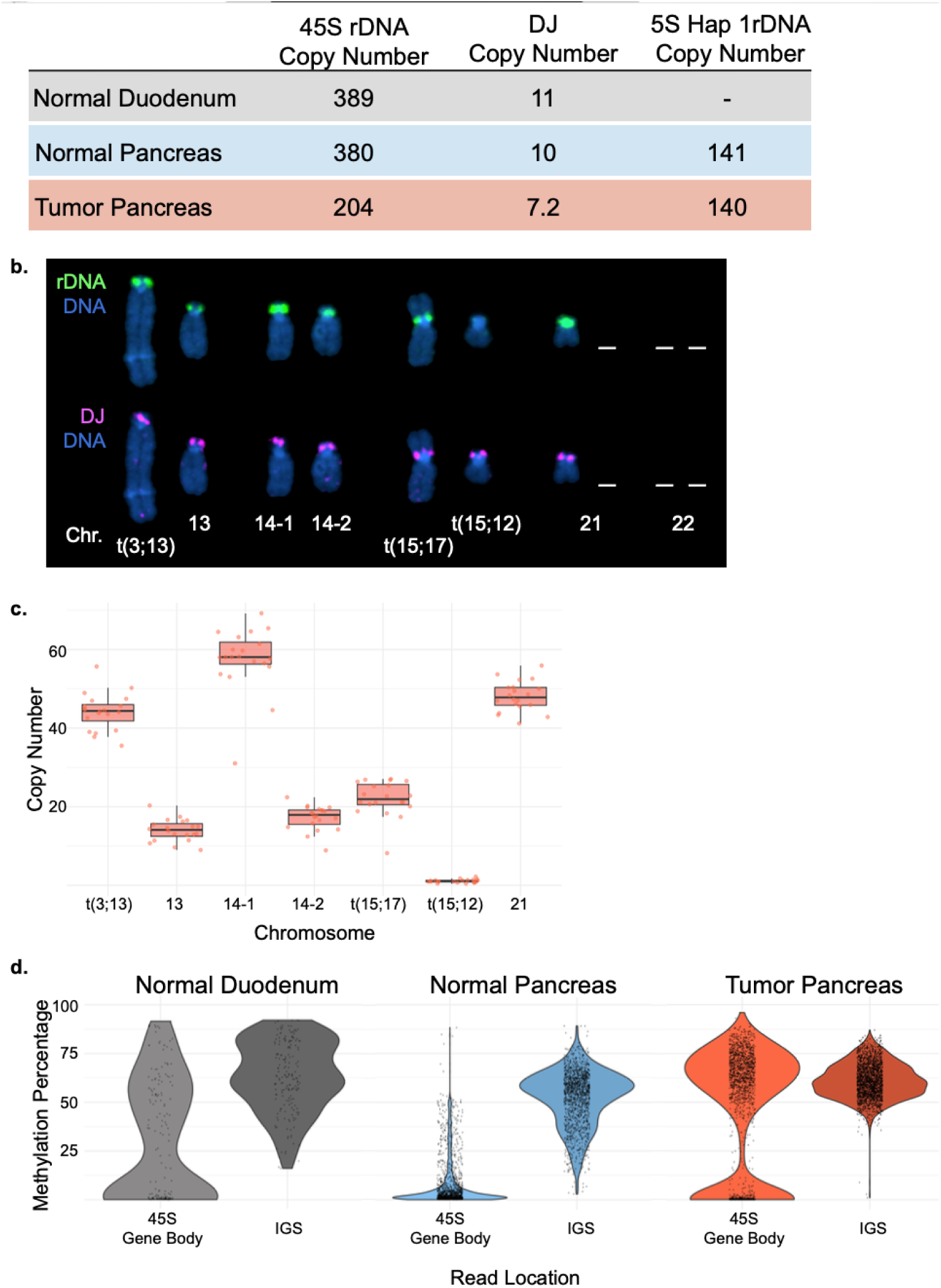
Ribosomal DNA Regions are Impacted in a Pancreatic Tumor Context. a. Copy number estimates of 45S (hypodiploid), DJ (hypodiploid), and 5S (hap 1) in the three samples are shown. Both 45S rDNA and DJ show a significant decrease in the tumor relative to either normal sample, which share a similar copy number. The 5S rDNA assembly for haplotype 1 shows loss of one gene copy in the tumor. b. Representative karyogram of 45S rDNA-containing chromosomes from hypodiploid cells. Mitotic chromosome spreads from these cells were labeled by FISH with 45S rDNA probe (green) and DJ probe (magenta). DNA was counterstained with DAPI (blue). White dashes indicate lost rDNA-bearing chromosomes. c. Distribution of copy number of 45S rRNA gene units per rDNA array, determined by fluorescent in-situ hybridization, for each of seven arrays in the pancreatic tumor line HG008-T. Each point is a single measurement (n = 20 per array); arrays are labeled by chromosome of origin, with translocation-derived arrays denoted t(a;b) and the two chromosome 14 arrays distinguished as 14-1 and 14-2. Boxes span the interquartile range and whiskers are 1.5 x interquartile range with the median indicated by the horizontal line. d. Violin plots depict the distributions of DNA methylation levels across the 45S rRNA gene body, and IGS, derived from ONT reads. Points represent per-read average methylation values within each region. Fraction of tumor reads with methylation >50% is significantly higher than either normal tissue (χ² Bonferroni-corrected p=2e-74 for normal pancreas and p=2e-11 for normal duodenum).

The 5S rDNA array on haplotype 1 was fully resolved in both tumor and normal samples and showed minimal copy number change (141 in normal and 140 in tumor), although haplotype 2 had a gap in the normal assembly. FISH demonstrates that both 5S arrays are present in the hypodiploid and genome doubled cancer cells, with the larger haplotype 1 on a normal copy of chromosome 1, and haplotype 2 on t(1:2) (Supplemental figure 46). The haplotype 1 tumor 5S gene bodies were identical to normal except for the lost monomer unit (2,232 bp) within the methylated region, and both tumor and normal had a similar unmethylated 40 kb region at the beginning of the array (Supplementary Fig. 46).

We next examined 45S DNA methylation in ONT reads from the HG008 normal pancreatic and duodenal tissue samples and tumor cell line. Previous work in cultured cell lines shows the 45S rRNA gene body is methylated when silent and unmethylated when transcribed, while the IGS remains consistently methylated.^40,41^ Here, per-read average methylation across the 45S gene body showed the tumor cell line possessing a larger fraction of highly methylated reads compared to normal tissues, suggesting unexpectedly that HG008 tumor cells transcribe fewer gene copies (Fig 6D).

Transcription of 45S rRNA genes requires Upstream Binding Factor (UBF) for RNA polymerase I loading.^42^ Cytogenetic work investigating UBF binding along the 45S rDNA arrays in the tumor sample showed while most arrays were UBF-positive, some cells exhibiting genome-doubling possessed a UBF-negative chromosome 21, indicative of transcriptional silencing (Supplementary Fig. 47), and consistent with a shift in methylation. However, we acknowledge silencing of one array is unlikely to account for the degree of shift observed and suggest in addition that the number of methylated genes within active arrays may increase.

Analysis of 45S rDNAs in the tumor assembly graph confirmed the cytogenetically-observed chromosomal fusions and matched the estimated 1.1-1.2 copies of the 45S rDNA repeat on t(15;12) (Supplementary Fig. 48). Across the remaining arrays, assembled and graph-predicted rDNA sequence variants (morphs) generally clustered by chromosome, consistent with localized repeat propagation on the DJ or proximal side (Supplementary Fig. 49; Supplementary Table 11).^43,44^. Shared morph clusters between chr14-2 and t(15;17) additionally suggested interchromosomal sequence exchange that we observed in the normal and tumor assemblies on these short arms (Supplementary Fig. 9).

In the tumor sample, loss of three 45S-rDNA arrays and changes in gene body methylation point to instability at the 45S rDNA arrays relative to the 5S rDNA arrays in HG008. These observations illustrate how near-complete assemblies enable interrogation of rDNA sequence, copy number, and epigenetic state, and reveal that the challenging acrocentric p-arm regions can harbor important structural and epigenetic changes in cancer. Although the consequences of these changes have not been evaluated in HG008, they may translate into cellular effects such as altering the pool of rRNAs available for ribosome biogenesis.^45,46^

### Curated somatic variant benchmarks

In addition to the more comprehensive SV benchmark on the normal DSA that includes centromeric SVs, we created a somatic SV and CNV benchmark on GRCh38 with deletions, duplications, insertions, VNTR expansions and contractions, and complex translocations and inversions (Supplementary Table S12). Important for use of this benchmark, v5 of the SV comparison tool truvari was developed to account for differences in how tools represent SVs as BNDs vs INV, DEL, and DUP. While SVs detectable on GRCh38 were generally detectable by both short and long reads outside TRs, one of the chr18/chrX balanced translocation pairs was missed by short reads in low mappability regions because the chr18 breakpoint was inside an L1HS and the chrX breakpoint was in an Alu between LTRs. In addition, short reads supported all L1 insertions except an L1 insertion inside a reference L1, but did not resolve the sequences of the LINE insertions. Common sources of false positives across all callsets included BNDs and DELs in tandem repeats and segmental duplications, often related to germline SVs. With availability of more comprehensive benchmarks like this, callers can improve in filtering false positives and calling SVs in repetitive regions.

Based on the benchmark small variants on the normal assembly (Supplementary Table S13), we developed a benchmark for truncal SNVs and INDELs on GRCh37, GRCh38, and T2T-CHM13v2.0, enabling reliable identification of false positives and false negatives in challenging regions, including homopolymers, tandem repeats, and segmental duplications (summaries and curation examples of challenging variants in Supplementary Figs 27-39, 50). Benchmark SNVs unique to the DSA vs. GRCh38 and T2T-CHM13 were concentrated in centromeres, while indels unique to GRCh38 vs. T2T-CHM13 were distributed across the chromosomes depending on the presence of germline variation relative to each reference (Supplementary Fig 51). FP SNVs were highly enriched in segmental duplications and tandem repeats, while FP indels were most enriched in homopolymers and tandem repeats (Supplementary Table S14 and Supplementary Fig 36). While FN SNVs were common in all three repeats, FN indels were most enriched in homopolymers and tandem repeats. Previous multi-technology ensemble benchmark approaches missed 8–40% of SNVs and 64–94% of indels in homopolymers, TRs, and difficult to map regions (Supplementary Fig. 52). A new DSA-based caller PRCGAP^47^ only missed 3% of SNVs, but still missed 80% of indels with default parameters due to noisy long reads in homopolymers.

We further compared the benchmarks on the normal assembly and GRCh38 against published mapping-based approaches that take the consensus of multiple technologies and callers. First, we compared SVs that could be represented on GRCh38 to a recent ensemble benchmark approach.^13^ The ensemble’s combination of short and long read technologies was effective at controlling false positives, with two-technology support retaining most TPs while substantially reducing FPs. However, increasing the stringency of the ensemble progressively reduced sensitivity, with curation showing these were true repetitive, complex, and subclonal SVs, with the previous publication’s requirement of 2 technologies and 4 callers missing 23 SVs modifying 32 kbp (Supplementary Tables S15 and S16), in addition to 126 SVs modifying 907 kbp that are in the DSA benchmark and not GRCh38. The >1/3 of SVs in CenSat and TR regions are challenging even for long reads mapped to the DSA, with 64 of 71 deletions and 43 of 46 insertions in these repeats missed or filtered by a new multi-technology DSA-based caller PRCGAP^47^ with baseline parameters, compared with 5 of 35 deletions and 3 of 15 insertions missed outside these repeats (Supplementary Fig 53 and Supplementary Table S17).

## Discussion

We demonstrate that near-complete assembly of all chromosomes except most rDNA arrays in an early-passage hypodiploid tumor cell line is possible. This assembly uncovers distinct centromeric structures in rearranged chromosomes as well as many small and large truncal somatic variants in repeats not detected by current reference-based methods. Even though this pancreatic cell line has different mutational signatures than a recently characterized melanoma cell line with a DSA of the normal,^2^ we find similar enrichment of somatic variants in centromeric satellites (particularly CDR and HSat2) and telomeres. While short reads detect most large somatic SVs in non-repetitive regions,^4^ the assemblies reveal many SVs in repetitive regions, with even multi-technology GRCh38-based ensemble approaches missing 11% of truncal and subclonal SVs on GRCh38 plus the 37% of truncal SVs not represented on GRCh38, and mapping-based DSA approaches missing 42% of all truncal SVs. More recent work showed that some of these SVs in repeats can be detected when assembling only the normal, but this normal DSA approach still can miss some SVs like large duplications/insertions in the centromere, and phasing and disentangling complex SVs is challenging. In addition, we find that the tumor assembly can help close gaps and correct errors in the normal assembly, particularly in regions with LOH or when ultralong reads only exist for the tumor cell line. While previous work has shown higher concordance for small variants than SVs, we find that polishing assemblies with high-accuracy reads like Element or Onso enables resolution of two times more small indels in homopolymers and TRs that are challenging to detect without the paired tumor and normal assemblies, even in this tumor without microsatellite instability. While most homopolymers occur outside coding regions, accurate calling of these can be useful for lineage analysis.^48^ In addition, somatic variants can appear to be SNVs when they are actually indels, or small indels when they are actually SVs, due to germline variation with respect to a non-DSA reference. 5,111 small variants on GRCh38 have representations differing from the true somatic change, affecting mutational signatures. These variants pose representation challenges and a need for standards for representation and tools to compare these variants across callsets and individuals. Ideally, somatic variants are called against the DSA rather than a reference like GRCh38 to avoid these representation challenges and enable robust somatic calling in repetitive and highly variable regions.

Stable persistence of derivative chromosomes composed of dynamic dicentric and fused centromeres across passages, together with critically eroded telomeres and chr13q/chr15p telomeric translocations, suggests this tumor has reached a structural endpoint of the breakage-fusion-bridge cycle. The putative kinetochores of the chr6/chr7 fusion were 1 Mbp apart, similar to the chr1-chr19 translocation observed in oligodendroglioma^49^ and dicentric Robertsonian translocations.^27^ In another case not previously identified, a single putative kinetochore was derived from kinetochores from two different chromosomes resulting in a fused centromere. Three pseudodicentrics exemplify different mechanisms for inactivation of one centromere in detail: deletion due to an interchromosomal translocation, deletion due to a translocation-associated inversion, and/or methylation. The tumor and normal assemblies also enabled discovery of exceptionally large somatic insertions in centromeric satellites caused by tandem duplications that are challenging without a tumor assembly, with insertions up to 7 times larger than the 19.4 kbp largest centromeric germline de novo insertion found in assemblies of a 17 member pedigree^34^. Although the assemblies precisely resolved all translocations causing large copy number or structural changes, including complex SVs in centromeres, additional insertions and deletions may exist in the centromeres that are not fully resolved in the normal assembly. While centromeres exhibit low recombination and instability in meiosis, these findings, alongside recent analysis of a melanoma cell line,^2^ point to centromeres as regions of genomic instability in cancer, with 39% of truncal HG008 SVs in centromeric satellites. Of 16 derivative chromosomes, 4 have breakpoints in a-satellites and 3 in other acrocentric or other centromeric satellites. More experiments and cancer types will need to be analyzed to assess the functionality of the novel centromeres, but here we establish the structural and epigenetic substrate to assess this.

Similar to the recently developed ‘genome benchmarking’ approach where the HG002 normal assembly was used as a benchmark instead of benchmarking variant calls, this paired tumor-normal assembly could serve as a prototype for developing genome benchmarking methods for somatic genomes. However, this work has several notable limitations. Most 45S rDNA arrays were not assembled, but we report array loss accompanied by increased methylation in the tumor, pointing to a substantially altered 45S-derived rRNA template pool in the tumor. In contrast, the 5S array on haplotype 1 was relatively stable in terms of copy number and DNA methylation. Here, we focus on truncal mutations to establish a baseline for future work on subclonal analyses, including analyzing cell lines derived from HG008 single cell clones. In contrast to benchmarks for which subclonal variants are typically the most challenging, this comprehensive, curated benchmark resource for cancer genomics contains challenging truncal variants that constitute more than one third of small indels, large insertions and deletions, and translocations. While the paired normal tissues for HG008 enabled us to avoid normal cell line artifacts previously observed and measure methylation in primary tissues, the quantity of normal tissues is limited. Because this is a single hypodiploid line with extensive LOH, our observations on centromere plasticity and somatic mutations are findings that motivate future work on more cancer genomes. We expect much of the methodology to extend to other tumors, but higher-ploidy or heterogeneous genomes with complex amplifications will involve more challenges resolving multiple somatic haplotypes and duplication-aware polishing. Additional genomes spanning diverse mutational and copy-number signatures will be needed to test generalizability. Therefore, in the future, it will be important for the community to develop new tumor cell lines with normal cell line pairs and broad consent for genomic data sharing and cell line distribution, enabling ongoing improvement of somatic calling in challenging regions.

## Methods

### Cell line availability and consent

The HG008-T tumor cell line is publicly available as CRL-3734 from ATCC. The ATCC cell line is derived from the same passage 13 stock as the MGH passage 23 cells (the primary source of data for this work) and the NIST passage 21 and 41 cells (used to confirm truncal variants). Therefore, we expect that truncal variants should generally be observed in cells from all sources, but subclonal variants will differ. While data for the assembly was derived from MGH passage 23 cells (0823p23), DiMeLo-seq, CUT&RUN, and acrocentric FISH measurements were from ATCC batch# 70079483 at around passage 28. Fiber-seq measurements were from 3 clones of HG008-T grown from single cells at NIST: HG008-SC14 P20+15 (PS01680), HG008-3E4 P15+12 (PS01682), and HG008-SC24 P15+9 (PS01684), where the first number after P indicates the passage at which single cell cloning was done and the number after + indicates the number of passages from initial growth of the clone. NIST and MGH are working towards depositing the clonal lines in a public repository. Normal cell lines and tissues are not available - only data from normal pancreatic and duodenal tissues.^5^

### Strategy for assembling the tumor and normal genomes

We hypothesized that a near-complete assembly of the average cell in this PDAC tumor cell line may be possible because most large somatic CNVs were deletions, with only a small number of large duplications, and most SVs were truncal or in a small fraction of cells. Accordingly, we adapted the process used in recent T2T normal genome efforts (Verkko-based assembly with PacBio HiFi, ONT, and Hi-C reads) to generate haplotype-resolved assemblies of the normal and average tumor genomes. We used Verkko so that we could curate the assembly graphs and create new assemblies with curated paths. This was important for the tumor to resolve duplications and for the normal because only the tumor had ultralong ONT reads due to degradation of DNA in the normal tissues.

### Initial Assembly Using Verkko

We used verkko v2.2 and v2.2.1 to assemble normal and tumor genomes, respectively, using HiFi and ONT reads with Hi-C for phasing.^17^

For the normal, data from both duodenal and pancreatic tissues were combined for ONT-std R10.4.1 25kb+ reads totaling about 16x coverage per haplotype under https://ftp.ncbi.nlm.nih.gov/ReferenceSamples/giab/data_somatic/HG008/Liss_lab/Northeastern_ONT-std_20240422/. Combined HERRO-corrected reads are under https://ftp.ncbi.nlm.nih.gov/ReferenceSamples/giab/data_somatic/HG008/Liss_lab/Northeastern_ONTstd_20240422/HERRO-corrected-20250218/. For PacBio HiFi, we used pancreatic tissue reads with 17x coverage per haplotype under https://ftp.ncbi.nlm.nih.gov/ReferenceSamples/giab/data_somatic/HG008/Liss_lab/PacBio_Revio_20240125/). For Hi-C, we used Arima HiC sequencing of pancreatic tissue with Illumina under https://ftp.ncbi.nlm.nih.gov/ReferenceSamples/giab/data_somatic/HG008/Liss_lab/Arima_HiC-ILMN_20240112/.

For the tumor, data from bulk passage 23 (batch 0823p23)^5^ was used. ONT reads were 27x coverage per haplotype with N50 84kb at https://ftp.ncbi.nlm.nih.gov/ReferenceSamples/giab/data_somatic/HG008/Liss_lab/Northeastern-ONT-UL-20241216/. HiFi Reads were all reads (58x per haplotype) from: https://ftp.ncbi.nlm.nih.gov/ReferenceSamples/giab/data_somatic/HG008/Liss_lab/PacBio_Revio_20240125/ as well as only the 20kb+ reads from https://ftp.ncbi.nlm.nih.gov/ReferenceSamples/giab/data_somatic/HG008/Liss_lab/BCM_Revio_20240313/HG008-T_ubams/. For Hi-C, we used Arima HiC sequencing with Illumina under https://ftp.ncbi.nlm.nih.gov/ReferenceSamples/giab/data_somatic/HG008/Liss_lab/Arima_HiC-ILMN_20240112/.

To correct errors in ONT reads prior to assembly, we used HERRO^50^

(https://github.com/lbcb-sci/herro) so that corrected ONT reads could be used by verkko to build the graph under the --hifi parameter, with uncorrected reads input as well under the --ont parameter. The version of HERRO used was the debug branch (commit on April 22, 2025) on the HERRO github repository. The Minimap2 parameters used for overlapping were -K8g -cx ava-ont -k25 -w17 -f 0.005 -e200 -r150,150 -z200 -m<X> where <X>=1500 for the tumor sample and <X>=100 for the normal sample. The command used for running the verkko assembler was like “verkko -d verkko_asm_out --hifi hifi.fq.gz ont_HERRO.fa.gz --hic1 HiC-R1.fastq.gz --hic2 HiC_R2.fastq.gz –nano ont_uncorrected.fq.gz --unitig-abundance 7”. Contiguity of automated assemblies of HG008 was limited by lack of ultra-long reads in the normal and by large somatic duplications and subclonal SVs in the tumor. Of the 46 normal chromosomes, verkko output 32 T2T scaffolds (4 T2T contigs), and of 35 tumor chromosomes, verkko output 21 T2T scaffolds (15 T2T contigs).

### Manual Resolution of Tumor Assembly Graph

To resolve large duplications and achieve a near-complete tumor assembly, we curated the assembly graph in https://ftp.ncbi.nlm.nih.gov/ReferenceSamples/giab/data_somatic/HG008/Liss_lab/analysis/Verkko_assemblies_05162024/HG008T/HG008T_verkko_v2.2.1_herro_corrected/PBhifi+20kbBCMhifi_UL_ONTq26_herro/. Most tandem duplications and the 23 Mbp duplication of the start of chr1 were resolved automatically by verkko with the ultralong reads. We manually resolved 3 large somatic tandem duplications larger than 1 Mbp by going around loops in the graph twice, randomly selecting which path in each bubble comes first when these variants between duplications were not automatically phased by verkko (Supplementary Fig 54). This choice means the smaller somatic variants occurring after the duplication may be on the wrong copy of the duplication, but when the tumor assembly is aligned to the normal assembly or the reference genome, somatic variants will still be called accurately. Variants were similarly assigned randomly on the very large 83 and 64 Mbp duplications of chr17 and chr20 if they were not phased by the assembler. We also needed to manually resolve paths due to a subclonal translocation connecting most of the chrX-chr18 hybrid to the chr2-chr20 hybrid (https://github.com/jzook/HG008SVcuration/issues/101). While this subclonal variant is in about half of the passage 23 batch of cells used for most of our sequencing, it is not detectable in other passages, so we chose the path without this variant for our tumor assembly, and output the subclonal chr2-chr20-chrX-chr18 hybrid chromosome in a separate fasta file that is not used further in this work. This subclonal translocation joins chr2_hap2 to chrX_hap1 so it causes a challenge in our phasing convention, since for truncal translocations we were able to arrange the haplotypes in the normal and tumor such that they match (except for chr19 where the tumor chromosome combines both haplotypes). This curation, as well as manual removal of some erroneous sequences after the telomeres, resulted in all 35 chromosomes being T2T scaffolds and 28 T2T contigs, with only 8 gaps, with all in acrocentric short arms around rDNA. The chr15_hap2 rDNA array was assembled because it was a single monomer, but other rDNA arrays contained gaps in the assembly. While not experimentally confirmed, the assembler uniquely scaffolded the telomeric distal bits for the acrocentric short arms in the tumor, except for the highly identical chr14 and chr15, which were manually resolved as described above. The manually resolved paths are listed in Table S18.

For regenerating the assembly sequences after each round of manual curation, we created a path file (manual.paths.gaf), and the command for re-running verkko was like “verkko -d verkko_asm_out_curated --assembly verkko_asm_out --paths manual.paths.gaf --hifi hifi.fq.gz ont_HERRO.fa.gz --hic1 HiC-R1.fastq.gz --hic2 HiC_R2.fastq.gz –nano UL-ont.fq.gz --unitig-abundance 7”, as suggested by the verkko developer.

### Manual Resolution of Normal Assembly Graph

The automated normal assembly contained many gaps due to the lack of ultralong reads (outputs in https://ftp.ncbi.nlm.nih.gov/ReferenceSamples/giab/data_somatic/HG008/Liss_lab/analysis/Verkko_assemblies_05162024/HG008_N_asm_hifiherrohic_verkko2.2_20250218/). To improve the normal assembly, we aligned the tumor assembly to the normal assembly graph using GraphAligner v1.0.20. We then manually chose paths in the normal matching haplotypes present in the tumor, except where somatic structural variation occurred. The paths of the tumor assembly through the normal graph resolved many gaps where the phasing had been ambiguous or the tangle had been too complex without long reads, and it also corrected phasing that verkko had assigned incorrectly without the additional information from the tumor. We manually chose the paths to avoid incorporating somatic variation from the tumor into the normal assembly.

Generally, somatic SVs were clear because they caused breaks in the alignment of the tumor to the assembly graph. We also scaffolded the normal acrocentric short arms based on the tumor, except for chr21_hap2 and chr22_hap1, which were deleted in the tumor and lacked sufficient evidence in Hi-C to scaffold.

Note that we could not resolve all regions in the normal assembly that were missing from the tumor, so more errors and gaps remain in them, but they do not affect the discovery of somatic variants. This curation, as well as manual removal of some erroneous sequences after the telomeres, resulted in 44 of 46 chromosomes being T2T scaffolds and 23 T2T contigs, with only 53 gaps, mostly in centromeres and rDNA tangles. We confirmed the structural and phasing accuracy of the assemblies by mapping Hi-C reads to the combined haplotypes, which demonstrated much higher connections within each haplotype-resolved chromosome (93.5% for normal and 94.2% for tumor) than between haplotypes or chromosomes, with no evidence of large structural or phasing errors (Supplementary Fig 3).

### Polishing

After manually resolving structural issues in the assemblies, we polished the assemblies by aligning short insert and long insert Element, HiFi, and ONT reads^5^ to the assemblies and calling variants with DeepVariant. The use of HERRO-corrected ONT and HiFi reads to form the consensus sequence may have resulted in increased homopolymer errors. HiFi and uncorrected ONT reads were mapped only to the combined haplotypes, while short and long insert Element reads were mapped to the combined haplotypes as well as to each haplotype separately because the shorter reads often cannot be mapped uniquely to one haplotype. Homozygous variants with GQ>4 and coverage less than two times the mean were candidates for correction, with additional filtering for long reads. Variants in homopolymers longer than 6 bp and dinucleotide TRs longer than 11 bp, as well as 1 bp and 2 bp indels and GQ<10, were filtered for ONT due to its higher error rates. HiFi variants were ignored in homopolymers longer than 6 bp and dinucleotide TRs longer than 11 bp if any Element callset had a 1 bp to 4 bp indel call, since Element is more accurate for small indels in these repeats if it is mappable. Finally, calls from all technologies were merged, calls within 5 bp of a somatic SV from the tumor to normal svim-asm calls were removed, and the remaining edits were applied to the assembly using bcftools consensus. The same process was applied for both tumor and normal assemblies.

We made 204,640 (552 SNV, 204,071 indel, and 17 SV) and 164,093 (694 SNV, 163,368 indel, and 31 SV) corrections to produce the v6.2 normal and v3.1 tumor assemblies, respectively, by aligning short and long-insert Element, PacBio HiFi, and ONT reads to the combined haplotypes, and Element to separate haplotypes. 99.6 % and 99.4 % of corrections were in homopolymers or diTRs, mostly from Element reads, which are more accurate in long homopolymers. While most errors in long homopolymers do not impact QV because they often do not contain unique *k*-mers and sequencing reads are noisy, the QV still increased from 54 to 59 using Illumina 31-mers or 65 to 66 using PacBio HiFi 31-mers. These QVs are near the limits of the utility of *k*-mer-based approaches, as many remaining *k*-mer errors were in regions with technology dropout. The lowest quality regions in the tumor assembly were those with highly identical somatic duplications, since short reads and most long reads cannot map uniquely to each copy. When identifying putative somatic variants by mapping the tumor to normal assemblies, the largest improvement after polishing was for 1 bp insertions, decreasing from 194,000 to 19,000, and 1 bp deletions, 2 bp insertions and deletions, and 3- to 49 bp insertions also decreased more than two fold. In addition, curation of variants called from Onso, Element, PacBio HiFi, and ONT reads aligned to their respective polished assemblies indicated most were not errors in the assembly, though we conservatively use some of these calls to exclude regions with potential errors or subclonal variation from the small variant benchmark.

### Polishing duplications for v3.4 assembly

Large, near-identical segmental duplications in the HG008-T v3.2 assembly cause reads to map with MAPQ 0 across both copies, so standard read-based polishing leaves these regions uncorrected. Supplementary Fig 55 details the process of identifying duplications and successively applying rules to edit and polish the assembly in these regions. It begins with identifying duplicated regions within each haplotype separately by minimap2 self-alignment.

Known assembly errors such as a false duplication flanked by a gap on chr18_chr20_hap1 and duplication regions with ≥3 copies, mostly rDNA, were excluded from masking. A second, independent source of duplications was obtained by lifting the manually curated tandem duplications >10 kb from the matched HG008-N v6.3 structural-variant benchmark onto the tumor assembly and retaining those that mapped as two adjacent copies on a haplotype. The final set comprised 52 duplications with 13 from self-alignment and 39 from the normal-benchmark liftover. For each duplication we designated copyA and copyB and built masked diploid references: set A, in which every copyA was replaced with N so reads from both copies map to copyB, and set B, which is the reverse.

Element AVITI short reads from standard and long-insert libraries were aligned with minibwa to each masked reference using either the full diploid reference or to each haplotype separately. PacBio HiFi reads from the HG008-T bulk sample were aligned with pbmm2 to each diploid masked reference. To limit compute, set B alignments used only reads overlapping footprint of the duplications extracted from the set A alignments padded by 20 kb. DeepVariant was run on each of these alignments with only FILTER=PASS records used. All variant calls were normalized with bcftools norm -a -m -any against the corresponding masked reference before comparison, so that homopolymer indels represented with different anchors by different callers were matched correctly.

Because masking collapses both copies onto one, a homozygous-alternate call indicates that reads from both copies disagree with the exposed copy’s sequence, whereas a heterozygous call indicates that the two copies should differ from each other at that position. For alignments to only one haplotype, homozygous-alternate calls were used as evidence, because a heterozygous call usually reflects germline heterozygosity rather than an assembly error.

Individual calls were ignored as evidence if any of the following held: the call’s DP exceeded twice the median DP of PASS records from the same VCF within the same duplication; the position fell in 1 kb windows identified by mosdepth as exceeding twice the median depth over the duplication footprint since the VCF DP is downsampled at high coverage; and DP < 10. A heterozygous Element call with zero reference-supporting reads was flagged as a multiallelic-split artifact and prevented from driving a one-copy decision. Sites that fell inside the 1 kb padding around a duplication interval but outside the interval itself were dropped as footprint-edge artifacts.

Each of the 139,756 candidate sites across both sets was classified into one of four decisions. Homozygous support was defined as a homozygous-alternate call with GQ > 10 from any Element configuration or from HiFi while heterozygous support as a heterozygous call with QUAL > 10 from a diploid Element configuration or from HiFi. Sites were first stratified by whether they fell within homopolymers longer than 6 bp or dinucleotide tandem repeats annotated on v3.2 because Element reads are more accurate than HiFi in these contexts.

Inside repeat annotations, Element decided the genotype: Element homozygous support resulted in a site classified as apply_both_copies and Element diploid heterozygous support resulted in apply_one_copy. A HiFi-only homozygous call in a repeat was accepted as apply_both_copies only if its GQ ≥ 10 and Element had made no PASS call of any allele at that position though a different-allele Element call at the same position indicates a conflicting representation rather than absence of evidence, and such sites were removed from the benchmark regions. A HiFi-only heterozygous call in a repeat gave apply_one_copy when under the same no-Element-call condition so was removed from benchmark regions.

Outside repeat annotations, homozygous support from Element or HiFi gave apply_both_copies. For heterozygous sites, SNVs required HiFi heterozygous support while indels required Element diploid heterozygous support corroborated by at least one further source such as a second Element configuration or HiFi. Otherwise, HiFi heterozygous support could corroborate a haplotype-split Element heterozygous call. Sites with a haplotype-split heterozygous call and no other support were rejected while all remaining sites were removed from the benchmark regions.

Decisions were converted to edits of the unmasked v3.2 assembly, or to entries in an exclusion list of sites we could not resolve, by the rules as detailed in Supplementary Fig 55. In total, 14,370 edits were applied and 3,642 sites were excluded. Edits were written as a VCF against the unmasked v3.2 assembly after confirming that every reference allele matched the v3.2 sequence, and applied with bcftools consensus. As independent checks that used no data involved in building v3.4, sequence differences between each duplication and its ortholog in the normal HG008-N v6.3 assembly fell from 10,237 to 2,110 across the 50 duplications whose copies derive from the same normal haplotype. 93.0% of edits lifted to normal coordinates fell within 50 bp of a site independently flagged as an error in the previous v3.2 HG008-T assembly.

### Checking Assembly Completeness

When using compleasm^51^ (https://github.com/huangnengCSU/compleasm, v0.2.7) with the primate gene database containing 11,834 genes^34,52,53^, the combined normal haplotypes have no missing or fragmented genes, 4 genes are on only one haplotype, 85 and 78 genes are duplicated and 89 and 91 genes have frameshift mutations on haplotypes 1 and 2, respectively. The genes missing from one normal haplotype were *STX1* (which appears correct in both haplotypes), C2orf80 (which has a true 25 kbp deletion on hap2), and *PPP1R12A* (which has a gap in hap1). *PLIN4* is fragmented because it has a true 396 bp deletion in an exonic TR. The highly polished HG002v1.1 assembly has similar numbers of genes with duplications (76 and 83) and frameshifts (90 on each). The combined tumor haplotypes are missing one gene entirely due to deletions of *CDKN2A* on both haplotypes and have one fragmented gene due to a translocation in *KDM2B* and deletion of the other haplotype, with about half of genes on only one haplotype due to large CNVs, and 136 genes have frameshift mutations. NucFlag identified potential errors with PacBio HiFi reads, which were fixed with a hifiasm assembly in the final v6.3 normal and v3.2 tumor assemblies when the hifiasm assembly reduced errors. We follow the same methods as in “Assembly repair within centromeric regions” in Ref ^34^. Nucflag version is 0.3.3 and MISJOIN, COLLAPSE, COLLAPSE_VAR, COLLAPSE_OTHER are considered errors (only HET were not considered errors). A detailed explanation can be found at the bottom of https://github.com/logsdon-lab/NucFlag/wiki#nucflag-v10-1. For normal, we repaired once with an ONT-only hifiasm release r799 with default parameters. For tumor, we repaired twice with two ONT-only hifiasm assemblies from releases r726 and r839 with default parameters. In the final assemblies, NucFlag statistics are in Supplementary Tables S19 and S20, and HMM-Flagger statistics are in Supplementary Table S21, with most errors identified around assembly gaps.

Finally, we used KaryoScope to assign each contig to its chromosome(s) of origin and to order and orient them based on CHM13 (Fig. 1a and Supplementary Fig 56). For tumor chromosomes derived from more than one chromosome, we include all chromosome numbers in the name, and orient by the largest sequence.

### Handling unassigned contigs

We removed the remaining unassigned contigs from the tumor assembly because these were redundant sequences or rDNA, but we kept 18 unassigned contigs with nonredundant sequence for the normal assembly, with 16 mostly centromeric contigs ranging in size from 111 kbp to 2.2 Mbp, and 2 contigs for the acrocentric short arms of chr21_hap2 and chr22_hap1. To assign potential chromosomes to the centromeric contigs, we mapped the unassigned contigs from HG008N_v6.3 to all 466 HPRCv2 haplotype assemblies using minimap2 (v2.30, asm5 preset).

For each contig, haplotype, and chromosome, we merged overlapping query-coordinate intervals to compute query coverage fraction, and separately computed a mean gap-compressed identity weighted by alignment length. For each contig and chromosome, we then computed the product of these two values per haplotype (0 if the contig had no alignments to that chromosome in a given haplotype) and ranked chromosomes by the mean of this score across all 466 haplotypes. Contigs were considered assigned to the top-ranked chromosome when at least 95% of haplotypes agreed on that chromosome (haplotype unanimity ≥ 0.95) and mean score ≥ 0.60. 13 contigs were assigned (Supplementary Table S22).

### Matching tumor haplotype assembly to normal haplotype assembly

The tumor contigs and scaffolds exactly match the chromosomes found in most cells with dGH SCREEN and traditional karyotyping (**Fig. 1a**). All large structural events found by dGH SCREEN in >80 % of cells are recapitulated in the tumor assembly, and other events were in only 1 or 2 of 80 cells. We also manually separated the assemblies into haplotype 1 and haplotype 2 to ensure the haplotype chromosome numbers of the tumor and normal are the same. The matched tumor and normal haplotypes were confirmed by haplotype-specific markers (Supplementary Fig 2c). This matching of haplotypes was possible for all truncal events in HG008 except the true haplotype switch on chr19. Interestingly, this matching would not be possible for all tumors and would not be possible for a subclonal chr2_hap2-chrX_hap1 translocation that occurs between different haplotypes in HG008 (https://github.com/jzook/HG008SVcuration/issues/101). The tumor assembled haplotypes were aligned to their corresponding normal haplotypes, so that differences are putative somatic variants (**Fig. 1**b). Potential haplotype switch errors or somatic changes were detected by looking for *k*-mers from each normal assembly haplotype in the tumor contigs, and the only evidence for a haplotype switch was in the chr19_chr22 hybrid chromosome that contains both chr19 haplotypes (Fig. 2), as described below. The alignments between tumor and normal haplotypes form the basis for comprehensive truncal somatic variant calls.

### Comparing normal haplotype assemblies to T2T-CHM13v2.0

The normal haplotype assemblies were compared to T2T-CHM13v2.0 using SyRI to illustrate the high level structural difference (Supplementary Fig. 2a). A known large polymorphic inversions on the p-arm of chromosome 8 for both HG008_hap1 and HG008N_hap2 were observed, which is similar to that in HG002^7^. Similar inversions were found in haplotype assemblies of chr9, chr15, and chr16. When matched tumor haplotype assemblies were aligned onto their respective normal assembly, we noticed that large inversions exist on tumor chr3_hap1, chr18_hap1, chr5_hap2, and chr7_hap2 (Supplementary Fig. 2b).

### Subclonal whole genome duplication

Karyotyping and dGH show that some cells have experienced a whole-genome duplication event (i.e., at some evolutionary epoch all chromosomes then present were duplicated), and almost all cells have some unique large copy number variation. While the fraction of whole genome doubled cells is low at the passages sequenced here (9 of 40 cells at passage 21), it increases with passages, making up 28 of 40 cells at passage 53, and 20 of 20 cells at passage 101, as seen previously.^15,54^ Because whole genome doubling and low frequency CNVs occurred more recently and vary between passages, we do not include these in our truncal benchmark, and normalize copy number to the hypodiploid state.

### Computational Karyotyping with KaryoScope

KaryoScope is a k-mer-based computational karyotyping framework that classifies assembled genome sequences in an alignment free manner to determine chromosomal composition and annotate structural features.^18^ We applied KaryoScope to both the normal and tumor assemblies to assign each contig to its chromosome(s) of origin and to order and orient them. KaryoScope uses a reference k-mer database derived from CHM13 to classify query sequences by chromosomal origin and annotate cytogenetic features such as centromeres, pericentromeric satellites, and subtelomeric repeats. Contigs containing interchromosomal rearrangements were assigned to all constituent chromosomes. To compare computational and cytogenetic results, we generated Fig. 1a displaying KaryoScope haplotypes as colored bars representing chromosomal composition alongside KromaTid dGH microscopy images for each chromosome, with ISCN annotations for both methods.

### Hybridization-based Karyotyping with KROMASURE SCREEN

Whole-genome cytogenetic structural variant analysis was performed using directional genomic hybridization (dGH) SCREEN to characterize chromosome-scale structure in HG008-T across early and late passage conditions, enabling assessment of both shared clonal architecture and passage-associated structural variation.

SCREEN is a single-cell, strand-specific cytogenetic method that enables genome-wide detection of numerical and structural chromosomal variation, including inversions, translocations, insertions, copy number alterations, and complex rearrangements. The method selectively removes one DNA strand prior to probe hybridization, enabling differential labeling of sister chromatids and direct visualization of strand orientation. This strand-specific resolution enables detection of intrachromosomal rearrangements, such as inversions, that are not readily resolved by conventional cytogenetic or sequencing-based approaches (Ray *et al.*, 2013; Robinson *et al.*, 2019, Cross and Bailey, 2026).^16,55,56^

Cells were cultured and harvested using standard cytogenetic procedures with incorporation of halogenated nucleotide analogs for one cell cycle, followed by mitotic arrest and metaphase preparation. Metaphase spreads were quality-filtered based on chromosomal morphology, hybridization performance, and resolution prior to analysis.

All human chromosomes were simultaneously assessed using a five-color high-density paint configuration. Individual metaphase cells were imaged, assembled into karyograms, and analyzed at single-cell resolution. Structural events were scored per cell and classified as whole-chromosome gains or losses, inversions, interchromosomal rearrangements, large deletions including truncations and homolog size asymmetry, sister chromatid exchanges, and complex multi-breakpoint events. Structural variation was detected at an effective resolution of approximately 20-30 kb. Interpretation of structural events incorporated prior cytogenetic characterization, including features established from G-banding datasets, together with an inverted DAPI channel to support chromosome identification, banding context, and breakpoint localization, particularly in regions of high structural complexity performed in concert with the SCREEN analysis.

To resolve complex and multi-chromosomal rearrangements, targeted follow-up hybridizations were performed using modified paint configurations in which subsets of chromosomes originally labeled with the same fluorophore were separated into distinct color channels. These follow-up assays were performed on the early passage sample and enabled unambiguous identification of chromosomal partners and source material within complex rearrangements, including marker chromosomes and inserted segments that could not be fully resolved using the standard five-color configuration. Information derived from these targeted analyses was incorporated into final structural annotations across both early and late passage datasets. Two modified screen assays were run with the following configurations: Modified assay #1 involved Chr.1 (yellow),Chr.3 (orange),Chr.7 (red), and Chr.19 (green). Modified assay #2 involved Chr.6 (yellow), Chr.11 (red), Chr.12 (green), Chr.18 (aqua), Chr.20 (orange). Events resolved by modified dGH SCREEN assay analysis are incorporated into the final annotations and include the removal of the “?” from the following preliminary calls: der(3)t(3q-11q-6q), der(5)t(1q-5q), der(6)t(7p-6p;11p-6q)inv(7(p), der(11)t(3q-11q), t(12;15)(15q-12q), t(Xq-18q), and der(22)t(19-22p).

Per-cell event calls were aggregated to generate population-level structural profiles, enabling quantification of clonal and subclonal variation across the cell population. The single-cell nature of the assay enables direct assessment of structural heterogeneity and identification of recurrent chromosomal configurations consistent with shared clonal ancestry.

SCREEN analysis was used to define chromosome-scale architecture and to validate structural features observed in haplotype-resolved tumor genome assemblies. Structural rearrangements identified by SCREEN were compared with sequencing-based, assembly-derived karyotypes to assess concordance in chromosome composition, translocation partners, and large-scale structural organization. Because SCREEN directly interrogates intact metaphase chromosomes, it provides an orthogonal measurement to sequencing of genome structure that is independent of sequence alignment and enables detection of rearrangements in repetitive and centromeric regions that remain challenging for sequence-based approaches.^57,58^

SCREEN analysis of the HG008-T cell line revealed a hypodiploid genome characterized by widespread chromosome arm loss, consistent with prior karyotyping and sequencing-based observations. Across analyzed metaphase cells, the majority of structural features were shared across cells, indicating a dominant clonal architecture, while additional variation was observed at lower frequency, consistent with subclonal diversification and passage-associated evolution.

Chromosome-scale losses were prevalent across the genome, frequently involving partial or complete loss of chromosomal arms. In a subset of cells, these losses co-occurred with genome doubling, resulting in mixed ploidy states across the population. Consistent with the underlying dataset, the proportion of genome-doubled cells increased substantially between early and late passage samples, reflecting expansion of this population during in vitro propagation. In addition to numerical abnormalities, extensive structural variation was observed across multiple event classes, including interchromosomal rearrangements, inversions, and complex multi-breakpoint configurations. Recurrent rearrangements involved multiple chromosomes, including chromosomes 1, 3, 6, 11, 12, 15, 18, 19, and 22, with frequent formation of derivative and dynamic dicentric chromosomes. These events were consistent with a highly rearranged but structured tumor karyotype, rather than random genome fragmentation.

Comparison of early and late passage samples demonstrated that the majority of large-scale structural features were conserved across passages, consistent with a shared clonal origin. Superimposed on this clonal backbone, additional lower-frequency structural and numerical variants were observed, with many events present in only one passage condition, consistent with ongoing genomic instability and selection during culture. Multiple complex chromosomal configurations were identified, including rearrangements involving three or more chromosomes and the presence of marker chromosomes with ambiguous origin based on conventional karyotyping. To resolve these events, targeted follow-up hybridizations using modified paint configurations were applied to deconvolute chromosomes initially labeled within the same fluorophore group. This approach enabled direct identification of chromosomal partners and assignment of source material within complex rearrangements, resolving previously ambiguous structural calls and refining the composition of derivative chromosomes.

All dGH results are available at https://ftp-trace.ncbi.nlm.nih.gov/ReferenceSamples/giab/data_somatic/HG008/NIST/HG008-T_bulk/20240508p21/KromaTiD-dGH_20241029/.

### Centromere mapping and annotation

Centromeres are specialized chromatin domains that mediate the segregation of chromosomes to daughter cells during cell division. In humans, centromeres are composed of near-identical tandem repeats known as α-satellites, which are organized in a head-to-tail fashion and assembled into arrays of higher-order repeats (HORs) that can span several million basepairs.

The epigenetic landscape of centromeres is critical to their identity and function, particularly a 75-300 kbp hypomethylated region, known as the “centromere dip region” (or CDR^59,60^), enriched with nucleosomes containing the histone H3 variant CENP-A.^34,60,61^ This combination of hypomethylated DNA and CENP-A chromatin is thought to define the site of kinetochore assembly during mitosis and meiosis, resulting in spindle microtubule attachment during metaphase and subsequent segregation of chromosomes to daughter cells during cell division.^62^

Although the genetic and epigenetic landscapes of human centromeres have been well-defined in the healthy population,^34,52,53^ they have not yet been completely assembled in any tumor cell line to our knowledge. Therefore, we sought to determine the complete genetic and epigenetic landscape of centromeres in both the normal and tumor samples and assess the events that may have led to changes within centromeres. We first identified each centromere within the normal and tumor genome assemblies using CenMAP,^34^ successfully identifying all 46 centromeres in the normal sample and all 35 centromeres in the tumor sample. The reduction in chromosome number in the tumor sample suggested chromosomal fusion events at or near centromeres, forming chromosomes with two putative functional centromeres (termed a dicentric). To test this hypothesis, we applied our centromere mapping and annotation pipeline (CenMAP^34^), which we refined to be able to detect chromosomes with more than one centromeric α-satellite HOR array.

To identify and annotate centromeric regions in the HG008 normal (HG008-N) and tumor (HG008-T) genome assemblies, we applied CenMAP, a centromere mapping and annotation pipeline (https://github.com/logsdon-lab/CenMAP; v1.2.0, commit c439499ca94227685e074256a102673f49eea400),^34^ which automatically identifies α-satellite-containing regions in a whole-genome assembly, annotates the sequence and structure of the centromeric region, determines the length of the active α-satellite HOR array, defines the location of the CDR, and generates plots for visualization. CenMAP was updated specifically for this study (v1.0 and greater) to enable the detection of multiple α-satellite-containing regions from different chromosomes on the same contig, as observed in the HG008 tumor assembly.

To detect α-satellite-containing regions, CenMAP (v1.0 and greater) runs three tools: K-Mer Counting^63^ (KMC), Satellite Repeat Finder^64^ (SRF), and Tandem Repeat Finder^65^ (TRF). First, CenMAP identifies repetitive sequences in a whole-genome assembly by identifying k-mers (k = 171 bp; the average length of an α-satellite repeat) that occur more than 10 times using KMC 3 (v3.2.4; Kokot et al., Bioinformatics, 2017). Then, satellite motifs are constructed using SRF^64^ (commit e54ca8c8eccf6b1f19428b0f862f2c90575290a0), and tandem repeats are annotated in each motif with a fork of TRF (trf-mod; commit 3e891db310124f7e5f7a630a1c006650be9d1f3a; github.com/lh3/TRF-mod). Using a custom script we developed to work with both SRF and TRF outputs, srf-n-trf (v0.1.1; https://github.com/koisland/srf-n-trf), we remove any SRF motif without a TRF motif and with a length of n x 170 bp, where n was <= 5 for α-satellite monomers or <= 42 for HSat1A monomers^66^ within 2% difference in length using the following command: srf-n-trf motifs -f ${srf_motifs} -m ${trf_monomers} -s 170 340 42 -d 0.02. With a candidate set of motifs, CenMAP maps each assembly to the motifs using minimap2^67^ (v2.29) with the following command: minimap2 -c --eqx -N 1000000 -f 1000 -r 100,100 <(srfutils.js enlong ${srf_motifs}) ${assembly.fasta}. The output PAF is then filtered to find monomers from aligned regions or CIGAR elements that contain an α-satellite or HSat1A monomer with the command: srf-n-trf monomers -p ${paf} -m ${trf_monomers} -s 170 340 42 -d 0.02 --max-seq-div 0.3. All α-satellite and HSat1A regions are iteratively merged with srf-n-trf regions if they are within 100 kbp of each other and contain at least one monomer using the following command: srf-n-trf regions -b ${monomers.bed} -m 30000 -d 100000 -s 170 340 --diff 0.02. This yields the coordinates of the centromeric α-satellite HOR array(s) greater than 30 kb. The coordinates of the α-satellite HOR array(s) are subsequently extended on either side by 1 Mb using BEDtools slop, producing the coordinates of the centromeric regions within each contig.

Following the detection of candidate centromeric regions, CenMAP then assigns chromosome names to each contig. CenMAP aligns each whole-genome assembly to the centromeric regions of the CHM13 reference genome (v2.0) using minimap2 (v2.28) with the following parameters: -x asm20 --secondary=no -s 25000 -K 15G. The resulting BAM files are then converted into BED files using rustybam (v0.1.34).^68^ Any contigs containing centromeric regions are reoriented from p to q arm relative to the CHM13 reference genome with the HG002 Y chromosome assembly (v2.0). Finally, contigs are renamed to indicate the chromosome they most accurately align to based on the percent of sequence aligned to the CHM13 reference genome (v2.0), with a minimum of 5% required.

To annotate centromere sequence, structure, and epigenetic features, CenMAP uses ModDotPlot^69^ (Implemented in CenStats v0.1.3), Repeatmasker^70^ (v4.1.2p1), HumAS-SD (https://github.com/logsdon-lab/Snakemake-HumAS-SD/tree/feature/multi-chr-lib; commit: 52455f930ddc1615b2751d7a0ca546232f9e7aa3), and CDR-Finder^26^ (commit: 18a8dc62523a2de75bdbce136561ce1a9ce6fed9; https://github.com/logsdon-lab/CDR-Finder). We modified the original HumAS-SD implementation to construct monomer libraries with multiple chromosomes in cases of chromosomal rearrangements within the centromere. In such situations, a mixed monomer library is built from the base monomer library to avoid incorrect/missing annotations.

### Centromere breakpoint analysis

To detect and validate centromere fusion or recombination breakpoints, both assembly- and read-based evidence were evaluated. First, we aligned the HG008 tumor assembly to the normal assembly using a custom mapping pipeline (https://github.com/koisland/asm-to-reference-alignment; commit d3610a8f8c4a3fddedb7b64078ffb65fe0853573), default minimap2 parameters (-x asm20 --secondary=no -s 25000 -K 8G -a --eqx --cs), and rustybam trim and break PAF parameters (rb trim-paf {input.paf} | rb break-paf --max-size 10000). The resulting PAF and trimmed PAF were used with SVbyEye^71^ (commit ea6d0ddef601fdb029d88cfeec66537809ac96b4) and SafFire^68^ (commit 63596fa6426b4562a9625077fde2300e567eeece), respectively, for visualization of centromeric breakpoints and homologous regions. Next, read alignments from publicly available PacBio HiFi and ONT datasets were used to determine exact breakpoints and manually inspected through IGV^72^ (v2.19.1). IGV sessions and screenshots can be found at https://github.com/koisland/HG008_TN_analysis/tree/main/data/igv.

Visualization of both tumor and normal centromeric regions and their breakpoints was performed with a modified fork of ModDotPlot^69^ v0.9.9 that allows for more colors and fixes custom breakpoints (https://github.com/koisland/ModDotPlot; commit: 3c174a4dbdab86a46f2ad9793238ca4e1705e838). Non-centromeric regions associated with centromeric translocation breakpoints were subset with SAMtools^73^ faidx (v1.23) and/or centromeric regions were taken directly from CenMAP’s output and concatenated. ModDotPlot was run with the following command: moddotplot static --compare-only -f ${multifasta} -o ${outdir} -id 70.0 -w 5000 --breakpoints ${breakpoints} --colors ${colors}. Combining these plots with those generated by CenMAP in Adobe Illustrator produced the final figures.

### NucFlag

To detect potential assembly errors in HG008-T and HG008-N with NucFlag, we first aligned both PacBio HiFi or ONT reads to their respective whole-genome assembly using minimap2 (v2.30; Li et al., Bioinformatics, 2018) with the following parameters: -a -x lr:hqae -y --eqx --cs -I 8g. Alignments were filtered using SAMtools^73^ v1.22 and the filter flag (-F) 2308 to remove secondary, supplementary, and low-quality alignments. Next, we ran NucFlag (v1.0.0-a3) with the following command: nucflag call -i ${read_alignments.bam} -f ${assembly.fasta} -p ${num_of_processes} -t ${num_of_threads} -x {ont_r9 or hifi} -o ${output.bed}. NucFlag uses peak detection in the coverage, mismatch, insertion, deletion, and softclip signals to detect collapsed and misjoined sequences as well as insertions, deletions, softclipping, low-quality regions, mismatches, and heterozygous sites within each genome assembly. In addition, NucFlag v1.0 annotates specific repeat-associated errors associated with long-read technologies, such as homopolymers and dinucleotide repeats, to allow downstream filtering.

### Long-read CUT&RUN

CUT&RUN was performed following the CUTANA ChIC/CUT&RUN Kit (14-1048, EpiCypher) protocol with some modifications as described in Ref. ^34^. For each reaction, 5.5 x 10^5^ cells were collected, resuspended in 100 μl of cold Nuclei Extraction Buffer [0.53 mM Spermidine (21-1005, EpiCypher), and 1x cOmplete, EDTAfree Protease Inhibitor Cocktail (11873580001, Roche) in Pre-Nuclei Extraction Buffer (21-1026, EpiCypher)], and incubated on ice for 10 mins. Nuclei were immobilized with activated Concanavalin A (ConA) beads and incubated overnight at 4°C in 50 μl of Antibody buffer [1x cOmplete, EDTAfree Protease Inhibitor Cocktail, 0.5 mM Spermidine, 0.05% w/v Digitonin, and 2 mM EDTA in Pre-wash buffer (21-1002, EpiCypher)] with either mouse anti-CENP-A antibody (ADI-KAM-CC006, Enzo) or rabbit anti-IgG antibody (ab171870, Abcam) at a 1:50 dilution. 2 μl of K-MetSat Panel were added to the IgG reaction. Samples were rotated overnight on a nutator at 15 rpm at 4 °C and then washed twice with Cell Permeabilization Buffer [1x cOmplete, EDTAfree Protease Inhibitor Cocktail, 0.5 mM Spermidine, and 0.05% w/v Digitonin in Pre-wash buffer]. pAG-MNase (15-1016, EpiCypher) was added at a 1:20 dilution with Cell Permeabilization Buffer and incubated for 10 mins at RT. Then, 100 mM CaCl_2_ (21-1007, EpiCypher) was added to activate chromatin-targeted digestion at 4 °C for 2 hours. To stop the digestion, Stop Buffer (21-1003, EpiCypher) was added and incubated at 37 °C for 10 min.

For DNA purification, each CUT&RUN DNA sample was brought up to 300 μl with 0.1x TE (21-1025, EpiCypher), followed by addition of 3 μl of 10% SDS (15553027, Invitogen) and 2.5 μl of 20 mg/ml Proteinase K (P8107S, NEB), then incubated at 70 °C for 10 minutes. An equal volume of phenol:chloroform:isoamyl alcohol (25:24:1 v/v; 15593031, Invitrogen) was added to the DNA sample. The mixture was transferred to a MaXtract High Density 2ml tube (129056, QIAGEN) and centrifuged at 16,000 ×*g* for 5 min at 4°C. Recovered aqueous phase was mixed with an equal volume of chloroform (C2432, Sigma-Aldrich) and transferred to another MaXtract High Density 2ml tuber, then phase-separated under the same centrifugation conditions. The aqueous phase containing the purified CUT&RUN DNA was precipitated with 2 μl of Glycogen (AM9510, AMB) and 2.5× volumes of 100% ethanol and stored at -20 °C overnight. Pelleted CUT&RUN DNA was washed with 100% ethanol and eluted in an appropriate volume of 0.1x TE.

For ONT library preparation, 5 ng of CUT&RUN-enriched DNA was amplified using CUTANA CUT&RUN Library Prep Kit (14-1001 & 14-1002, EpiCypher). After performing end repair, adapter ligation and U-excision, DNA cleanup, KOD Xtreme Hot Start DNA polymerase (71975-3, SIGMA) was used for fragment enrichment at the indexing PCR step. Each 50 μl of PCR reaction contains 2x KOD PCR buffer, 0.4 mM dNTPs, 1.5 μl of i7 primer, 1.5 μl of i5 primer, and 1 μl of KOD polymerase. PCR cycling conditions were: 2-min initial denaturation at 94°C; 18 cycles of 10-second denaturation at 98°C, 30-s primer annealing at 55 °C, and 5-min extension at 68°C; followed by a final 5-min extension at 68°C. For PCR cleanup, 1x volume of SPRIselect beads (21-1405, EpiCypher) was used, and CUT&RUN-enrichment DNA was eluted in 12 μl of 0.1x TE. The DNA library was prepared using the Oxford Nanopore Technologies (ONT) Ligation sequencing gDNA-Native Barcoding Kit 24 V14 (SQK-NBD114.24). Appropriate amounts of CENP-A and IgG CUT&RUN DNA samples underwent DNA repair and end-prep, then were ligated with individual barcodes and subsequently with the native adapter for subsequent sequencing. An appropriate amount of library DNA was mixed with Sequencing buffer (SB) and Library Beads (LIB), then loaded onto a PromethION 24 using a R10.4.1 flow cell (FLO-PRO114M, ONT) and sequenced with MinKNOW v26.01.15 (ONT). To achieve sufficient sequencing depth, the flow cell was washed using Flow cell Wash Kit XL (EXP-WSH004-XL, ONT) and was reloaded with a fresh library.

CUT&RUN long-read sequencing data was basecalled with dorado v1.1.1 (commit e72f1492) using the super accuracy (SUP) basecalling model, with simultaneous detection of 5mC, 5hmC, and 6mA signals, using the following parameters and models: sup,5mCG_5hmCG,6mA --min-qscore10; basecall_model=dna_r10.4.1_e8.2_400bps_sup@v5.0.0, modbase_models=dna_r10.4.1_e8.2_400bps_sup@v5.0.0_5mCG_5hmCG@v3, dna_r10.4.1_e8.2_400bps_sup@v5.0.0_6mA@v3. The resulting CUT&RUN ONT data were processed using Snakemake-CutNRun (https://github.com/logsdon-lab/Snakemake-CutNRun) as described in ^34^. In this pipeline, unwanted adapters were trimmed using cutadapt (v5.1).^74^ Reads were filtered to retain fragments between 100 bp to 10 kb using seqkit (v2.10.0),^75^ and aligned to the HG008-T genome assembly using BWA-MEM (v0.7.19)^76^ with the following parameters: -k 50 -c 1000000 {ref.fasta} {reads.fastq}. CENP-A aligned read depth was normalized against IgG aligned read depth using deepTools^77^ (v3.5.6) with the following command: bamCompare -b1 {bam_CENPA} -b2 {bam_IgG} --operation ratio --binSize 5000 --minMappingQuality 0. The normalized CENP-A enrichment profile in centromere regions was extracted from the bigWig file (read_to_ref.bw) and converted to a bedGraph. Finally, we used CenPlot (v0.1.4; https://github.com/logsdon-lab/CenPlot) to visualize the following tracks: CENP-A enrichment profile from CUT&RUN, and CDR location, CpG methylation, and α-satellite HOR array annotation generated from CenMAP pipeline.

### DiMeLo-seq

DiMeLo-seq was performed as described in Ref. ^34^ to validate the CENP-A chromatin profiling on HG008-T tumor cell line. For each reaction, 7 x 10^6^ cells were resuspended in 1 ml of Dig-Wash buffer containing 0.02% Digitonin and Wash buffer [20 mM HEPES-KOH, pH 7.5, 150 mM NaCl, 0.5 mM Spermidine, 0.1% w/v BSA, and 1 tablet of cOmplete, EDTAfree Protease Inhibitor Cocktail (11873580001, Roche) per 50 ml buffer] and incubated on ice for 5 mins to isolate nuclei. Nuclei were pelleted and resuspended in 200 μl of Tween-Wash buffer [0.1% Tween-20 and Wash buffer] and supplemented with a primary antibody at a 1:50 dilution. Primary antibodies used were mouse anti-CENP-A antibody (ADI-KAM-CC006, Enzo) or rabbit anti-IgG antibody (ab171870, Abcam). Samples were rotated overnight on a nutator at 15 rpm at 4°C to allow for antibody-binding. After washing twice with Tween-Wash buffer, samples were resuspended in 200 μl of secondary antibody buffer containing 200 nM of mouse nanobody-Hia5 (Nb-Hia5) gifted by Nicolas Altemose (Stanford University) within Tween-Wash buffer. Then, the samples were rotated for 2 hours on a nutator at 15 rpm at 4°C. After washing twice with Tween-Wash buffer, samples were resuspended in 100 μl of SAM activation buffer [15 mM Tris, pH 8.0, 15 mM NaCl, 60 mM KCl, 1 mM EDTA, pH 8.0, 0.5 mM EGTA, pH 8.0, 0.05 mM Spermidine, and 0.1% w/v BSA] containing 800 μM of methyl donor S-Adenosyl Methionine (SAM; B9003S, NEB). Samples were incubated for 2 hr at 37 °C to activate the Hia5 adenine methyltransferase activity. An additional 800 μM of SAM was added to replenish the activity after one hour incubation. Samples were collected and resuspended in 400 μl of cold PBS for genomic DNA (gDNA) extraction using using the Monarch HMW DNA Extraction Kit for Cells & Blood (T3050S, NEB) by following the FindingNemo protocol.^78^ The extracted gDNA was eluted in 760 μl of extraction EB (EEB; ONT).

Using the Ultra-Long DNA sequencing Kit V14 (SQK-ULK114), 8 μl of FRA (transposase fragmentation mix) was added for optimal DNA fragmentation following the FindingNemo protocol.^78^ After the FindingNemo library clean-up step, the library was eluted in 100 μl of 10 mM Tris-HCl (pH 9.0) for a single library load. Libraries were loaded onto a PromethION 24 R10.4.1 flow cell (FLO-PRO114M) and sequenced with MinKNOW (v26.01.15). To enrich for centromeric sequences during long-read sequencing, the deplete mode of adaptive sampling enabled in MinKNOW was executed to exclude sequencing reads originating from outside centromeric regions. The depletion BED file was made by extending the coordinates of the centromere annotations of the CHM13 reference genome (v2.0) by 2 Mb on either side. To obtain enough coverage, two libraries were made for each treatment, anti-CENP-A and anti-IgG. Depending on pore availability, the flow cell was washed using Flow cell Wash Kit XL (EXP-WSH004-XL, ONT) and libraries were retrieved and reloaded every 20-24 hours. DiMeLo-seq data was basecalled with the same parameters as described in our CUT&RUN experiments. To visualize the DiMeLo-seq sequencing data, we used the Snakemake-DiMeLo-seq-Viz (v0.1.0, https://github.com/logsdon-lab/Snakemake-DiMeLo-seq-Viz) pipeline as described in Ref. ^34^. Basecalled reads were aligned to the HG008-T whole-genome assembly using minimap2 (v2.28)^67^ with following parameters: -y -a --eqx --cs -x lr:hqae -I8g -s 4000 and filtered using SAMtools (v.1.22)^73^ with the filter flag argument: -F 2308. To quantify 6mA methylation across centromeres, we filtered base modification calls to only high-confidence calls with a base modification probability of ≥99% with modkit (v0.3.2, https://github.com/nanoporetech/modkit) and the following pileup command: modkit pileup {input.bam} {output.bed} --ref {input.fa} --motif AT 0 --filter-threshold 0.99. To analyze the CENP-A enrichment profile, the relevant genome was divided into 5 kbp non-overlapping windows, and the average 6mA signal was calculated for each window by subtracting the average of control IgG signal from the average of targeted CENP-A signal. The resulting TSV files were converted to a bedGraph to visualize the CENP-A enrichment profile using CenPlot (v0.1.4; https://github.com/logsdon-lab/CenPlot) along with the tracks of annotated centromere sequence, structure, and epigenetic features.

### Constructing Hi-C contact matrices

We construct Hi-C matrices to observe support for large SVs (Supplementary Fig 12) by aligning the raw arima paired end reads of both the tumor and the matched normal tissue onto the GRCh38 reference assembly using the dockerized pipeline (https://github.com/4dn-dcic/docker-4dn-hic) provided by the 4DN consortium. The pipeline generates .pairs files for both the tumor and the matched normal that we further process with Juicer^79^ (https://github.com/aidenlab/juicer) to produce .hic files. To visualize Hi-C matrices we use the Juicebox toolkit (https://aidenlab.org/juicebox/).

### Tumor/normal haplotype assembly-based somatic SV calling

For tumor-normal haplotype assembly-based somatic SV identification, we used SVIM-asm,^80^ SyRI,^81^ and PAV^82^ to generate initial SV sets. We also selected mapping-based calls from severus,^13^ savana,^83^ cutesv,^84^ nanomonsv,^85^ and sniffles2^86^ to ensure the assemblies did not miss variants. We manually inspected each of those candidate SVs using IGV and svviz2,^87^ and retained likely truncal SVs with supporting reads in the tumor but not in the normal.

We lifted the tumor vs. normal assembly-based SV breakpoint regions to the GRCh38 reference and curated each SV, as well as CNVs caused by them, to develop a confident benchmark for SVs and CNVs with precise breakpoints. To ensure the applicability of the truncal variants across passages, we required support for variants in passage 41 HiFi reads for the truncal benchmark, a passage that originated from the passage 13 stock of cells that we expect all bulk HG008-T cells to have in common in the future. While we include many subclonal SVs in the vcf, we did not focus on characterizing low frequency SVs in this work.

### Structural and copy number variant curation

We used an SV curation process documented as issues in a Github repository (https://github.com/jzook/HG008SVcuration) which involved generating issues for all of the SVs then requesting community members to curate the SVs so that all were curated by at least two individuals. Ribbon and splitthreader^88,89^ were used to identify whether a similar SV existed with a few kb, the SV is a true translocation, it causes a CNV, the SV is part of a complex set of variants within 5 Mbp, and/or the SV is inside a larger CNV or LOH region. Within IGV, we determined if the breakpoints appear exactly correct for the SV. If the variant was an insertion we used BLAT to determine if it came from somewhere else in the genome or is a tandem duplication. Finally, the curator used IGV and ribbon/splitthreader images to determine if the SV appeared to be in all cells or likely to be subclonal. The Github repository has 306 issues and the results of this curation updated criteria for inclusion in the benchmark. We follow the VCFv4.5 specification in describing translocations and complex inversions as breakend (BND) calls with an EVENTTYPE that gives more detail about the type. Variants in VNTRs are represented as close to their true somatic change as possible even if they modified germline variants. In the draft GRCh38 V0.6 SV+CNV somatic benchmark, the FILTER field has values of CENSAT for centromeric and heterochromatic SVs, LT50 for variants close to but smaller than 50 bp, and VAFlt5percent for subclonal SVs at <5% VAF in batch 0823p23. BED files are included that include or exclude subclonal variants with VAF >5%. SVVIZ_VAF_ALL INFO field gives the VAF estimated by svviz2 for combined Element, HiFi, and ONT datasets. The SVVIZBYDATASET INFO field gives the REF and ALT counts and VAF estimated by svviz2 for Element, HiFi, and ONT datasets individually for GRCh38, but these results are generally not reliable for SVs in TRs due to germline SVs. We also ran svviz2 on single cell WGS using ∼20x coverage of Ultima reads per cell on GRCh38, and re-curated any SVs with support in <80% of cells to determine if they might be subclonal. In addition, to develop an initial set of calls on the normal assembly not in GRCh38, we used svviz with HiFi and ONT reads from the tumor and normal aligned to the HG008Nv5 assembly to evaluate support for SVs called by svim-asm when aligning the HG008Tv3.1 assembly to HG008Nv5. These images are included in the GitHub issues. These initial SVs on the v5 assembly were lifted to v6.3 for the final calls. Several HG8N6.3 INFO fields give calls from tumor vs normal assembly alignments in HG008Nv6.3 coordinates. Phasing information is based on the HG008Nv6.3 and HG008Tv3.4 assemblies.

For somatic CNVs, we curated all SV breakpoints and added CNVs with breakpoints matching the SVs if the SV caused a copy number change. In addition, if a large CNV had a breakpoint in the centromere, we added an SV based on the alignment of the tumor assembly to GRCh38. Unlike somatic breakpoints in the normal assembly centromeres, typically these GRCh38 SVs and CNVs have annotations indicating large uncertainty in the breakpoints because centromeres are not well-represented in GRCh38 and centromeres are highly polymorphic between individuals. Links to curations for all SVs and CNVs are in Supplementary Table S12.

To define the GRCh38 SV benchmark regions, the starting regions were the dipcall dip.bed for HG008-N v6.3 aligned to GRCh38 (chr1-22 and chrX regions with 1:1 assembly alignment). The region around SV_119 was trimmed at a known false duplication in the v6.3 assembly (chr15:31,455,163-31,671,392). Every non-PASS record in the ALL VCF was expanded into an exclusion region. For BND records, both breakends (POS and the mate position in ALT), for DEL/DUP/INS records: POS-END. Each region was expanded to any overlapping GRCh38 tandem repeat or homopolymer (GIAB stratification, 5 bp slop), padded by ±50 bp, and merged with other exclusions within 200 kb. This removes whole complex loci. The exclusions were subtracted from the starting regions. Two regions were then extended by hand to cover PASS SVs whose breakpoints dipcall had split (chr7:404,087 to 405,257 and chr19:38,774,993 to 38,787,017).

To define the HG008Nv6.3 SV benchmark regions, the starting regions were all of chr1-22,X on both haplotypes (6,023.3 Mb). The union of the following tracks was subtracted: 1. No tumor coverage on the matching haplotype. Each alignment was first padded 20 kb past its end, so a contig that stops at a deletion breakpoint still covers the far side. Each gap was padded by 10 kb. Haplotypes with <1% coverage (loss of heterozygosity) were excluded. Curated SV footprints (±1 kb) were subtracted back out of this track, because true breakpoints are exactly where colinear alignment ends. 2. SVs in the dipcall VCF not in our curated benchmark, which may be subclonal or errors in the tumor or normal assemblies. Specifically, we excluded indels ≥50 bp on the matching tumor haplotype, unless they lie within max(1 kb, |SVLEN|) of a curated SV’s POS or END, or, for insertions, of POS+|SVLEN|. 3. Potential assembly errors in the normal assembly identified as NucFlag (HiFi and ONT) structural-error labels or HMM-Flagger v1.2 conservative ‘Err’;/‘Dup’;/‘Col’ calls from four datasets (HiFi and ONT, “D=duodenum as normal” and “P=pancreas as normal”). Error regions containing a curated SV were kept. 4. Curated subclonal SVs or 40-50bp indels that should not be in the benchmark regions (footprints ±1 kb): GRCh38-only SVs lifted to v6.3 with liftOver (‘-multiple -minMatch=0.1’; lifted spans capped at 5 Mb); SVs curated uncertain or incorrect; LT50 records; all subclonal SVs. 5. High same-haplotype divergence between tumor and normal outside of curated SV regions. A 100 bp bin was divergent if any matching-haplotype alignment over it had ≥3 SNV/indel differences per 10 kb within ±5 kb. The tumor and normal come from the same individual, so dense differences on the same haplotype point to an assembly error, a haplotype switch or a mis-placed repeat copy. Divergent bins were merged across gaps <10 kb.

### Mechanism of somatic insertions in centromeric satellites and alignment optimization

We selected all PASS, clonal insertions of at least 2 kb that overlap cenSat annotations in the HG008Nv6.3-based benchmark, including those represented as tandem duplications, for a total of 39 sites. For each site, we took a window of POS ± 5 × SVLEN on the normal haplotype and projected it onto the matching HG008-T v3.4 haplotype contig using the dipcall tumor-vs-normal alignment. We confirmed that the difference in length between the tumor and normal windows matched the curated SVLEN.

We aligned each pair of tumor and normal windows with minimap2 2.30-r1287 and took the alignment gap closest in length to SVLEN. We then refined its position and length with edlib 1.3.9 and tested whether the inserted sequence of length L is a tandem copy of the adjacent normal segment. An insertion was called a tandem duplication (TD) if its local fit to this model at the junction was at least 0.99 and the two adjacent tumor copies were at least 98% identical. Where minimap2 split the window into fragments or merged nearby indels, we instead aligned the inserted bases to the normal and placed both tumor copies of the matching normal segment directly with edlib.

For each site, we recorded the length of identical sequence at the novel junction, which also defines the range of equivalent insertion positions. We also recorded the repeat period, taken as the most common 12-mer recurrence distance in the window, and the identity between the 1 kb flanking each end of the duplicated segment. To count novel junctions, we split the inserted sequence as placed into gap-free pieces of its alignment to the normal. A TD whose length was a whole number of repeat periods (within 2%) and whose end homology was at least 95% was classed as an in-register repeat duplication. TDs with identical junction sequence longer than the 5 kb scan were classed as having long ambiguous homology. Insertions made of exact pieces of nearby normal sequence joined at more than one novel junction were classed as multi-junction duplications, and curated LINE-1 insertions were classed as mobile element insertions.

To evaluate each sequence-resolved benchmark record, we inserted its ALT at POS into the normal sequence and measured the identity to the tumor assembly with edlib. We also checked whether POS was the leftmost equivalent position. Based on these results, eight records were corrected in later curation releases. The corrections moved POS to the leftmost equivalent position, set SVLEN to the exact duplicated length, merged records that one TD had been split into, or represented long events as tandem duplications. Each corrected record was checked by confirming that 5 kb of reference flank on each side plus the insertion reproduced the tumor assembly with no edits, and that the REF base matched HG008Nv6.3. In the final benchmark, all 34 sequence-resolved TD records reconstruct the tumor assembly exactly.

### Somatic LINE insertions

Candidate MEIs from the 50–7,000 bp insertions were manually curated based on alignment to L1HS, AluYa5, or SVA_F mobile element sequences, presence of poly-A tracts, and/or target site duplications (TSDs). Putative source (driver) full-length L1 elements were identified in GRCh38 and CHM13 (HS1) reference genomes using UCSC BLAT and the UCSC Genome Browser, and by interrogating the matched normal VCF. All insertion sequences containing LINE-derived sequences were aligned to a reference L1 element for structural characterization and poly-A tail analysis (Supplementary Fig 43). Mobile element annotation tools used to supplement manual curation were TraFIC,^90^ SVAN (<u>SVAN:</u> https://github.com/REPBIO-LAB/SVAN), and MEIGA (<u>MEIGA:</u> https://gitlab.com/mobilegenomesgroup/MEIGA_LR).

To identify differences in methylation and chromatin accessibility between source L1s and other L1s with both intact ORFs, we first identified the 8 source-L1 locations (GRCh38) in HG8Nv6.3 coordinates. The chr14 and chr15 sources are germline insertions that are not in GRCh38. Query element, L1.3 (GenBank L19088, 6,059 bp), is a full-length active L1HS: ORF1 338 aa (start at 910), ORF2 1,275 aa (start at 1,990). Dfam splits L1HS into separate 5’-end and 3’-end models, so it was not used. HG008N_v6.3.fasta was cut into 20 kb windows at a 10 kb step. These are the minimap2 query and L1.3 is the target, run as minimap2 -cx asm20 --secondary=yes -N 10 -p 0.1. Colinear hits in a window are merged, then loci are deduplicated across the overlapping windows. Output in l1_hits_ge1kb.tsv (57,119 loci with ≥ 1 kb of L1.3 aligned). Full-length means ≥ 95% of L1.3 is covered: 9,724 loci across both haplotypes. For each full-length locus, the sense sequence is taken and the longest ATG-initiated ORF is found within ±400 bp of the expected ORF1 and ORF2 starts. The 1 kb flanks of each locus are lifted to GRCh38 with transanno (minMatch 0.5). Each GRCh38 source coordinate is lifted to hap1 and hap2 separately, with the GRCh38→hap chains and liftOver -multiple -minMatch=0.1. If a single base does not lift, it is lifted with ±100 bp padding and the midpoint is used. Each lifted point is assigned to the nearest ≥ 1 kb L1 locus within 10 kb on the same contig. Only full length L1s with both ORFs intact were used for downstream analyses.

### Fiber-seq analysis of full-length L1 elements

Fiber-seq^35^ was performed on isolated nuclei. For each reaction, 1 × 10⁶ cells were pelleted (250 × g, 5 min), washed in 1 mL PBS (250 × g, 5 min), and resuspended in 60 µL of Buffer A. Buffer A contained 15 mM Tris-HCl pH 8.0, 15 mM NaCl, 60 mM KCl, 1 mM EDTA pH 8.0, 0.5 mM EGTA pH 8.0 and 0.5 mM spermidine. Cells were lysed by adding 60 µL of 2× lysis buffer (Buffer A with 0.2% IGEPAL CA-630; 0.1% final), mixing by gentle flicking and incubating on ice for 10 min. Nuclei were pelleted at 350 × g for 5 min, and the supernatant was removed. Nuclei were then resuspended in 60 µL reactions containing Buffer A pre-warmed to 37 °C, 0.8 mM S-adenosylmethionine (1.5 µL of 32 mM SAM; New England Biolabs), and either 1 µL (100 U) or 0.5 µL (50 U) of FPLC-purified Hia5 (100 U/µL in 10% glycerol). Reactions were incubated for 10 min at 25 °C and stopped by adding 3 µL of 20% SDS (∼1% final). They were mixed by pipetting and transferred with wide-bore tips to 1.5 mL tubes. High-molecular-weight DNA was extracted with the MagAttract HMW DNA Kit (Qiagen). Each 63 µL reaction was brought to 180 µL with Buffer ATL, digested with 20 µL Proteinase K for 1 h at 56 °C with shaking at 500 rpm, and transferred with a wide-bore tip to a 2 mL DNA LoBind tube (Eppendorf). It was then treated with 4 µL RNase A for 2 min at room temperature. Buffer AL (150 µL) and Buffer MB (280 µL) were added in turn, each mixed by inverting 10 times. DNA was bound to 40 µL MagAttract Suspension G for 3 min at 25 °C with shaking at 500 rpm. Beads were washed twice with 700 µL Buffer MW1 and twice with 700 µL Buffer PE (1 min each, 25 °C, 900 rpm), and residual buffer was removed after brief centrifugation. Beads were then rinsed twice on the magnet with 700 µL distilled water for 1 min without resuspension. DNA was eluted in 100 µL Buffer AE (10 mM Tris-HCl pH 9.0, 0.5 mM EDTA) by gentle mixing with a wide-bore tip, incubation overnight at 4 °C and then 5 min at 56 °C with shaking at 500 rpm. The eluate was transferred with a wide-bore tip to a new 2 mL DNA LoBind tube. PacBio SMRTbell libraries were generated using the Fiber-seq treated gDNA following the manufacturer’s SMRTbell prep kit 3.0 procedure https://www.pacb.com/wp-content/uploads/Procedure-checklist-Preparing-whole-genome-and-metagenome-libraries-using-SMRTbell-prep-kit-3.0.pdf.

Full-length L1 elements annotated as described above on the HG008-T v3.2 diploid assembly were restricted to elements with intact ORF1 and ORF2 (n = 206). Elements were grouped as source (n = 5) or non-source (n = 201) L1s based on whether they matched an annotated source. For each clone, Fiber-seq reads were aligned to HG008-T v3.2 with pbmm2, and only primary alignments overlapping each element ± 1 kb were kept. Fiber-seq Inferred Regulatory Elements (FIREs) were identified using ft fire (https://github.com/fiberseq/FIRE) with default settings and reference positions of m6A and CpG 5mC calls and FIREs were then extracted with ft extract --all --ml 0 (fibertools-rs v0.6.4).^91^ Each element was oriented by its strand, with the transcription start site (TSS) defined as its 5′ end and the transcription end site (TES) as its 3′ end. Each element was divided into 200 bins: 50 bins of 20 bp covering the 1 kb upstream of the TSS, the L1 element body scaled to 100 bins, and 50 bins of 20 bp covering the 1 kb downstream of the TES. In each bin, CpG methylation was calculated as the number of 5mC calls with ML ≥ 128 divided by all CpG 5mC calls, combining calls from both strands. m6A methylation was calculated as the number of m6A calls with ML ≥ 125 divided by the number of reference A and T bases covered by the aligned molecules. 95% confidence intervals were obtained from 1,000 bootstrap resamplings of source and non-source elements, respectively. Promoter accessibility was quantified as the percentage of molecules spanning the entire TSS ± 500 bp window that carried at least one FIRE overlapping it. A FIRE was defined as an MSP with a FIRE score ≥ 229. These percentages were pooled per group, with bootstrap confidence intervals computed as above. Per-element percentages, for elements with ≥ 5 spanning molecules, were compared between source and non-source L1s within each clone using a two-sided Mann–Whitney U test.

### Truncal small variant benchmark development

To identify true truncal variants, we used repun^92^ to evaluate support for each putative assembly-based somatic variant by short and long reads from both tumor and normal mapped to the normal assembly. To be included in the truncal somatic benchmark, we required support in >50% of tumor but <10% of normal reads mapped to combined normal assembly haplotypes with coverage > 5 (ignoring HiFi and ONT for homopolymers covered by short reads, and ignoring Element and Onso for indels >10bp or low MQ). In regions with somatic duplications, we required >30% of tumor but <10% of normal reads. For short reads aligned separately to each haplotype, we required >30% of tumor reads and <10% of normal reads, or >90% of tumor reads and <70% of normal reads to support the variant.

The GRCh38 benchmark is generated by first lifting coordinates to GRCh38. Each somatic variant is given ±50 bp of slop on the assembly, assembly-gap regions are excluded, then coordinates are lifted from the HG008-N v6.3 assembly to GRCh38 using liftOver (minMatch=0.1) with chains derived from minimap2 alignments. Lifted regions are expanded to overlapping repeats. The somatic candidate VCF is split by haplotype and rtg vcfeval (https://github.com/RealTimeGenomics/rtg-tools) is run for each haplotype, comparing HG008-T calls against HG008-N calls. False positives (tumor-unique variants), false negatives (normal-unique variants), and true positives (germline variants shared between tumor and normal) are produced for each haplotype. To generate GRCh38-based somatic variants, vcfeval FP and FN results are filtered to the somatic candidate regions lifted from calls on the HG008N v6.3 assembly, producing the per-haplotype somatic variant set on GRCh38. Regions excluded from the benchmark BEDs include subclonal variants, structural-variant breakpoints (±50 bp), assembly errors (variants flagged by DeepVariant calls on the normal assembly), and complex repeats where the assembly alignment to GRCh38 is unreliable. Phasing information is based on the HG008Nv6.3 and HG008Tv3.2 assemblies.

Three versions of benchmark beds are produced: ‘all.bed’, ‘nogermlineinterference.bed’, and ‘nogermlinewithin50bp.bed’. HG008-T_somatic_smvar_benchmark_v0.3_all.bed includes all somatic variants that are confidently called, but current comparison tools may not handle different representations of somatic variants that modify germline variants. HG008-T_somatic_smvar_benchmark_v0.3_nogermlineinterference.bed is the all.bed except with regions excluded where somatic variants modify germline variants in a way that current comparison tools cannot resolve correctly. HG008-T_somatic_smvar_benchmark_v0.3_nogermlinewithin50bp.bed is the strictest set: every region in the benchmark is at least 50 bp away from any germline variant. This is not recommended as the primary benchmark bed but rather to understand the effect of germline variants on FP and FN somatic calls.

The v0.1 draft benchmark was compared to nine community-contributed somatic small variant callsets using both aardvark^93^ (https://github.com/PacificBiosciences/aardvark) and rtg vcfeval.^94^ False positives supported by two or more callers were manually curated; some FPs were genuine somatic variants at low VAF that were removed from the benchmark regions as filtered subclonal variants in v0.2, while others resulted from caller errors or representation differences. False-negative rates were stratified by whether the assembly variant has the same or a changed representation on GRCh38; this stratification motivated the multi-tier BED design.

### Mapping & Variant calling for small variant comparisons

Short read data from the different submitters was analysed with community standard computational pipelines to detect somatic variation to build and test the variant benchmarks.

#### OncoAnalyser

Genomic short reads were analyzed using OncoAnalyser v2.3.0 [http://github.com/nf-core/oncoanalyser/releases/tag/2.3.0]. The nf-core page provides detailed parameter settings, resources, and software versions. Briefly, DNA reads were mapped to the reference genome GRCh38 using BWA-MEM2 and processed downstream using the Nextflow implementation of the open-source, multi-platform Hartwig Medical Foundations tool set WiGITS [http://github.com/hartwigmedical/hmftools]. This resulted in a comprehensive (SNV, MNV, CNV, SV, viral integrations) dataset.

#### Ultima Genomics small variant calling

Ultima genomics short reads were aligned with Ultima Aligner (UA) with default parameters (DockerHub: ultimagenomics/alignment), Somatic short variants were then called using Ultima-adapted somatic DeepVariant (https://github.com/Ultimagen/healthomics-workflows/tree/main/workflows/efficient_dv) using the default WGS somatic calling model for matched samples (https://github.com/Ultimagen/healthomics-workflows/blob/main/workflows/efficient_dv/input_templates/efficient_dv_template-WGS-somatic-matched-samples-v1_3-40X-200X.json).

#### DeepSomatic ensemble was performed on tumor-normal Illumina, ONT, and PacBio HiFi reads

Calls used DeepSomatic^95^ from https://github.com/google/deepsomatic (Docker: google/deepsomatic:1.8.0). Bam files used:

ONT:

HG008-N-D_GRCh38-GIABv3_ONT-R1041-dorado_0.5.3_5mC_5hmC_13x_gt25kb.bam

HG008-N-P_GRCh38-GIABv3_ONT-R1041-dorado_0.5.3_5mC_5hmC_20x_gt25kb.bam

HG008-T_GRCh38_GIABv3_ONT-R10.4.1-doradov0.3.4-5mCG-5hmC-63x_UCSC_20230905.bam

PacBio:

HG008-N-D_PacBio-HiFi-Revio_20240313_68x_GRCh38-GIABv3.bam

HG008-N-P_PacBio-HiFi-Revio_20240125_35x_GRCh38-GIABv3.bam

HG008-T_PacBio-HiFi-Revio_20240125_116x_GRCh38-GIABv3.bam

HG008-T_PacBio-HiFi-Revio_20240313_106x_GRCh38-GIABv3.bam

Illumina:

HG008-N-P_Illumina_150x_GRCh38-GIABv3_sorted.

HG008-T_Illumina_195x_GRCh38-GIABv3.bam

Generating benchmarking set VCF files and high confidence region BED files using DeepSomatic multi-cancer models:

VCF files - A set of high confidence variants using the multi-cancer models was generated using the same method that was used for generating a high confidence set using the single-cancer models. A union set was generated with all somatic variants from the three DeepSomatic multi-cancer models for each sequencing technology. The union set was then filtered for high-confidence variants against a criteria of: 1) for SNVs, called by at least two out of the three multi-cancer models 2) for indels, called by the multi-cancer Illumina model and also called by at least one other multi-cancer model. High-quality germline variants were also called using DeepVariant’s Illumina+PacBio hybrid model and added to the final training set.

BED files - High confidence regions for the multi-cancer models were defined using the same method that was used to generate high confidence regions for the single-cancer models described above. The only difference between these high confidence regions are the “low confidence variant regions”, which were generated using variants called by the multi-cancer models instead of the single-cancer models.

Method that was used to generate high confidence regions for the single-cancer models: BED files - Steps to generate high confidence region BED files were adapted from the methods used by SEQC2 in generating their HCC1395 benchmarking set (Supplementary Figure 2). For each cell line across all three sequencing technologies, GATK (broadinstitute/gatk3:3.8-1) CallableLoci was used to identify the callable regions for both tumor and normal bams.

Parameters were set to ‘--maxDepth [8*average_coverage] --maxFractionOfReadsWithLowMAPQ 0.1 --maxLowMAPQ 1 --minMappingQuality 20 --minDepth 10 --minDepthForLowMAPQ 10’.

‘--minBaseQuality’ was set to 20 for Illumina and PacBio HiFi reads, and to 7 for ONT reads.

Additional flags were also added: ‘--filter_reads_with_N_cigar

--filter_mismatching_base_and_quals --filter_bases_not_stored

--allow_potentially_misencoded_quality_scores’.

First, the callable regions for tumor and normal reads within each technology type were intersected. Then, the intersected set of callable regions for each sequencing technology were merged together as a union set of all callable regions. A series of three filtering steps were performed to remove these regions: 1) “Low confidence variant regions” identified by subtracting the high confidence variants included in the training sets from the union set of variants, 2) structural variant (SV) regions identified by Severus (+/- 1000bp removed from start/end of each SV), 3) segmental duplication regions published by GIAB. To generate benchmark VCF for HG008 with criteria described above, we used https://github.com/jimin001/DeepSomatic_manuscript/blob/main/deepsomatic_benchmarks/generate_benchmarks_vcf.sh. To generate benchmark BED for HG008 with criteria described above, we used https://github.com/jimin001/DeepSomatic_manuscript/blob/main/deepsomatic_benchmarks/generate_benchmarks_bed.sh. The vcf was intersected with the bed to produce the final vcf for comparison.

#### NeuSomatic ensemble was performed on tumor-normal Illumina, ONT, and PacBio HiFi reads

First, for the Illumina sample (NYGC_Illumina_NovaSeq6000), we used the existing NeuSomatic Illumina model. For PacBio and ONT, we trained technology-specific models using the data from five cancer samples provided in the deepsomatic study^95^. We then applied these models to multiple HG008 tumor-normal sample pairs. For ONT, these included UCSC_ONT_20231003 and UCSC_ONT-UL_20231207 as tumor samples, paired with Northeastern_ONT-std_20240422 ND and NP normal samples. For PacBio, we used tumor and normal samples from PacBio_Revio_20240125 and BCM_Revio_20240313.

In addition, we trained a multi-technology model (IPO) that jointly uses Illumina, PacBio, and ONT data as inputs to the network. Based on our previous experience with germline variant calling in the precisionFDA challenge, integrating multiple sequencing technologies may improve calling accuracy and robustness. The IPO model was also trained using data from the five cancer samples in the deepsomatic study. We then evaluated this model on eight different combinations of HG008 tumor and normal samples across the three sequencing technologies.

Finally, we integrated the predictions from the technology-specific and IPO models to generate a consensus VCF. A variant was included in the final call set if it was detected by the IPO model in more than 75% of the evaluated tumor-normal sample combinations, or if it was detected by at least two individual technology-specific models in more than 75% of their corresponding tumor-normal sample combinations. More than 97% of the variants in the final call set were supported by the multi-technology IPO model.

### Small Variant Curation

We used IGV 2.19.7 with a session that contained links to the HG008 v0.2 draft somatic small variant benchmark, one of nine different callsets submitted from collaborators for evaluation, and a variety of short and long read alignments as well as assemblies of both tumor and normal aligned to GRCh38. Using aardvark for comparison of each submitted callset to the draft benchmark, we randomly selected at least five SNVs and INDELs from each set of false positives and false negatives. Also, 5 false negative SNVs and INDELs that were common to all callsets were curated. For a discrepant variant, we examined both the local 48 bp window along with a larger context view of around 10 kb. Along with noting genomic context around the variant, we curated with the goal of determining whether the benchmark variant was correct at a given loci and if the comparison variant was correct or not. The IGV session included 68x HG008-N-D PacBio HiFi, 106x HG008-T PacBio HiFi, 157x HG008-T Ultima, 48x HG008-N-D Pacbio Onso, 136x HG008-T Pacbio Onso, 77x HG008-N-D Element, 111x HG008-T Element, 118x HG008-N-D Illumina, 161x HG008-T Illumina, 20x HG008-N-P ONT 25Kb, 54x HG008-T ONT-UL, HG008-N Haplotype 1 assembly, HG008-N Haplotype 2 assembly, HG008-T Haplotype 1 assembly, HG008-T Haplotype 2 assembly, a BED file denoting repeats in GRCh38, and a BED file denoting Segmental Duplications. The alignments were hosted on an NCBI-hosted FTP site for GIAB data or an S3 bucket.^5^ Curation results for the FPs and FNs are in Supplementary Table S14. Based on the curations, somatic variants or potential errors that lifted to multiple GRCh38 locations were excluded from the v0.3 small variant benchmark regions.

### PRCGAP assembly-based small variant and SV calling Methods

Somatic variants were called with PRCGAP v0.1.1 <u>(Sakamoto et al. 2026)</u> from tumor and matched normal pancreas PacBio HiFi and ONT reads against the HG008-N v6.3 assembly, using DeepSomatic (v1.8.0) for small variants and nanomonsv (v0.8.0) for SVs. The PRCGAP post-processing modules, mutation_postprocess and nanomonsv_postprocess, were updated to v0.1.3 and v0.2.6 for this study. Calls were removed when their supporting reads matched, on average, less than 80% (60% for ONT SVs) of the reference haplotype-specific k-mers they covered, and the remaining calls were compared with the small-variant benchmark v0.4 and the SV benchmark v0.6.

Small variants were normalized and matched exactly to the benchmark. In centromeric and satellite arrays, reads spanning SVs can be aligned with mismatches instead of as SVs, producing false-positive SNV clusters. In an SNV-cluster triage, we assembled tumor HiFi reads within 50 kb of each region with at least three SNVs per kb using hifiasm (v0.25.0), aligned the assemblies with mm2-ivh <u>(Suzuki et al. 2026)</u>, and removed the SNVs when the assembly showed an event of at least 100 bp within 2 kb or did not reproduce them. Variants filtered by DeepSomatic as RefCall or GERMLINE were rescued when samtools v1.17 mpileup showed a tumor depth of at least 20 with a VAF above 0.5, and a normal depth of at least 10 with fewer than three alternative reads.

SVs matched the benchmark when both breakpoints were within 500 bp with matching orientation. In centromeric arrays, one breakpoint could be up to 10 kb away if the other was within 500 bp. A tandem duplication matched an insertion of similar size (size ratio of at least 0.7) within 500 bp of either end. To find the same SV placed on another copy of a tandem or HOR array, we also applied each deletion, insertion or duplication of similar size within 10 kb to the reference, and accepted the match when the two resulting sequences differed by an edit distance of at most 0.3 times the larger SV size.

To recover SVs filtered by the default pipeline, we generated three sets of rescue calls genome-wide, without reference to the benchmark, and scored them with the default calls. First, nanomonsv was re-run without its matched-normal realignment filter, which rejects somatic SVs in repeat arrays because normal reads realign to the SV allele. Calls were instead required to have at most one supporting read in the normal nanomonsv parse output and to be detected by both HiFi and ONT. Second, SVs from the triage assemblies were added. Third, single-breakend calls were converted into two-breakpoint SVs when their consensus sequence had a bwa mem (v0.7.19) alignment of at least 2 kb with MAPQ of at least 40 at the breakpoint. Likely subclonal PRCGAP-only calls (supported by fewer than 20% of tumor reads) were labeled.

### Structural variant calling and ensemble benchmarking

We used matched tumor–normal sequencing data from the HG008 sample, generated using three sequencing technologies: PacBio HiFi, Oxford Nanopore (ONT), and Illumina. All datasets were aligned to the GRCh38 reference genome. The following BAM files were used:

● PacBio HiFi

○ Tumor: HG008-T_GRCh38_GIABv3_120x_PB-HiFi-Revio_mm2.bam
○ Normal: HG008-N-P_GRCh38_GIABv3_37x_PB-HiFi-Revio_mm2.bam
● Oxford Nanopore (ONT)

○ Tumor: HG008-T_GRCh38_GIABv3_ONT-R10.4.1-doradov0.3.4-5mCG-5hmC-63x_UCSC_ 20230905.bam
○ Normal: HG008-N-P_GRCh38-GIABv3_ONT-R1041-dorado_0.5.3_5mC_5hmC_41x.bam
● Illumina

○ Tumor: HG008-T_Illumina_191x_GRCh38-GIABv3_sorted.bam
○ Normal: HG008-N-P_Illumina_150x_GRCh38-GIABv3_sorted.bam

To evaluate the performance and limitations of different sequencing technologies and SV callers, we performed somatic structural variant (SV) calling using multiple methods. PacBio HiFi and ONT data were analyzed using Nanomonsv v0.9.0, Savana v1.3.2, Sniffles2 v2.2, and Severus v1.7, while Illumina data were analyzed using SvABA v1.2.0, GRIDSS v2.13.2, Manta v1.6, and DRAGEN v4.2.4.

SV calls from individual callers and sequencing technologies were combined using Minda to generate ensemble VCFs. We tested different merging parameters and varied the minimum number of supporting callers (1–12) and sequencing technologies (1–3). Each ensemble callset was compared against the HG008 assembly-based somatic SV benchmark using GRCh38_HG008-T-V0.6_somatic-stvar_ALL.draftbenchmark.vcf and GRCh38_HG008-T-V0.6_somatic-stvar-clonal_and_subclonal.draftbenchmark.bed. For each configuration, we quantified the numbers of true-positive (TP) and false-positive (FP) SVs to assess how increasing the required number of supporting callers or technologies affected TP retention and FP reduction.

To further investigate the limitations of individual callers and ensemble-based approaches, we identified benchmark SVs that were not detected by any caller, detected by only a single caller, or missed by multiple callers. We then examined these variants based on their structural complexity and clonality to determine whether particular types of SVs were consistently missed across methods. These analyses allowed us to assess the trade-off between sensitivity and specificity under different ensemble configurations and investigate potential biases in consensus-based SV benchmarking, particularly for complex and subclonal variants.

### Telomere Length Estimation Methods

We analyzed Nanopore long reads aligned to chromosome ends (*i.e.* first and last 10 kbp) of CHM13v2.0. Those reads were analyzed using Topsicle<u>(Nguyen and Choi 2025)</u> by quantifying 4-mer repeats based on human telomere pattern (CCCTAA), with TRC cutoff of 0.7, minimum length of long read as 7 kbp and the longest possible telomere length is 30 kbp. The same reads were also processed with Telogator2 (https://github.com/zstephens/telogator2), run with default parameters except for specifying a custom reference containing HG008 subtelomeres via the -t input option.

### Analysis of Ribosomal DNA Regions

Despite their central role in cellular physiology, rDNA loci have historically been excluded from genome-wide analyses because their repetitive structure prevents accurate mapping and assembly using short-read sequencing. This limitation persisted until the CHM13 T2T assembly, which finally resolved the p-arms of the acrocentric chromosomes, yet the 45S rDNA arrays themselves have still not been fully assembled.^6^ As a result, the stability and variation in the rDNA regions have been challenging to characterize, further preventing our understanding of how these arrays change in disease. Long-read sequencing and near-complete genome assemblies as well as new analysis methods now allow for improved interrogation of these regions, enabling simultaneous analysis of rDNA sequence, copy number, and epigenetic state in matched tumor and normal genomes. We examined rDNA copy number, methylation, and the variant profile in the normal duodenal and pancreatic tissues and pancreatic tumor cell line from HG008. As all of these measurements vary between individuals to collectively form a unique rDNA fingerprint, it is critical to use matched samples for comparisons.^40^ As the 45S rDNA arrays mostly remain as gaps, we used a k-mer based approach for copy number quantification from short read data,^40^ normalizing estimates to coverage from chromosome 4.

### rDNA Copy Number Analysis

#### 45S rDNA

Copy number of the rRNA gene was estimated using a Snakemake pipeline called CONKORD (Version 8 https://github.com/borcherm/CONKORD). This pipeline identifies bins in a reference genome that are of similar G/C percentage to the provided feature and on the same chromosome. K-mers from the provided feature and matched bins are counted in short-read whole genome sequencing data, then filtered to remove any which occur at counts greater than 3 standard deviations from the mean of the set, as well as those that do not appear at all. K-mers which are not exclusive to the matched bins are removed from the analysis, and those that occur multiple times in the reference have their counts divided by their genomic multiplicity. Copy number is calculated by dividing the median count of feature k-mers by the median count of matched bin k-mers, which effectively gives the multiple of times that the target feature occurs relative to the average genomic background from a similar sequence context. The bin size used was 2 kb, and k-mers were of size 31. The 31-mers were extracted from the 18S portion of rDNA reference sequence KY962518.1 and counted in HG008 tumor and normal samples.

Chromosome 4 of human reference genome CHM13 v2.0 was used for background k-mer normalization. This chromosome was chosen as it was consistently haploid in the tumor sample, allowing for a stable expected k-mer count at half the value of the typical genomic background. The haploid status of chromosome 4 was confirmed through CONKORD, which showed mean tumor k-mer counts for chromosome 4 at ∼28, roughly half of other chromosome k-mer counts at ∼52. Normal pancreas and duodenum counts were consistent across all chromosomes. K-mer spectra further validated this, with a distinct haploid peak using chromosome 4 k-mers in the tumor sample, leading to division of the CONKORD output by 2 for just the tumor sample.

#### 5S rDNA

Haplotype resolved genome assemblies for HG008 normal and tumor samples were used to characterize the 5S rDNA array on chromosome 1, haplotype 1. CONKORD can not distinguish between 5S genes and pseudogenes in short read data, so the 5S array copy number was based on assemblies. For each haplotype assembly, the chromosome 1 sequence was extracted using samtools faidx. The 121 bp 5S gene body sequence within the 2240 bp 5S rRNA gene monomer (NCBI accession NR_023366) was used as a BLASTN query against each haplotype’s chromosome 1 to identify individual repeat copies. Array boundaries were established in two steps. First, raw BLAST hits were filtered to keep those with at least 98 percent identity and alignment across 100 percent of the query. This produced a high-confidence subset of hits, which was used to distinguish the true tandem array — a contiguous cluster of hits with spacing consistent with the known repeat unit length (∼2.2 kb) — from isolated, lower-identity matches elsewhere on the chromosome consistent with processed pseudogenes. Copy number for assembled haplotype 1 was determined by restricting the complete, unfiltered set of BLAST hits to the confirmed array boundary coordinates.

### DJ Copy Number Analysis

A sequence known as the distal junction (DJ) is present on each acrocentric chromosome,^96^ distal to the rDNA array; a normal diploid genome contains 10 copies, with variation ranging from 9 to 11 copies.^39^ The Illumina short read sequencing data for the HG008 normal duodenum, normal pancreatic, and tumor pancreatic samples was used for estimating DJ copy number with a reference free k-mer based approach in the DJCounter framework (https://github.com/marbl/DJCounter). A curated set of DJ specific 31-mers, present once per distal junction and shared across the five acrocentric chromosomes in CHM13 v2.0 reference assembly, was used to quantify DJ abundance. K-mer counts were generated from sequencing reads and normalized to the diploid coverage peak derived from genome wide k-mer frequency distributions. This approach enables robust quantification of DJ copy number independent of read alignment and is described in detail recently.^39^

### Chromosome spreads, Fluorescent In-Situ Hybridization (FISH), and immuno-FISH

For the preparation of chromosome spreads, cells were blocked in mitosis by the addition of Karyomax colcemid solution (0.1 µg/ml, Thermo Fisher) for 6-7h and collected by trypsinization. Collected cells were spun down, resuspended in hypotonic 0.4% KCl solution, incubated for 10-12 minutes and pre-fixed by addition of methanol:acetic acid (3:1) fixative solution (1% total volume). Pre-fixed cells were spun down and resuspended in methanol:acetic acid fixative. Spreads were dropped on a glass slide and incubated at 65°C overnight. Before hybridization, slides were treated with 0.1mg/ml RNAse A (Qiagen) in 2xSSC for 45 minutes at 37°C and dehydrated in a 70%, 80%, and 100% ethanol series for 2 minutes each. Slides were denatured in 70% deionized formamide/2X SSC solution pre-heated to 72°C for 1.2 minutes. Denaturation was stopped by immersing slides in 70%, 80%, and 100% ethanol series chilled to -20°C. Labeled DNA probes were denatured separately in a hybridization buffer by heating to 80°C for 7 minutes before applying to denatured slides. Labeled rDNA BAC probe RP11-450E20 was from Empire Genomics. Whole chromosome paints for acrocentric chromosomes were from Applied Spectral Imaging (FPRPR0167). The OligoMix probe for DJ loci consisting of pooled 100bp biotinylated oligos covering the entire DJ region was synthesized by LC Sciences. 5S rDNA probe was produced by cloning a synthetic DNA fragment containing an entire 5S gene unit (coding region and intergenic spacer, 2240bp) into pUC19 vector, which was biotin-labeled using the nick translation kit (Enzo Life Sciences, Cat.# ENZ-GEN111-0050). Specimens were hybridized to the probes under a HybriSlip hybridization cover (GRACE Biolabs), sealed with Cytobond (SciGene) and incubated in a humidified chamber at 37°C for 48-72 hours. After hybridization, slides were washed in 50% formamide/2X SSC 3 times for 5 minutes per wash at 45°C, then in 1x SSC solution at 45°C for 5 minutes twice, and at room temperature once. For biotin detection, slides were incubated with streptavidin conjugated to Cy5 (Thermo Fisher) for 2-3 hours in PBS containing 0.1% Triton X-100 and 5% bovine serum albumin (BSA), and then washed 3 times for 5 minutes with PBS/0.1% Triton X-100.

For rDNA and UBF immuno-FISH, slides labeled by FISH were subjected to antigen unmasking in 65°C Citrate buffer, pH 6.0, for 45 minutes before processing for immunofluorescence. Slides were blocked with 5% bovine serum albumin (BSA) in PBS/0.1% Triton X-100 for 1hour. Primary antibody (rabbit polyclonal anti-UBF, Novus Biologicals, cat.# NBP1-82545) and secondary antibody (goat anti-rabbit Alexa Fluor 647, Thermo Fisher) were diluted in 2.5% (weight/volume) BSA/PBS/0.1% Triton X-100. Specimens were incubated with primary antibody at 4°C overnight, washed 3 times for 5 minutes, incubated with secondary antibody for 2-4 hours and washed again 3 times. All washes were performed with PBS/0.1% Triton X-100. 1µg/ml DAPI was added to final washes to label chromosomes. Slides were mounted in ProLong Gold or ProLong Glass (Thermo Fisher).

Confocal Z-stack images were acquired on the Nikon TiE microscope equipped with PlanApo 100x oil immersion objective NA 1.45, Yokogawa CSU-W1 spinning disk, ORCA-Fusion BT sCMOS camera (Hamamatsu), and NIS Elements software.

### Estimating rDNA array copy numbers from FISH images

All image processing was performed using ImageJ/FIJI. Maximum intensity projections of confocal Z-planes were generated, and rDNA-containing chromosomes segmented. For chromosome identification, combination of whole chromosome paints and morphological features was sufficient to identify all rDNA-containing chromosomes. Then each rDNA signal encompassing both sister chromatids was segmented as a single region based on the threshold applied to the entire image. The region of the same chromosome outside the rDNA was used for the local background subtraction. The background-subtracted integrated intensity was measured for each array. The sum of all background-subtracted intensities of all rDNA arrays represented the total amount of rDNA per cell. The total rDNA copy number was estimated from short-read DNA sequencing data (see “rDNA Copy Number Analysis”). The fraction of the total rDNA fluorescence intensity was used as a proportion of the total rDNA copy number to determine the number of rDNA copies on specific chromosomes in each chromosome spread. 20 hypodiploid spreads were analyzed this way, genome doubling spreads were excluded from the analysis.

### rDNA methylation Analysis

For these samples, we cannot assign reads to specific chromosomes, and therefore collapsed all the reads in each sample into a single repeat unit to examine the average methylation level.

Oxford Nanopore Technologies (ONT) whole-genome sequencing data for the HG008 normal duodenum, normal pancreatic, and tumor pancreatic samples was used for rDNA methylation analysis. Raw BAM files containing per-read modification tags were converted to FASTQ format using samtools (v1.23.1) with retention of base modification tags (samtools fastq -T “*”) to preserve methylation information. Reads were aligned to the human 45S rDNA reference sequence (NCBI accession KY962518.1) using minimap2 (v2.26) with ONT specific parameters (-x map-ont).

Alignments were sorted and indexed using samtools. To restrict analyses to primary alignments, supplementary and secondary reads were removed (-F 0x900). CpG methylation frequencies were then quantified using modkit (v0.6.0) with the pileup function, specifying CpG context and combining strands. Resulting per-site methylation calls were processed in R, where sites with coverage <100 reads were excluded. Methylation levels were subsequently aggregated into 200 bp bins across the rDNA reference, and mean methylation per bin was calculated as the ratio of modified to total reads.

#### 45S Methylation Analysis

To investigate the distribution of methylated reads, reads were then aligned to human rDNA reference sequence KY962518.1 using minimap2 (v2.26), and were filtered to keep only alignments covering greater than 70% of the reference sequence at greater than 90% identity. These alignments were separated into individual bam files, which split very long reads that spanned multiple rDNA units, and provided one file per reference-spanning segment. Filtering and separation of reads was performed using samtools (v1.23.1), bedtools (v2.30.0), and python scripts (v3.9.12). The tool modkit (v0.2.0) was used to translate methyl tags into a bedmethyl format, using the filtering parameter to remove the 10% least-confident methylation calls. Average methylation was compiled per-read over the gene body, intergenic spacer, and promoter regions.

#### 5S Methylation Analysis

To investigate methylation across the assembled 5S rDNA array on haplotype 1, reads were aligned to the samples respective assembly using minimap (v2.26) with ONT specific parameters (-x map-ont). Alignments were sorted and indexed using samtools. Secondary reads were removed (-F 256). CpG methylation frequencies were then quantified using modkit (v0.6.0) with the pileup function, specifying CpG context. Reads were sorted and split into per-haplotype methylation calls using the haplotype resolved assembly bed files. For the assembled array on haplotype 1, resulting per-site methylation calls were processed in R, where 5hmC calls were excluded. Methylation levels were subsequently aggregated into 2240 bp bins, matching single monomer unit size, across the rDNA array, and mean methylation per bin was calculated.

#### 5S Array Sequence Identity

Individual repeat copies were extracted as separate FASTA records using bedtools getfasta, with one BED interval per BLAST hit, and aligned within each haplotype independently using MUSCLE (v5.3; default parameters, -align/-output). Aligned sequences were collapsed to single-line FASTA format and loaded into R using seqinr::read.fasta() for all downstream analyses. Repeat copies within each haplotype were reordered according to the strand orientation established above, such that array position 1 corresponds to the telomere-proximal, first-transcribed copy. Per-copy sequence variation was further summarized as pairwise sequence identity between all repeat copies within a haplotype and visualized as a heatmap ordered by array position.

#### 45S rDNA morph identification and prediction

The nodes of the tumor assembly graph were screened for rDNAs using verkko-fillet screen-assembly.sh. It runs MashMap v3.1.1 to find matches with segments over 5kb to human 45S rDNA reference KY962518.1 in nodes over 10kb. Nodes were colored with color_nodes.sh and manually inspected with BandageNG v2025.4.1.

bin/screen-assembly.sh PBhifi+20kbBCMhifi_UL_ONTq26_herro 8

bin/color_nodes.sh assembly_screen.rDNA.ids rDNA “#A020F0” > assembly_screen.rDNA.csv

Additional annotation of the assembly graph was performed using analyse.sh. In brief, nodes are mapped to a homopolymer compressed version of CHM13v2 with MashMap, and for each chromosome, the largest node with mapped size over 5 Mbp and 99% identity is reported in a chromosome assignment csv file, loadable to BandageNG. A secondary search is performed after excluding nodes connected within 1,000 hops from an annotated node, for nodes over 200 kb in length and 99% identity and appended to the csv file. Assembled sequences are searched for telomere motifs with seqtk telo -d50000 option and annotated with an additional telomeric node at the terminus of the assembly graph component (green triangle in Supplementary Fig. 48).

Five rDNA tangles were identified based on the colored rDNA nodes. From the verkko output, assembly.scfmap and assembly.paths.tsv were examined to confirm nodes left as internal 45S rDNA gaps in the final assembly. Node names in each tangle were collected by selecting them in BandageNG and provided as input for manual tangles in Ribotin v1.5, MBG from Verkko v2.3.2, GraphAligner bioconda v1.0.19 and liftoff v1.6.3.

ribotin-verkko \

--graphaligner /path/to/GraphAligner \

--mbg /path/to/verkko/bin/MBG \

-t $cpus \

-x human \

--guess-tangles-using-reference ./template_seqs/chm13_rDNAs.fa \

--orient-by-reference ./template_seqs/rDNA_one_unit.fasta \

--annotation-reference-fasta ./template_seqs/rDNA_one_unit.fasta \

--annotation-gff3 ./template_seqs/rDNA_annotation.gff3 \

-i HG008T_verkko_download/PBhifi+20kbBCMhifi_UL_ONTq26_herro \

-c ../t1.txt -c t2.txt -c t3.txt -c t4.txt -c t5.txt \

-o given_tangle

The predicted morphs from Ribotin were further examined for its validity. The morphgraph.gfa file was examined to exclude spurious rDNA morphs that were truncated, too low in coverage (<30) or disconnected from the heavy rDNA path. These morphs are built from sequencing errors and falsely recruited hifi and ont reads. Partial morphs mapping to the DJ and PJ junction were also excluded.

Acrocentric repeats and rDNAs were annotated with <u>annotate.sh</u> (https://github.com/arangrhie/Scratch/blob/master/Acro/src/annotate.sh) from Ref. ^39^. The rDNA is a rotated version of KY962518.1, which begins at base 41,096 to reflect the actual rDNA position next to the DJ. The $in_fa.fofn contains the path to HG008Tv3.3 split fasta files. annotate.sh $tools/Scratch/Acro/input/annotate_regions.txt HG008T $in_fa.fofn HG008Tv3.3 cat HG008Tv3.3.*/HG008T/HG008T_*.bed > ../HG008Tv3.3.acro_annotation.bed

Full-length rDNA coordinates were extracted and renamed from the HG008Tv3.3 assembly. Full rDNAs were defined when both the mapped reference and query coverage were over 90%. The rDNAs were labeled based on the distance from the DJ and PJ. In case there were more than 2 gaps, rDNA copies in the middle were labeled with M. The rDNAs were named after DJ only when there was no gap inside the assembled rDNA array.

Likewise, acrocentric repeat annotation was performed on CHM13v2 assembly and rDNAs were examined. Only the first rDNA morph of the multiple model rDNA morph copies was retained along with the assembled, diverged full rDNAs.

The collected rDNAs from the predicted Ribotin morphs, HG008Tv3.3 and CHM13v2.0 were split and mapped to the rotated rDNA reference with parsnp v2.1.4 using default parameters. Parsnp builds a phylogenetic tree based on SNPs, which is important because the Indels are the most prominent error types in the Ribotin morphs and the assembled rDNAs (parsnp -r rDNA_ROT.fa -d template_seqs/chm13/*.fa v3.3_units/*.fa morphs/*.fa -p 24 -o parsnp_wi_chm13_to_rot).

## Data availability

All sequencing data presented are publicly accessible in NCBI BioProject (PRJNA200694)^17^. HG008 benchmark data, related alignments, and assembled genomes are also accessible from an NCBI-hosted GIAB FTP site (https://ftp.ncbi.nlm.nih.gov/ReferenceSamples/giab/data_somatic/HG008/). Curations of SVs are in https://github.com/jzook/HG008SVcuration/issues and in Zenodo.^97^ Draft V0.6 somatic SV and CNV benchmark is at https://ftp.ncbi.nlm.nih.gov/ReferenceSamples/giab/data_somatic/HG008/Liss_lab/analysis/NIST_HG008-T_somatic-stvar-CNV_DraftBenchmark_V0.6-20261006. Draft V0.4 somatic small variant benchmark is at https://ftp.ncbi.nlm.nih.gov/ReferenceSamples/giab/data_somatic/HG008/Liss_lab/analysis/NIST_HG008-T_somatic-smvar_DraftBenchmark_V0.4-20261006. The haplotype-resolved HG008 assemblies have been deposited in GenBank. The normal assembly HG008-N v6.3 (BioSample SAMN43682936) is deposited as two chromosome-level haplotype assemblies (hap1: BioProject PRJNA1489689, WGS accession JCANOR000000000, assembly GCA_061372825.1; hap2: BioProject PRJNA1489688, WGS accession JCANOS000000000, assembly GCA_061372785.1.).

The tumor assembly HG008-T v3.2 (BioSample SAMN43682937) is deposited as two scaffold-level haplotype assemblies: (hap1: BioProject PRJNA1489687, WGS accession JCANOP000000000, assembly GCA_061372725.1; hap2: BioProject PRJNA1489686, WGS accession JCANOQ000000000, assembly GCA_061372705.1.) New Fiber-seq and CENP-A DiMeLo-seq and CUT&RUN will be deposited in NCBI BioProject (PRJNA200694).

## Code availability

Custom code for creating benchmarks and analysis scripts, as well as HG008-T_SV_curation_issues.html, which contains an archive of curations of all benchmark SVs, is in Zenodo.^97^ To make this resource more reusable to the community, we have also created skills and agents using the recent Paper2Agent publication’s methods,^98^ which allow anyone to confirm the results in the paper and ask different quantitative questions about the benchmarks and other results. These skills and agents are also available in the Zenodo repository.

## Acknowledgements

Certain commercial equipment, instruments, or materials are identified to specify adequately experimental conditions or reported results. Such identification does not imply recommendation or endorsement by the National Institute of Standards and Technology, nor does it imply that the equipment, instruments, or materials identified are necessarily the best available for the purpose. This work was supported in part by the National Center for Biotechnology Information of the National Library of Medicine (NLM), National Institutes of Health (NIH). This research was supported in part by the Intramural Research Program of the National Institutes of Health (NIH). The contributions of the NIH author(s) are considered Works of the United States Government. The findings and conclusions presented in this paper are those of the author(s) and do not necessarily reflect the views of the NIH or the U.S. Department of Health and Human Services.

## Author Contributions

JMZ, JW, FPB, CX, GAL, and MK designed the study. ASL, HJH, and ZH developed the HG008 cell line. JW, SK, DA, AP, HC, FT, JMZ, JZS, MA, SG, GAL, and CX contributed to genome assembly, evaluation, and correction. MiJ, KHM, MS, DL, and YX generated, basecalled, and error-corrected nanopore sequencing data. SE, GM, JZS, and EMM analyzed Hi-C. ND developed assembly stratifications. NDO generated assembly-based variant calls and regions. AMW, BP, FJS, GN, RLM, HB, JP, JW, LFP, SMES, SC, LM, RL, ZZ, KS, DEC, PCC, AC, PCB, YP, TNY, MFEM contributed to small variant calling and benchmarking. ACE developed Truvari for SV benchmarking. AB, AGK, MK, TA, HB, FT, JeM, ST, DC, YL, JdL, RH, and MFEM contributed to SV and CNV calling and benchmarking. YS and YSa performed benchmark evaluation. JMZ, JW, CX, SMES, ST, SC, DC, YL, LM, RL, ZZ, LFP, MaJ, CB, and MFEM performed variant curation. ECr, MV, and MMc designed and analysed dGH SCREEN experiments. AG performed MHC analysis and contig partitioning. GAL, KKO, and SCC performed centromere analysis. SCC performed CENP-A DiMeLo-seq and CENP-A CUT&RUN. ABS and JR generated Fiber-seq data. ABS and DD performed Fiber-seq analysis, including L1 insertion and centromere analyses. JoM performed mobile element analysis. FPB and TRR performed KaryoScope analysis. JLG, PK, MB, TAP, and AR performed rDNA analysis. JYC, LN, ZS, and JPK performed telomere length analysis and validation. AVZ performed gene annotation for normal and cancer genomes. PV, JNV, and DRR performed RNA analysis. MW contributed to analysis of sex chromosomes. CX, JMZ, JW, MK, CEM, AG, GAL, KKO, SG, JoM, FPB, and TRR wrote and edited the manuscript. All authors reviewed and approved the final manuscript.

## Ethics

Collection of tissue and blood from patients with pancreatic disease was approved by the Mass General Brigham IRB under protocol # 2003P001289. The MGH Data and Tissue Sharing Committee and NIST Research Protections Office determined that the signed consent explicitly allows for sharing of the participants’ genomic data and development and distribution of cell lines. Relevant excerpts from the patient consent were published with the previous data for HG008.^5^

## Corresponding Author

Correspondence and requests for materials should be addressed to Justin M. Zook.

## Generative AI Use

Analyses for and editing of this manuscript were assisted by Claude Code (Sonnet 4.6 to 5.5 and Opus 4.6 to 5.5), ChatGPT 5.6 Sol, and Gemini 3.1 Pro. We have created a Zenodo repository containing the scripts and analysis logs from using Claude Code, curations, and Paper2Agent results, so that the methods can be checked by other humans and agents. To help ensure accuracy of the results, humans monitored the work and manually checked a subset of sites. We also used different models (e.g. GPT 5.6 Sol) to critically evaluate the scripts and results, and relevant suggestions were incorporated. LLMs assisted in writing some methods sections performed by Claude Code, but were reviewed by humans. For the benchmarks, we performed additional human evaluations, including curation of differences between other callsets and the benchmarks by multiple laboratories. We have also made multiple versions of the benchmarks available to the public and received both positive feedback and suggestions for improvements. All content, scientific claims, and conclusions have been reviewed and verified by the authors to ensure accuracy and originality.

## Extended Data

**Extended Data Fig. 1:**
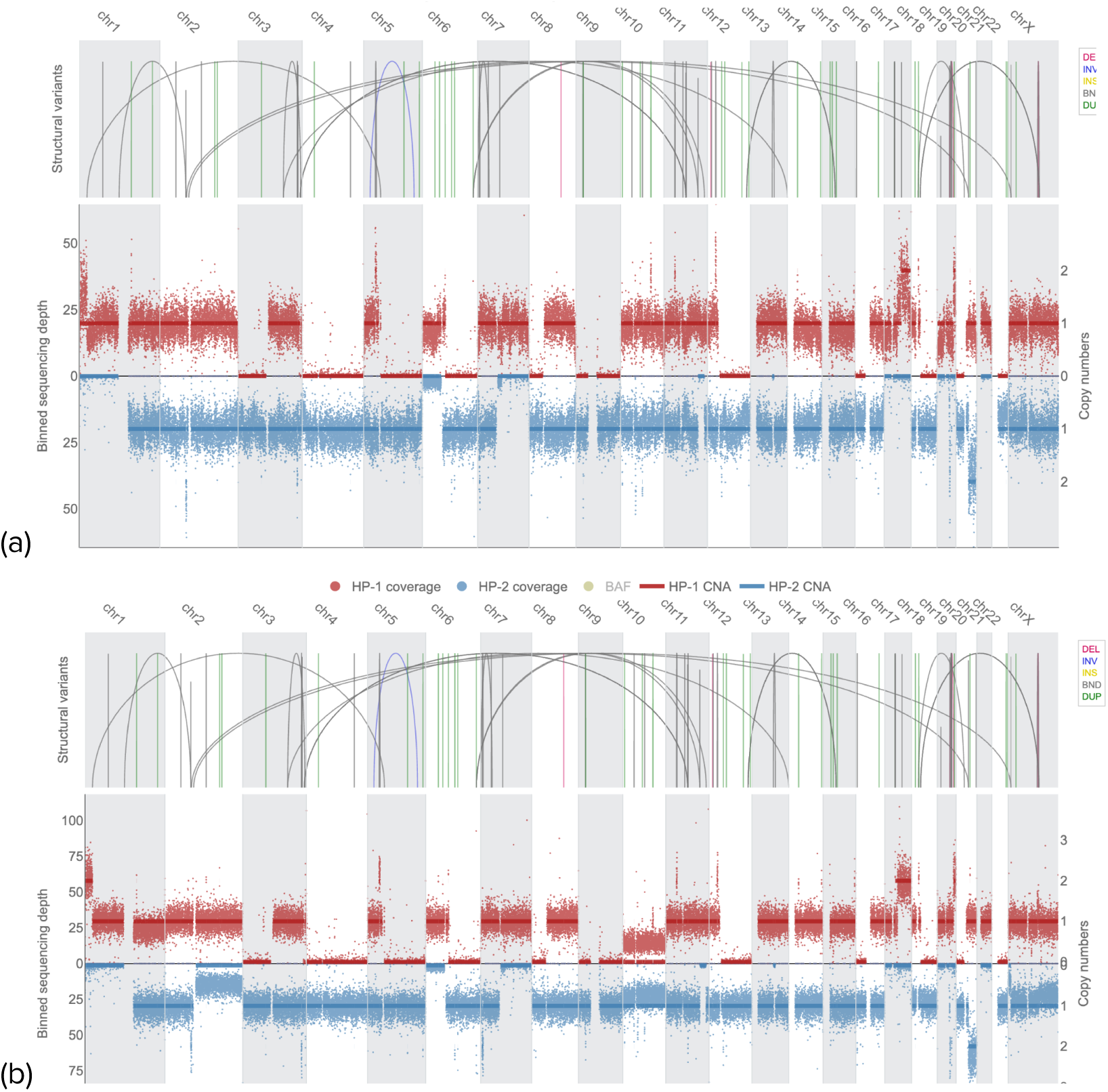
Haplotype-resolved copy number from long reads demonstrates many large deletions and fewer large duplications and copy neutral loss of heterozygosity. (a) Copy number from NIST passage 21 HiFi data, which contains no large high frequency subclonal CNVs and closely matches the assembly. (b) Copy number from MGH passage 23 ONT data, which contains some large high frequency subclonal CNVs (e.g., 1q, 2q, 10, X) that are not contained in the assembly because they are not truncal. While most truncal chromosome-scale CNVs were deletions, a large duplication on chr20_hap2 resulted in 3 total copies. Also, the only remaining haplotype of chr1p_hap1 and chr17q_hap1 had large partial duplications caused by unbalanced translocations that result in CNLOH. Smaller copy number changes were caused by tandem duplications, large deletions, or loss or gain of sequences between translocation breakpoints. Chr19 has the largest variation in copy number due to nested foldback inversions. Phasing copy number changes enables us to see that chr6_hap2 lost most of the p arm in most cells due to a translocation with chr8_hap2 and chr6_hap1 lost most of the q arm due to a translocation with chr7_hap1 in the centromeres.

**Extended Data Fig. 2:**
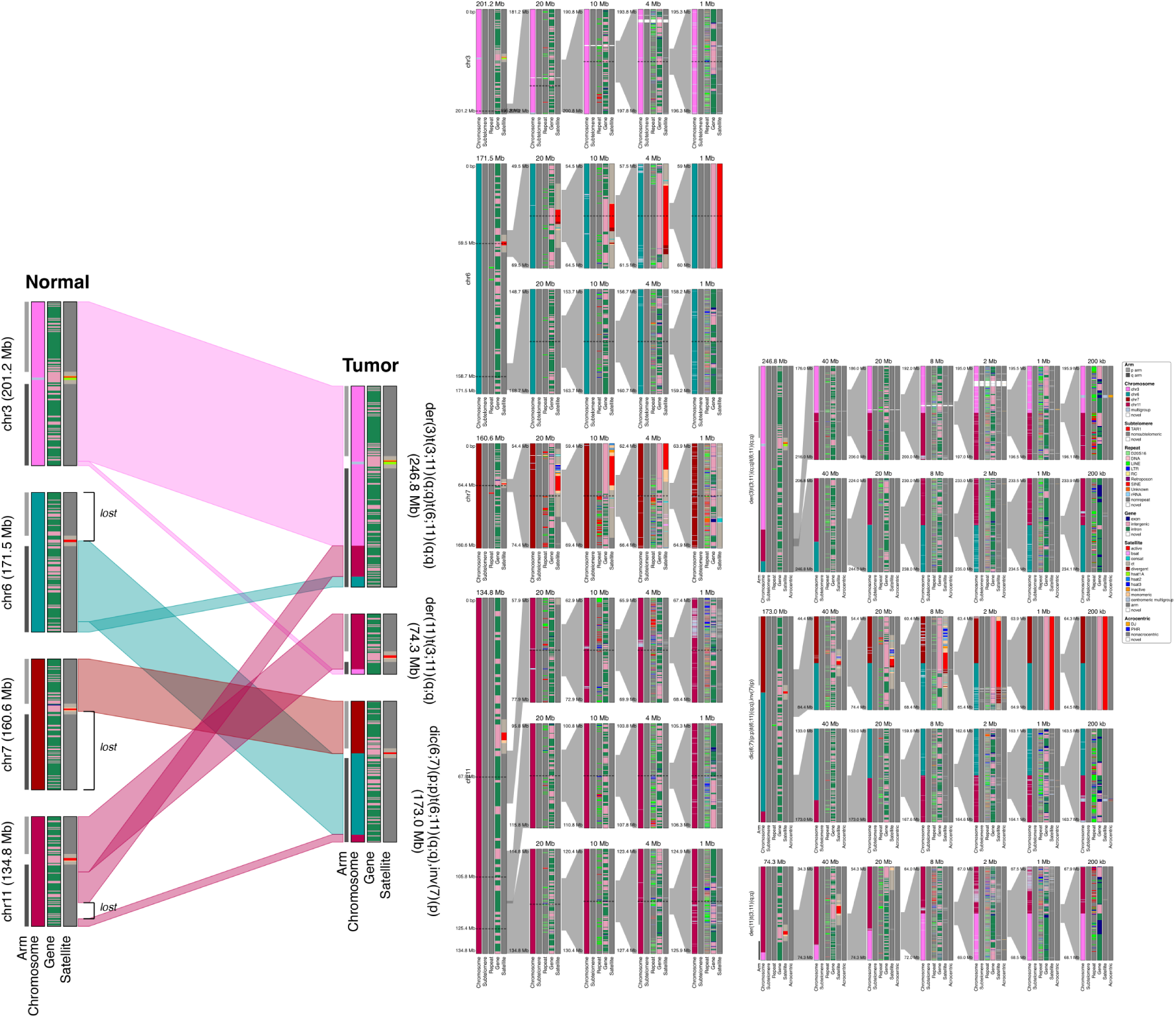
Chromoplexy causing a complex series of events linking chromosomes 3, 6, 7, and 11 in three hybrid tumor chromosomes. Karyoscope’s chromosome, subtelomere, repeat, gene, satellite annotations are shown across the entire chromosomes and at different resolutions around each interchromosomal breakpoint. Note that inversions, including the one on chr7p, are not shown by karyoscope.

**Extended Data Fig. 3:**
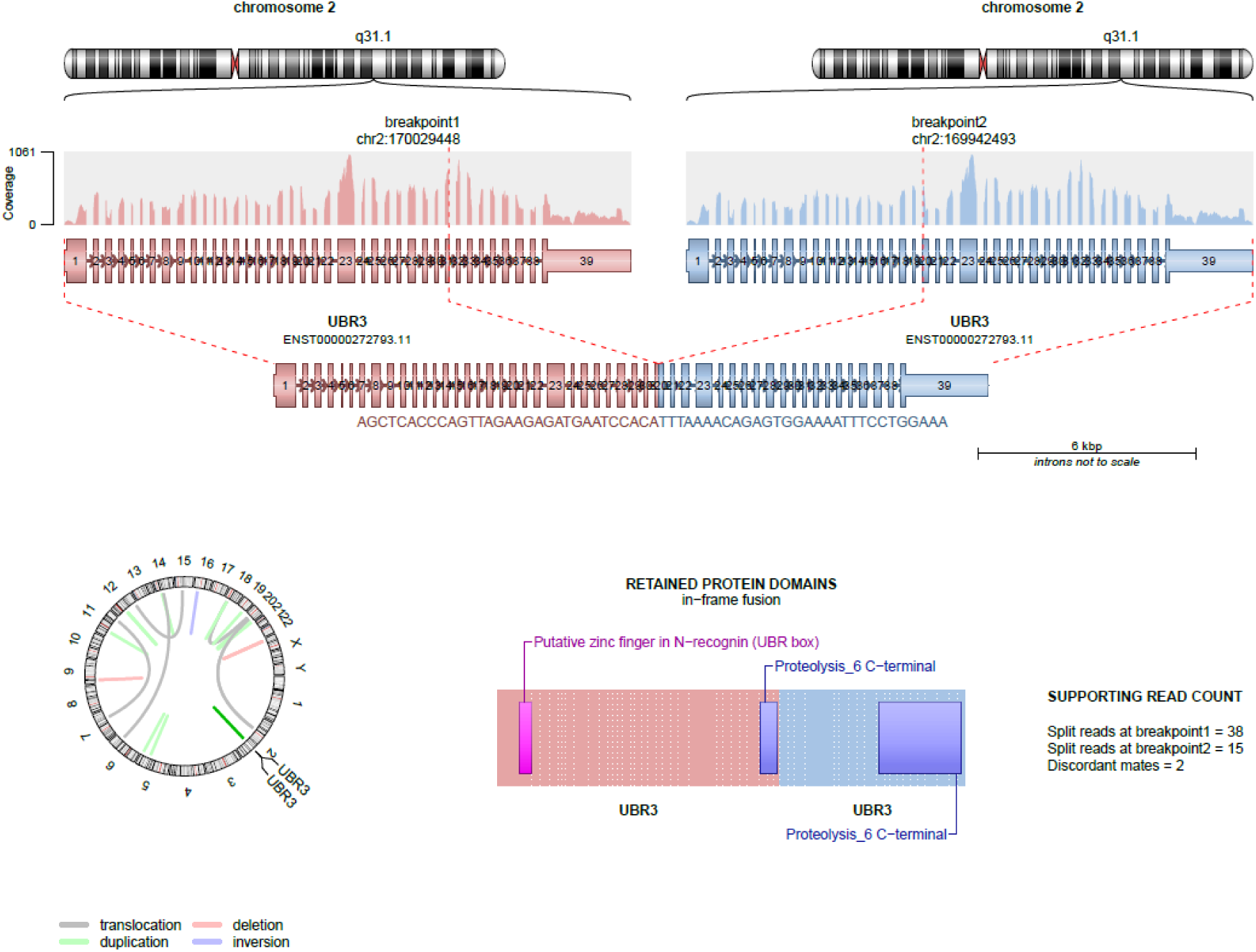
Expression of 92 kbp internal tandem duplication in *UBR3*. Internal tandem duplications in genes include *NFE2L2*, which has ITDs in NSCLC,^99^ *UBR3*, which can be important for genome stability,^100^, and *BBS9*. Fusion analysis of HG008-T RNA-seq also supports the detection of *UBR3* internal tandem duplication.

**Extended Data Fig. 4.**
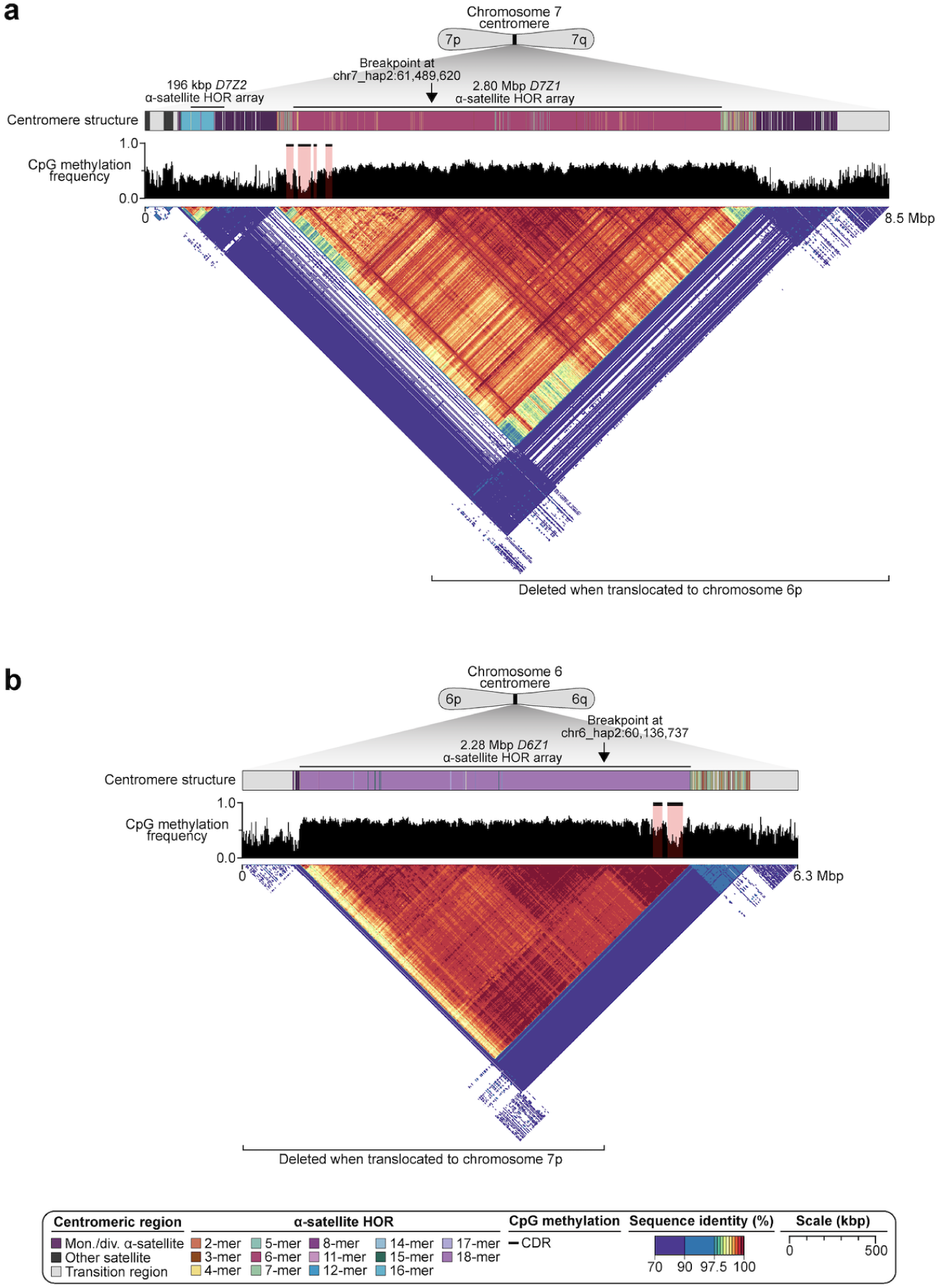
The sequence, structure, and epigenetic landscape of the chromosome 7 and 6 centromeres in normal cells. For both chromosomes a) 7 and b) 6, the CDR sites are retained upon translocation, while most of the α-satellite HOR array is deleted. This creates a dynamic dicentric chromosome with one or two active centromeres (Fig. 4a).

**Extended Data Fig. 5:**
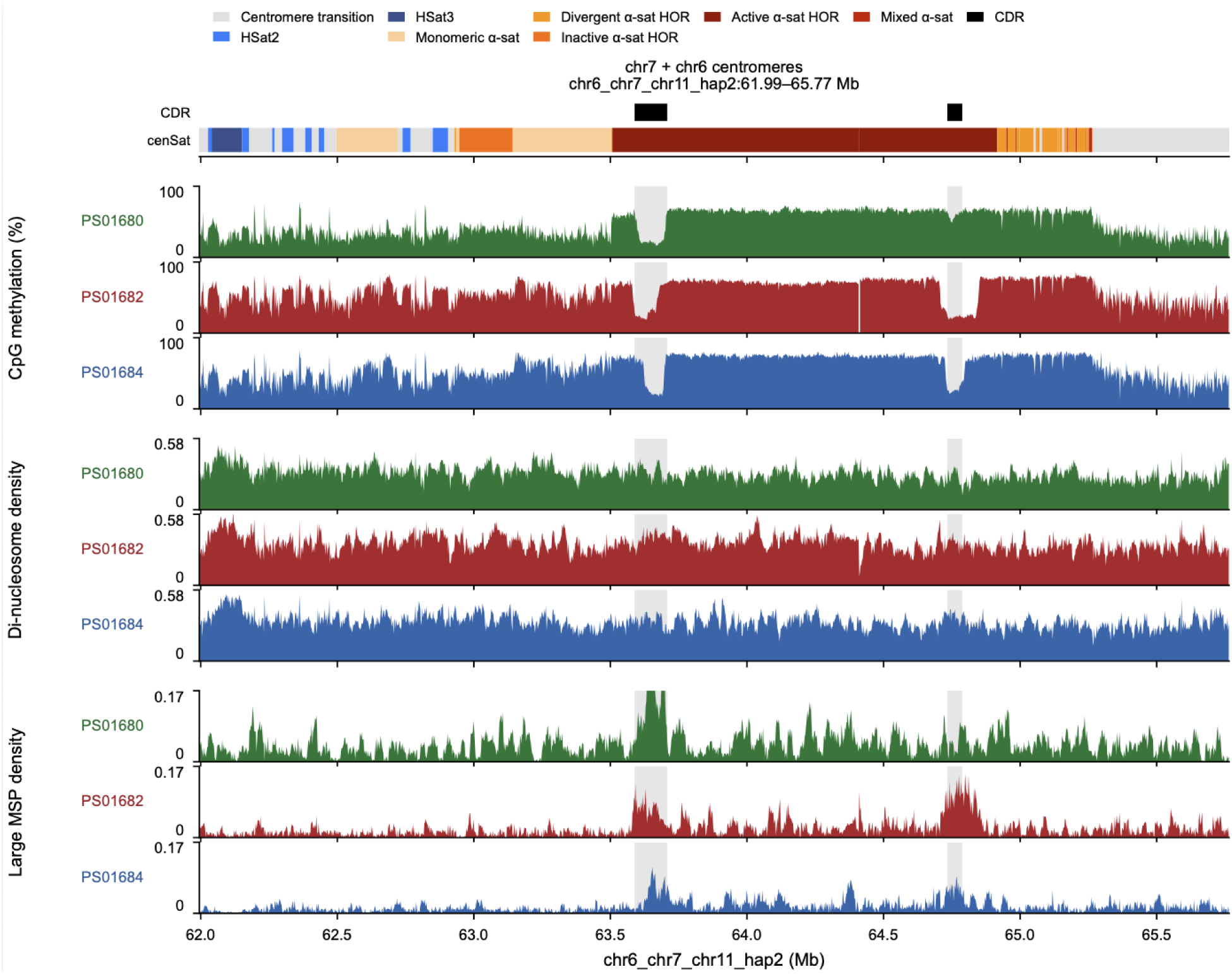
CpG methylation and Fiber-seq chromatin features across the dynamic dicentric chr6–chr7 centromere fusion in three HG008-T clones. Genome browser view of the HG008-T assembly contig chr6_chr7_chr11_hap2 (61.99–65.77 Mb). The view spans the fused chr7 and chr6 α-satellite arrays and 500 kb of flanking sequence on each side. Top, centromeric satellite (cenSat) annotation coloured by satellite class. The chr7 (63.51–64.41 Mb) and chr6 (64.41–64.92 Mb) active α-satellite higher-order repeat (HOR) arrays each contain a centromere dip region (CDR; black bars; 63.59–63.71 Mb and 64.73–64.79 Mb), and grey shading marks the CDRs in every track. Below, one track per clone (PS01680, green; PS01682, red; PS01684, blue) shows three measurements. CpG methylation is the percentage of 5mC calls with probability ≥ 0.5 at each CpG, combining both strands and averaged over the CpGs in a bin. Di-nucleosome density is the number of nucleosome footprints longer than 210 bp covering a base, divided by the number of molecules covering it. Large methyltransferase-sensitive patch (MSP) density is calculated the same way for MSPs longer than 150 bp. Values are means in 2.5-kb bins without smoothing. PS01680 has the CpG methylation profile most similar to that of the cells used for CENP-A measurements. PS01682 is a unique clone different from most bulk cells that does not have the chr6-chr7 translocation, and has large signals associated with kinetochores for both centromeres as expected.

**Extended Data Fig. 6:**
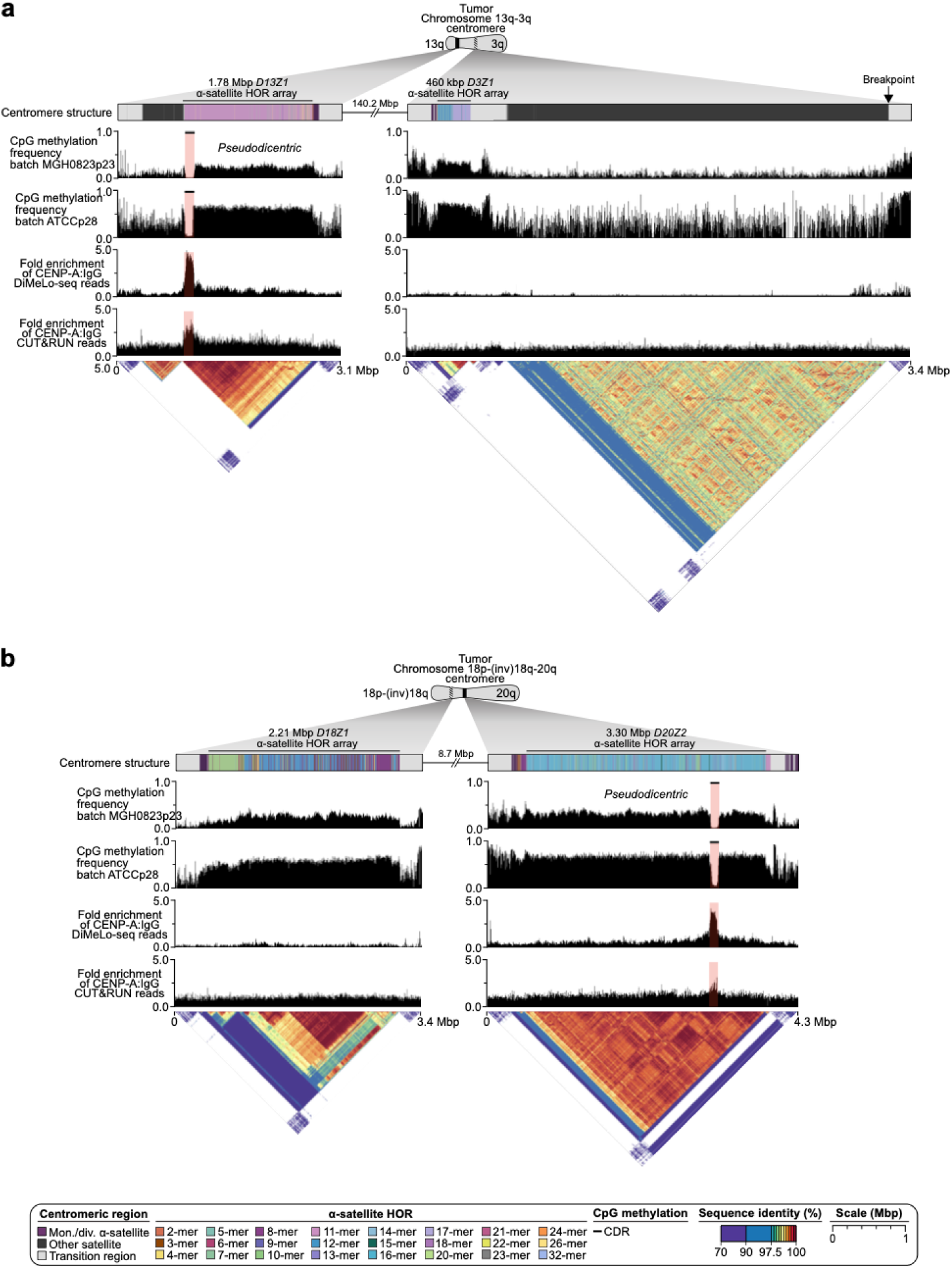
Plasticity of human centromeres during tumorigenesis in translocations with inversions. The top methylation track matches cell batch MGH0823p23 used for the assembly, whereas the second track matches the ATCC p28 cell batch used for CENP-A DiMeLo-seq and CUT&RUN measurements of kinetochore protein binding. CDRs are low methylation regions associated with the site of kinetochore assembly, highlighted by black bars and pink rectangles in the CpG methylation track. (a) The chr3 CDR is deleted by the translocation+inversion, and CENP-A only binds to the chr13 CDR. (b) The chr18 CDR is methylated and inside the inverted region, and CENP-A only binds to the chr20 CDR.

**Extended Data Fig. 7:**
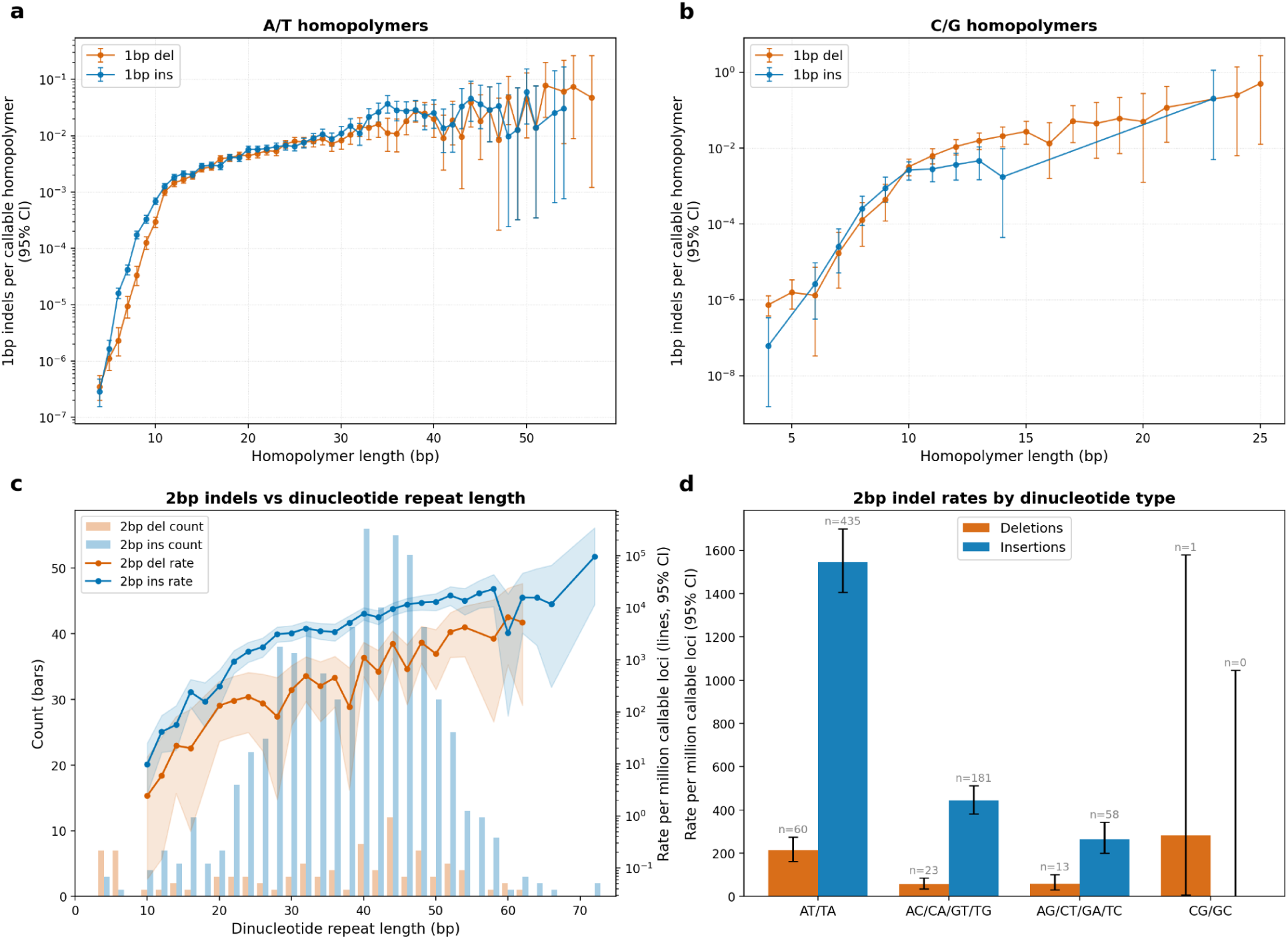
Somatic indel rates increase with homopolymer and dinucleotide TR length. Primary 1 bp indel rates per callable homopolymer versus homopolymer length for (a) A/T and (b) C/G homopolymers. Rates increase roughly exponentially from 4 to 11 bp (>3,000-fold for A/T and >8,000-fold for C/G deletions). At the same homopolymer length, deletion rates are higher for C/G than for A/T, though far fewer C/G homopolymers are in the human genome. For 1 bp indels occurred in C/G homopolymers, there were 139 deletions and 57 insertions; overall DEL/INS = 2.44, 95% CI [1.78–3.38]. Short (4 to 6 bp) C/G homopolymers showed a strong deletion bias (DEL/INS = 6.33, 95% CI [1.86–33.4]), as did long (>10 bp) ones (DEL/INS = 4.09, 95% CI [2.57–6.76]). 10 bp C/G homopolymers were balanced (DEL/INS = 1.21, 95% CI [0.56–2.66]). The 7 to 9 bp range had a non-significant trend toward insertion bias (INS/DEL = 1.89, 95% CI [0.80–4.81]). For 1 bp indels occurred in A/T homopolymers, there were 3,831 deletions and 4,685 insertions; overall INS/DEL = 1.22, 95% CI [1.17–1.28]. Short A/T homopolymers (≤13 bp) had a significant insertion bias (INS/DEL = 1.76, 95% CI [1.62–1.92]), strongest at 6 bp (INS/DEL = 7.0, 95% CI [3.9–13.6]). In contrast, longer A/T homopolymers (≥15 bp) showed only a slight insertion bias (INS/DEL = 1.07, 95% CI [1.01–1.12]). Homopolymers of 33–37 bp showed an unexpected insertion excess (INS/DEL = 1.95, 95% CI [1.46–2.61]). (c) Counts (bars, left axis) and rates per million callable loci (lines with 95% CI bands, right log axis) of 2 bp insertions and deletions versus dinucleotide repeat length; rates increase with repeat length, and insertion rates exceed deletion rates at most lengths. (d) 2 bp indel rates per million callable dinucleotide-repeat loci by dinucleotide type (95% CI). Insertions exceed deletions for AT/TA (435 vs. 60), AC/CA/GT/TG (181 vs. 23) and AG/CT/GA/TC (58 vs. 13). The AT/TA insertion rate is highest, at about 1,550 per million loci. Error bars and bands show 95% exact-Poisson CIs on rates; INS/DEL ratio CIs are 95% exact conditional-binomial (Clopper–Pearson) intervals.

**Extended Data Fig S8:**
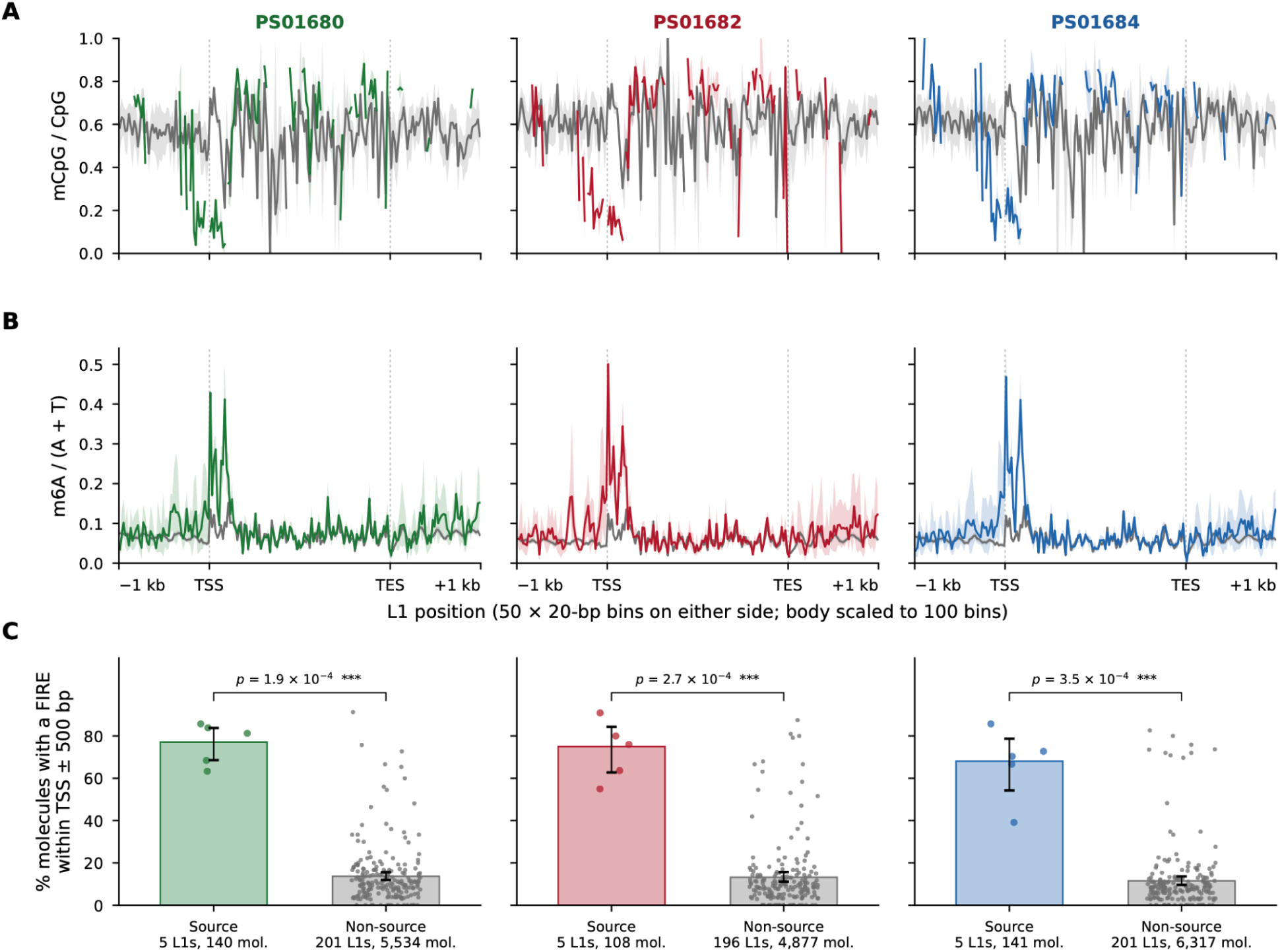
CpG methylation, chromatin accessibility and promoter FIRE occupancy at source and non-source full-length L1 elements using Fiber-seq. Columns show three HG008-T tumor cell line clones (PS01680, green; PS01682, red; PS01684, blue). Source L1s (n = 5) are drawn in each clone’s colour and non-source L1s (n = 201) in grey. (A, B) Strand-oriented metaplots across full-length L1s of (A) the fraction of methylated CpGs (mCpG/CpG) and (B) the fraction of adenines carrying m6A (m6A/(A+T), Fiber-seq), a measure of chromatin accessibility. Each plot spans 1 kb upstream of the transcription start site (TSS; L1 5′ end) through 1 kb downstream of the transcription end site (TES; 3′ end). The flanks are shown in 20-bp bins and the element body is scaled to 100 bins. Lines show values pooled across all molecules and elements of each group, and shaded bands show 95% bootstrap confidence intervals over elements. Note increased accessibility (m6A) corresponding to hypomethylation (mCpG) in source L1s proximal to TSS. (C) Percentage of Fiber-seq molecules spanning the TSS ± 500 bp that carry a Fiber-seq inferred regulatory element (FIRE; FIRE score ≥ 229) within that window. Bars show the pooled percentage per group, with error bars giving 95% bootstrap confidence intervals over elements. Dots show individual source or non-source L1 elements with ≥ 5 spanning molecules, and labels under the bars give the number of elements and molecules analysed. P values are from two-sided Mann–Whitney U tests between source and non-source elements within each clone. Thresholds and calculations are described in Methods.

**Extended Data Fig. 9:**
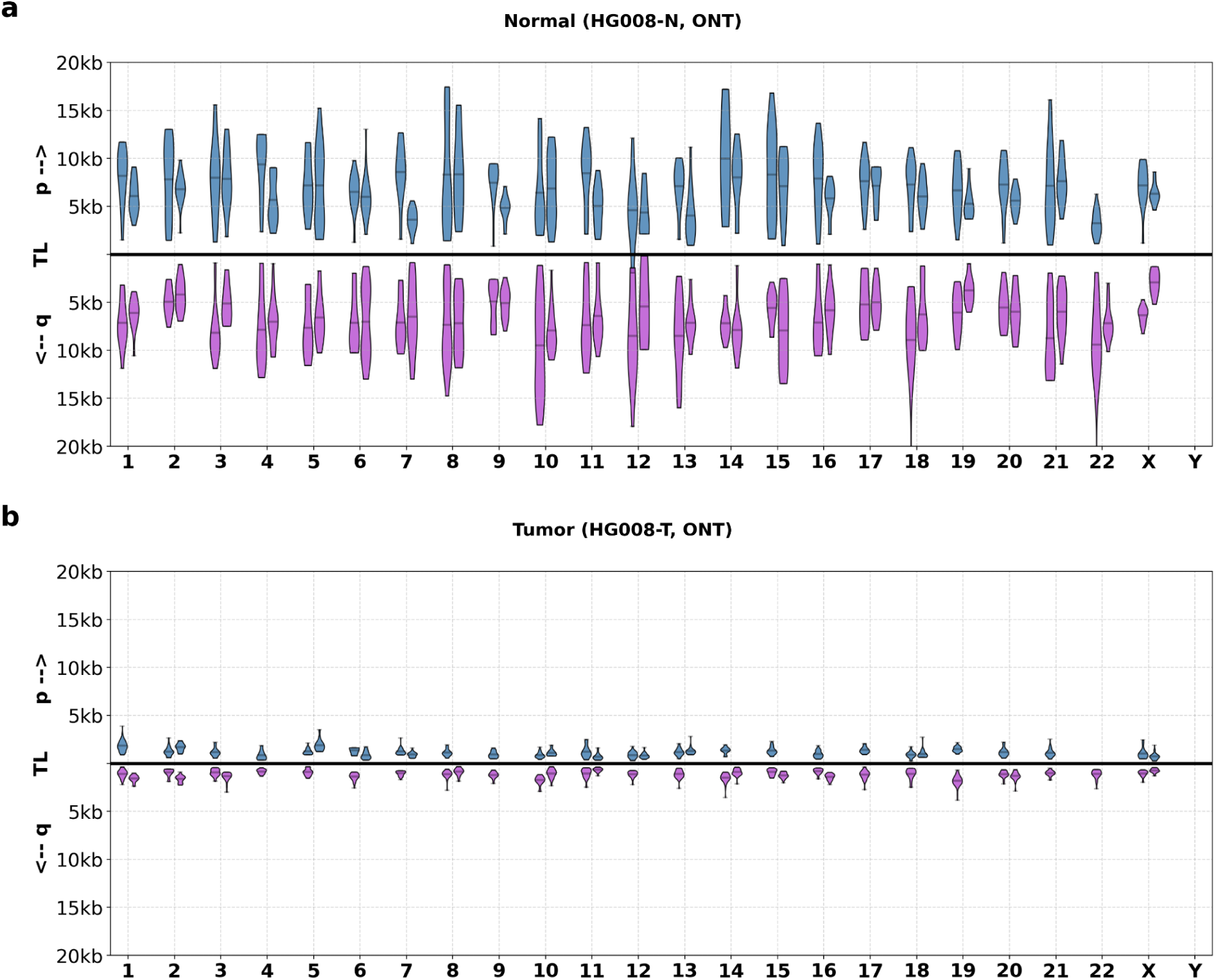
Estimated lengths of telomeres in normal pancreatic tissue and tumor cell line using Telogator with telomeres assigned based on alignments to the assemblies. (a) Telomere lengths of all individual alleles (except unresolved 21/22p) in HG008-N-P, as measured from ONT long reads. (b) Telomere lengths of 68 alleles identified in HG008-T, as measured from ONT long reads. The absent alleles correspond to telomere ends that are not present in the tumor genome due to deletions or chromosome fusions.

