## Supplementary Figures 1-56 for "A near-complete human pancreatic cancer genome"

### A near-complete human pancreatic cancer genome Supplemental Materials

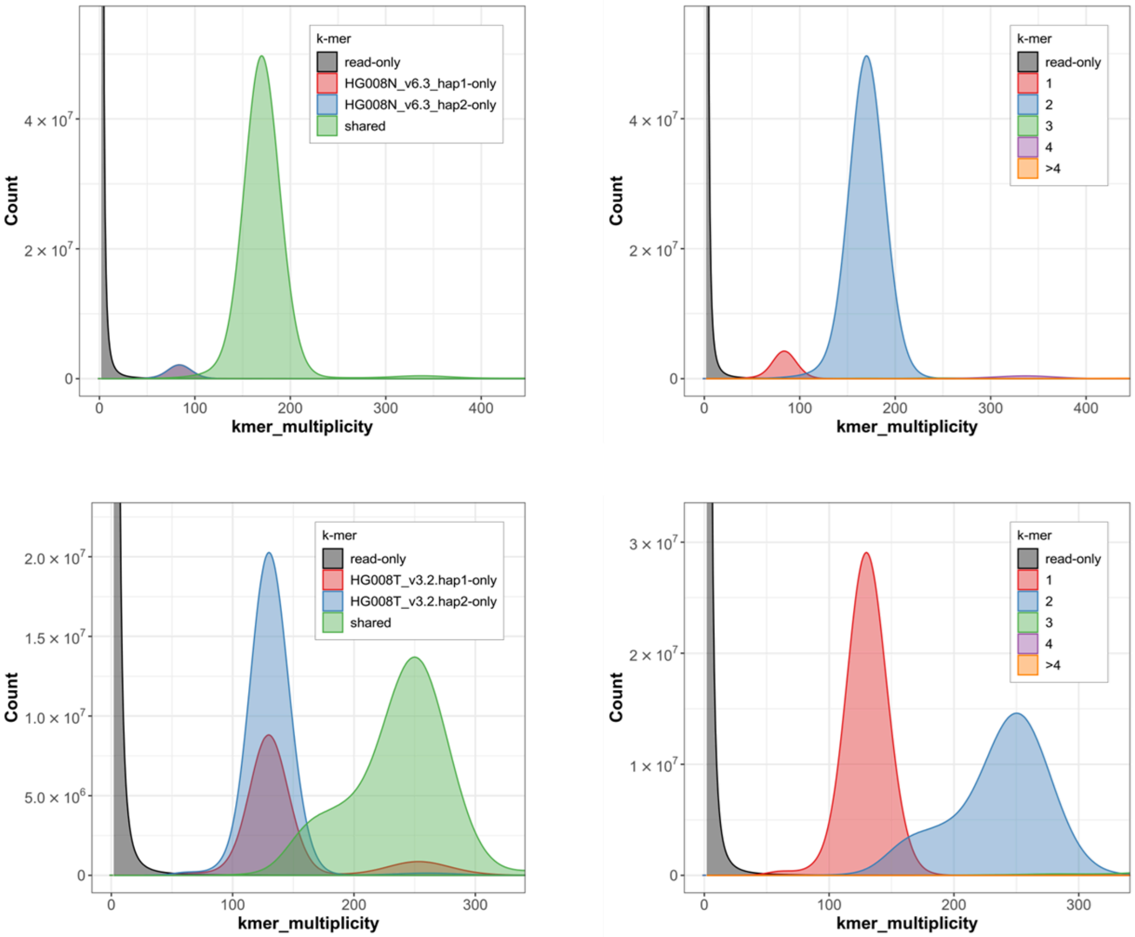


**Supplementary Fig. 1. Kmer-based assembly quality evaluation (QV=70 for HG008-N, top, and QV=72 for HG008-T, bottom) using Merqury[101] (k=31 with hybrid database from PacBio HiFi and Illumina reads).**

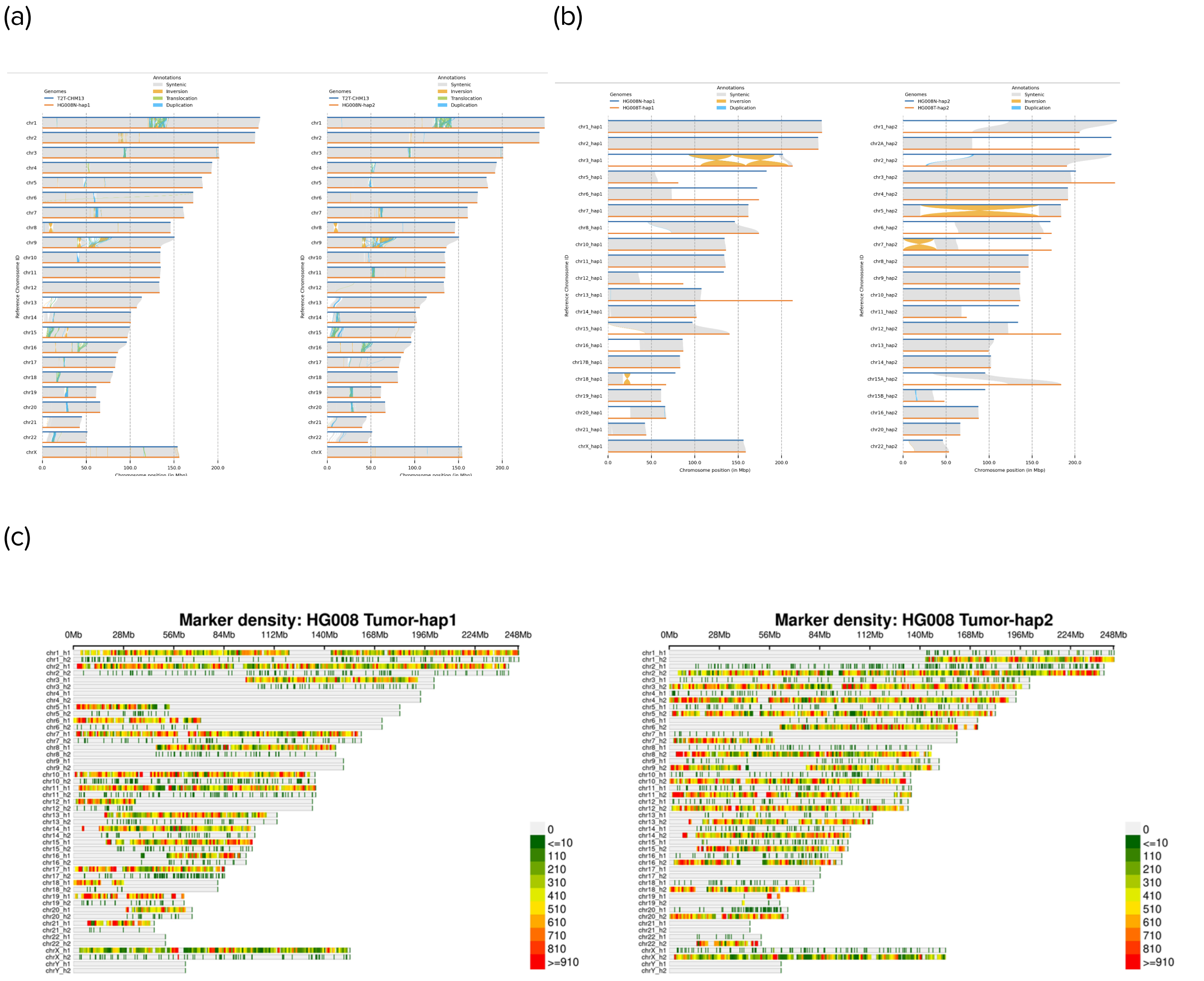


**Supplementary Fig. 2: Tumor and normal assembly evaluations.** (a) Assembly alignments of HG008N_hap1 and HG008N_hap2 to T2T-CHM13 reveal both structural similarities and dissimilarities by SyRI plot; (b) Alignments of matched tumor haplotype with normal haplotype assemblies shows large structural variations by SyRI; and (c) Distribution of normal haplotype-specific markers in each HG008T haplotype. Haplotype markers match between tumor and normal except for chr19, where the last 8 Mbp on tumor hap2 matches normal hap1 after BFB (red markers on chr19_h1 in the right plot for Tumor-hap2). This tumor contig derived from different normal haplotypes was supported by Hi-C reads and CNLOH in this region.

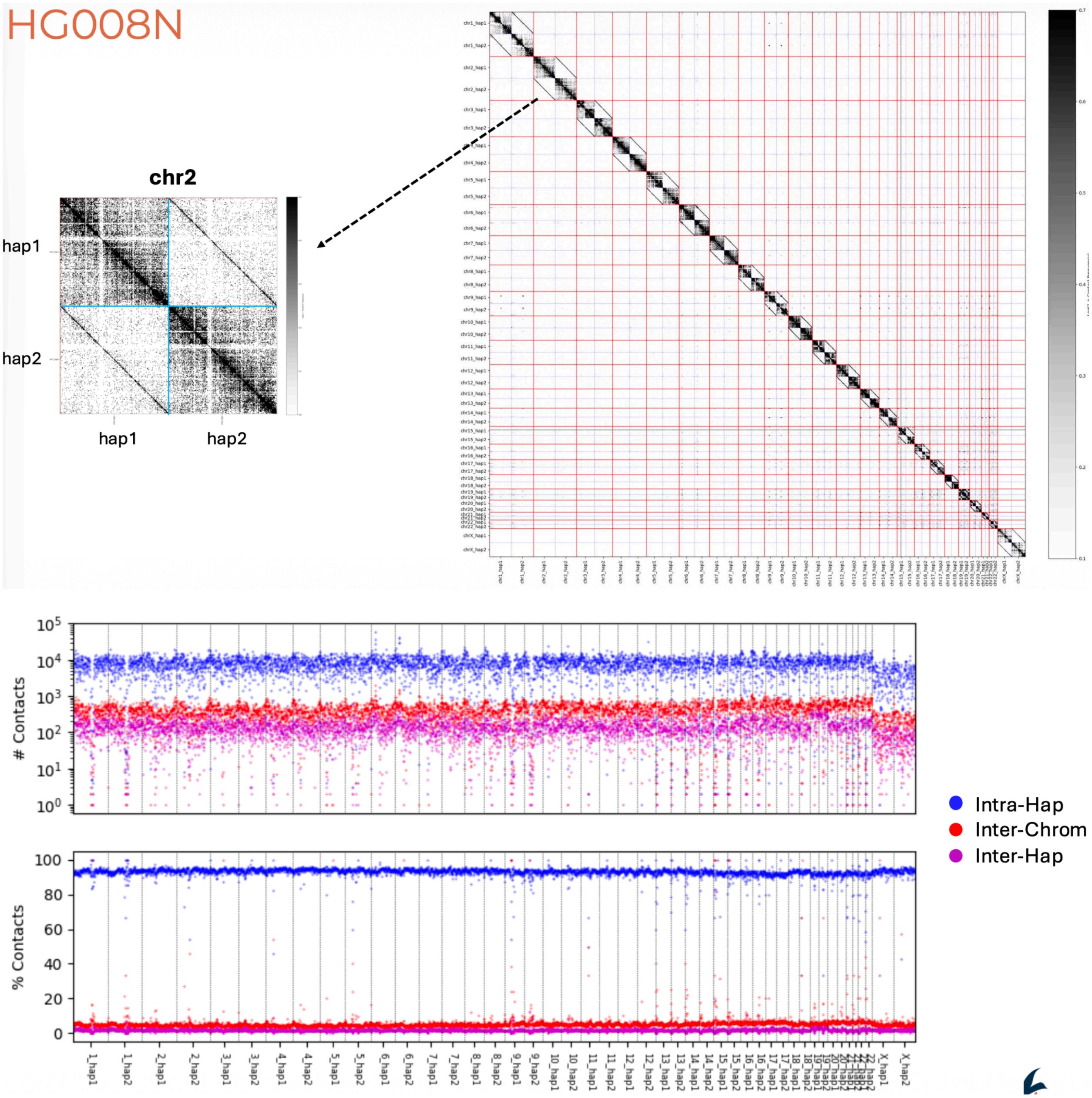


**Supplementary Fig. 3: Phased mapping of Dovetail Hi-C to diploid normal assembly confirms structural and phasing accuracy.**

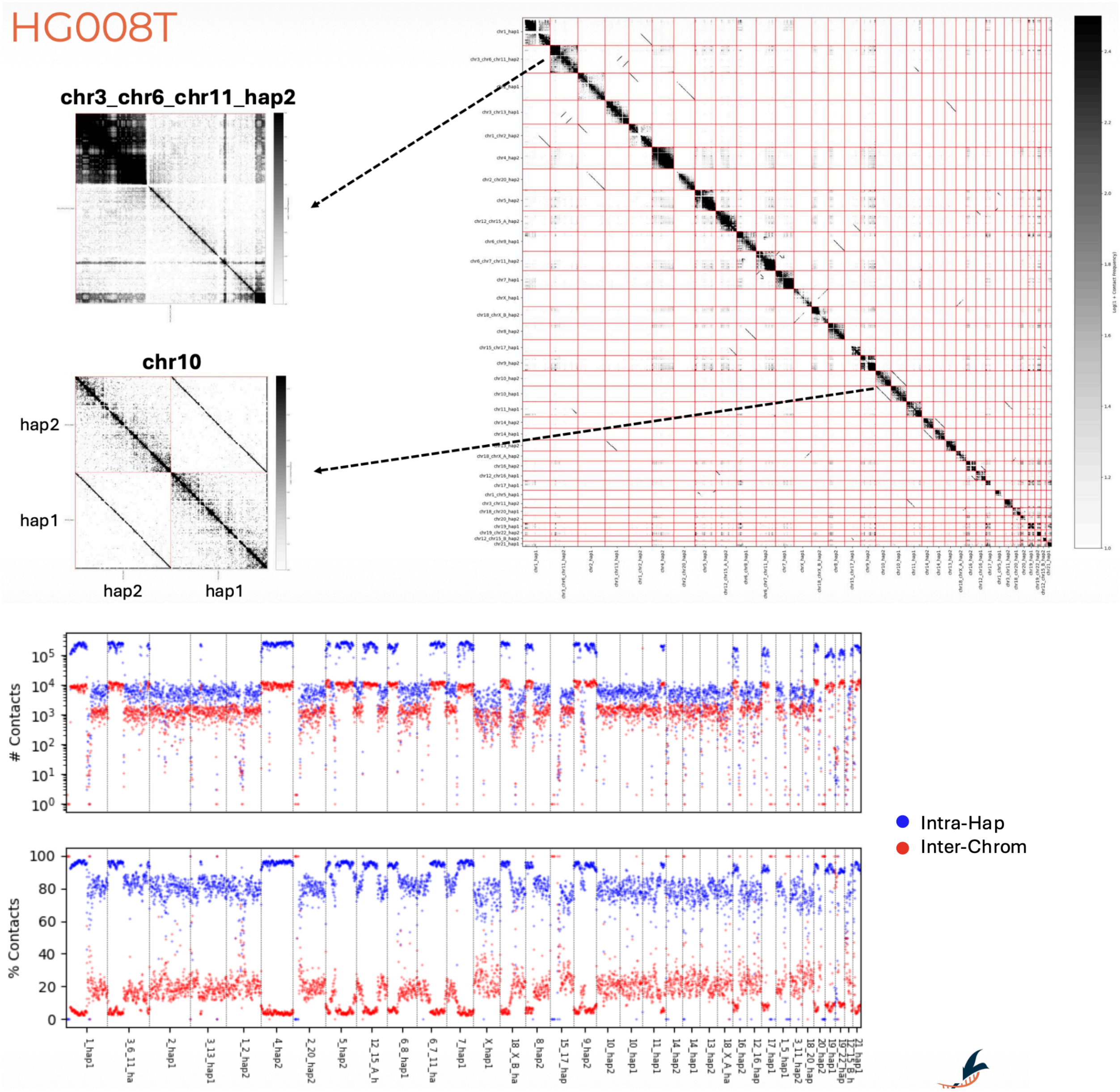


**Supplementary Fig. 4: Phased mapping of Dovetail Hi-C to diploid tumor assembly confirms structural and phasing accuracy.**

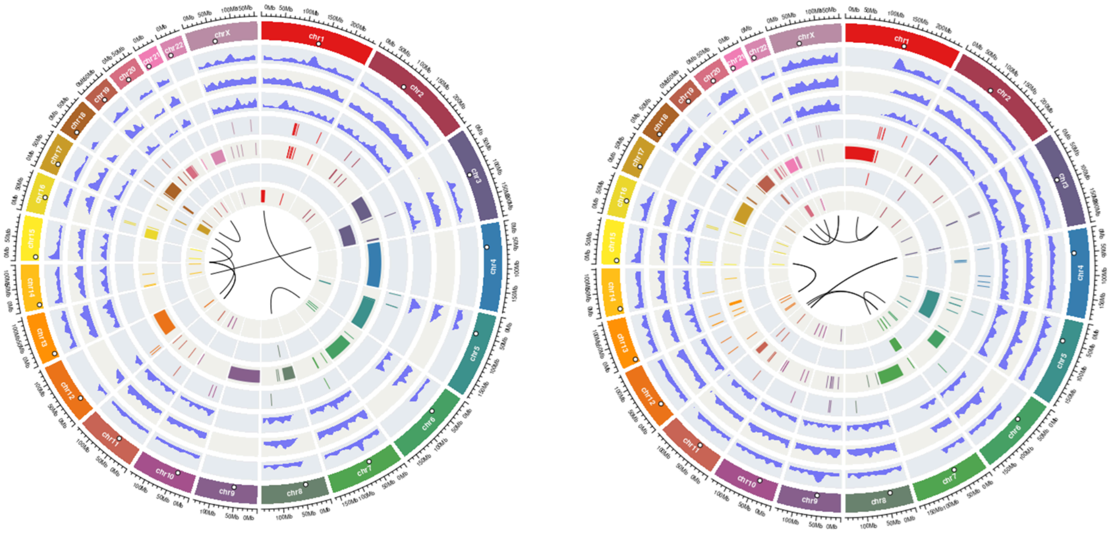

**Supplementary Fig. 5:** Overview of haplotype-specific somatic SNVs (density plot with default 10 Mbp window) and SVs from HG008T-hap1 (left) and HG008T-hap2 (right). HG008N centromere positions are indicated by white circles on each chromosome. The track order (from outer to inner) is density of somatic SNVs, insertions <50 bp, deletions <50 bp, insertions ≥50 bp, deletions ≥50 bp, inversions, duplications, and translocations, respectively.

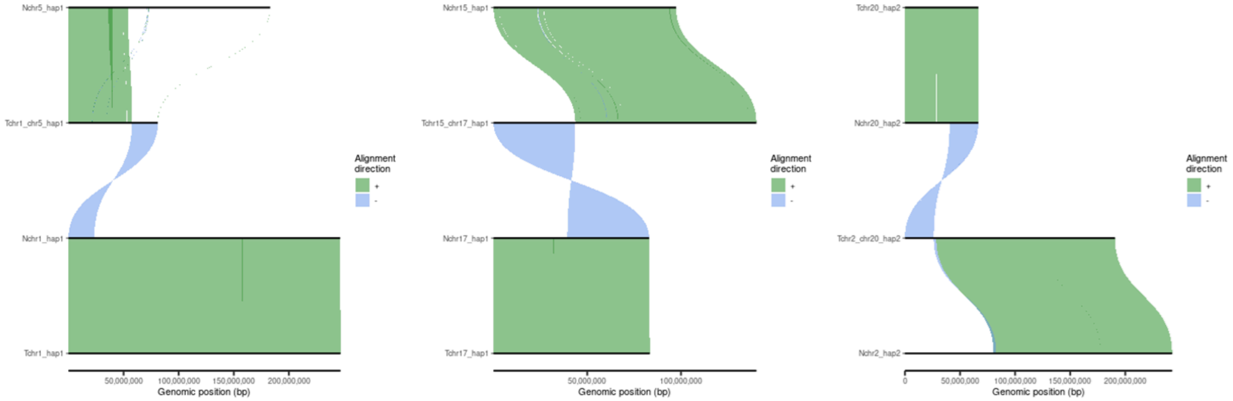


**Supplementary Fig. 6: While most large somatic CNVs were deletions, large duplications caused by translocation on chr1_hap1 and chr17_hap1 caused CNLOH, and a duplication on chr20_hap2 resulted in 3 total copies.**

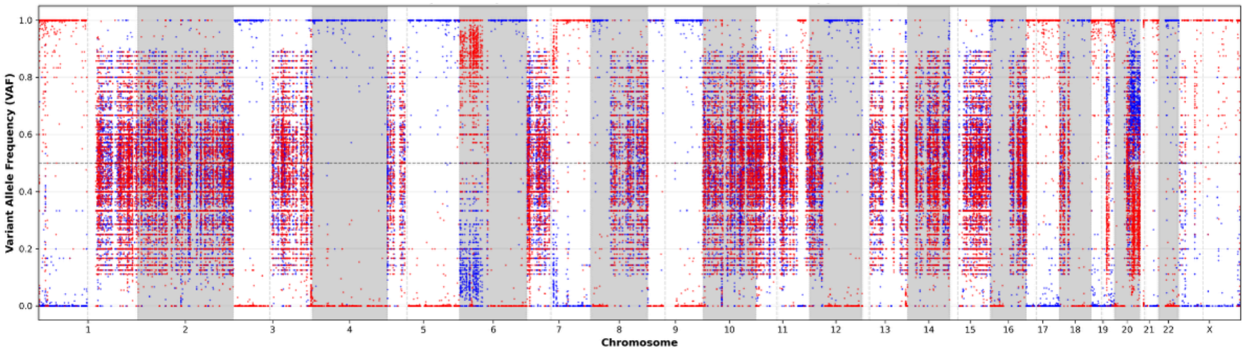


**Supplementary Fig. 7: VAF of heterozygous SNVs in RNA sequencing shows allele-specific expression of haplotype 1 (red) and haplotype 2 (blue).** RNA sequencing was performed on NIST passage 21 cells, and phase information for the SNVs was inferred from genomic data. The complete silencing across chromosome X demonstrates that the tumor cells retained silencing despite a balanced chr18_hap2/chrX_hap2 translocation involving the silenced copy of chrX. Because XIST is on the portion of chrX attached to chr18q, we initially hypothesized that chr18q haplotype 2 might also be silenced; however, silencing did not spread to most of the autosome, as chr18q retains expression of many haplotype 2 genes. The absence of haplotype 1 expression on chr18q is explained by a separate event: a complex unbalanced translocation between chr18 and chr20, together with a large inversion on haplotype 1, deletes chr18q on haplotype 1. Interestingly, the chr18/chr20 breakpoint lies only ~30 kbp from the chr18/chrX translocation breakpoint on haplotype 2 (https://github.com/jzook/HG008SVcuration/issues/272). In addition, chrXp haplotype 2 expression remains repressed even though XIST is no longer on this hybrid chromosome, consistent with maintenance of the established epigenetic state. Overall, the VAF from RNA-seq is consistent with the VAF from genome sequencing.

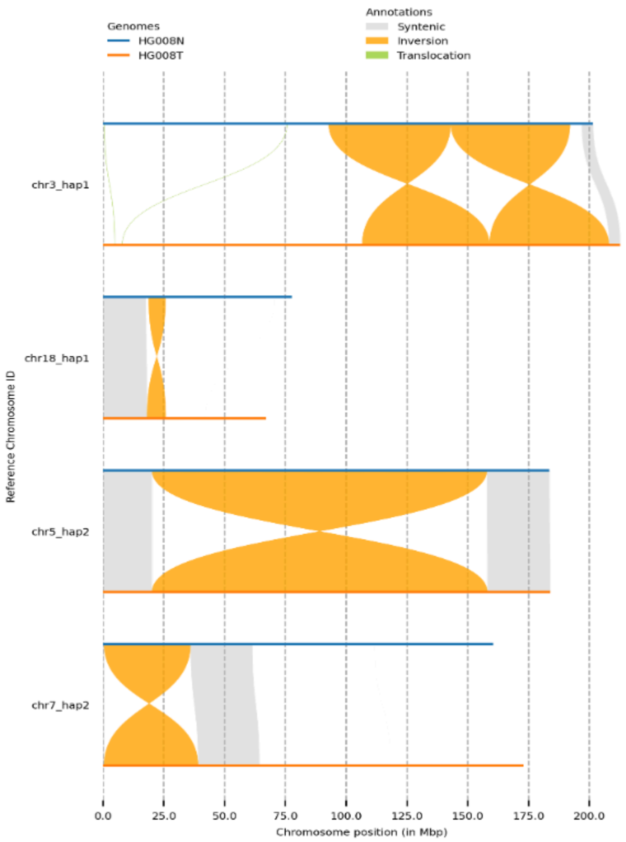


**Supplementary Fig. 8: HG008 contains 3 reciprocal inversions, nine non-reciprocal foldback inversions, five non-reciprocal inversions that do not cause duplications, and one inversion related to a tandem duplication.** The reciprocal inversions are 2.7 kbp (chr3_hap2), 36 Mbp (chr7_hap2), and 138 Mbp (chr5_hap2) in size, and all result in loss of 1 bp to 363 bp of sequence at each breakpoint. 9 foldback inversions cause 537 bp to 2.32 Mbp of duplicated sequence near interchromosomal translocation breakpoints, including 4 nested events on chr19 discussed below. Also, two occur in chr1 and chr8 centromeric active HORs and one in acrocentric short arm repeats near the beginning of chr15. 5 inversions associated with translocations do not cause duplications, with 2 smaller than 1 kbp and others 8, 54, and 99 Mbp. The 8 Mbp inversion at chr18_hap1 translocation with chr20_hap1 can only be seen precisely in the normal assembly because one breakpoint of the inversion is in the chr18 centromeric active HOR. The chr3-chr13 translocation has complex 54 and 99 Mbp inversions on chr3_hap1, also with one breakpoint in centromeric HSat1A.  While not a translocation-associated inversion, there is also a 4,577 bp inverted duplication of sequence starting approximately 3 kbp past the end of a 365 kbp tandem duplication on chr17 that is inserted between the two copies of the tandem duplication (https://github.com/jzook/HG008SVcuration/issues/266). The only non-complex translocations are the chr1_hap1/chr5_hap1 and chr12_hap1/chr16_hap1 unbalanced translocations as well as chr12_hap2/chr15_hap2 (which disrupts the KDM2B gene) and chr18_hap2/chrX_hap2 balanced translocations (Fig. 1b).

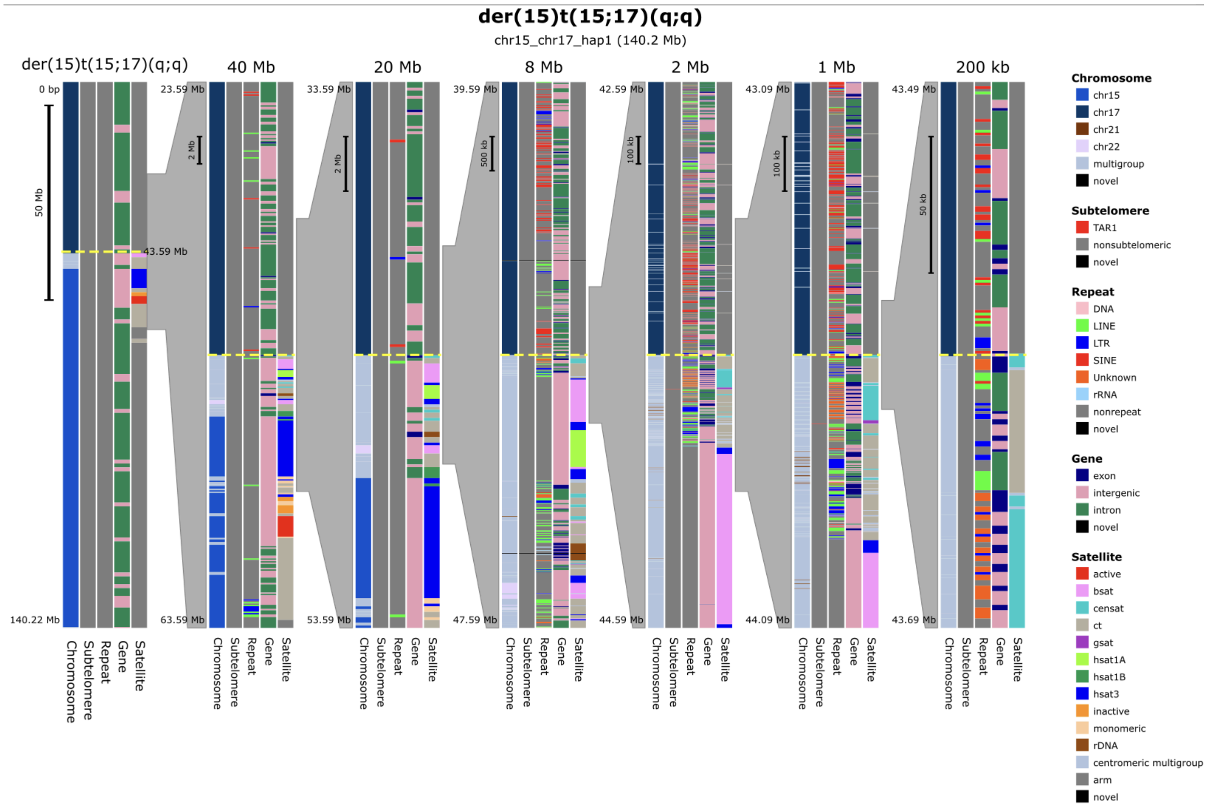


**Supplementary Fig. 9: der(15)t(15;17)(q;q) example of acrocentric breakpoints not resolvable without a normal assembly.** This event is caused by a translocation occurring in an acrocentric short arm near the p arm telomere of chr15_hap1, which is attached to a duplication of the only remaining copy of chr17. There is an inverted duplication of 25 kbp of chr15 starting 40 kbp from the telomere. Interestingly 210 bp from the middle of chr11 is duplicated in between the copies of the inverted duplication. This translocation is only clear using matched normal and tumor assemblies because acrocentric short arms frequently exchange material between chromosomes. Furthermore, the HG008 normal short arms of chr15 haplotype 1 and chr14 haplotype 2 are nearly identical in the first 4.1 Mbps from the telomere through the rDNA arrays, with only one SNV and one indel difference in this region outside the rDNA arrays. This region includes the rDNA arrays, though these arrays were not fully resolved in the assemblies. The tumor assembly contains 11 somatic SNVs on chr15 and 12 somatic SNVs on chr14 in this region supported by HiFi reads, illustrating the unusual situation of far fewer germline SNVs than somatic SNVs in this region. Karyoscope's chromosome, subtelomere, repeat, gene, satellite annotations are shown across the entire chromosomes and at different resolutions around each interchromosomal breakpoint. Note that inversions are not shown by karyoscope.

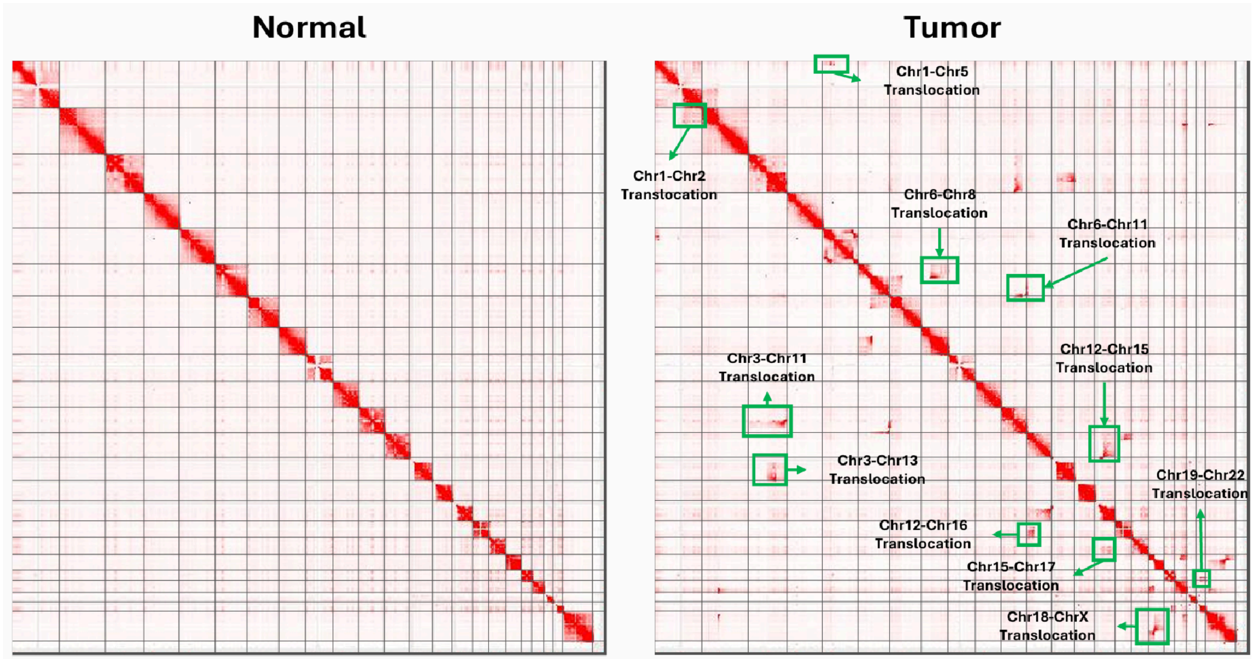


**Supplementary Fig. 10: Hi-C Contact maps for Tumor and Normal on GRCh38.** We highlight the impact of a few large interchromosomal translocations and inversions evident when comparing the Hi-C contact map aligned onto the GRCh38 reference assembly of the tumor sample against its matched normal tissue sample.

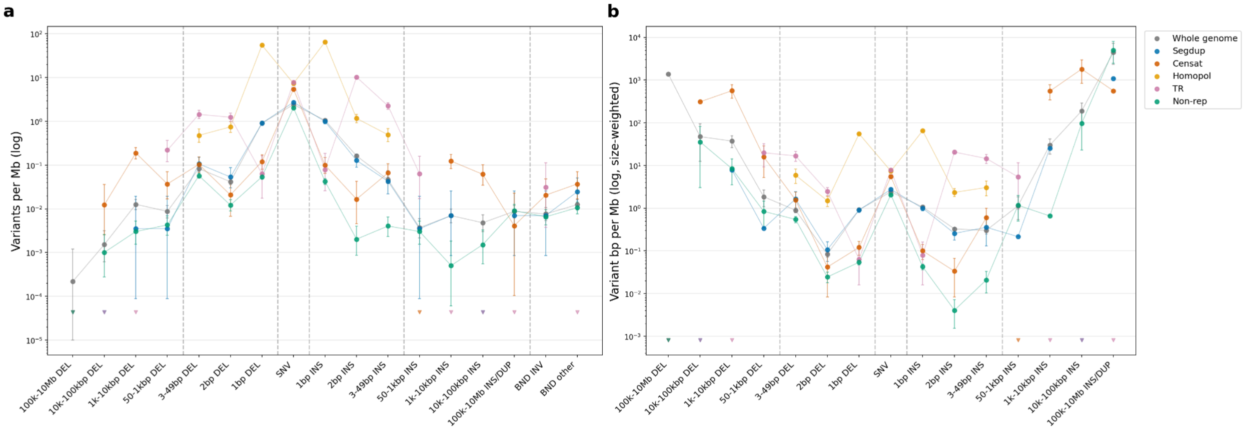


**Supplementary Fig. 11: Small variant and SVs are generally enriched in repetitive regions.** (a) Count-based variant rates per Mbp in each region. (b) Size-weighted variant rates per Mbp in each region. There are more 1-2 bp insertions than deletions, but more 3 bp to 10 kbp deletions than insertions, and more insertions than deletions larger than 10 kbp. Comparing repeat unit size, repeat length, and variant type we observe the following trends: 1 bp and 2 bp indels predominate in homopolymers and dinucleotide TRs; larger insertions and deletions predominate in STRs, VNTRs, and satellites; 10 kbp to 100 kbp insertions/duplication predominate in satellites; and tandem duplications > 100 kbp predominate in non-repetitive regions. Large homopolymers exhibit interesting characteristics including: 14 1bp indels in >50bp homopolymers, 7 contractions of 10-30bp in homopolymers longer than 50bp in our benchmark, and 14 deletions >=10bp in homopolymers <50bp. Unlike the 21 contractions of at least 10 bp in homopolymers, which appeared true upon curation, the 5 putative expansions of at least 10bp in homopolymers appeared likely to be assembly errors, mostly haplotype switches in the normal assembly. We observed 607 2bp ins and 80 2bp del in diTRs with 232 SNVs. Similar to previous analysis of mutation rates,[102] there are many more deletions (14) than insertions (2) larger than 9bp. Somatic mutation rates are lower for diTRs than A/T or G/C homopolymers of similar lengths and type, which is opposite what was recently found when merging all lengths and types for a smaller number of germline de novo mutations with similar accurate short reads.[19] However, it is similar to a previous analysis of germline variants a large number of number of samples comparing rates at different lengths of homopolymers and diTRs (Extended Data Fig 2 in [103]). Adjacent TRs and homopolymers show distinct patterns. For variants occurring in between two homopolymers each longer than 6bp, there are 65 SNVs, 108 1bp INS, and 84 1bp DEL. Interestingly, within adjacent homopolymers there are 39 5′−TTA−3′ ↔ 5′−TAA−3’ vs 13 5′−TAA−3′ ↔ 5′−TTA−3, similar to asymmetry of spontaneous mutations in *C. elegans*.[104] For adjacent diTRs, SNVs are enriched with 256 SNVs, 67 2bp INS, and 3 2bp DEL.

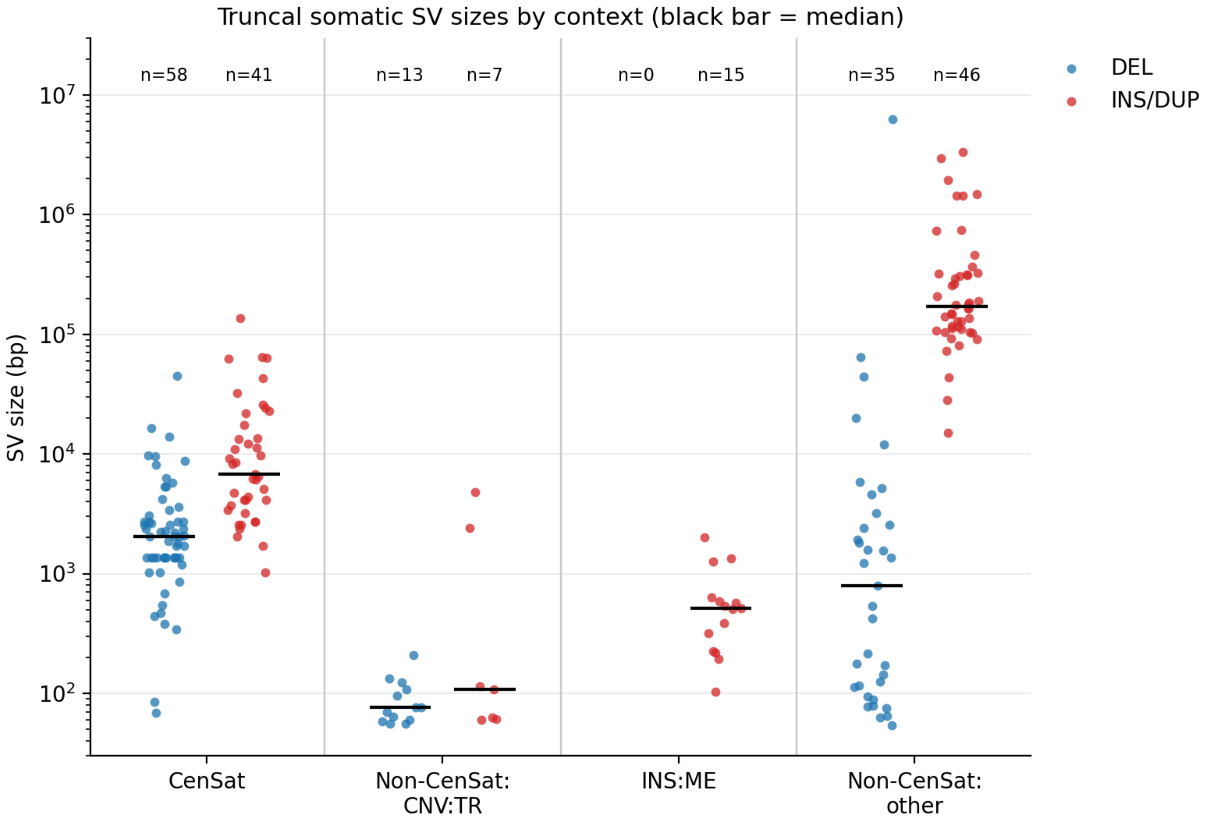


**Supplementary Fig. 12: SV size distributions by repeat SV type.** While germline variants typically have balanced insertion and deletions with respect to an accurate reference like T2T-CHM13, our tumor/normal assemblies reveal HG008's somatic SVs have distinct deletion and insertion profiles in different types of genomic repeats.

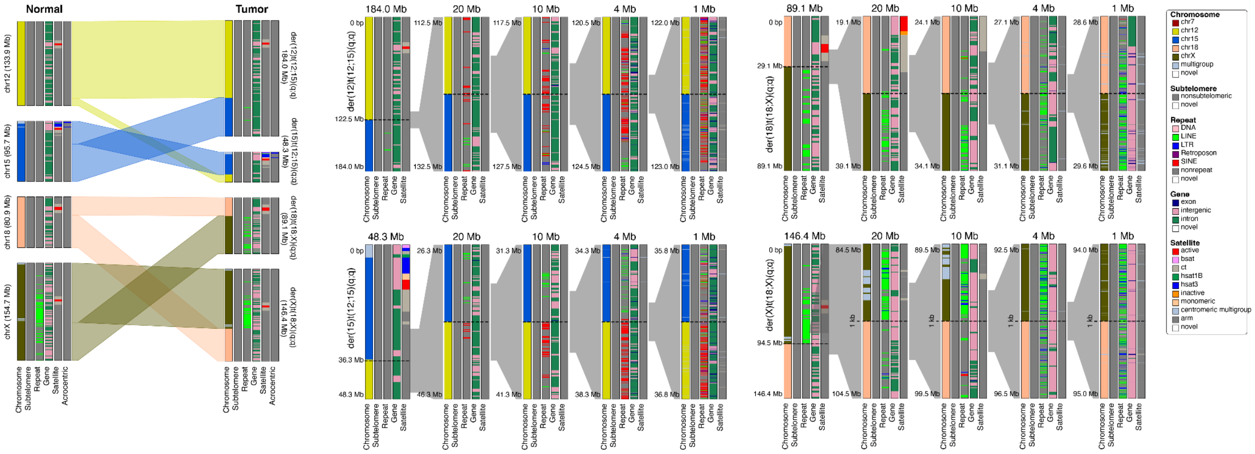


**Supplementary Fig. 13: Reciprocal translocations between chr12 and chr15 and chr18 and chrX on haplotype 2.** These translocations do not occur in the centromeres and so retain a single copy of a centromere in the resulting tumor chromosomes. Karyoscope's chromosome, subtelomere, repeat, gene, satellite annotations are shown across the entire chromosomes and at different resolutions around each interchromosomal breakpoint. Note that inversions, including the one on chr18, are not shown by karyoscope.

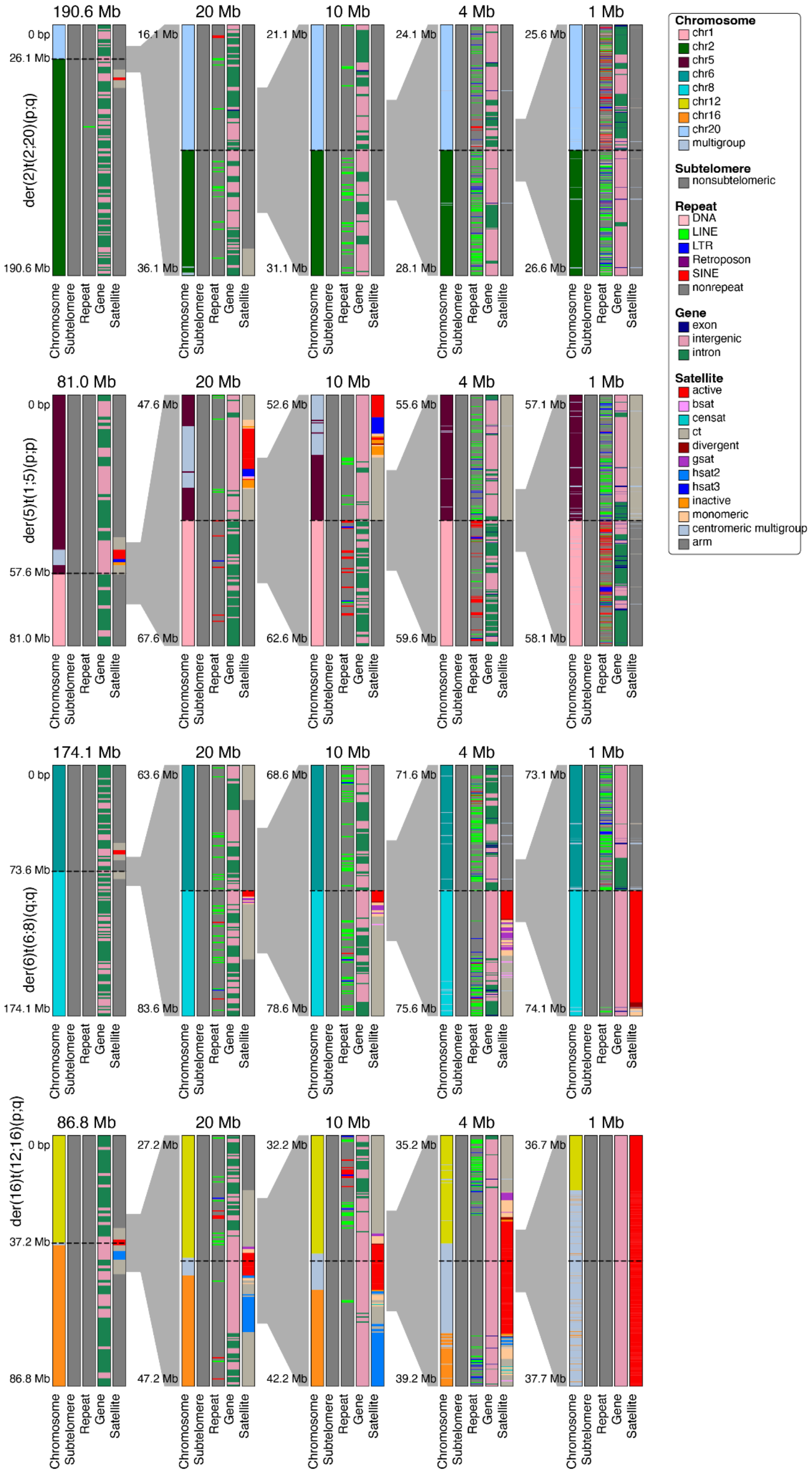

der(2)t(2;20)(p;q)
der(5)t(1;5)(p;p)
der(6)t(6;8)(q;q)
der(16)t(12;16)(p;q)
190.6 Mb
20 Mb
10 Mb
4 Mb
1 Mb
81.0 Mb
174.1 Mb
86.8 Mb
0 bp
26.1 Mb
190.6 Mb
16.1 Mb
36.1 Mb
21.1 Mb
31.1 Mb
24.1 Mb
28.1 Mb
25.6 Mb
26.6 Mb
57.6 Mb
81.0 Mb
47.6 Mb
67.6 Mb
52.6 Mb
62.6 Mb
55.6 Mb
59.6 Mb
57.1 Mb
58.1 Mb
73.6 Mb
174.1 Mb
63.6 Mb
83.6 Mb
68.6 Mb
78.6 Mb
71.6 Mb
75.6 Mb
73.1 Mb
74.1 Mb
37.2 Mb
86.8 Mb
27.2 Mb
47.2 Mb
32.2 Mb
42.2 Mb
35.2 Mb
39.2 Mb
36.7 Mb
37.7 Mb
Chromosome
Subtelomere
Repeat
Gene
Satellite
Chromosome
chr1
chr2
chr5
chr6
chr8
chr12
chr16
chr20
multigroup
Subtelomere
nonsubtelomeric
Repeat
DNA
LINE
LTR
Retroposon
SINE
nonrepeat
Gene
exon
intergenic
intron
Satellite
active
bsat
censat
ct
divergent
gsat
hsat2
hsat3
inactive
monomeric
centromeric multigroup
arm

**Supplementary Fig. 14: Karyoscope's annotations for non-reciprocal translocations. Translocations between chr1 and chr5 and between chr2 and chr20 cause duplications of parts of chr1p and chr20q. These translocations do not occur in the centromeres and retain a single copy of a centromere in the resulting tumor chromosomes originating from the non-duplicated chromosome. Translocations between chr6 and chr8 and between chr12 and chr16 occur in one or both centromeres.**

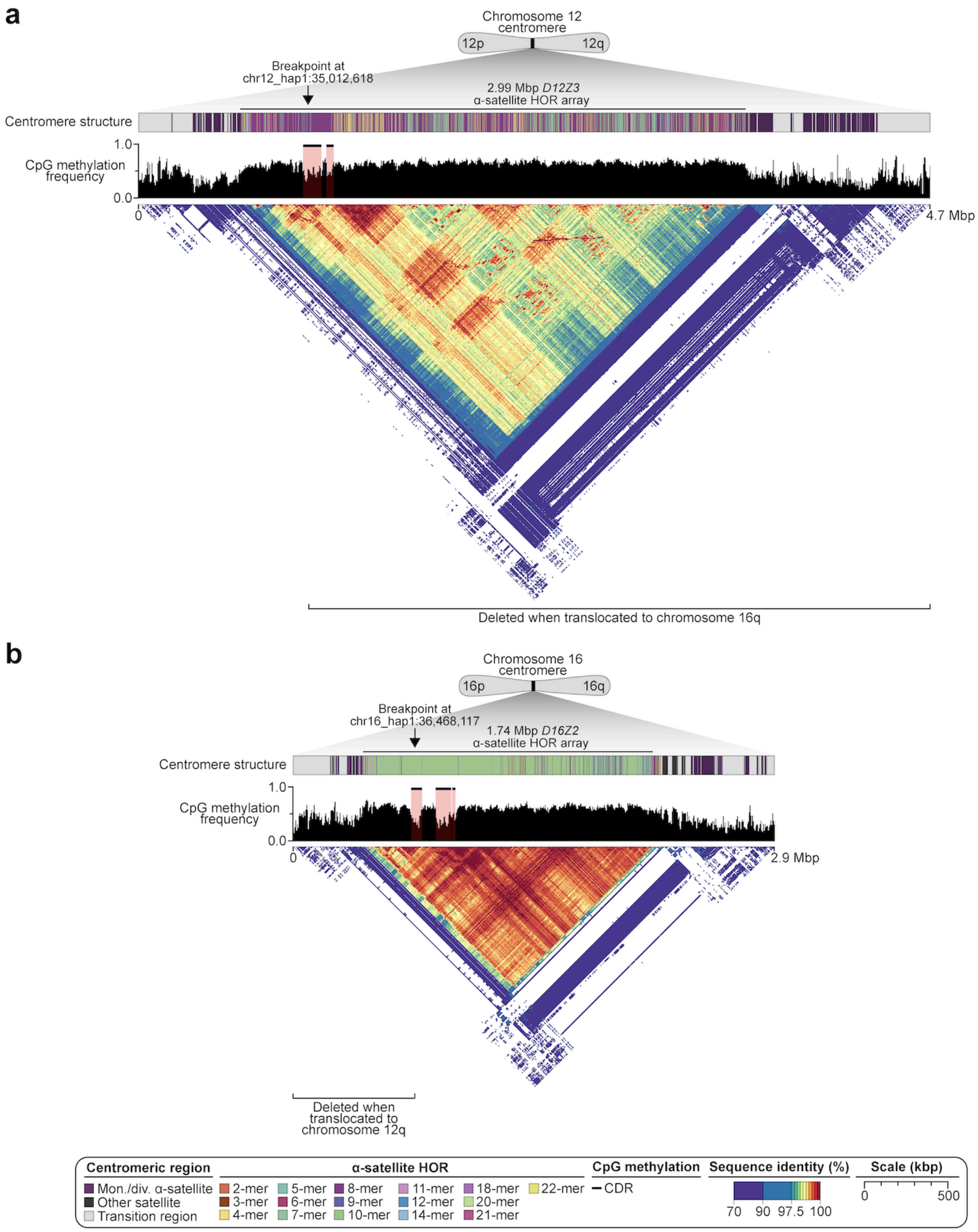
a
Chromosome 12 centromere
12p
12q
Breakpoint at chr12_hap1:35,012,618
2.99 Mbp D12Z3 α-satellite HOR array
Centromere structure
CpG methylation frequency
1.0
0.0
0
4.7 Mbp
Deleted when translocated to chromosome 16q
b
Chromosome 16 centromere
16p
16q
Breakpoint at chr16_hap1:36,468,117
1.74 Mbp D16Z2 α-satellite HOR array
Centromere structure
CpG methylation frequency
1.0
0.0
0
2.9 Mbp
Deleted when translocated to chromosome 12q
Centromeric region
Mon./div. α-satellite
Other satellite
Transition region
α-satellite HOR
2-mer
3-mer
4-mer
5-mer
6-mer
7-mer
8-mer
9-mer
10-mer
11-mer
12-mer
14-mer
18-mer
20-mer
21-mer
22-mer
CpG methylation
CDR
Sequence identity (%)
70
90
97.5
100
Scale (kbp)
0
500

**Supplementary Fig. 15. The sequence, structure, and epigenetic landscape of the chromosome 12 and 16 centromeres in normal cells.** For both chromosomes a) 12 and b) 16, a portion of the CDR sites is retained upon translocation, while the rest of the CDR site and most of the α-satellite HOR array is deleted. This creates a fused chromosome with a hybrid CDR site residing on two α-satellite HOR arrays (*D12Z3* and *D16Z2*; Fig. 4b).

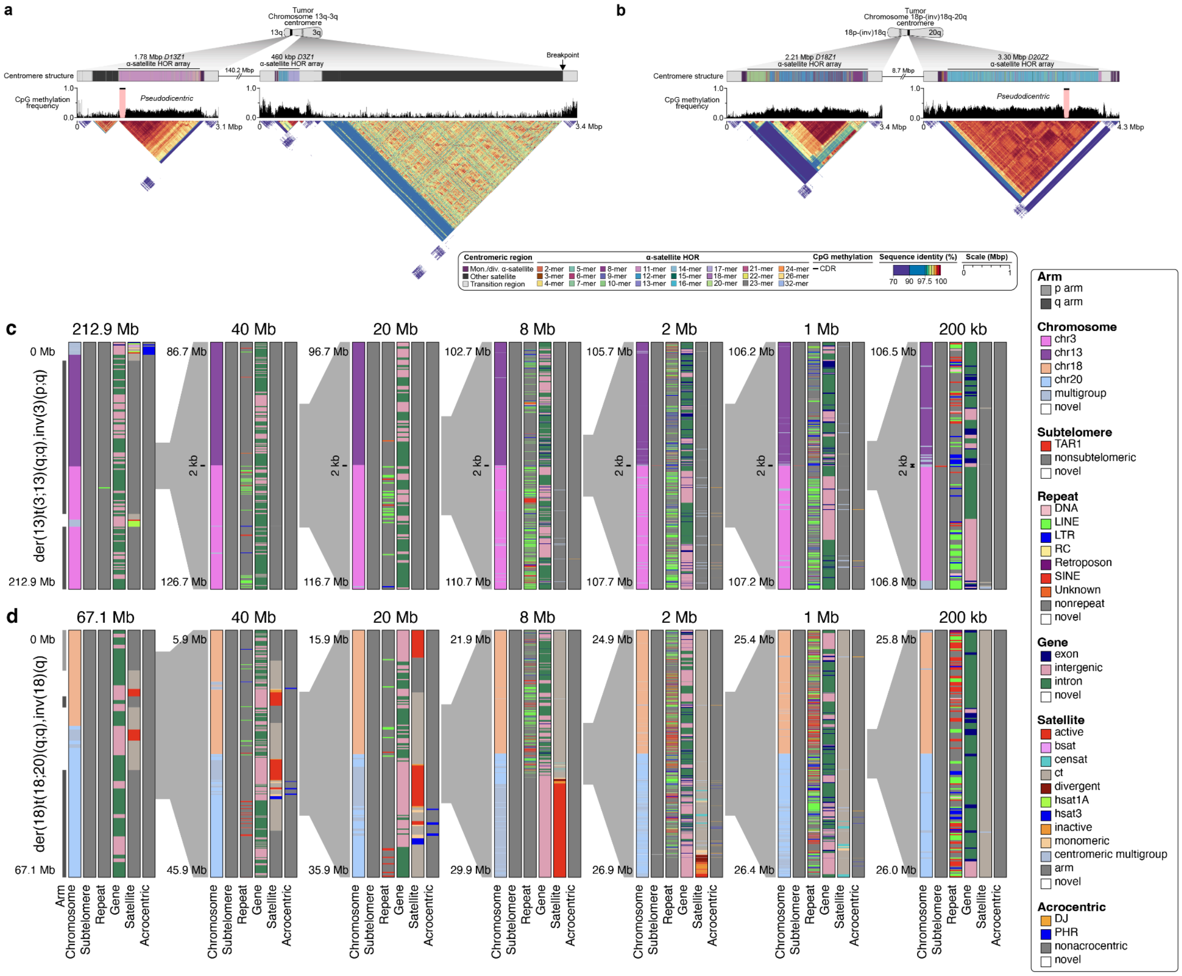

**Supplementary Fig. 16. The sequence, structure, and epigenetic landscape of the chromosome 13q-3q and 18p-(inv)18q-20q centromeres in the tumor.** a) Upon translocation of chromosome 13q to 3q, the chromosome 3 centromere is inactivated by the complex inversions in Supplementary Fig 8 that delete the CDR, while the chromosome 13 centromere retains the CDR. b) Upon translocation of 18p-(inv)18q to 20q, the chromosome 18 CDR is inactivated by methylation though the inversion in Supplementary Fig 8 also deletes part of the centromere, while the chromosome 20 centromere retains its function. c-d) Karyoscope's chromosome, subtelomere, repeat, gene, satellite annotations are shown across the entire chromosomes and at different resolutions around each interchromosomal breakpoint.

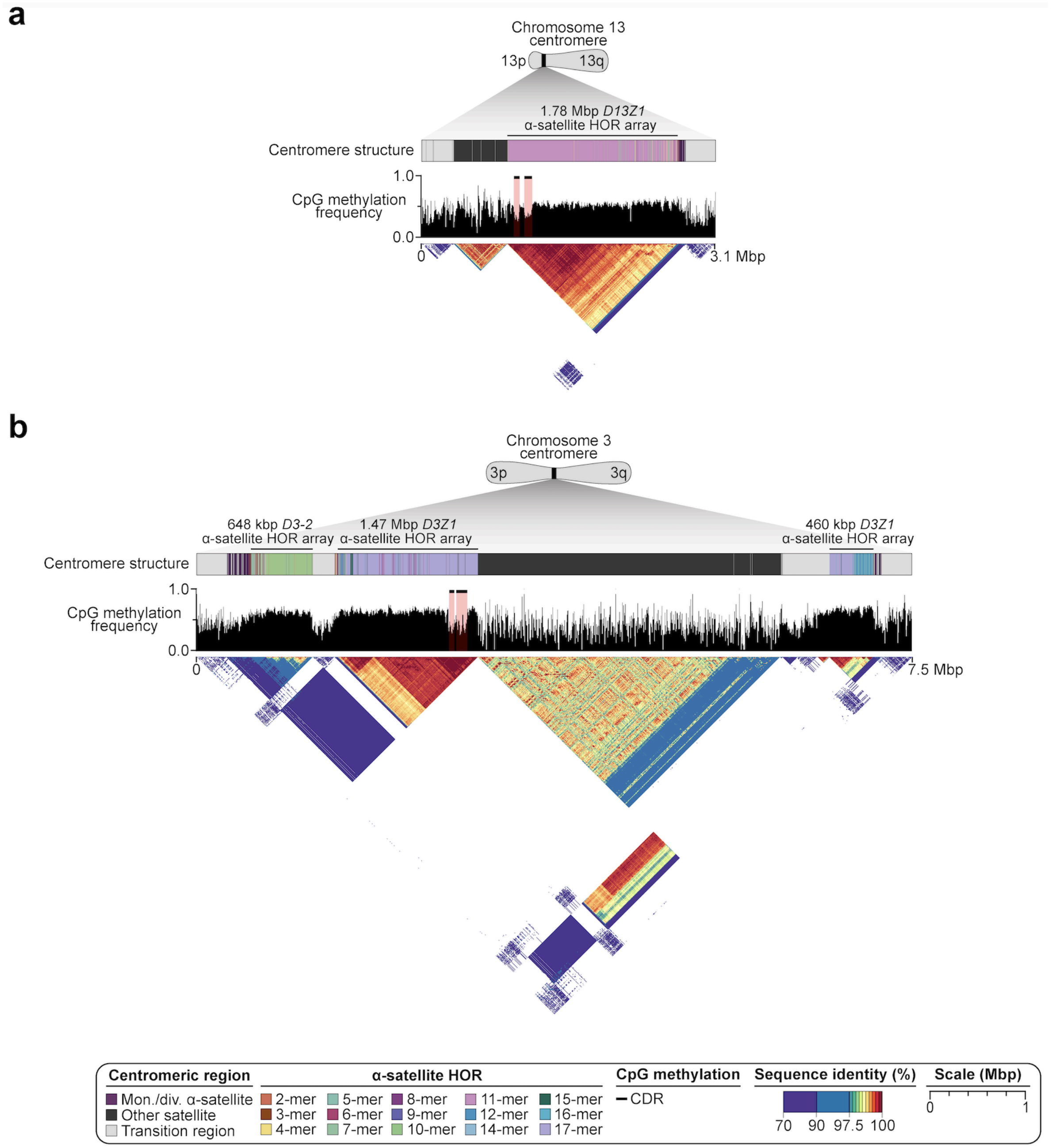


**Supplementary Fig. 17. The sequence, structure, and epigenetic landscape of the chromosome 13 and 3 centromeres in normal cells.** a) For chromosome 13, the CDR site is retained upon translocation. b) For chromosome 3, however, the CDR-containing centromeric α-satellite HOR array is deleted, and the remaining sequence is inverted upon translocation.

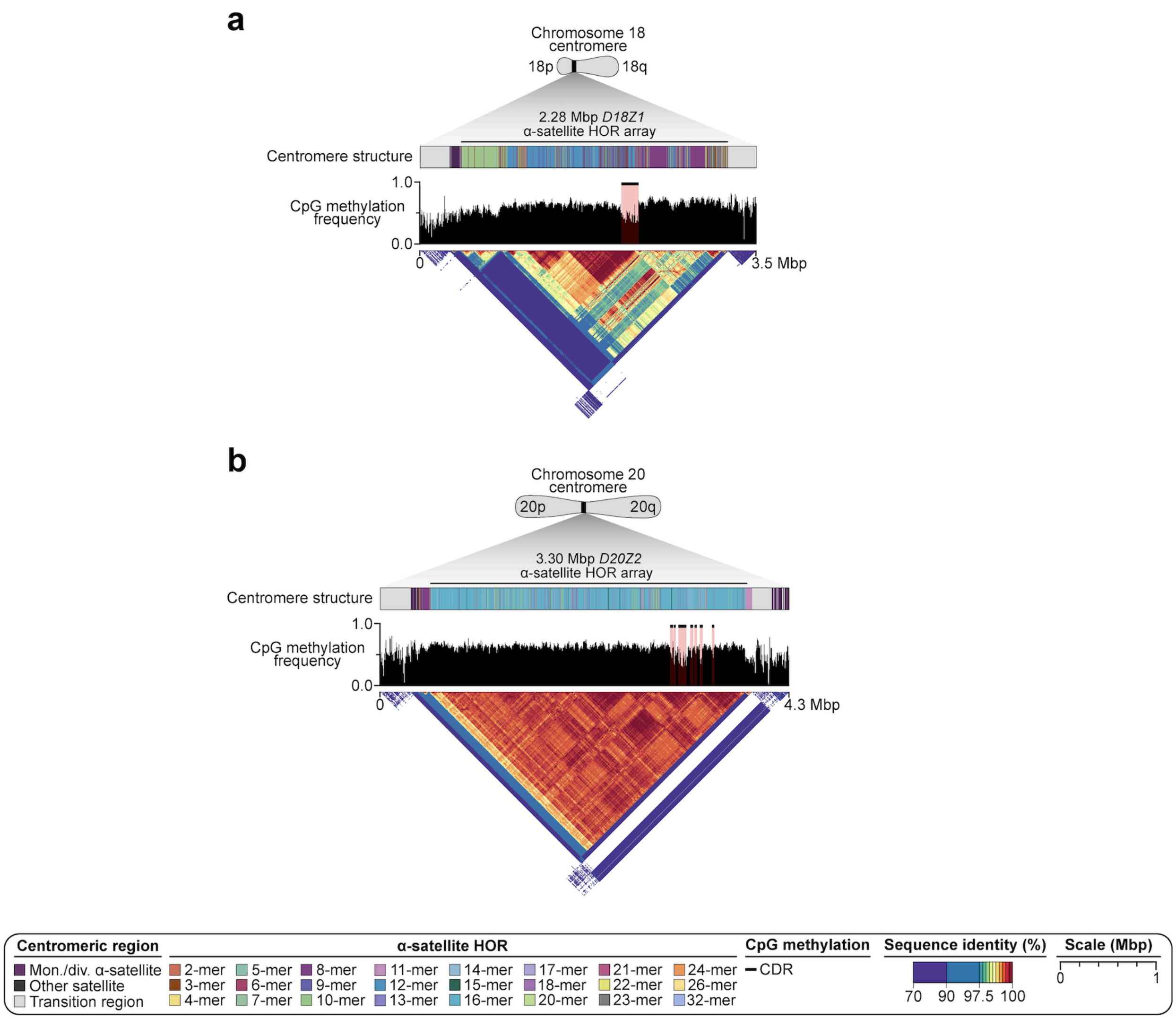


**Supplementary Fig. 18. The sequence, structure, and epigenetic landscape of the chromosome 18 and 20 centromeres in normal cells.** a) For chromosome 18, the CDR site becomes methylated during tumorigenesis, resulting in inactivation of the centromere, though part of the centromere is also deleted by a large inversion related to the translocation. b) For chromosome 20, however, the CDR position is retained but changes in methylation pattern during tumorigenesis.

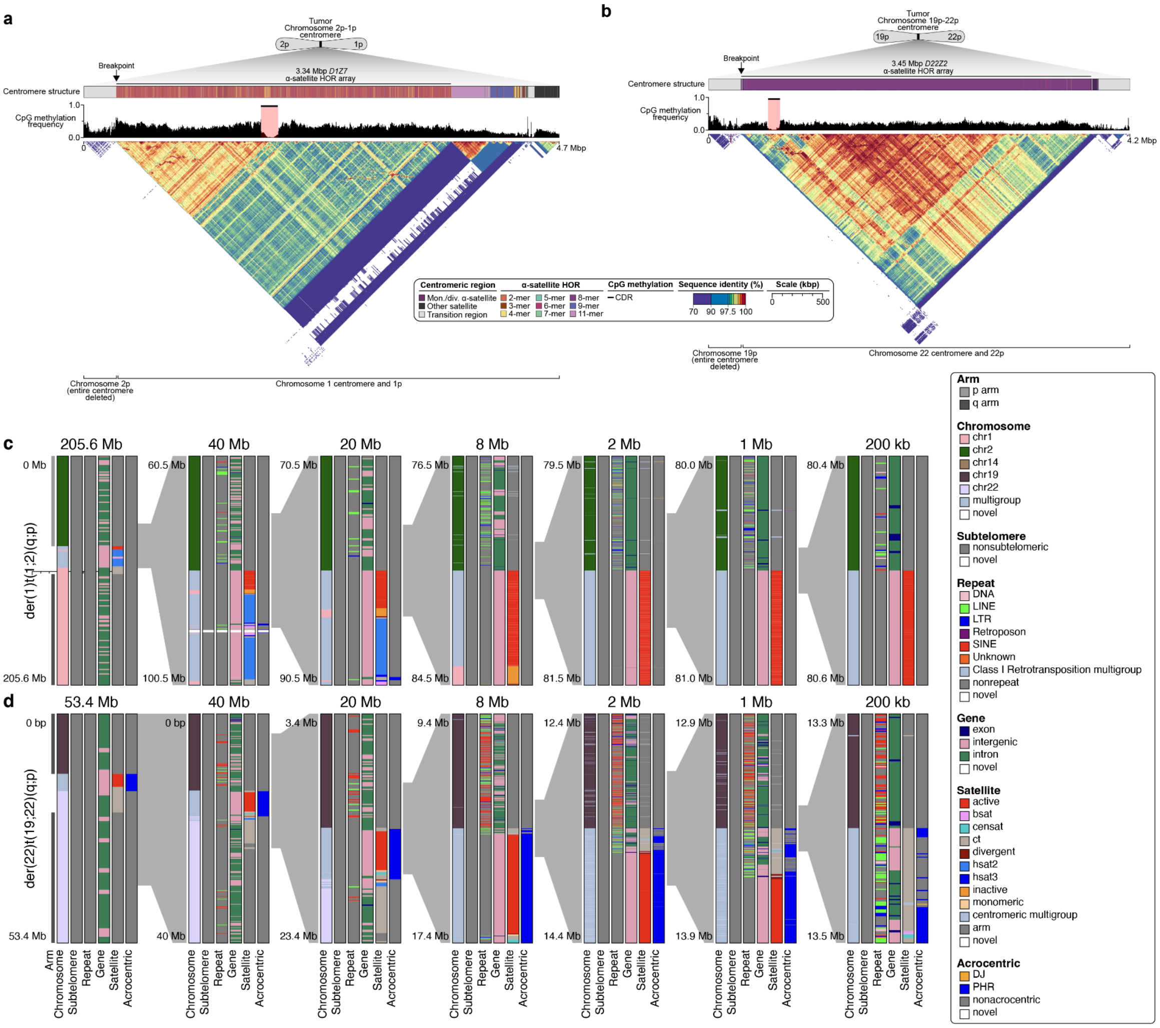


**Supplementary Fig. 19. The sequence, structure, and epigenetic landscape of the chromosome 2p-1p and 19p-22p centromeres in the tumor.** a) Upon translocation of chromosome 2p to 1p, the chromosome 2 centromere is deleted while the chromosome 1 centromere is retained, albeit truncated on the p arm-proximal side. b) Similarly, upon translocation of 19p to 22p, the chromosome 19 centromere is deleted, while the chromosome 22 centromere is retained, also with a truncation on the p arm-proximal side. c) Karyoscope's chromosome, subtelomere, repeat, gene, satellite, and acrocentric annotations are shown across the entire chromosomes and at different resolutions around each interchromosomal breakpoint.

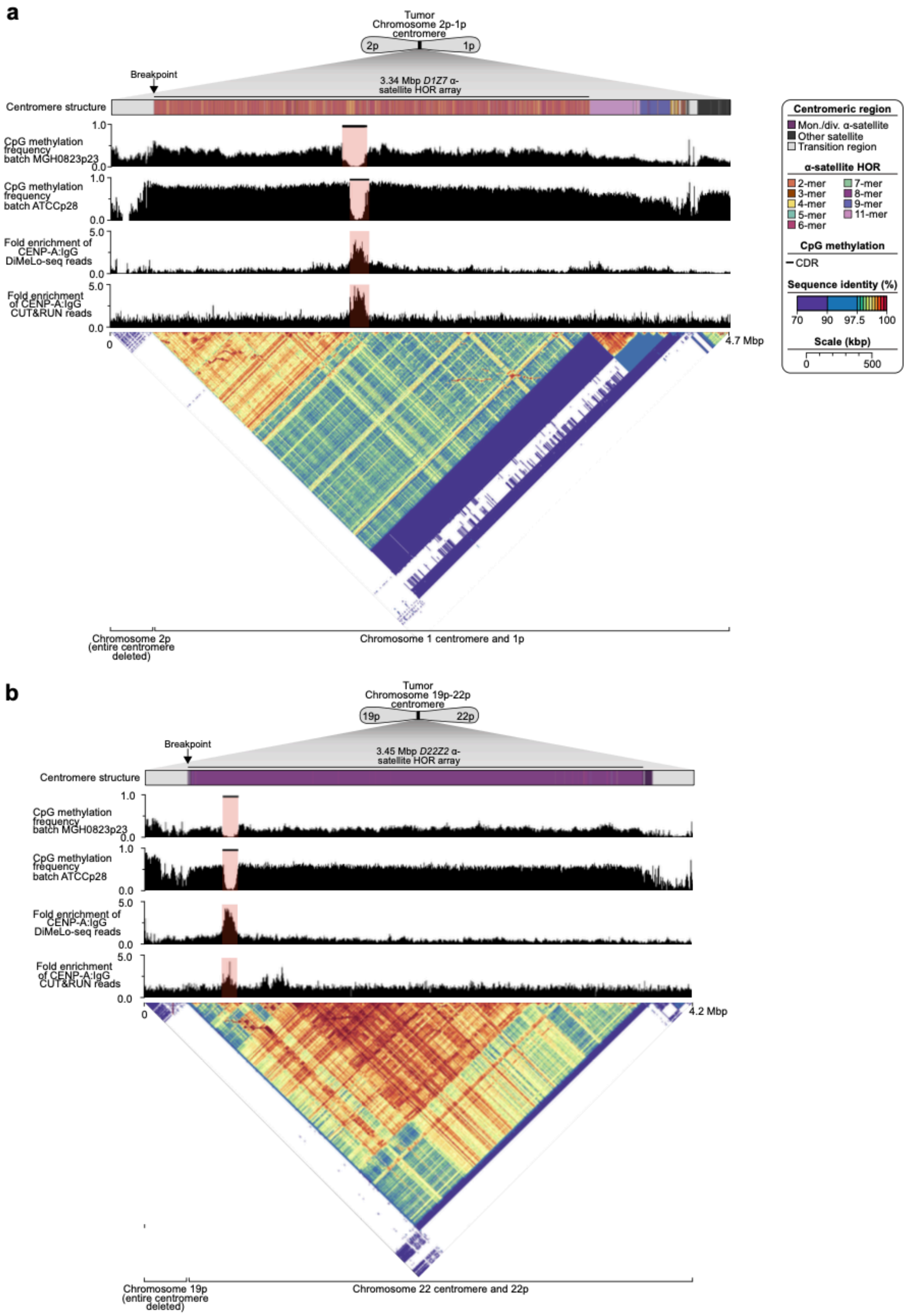
a
Tumor
Chromosome 2p-1p centromere
2p
1p
Breakpoint
3.34 Mbp D1Z7 α-satellite HOR array
Centromere structure
CpG methylation frequency batch MGH0823p23
CpG methylation frequency batch ATCCp28
Fold enrichment of CENP-A:IgG DiMeLo-seq reads
Fold enrichment of CENP-A:IgG CUT&RUN reads
1.0
0.0
5.0
0
4.7 Mbp
Chromosome 2p (entire centromere deleted)
Chromosome 1 centromere and 1p
Centromeric region
Mon./div. α-satellite
Other satellite
Transition region
α-satellite HOR
2-mer
3-mer
4-mer
5-mer
6-mer
7-mer
8-mer
9-mer
11-mer
CpG methylation
CDR
Sequence identity (%)
70
90
97.5
100
Scale (kbp)
0
500
b
Tumor
Chromosome 19p-22p centromere
19p
22p
Breakpoint
3.45 Mbp D22Z2 α-satellite HOR array
Centromere structure
CpG methylation frequency batch MGH0823p23
CpG methylation frequency batch ATCCp28
Fold enrichment of CENP-A:IgG DiMeLo-seq reads
Fold enrichment of CENP-A:IgG CUT&RUN reads
4.2 Mbp
Chromosome 19p (entire centromere deleted)
Chromosome 22 centromere and 22p

**Supplementary Fig. 20. The sequence, structure, and epigenetic landscape of the chromosome 2p-1p and 19p-22p centromeres in the tumor.** a) Upon translocation of chromosome 2p to 1p, the chromosome 2 centromere is deleted while the chromosome 1 centromere is retained with CENP-A binding, albeit truncated on the p arm-proximal side. b) Similarly, upon translocation of 19p to 22p, the chromosome 19 centromere is deleted, while the chromosome 22 centromere is retained with CENP-A binding, also with a small truncation on the p arm-proximal side.

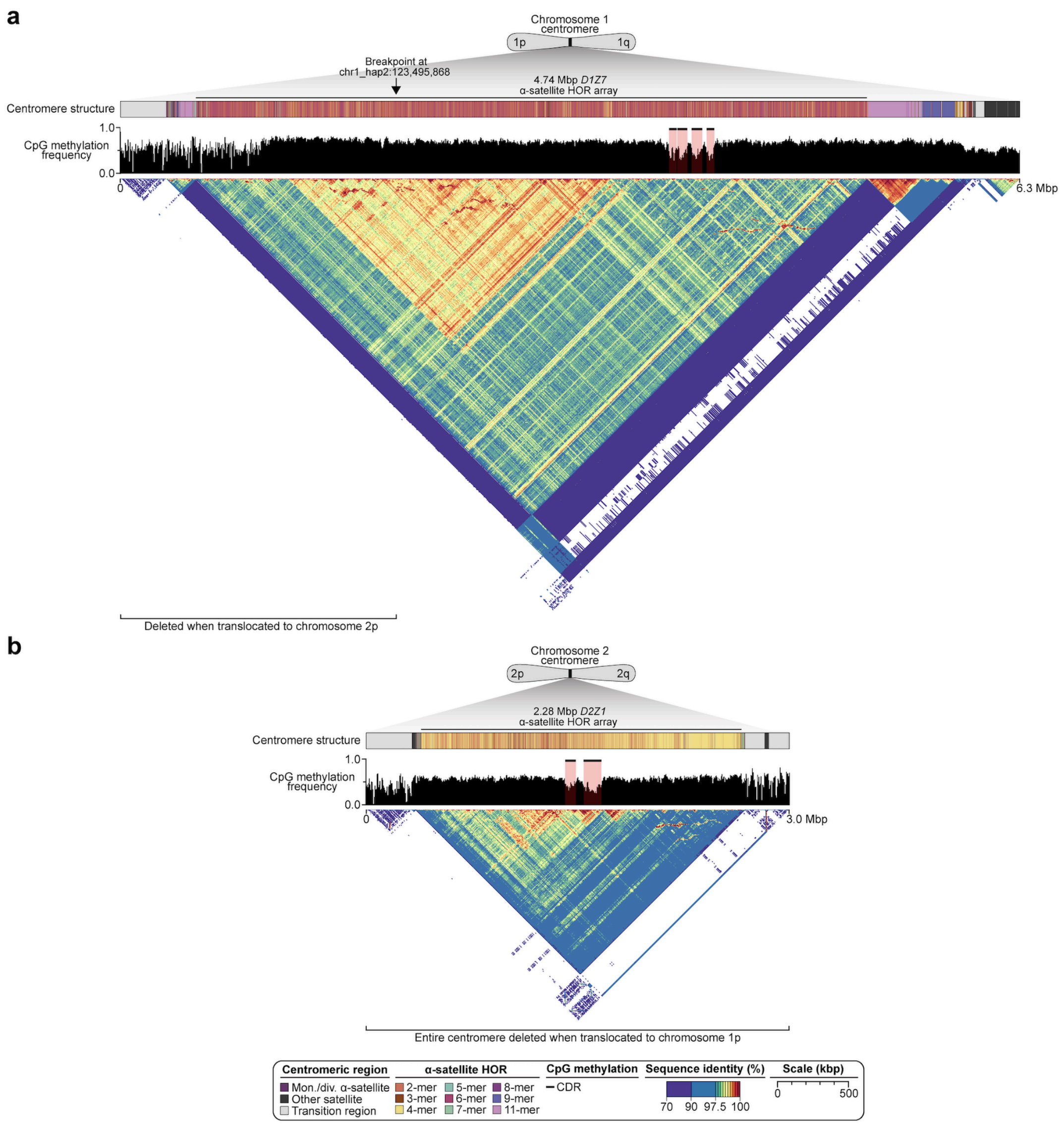


**Supplementary Fig. 21. The sequence, structure, and epigenetic landscape of the chromosome 2 and 1 centromeres in normal cells.** a) For chromosome 1, the CDR site is retained upon translocation. b) For chromosome 2, however, the entire centromeric α-satellite HOR array, including the CDR, is deleted upon translocation.

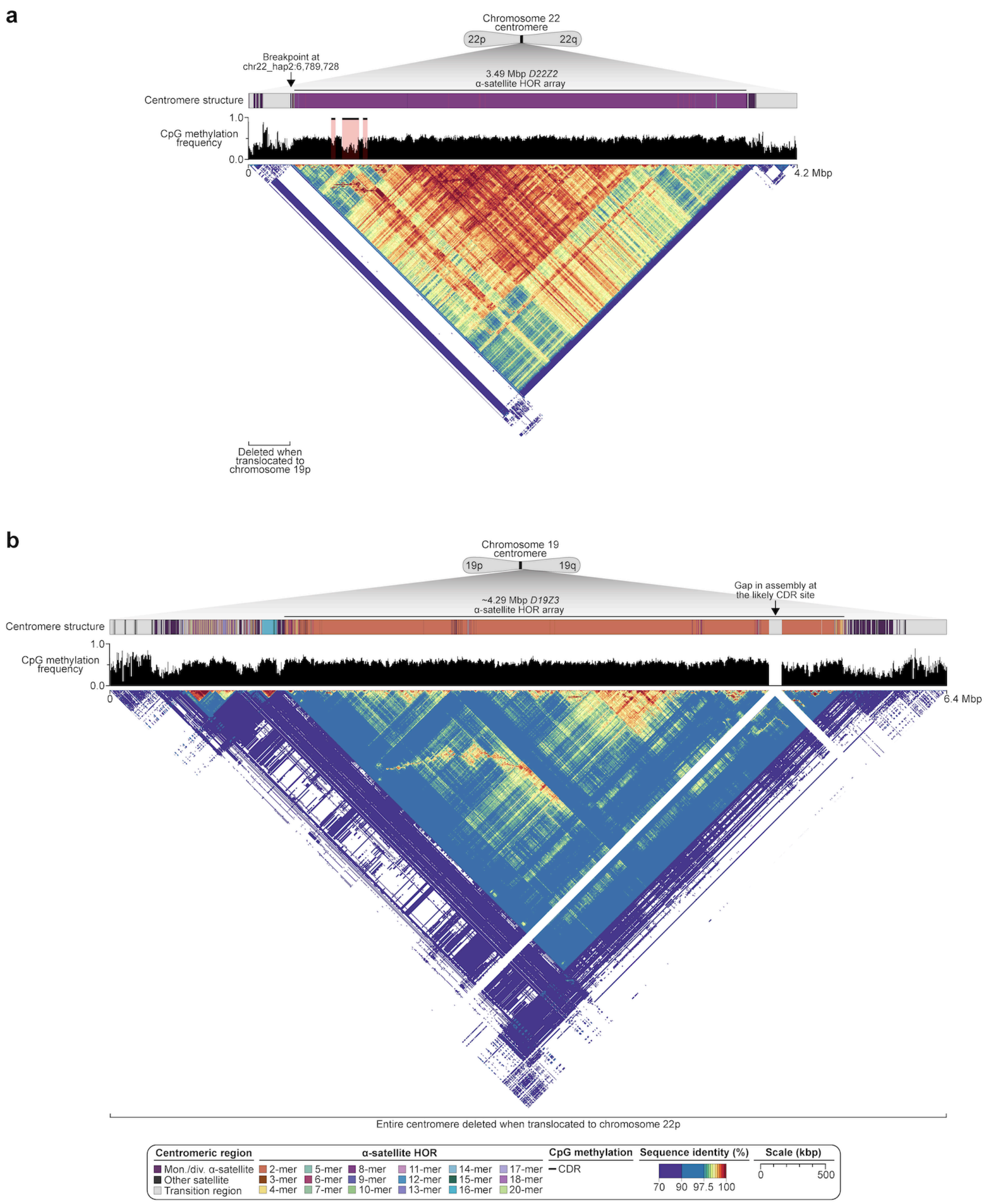


**Supplementary Fig. 22. The sequence, structure, and epigenetic landscape of the chromosome 22 and 19 centromeres in normal cells.** a) For chromosome 22, the CDR site is retained upon translocation. b) For chromosome 19, however, the entire centromeric α-satellite HOR array, including the CDR, is deleted upon translocation. The location of the CDR site likely resides in a gap in the assembly.

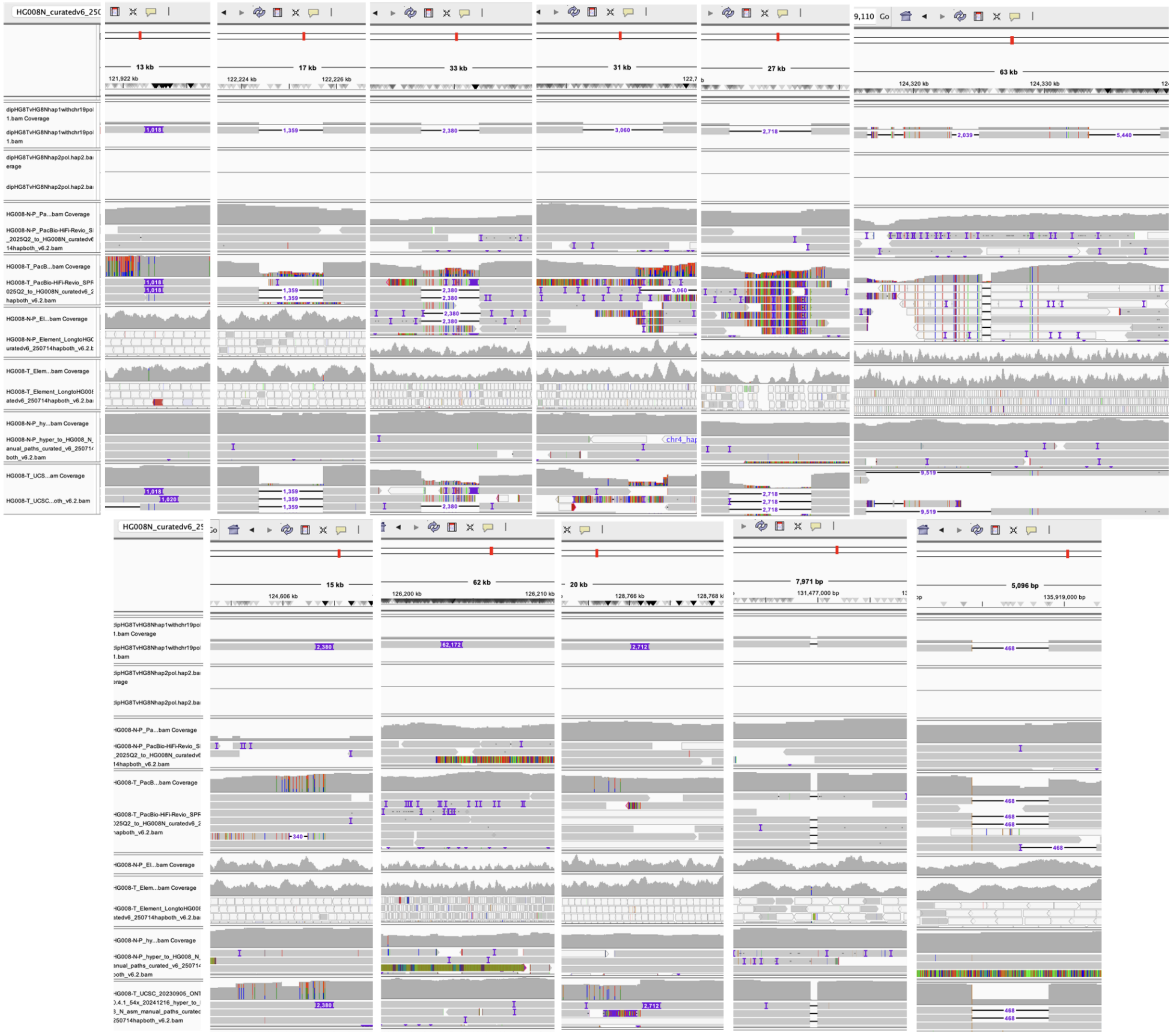

**Supplementary Fig. 23: Complete chromosome 1 centromere comparison reveals four times more somatic SVs than in entire GRCh38 chromosome 1.** While only three truncal SVs were identified on chromosome 1 haplotype 1 in GRCh38, the tumor vs normal assembly alignment of these haplotypes identified 12 additional likely truncal SVs in the CenSat region (as well as one in a small non-centromeric HSat3 array and one in the 5S rDNA array). We examined variants in the ~17 Mbp CenSat region of chromosome 1 haplotype 1, which is complete in both the tumor and normal. In the α-satellite HOR array, there are 4 insertions (1,018, 2,380, 12,240, and 62,172 bp in length) and 6 deletions (1,359, 2,039, 2,380, 2,718, 3,060, and 9,519 bp in length). There's also a 69 bp deletion of one β-satellite monomer, and a 2,712 bp insertion and 468 and 1,776 bp deletions in the HSat2 array. Four of the SVs have phased SNVs near the breakpoints, with many deletions having a SNV at the base immediately before the start of the deletion. In addition, there are 66 confirmed SNVs and 5 confirmed 1 bp indels in this ~17 Mbp CenSat region though some true variants may not have been confirmed due to mapping challenges. See support for SVs in https://github.com/jzook/HG008SVcuration/issues/327-338. A CenSat subclonal DEL is in https://github.com/jzook/HG008SVcuration/issues/340. While chr1 has a particularly large CenSat region with 12 SVs, this CenSat region is highly enriched for somatic SVs, while having similar SNV rate and lower indel rate than the rest of the chromosome. While acrocentric rDNA arrays are not resolved in the assemblies, the ~300 kbp 5S rDNA array on chr1 haplotype 1 (but not haplotype 2) is resolved in the tumor and normal assemblies, and it has a truncal somatic 2,232 bp deletion of one monomer unit (which includes a ~120 bp 5S rRNA gene and a 2.1 kbp intergenic spacer). The region near the middle of the array with the deletion is methylated in both tumor and normal, so it probably is inactive and unlikely to have an effect. Interestingly, the normal assembly enables us to see that while most of the array is nearly fully methylated, the ~40 kbp on the right side of the array is unmethylated, similarly in both tumor and normal https://github.com/jzook/HG008SVcuration/issues/339.

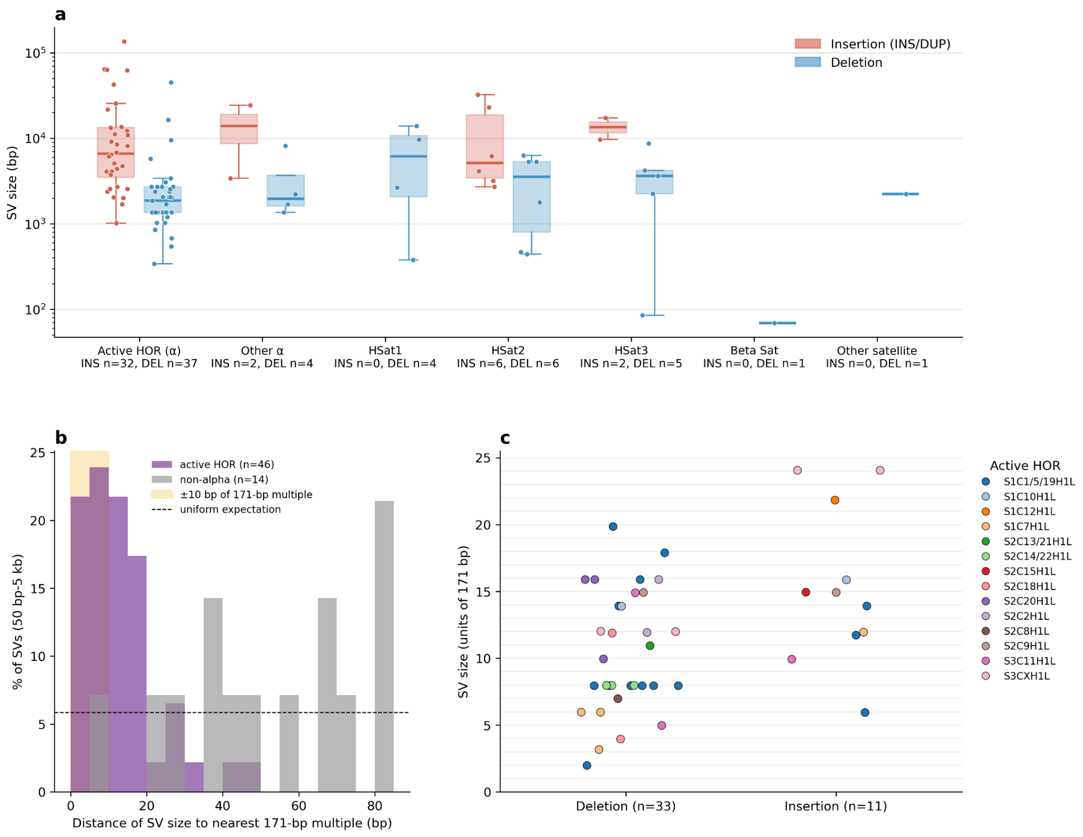


**Supplementary Fig. 24: Centromere satellite SVs vary by repeat type and follow expected alpha satellite periodicity.** (a) Analysis of 100 somatic insertions and deletions in centromeric and satellite DNA regions of HG008-T revealed two distinct patterns. First, somatic insertions were significantly larger than deletions pooled across all satellite types (median: 6,636 bp vs 2,046 bp; Mann-Whitney U test, $p = 3.1\times10^{-8}$). The four largest confirmed somatic expansions of 62 kbp, 63, kbp, 65 kbp, and 136 kbp are larger than germline or somatic de novo centromeric expansions seen in recent studies. This pattern may suggest that large insertions in HG008-T arise through tandem duplication mechanisms similar to those observed in non-repetitive regions. However, these are also challenging to detect without assembling both tumor and normal reads so may have been invisible to mapping-based approaches used in previous somatic studies. (b) Smaller somatic SVs (50 bp to 5 kb) in active higher-order repeat (HOR) α-satellites showed strong periodicity at 171 bp intervals, with 54.3% of variants falling within ±10 bp of an exact 171 bp multiple—a 4.4-fold enrichment over the 12.3% expected by chance ($\chi^2 = 75.6$, $p = 3.5\times10^{-18}$). This periodicity was specific to α-satellites, as non-α-satellite variants showed no such pattern (11.1% near 171 bp multiples), consistent with replication slippage or unequal sister chromatid exchange involving whole α-satellite monomers (171 bp) or their higher-order multiples. (c) 33 deletions and 11 insertions in active HOR are 25 units or smaller. The other 25 active-HOR SVs are larger and not shown. These findings demonstrate that somatic structural variation in satellite DNA follows mechanistic constraints imposed by the underlying repeat structure.

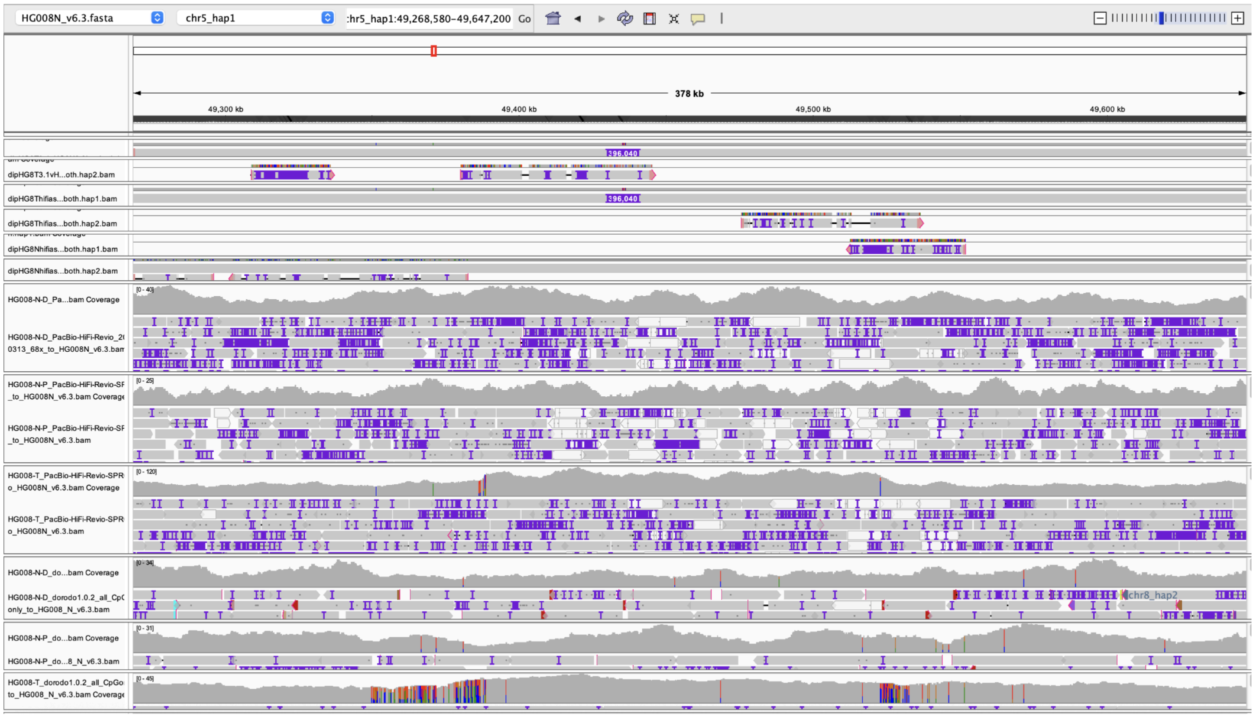


**Supplementary Fig. 25: Large CenSat insertions**. The largest CenSat insertion is a 136 kbp tandem duplication on chr5_hap1 (https://github.com/jzook/HG008SVcuration/issues/351). The hifiasm assemblies match the v3.1 tumor and v6.3 normal assemblies, and it is also supported by the ~2 times higher coverage of the 136 kbp region in the middle, and clipped ONT reads in the tumor and not normal. A 375 kbp ONT read traverses one copy of the duplicated region and aligns ~60 kbp into the second copy since it noisily aligns ~60 kbp past the left breakpoint. Other large CenSat insertions >30 kbp include https://github.com/jzook/HG008SVcuration/issues/445, https://github.com/jzook/HG008SVcuration/issues/396, https://github.com/jzook/HG008SVcuration/issues/381, https://github.com/jzook/HG008SVcuration/issues/379, and https://github.com/jzook/HG008SVcuration/issues/334.

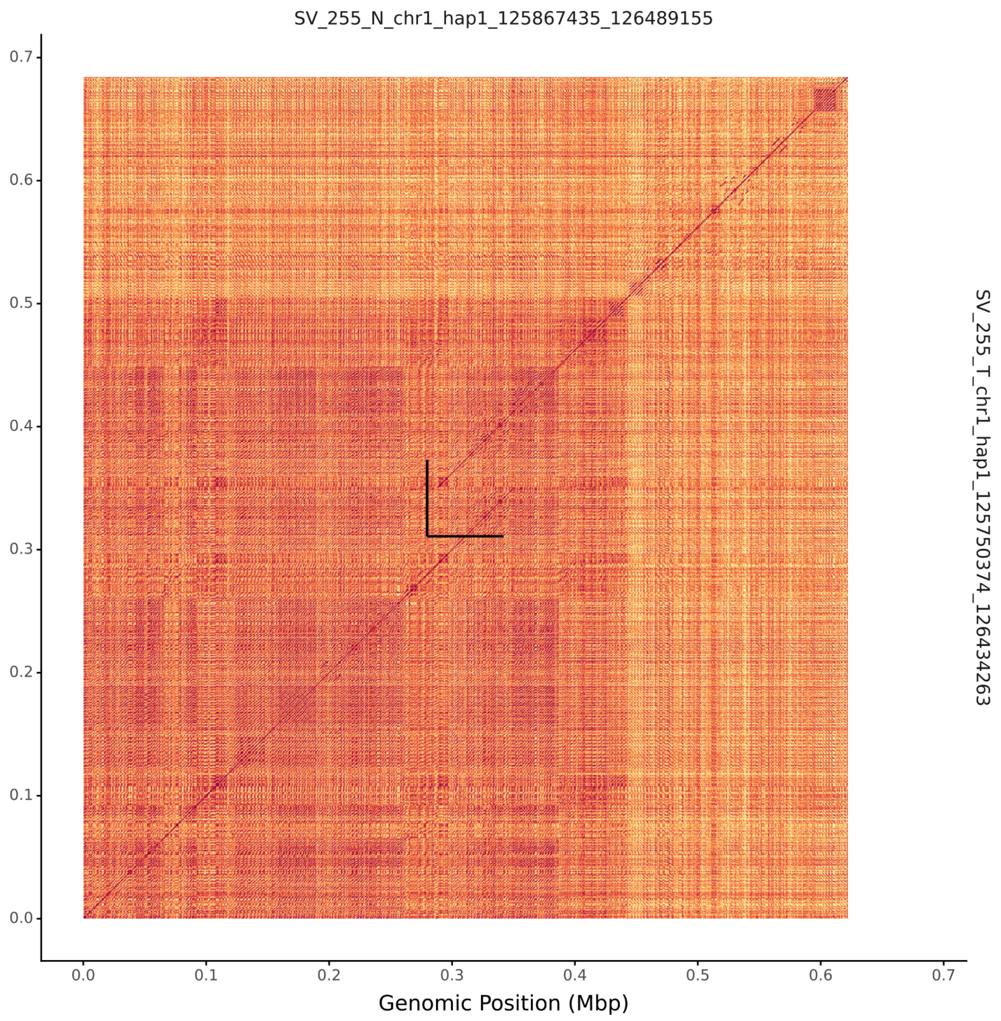


**Supplementary Fig. 26: ModDotPlot showing a 62,172 bp tandem duplication in the chr1 active α-satellite HOR array (active_hor-S1C1-5-19H1L).** The break in the diagonal 100% identity red line with 62 kb of parallel sequence indicates this is a tandem duplication of sequence rather than an expansion of a larger array of many copies of the same sequence, like most CenSat insertions >2kb. It starts at a smaller high identity repeat (small red box where the top diagonal alignment starts). The x coordinates are in the HG008Nv6.3 normal assembly between chr1_hap1:125867435-126489155, and the y coordinates are in the HG008Tv3.3 tumor assembly between chr1_hap1:125750374-126434263. Black bars show the size (but not exact position) of the duplication.

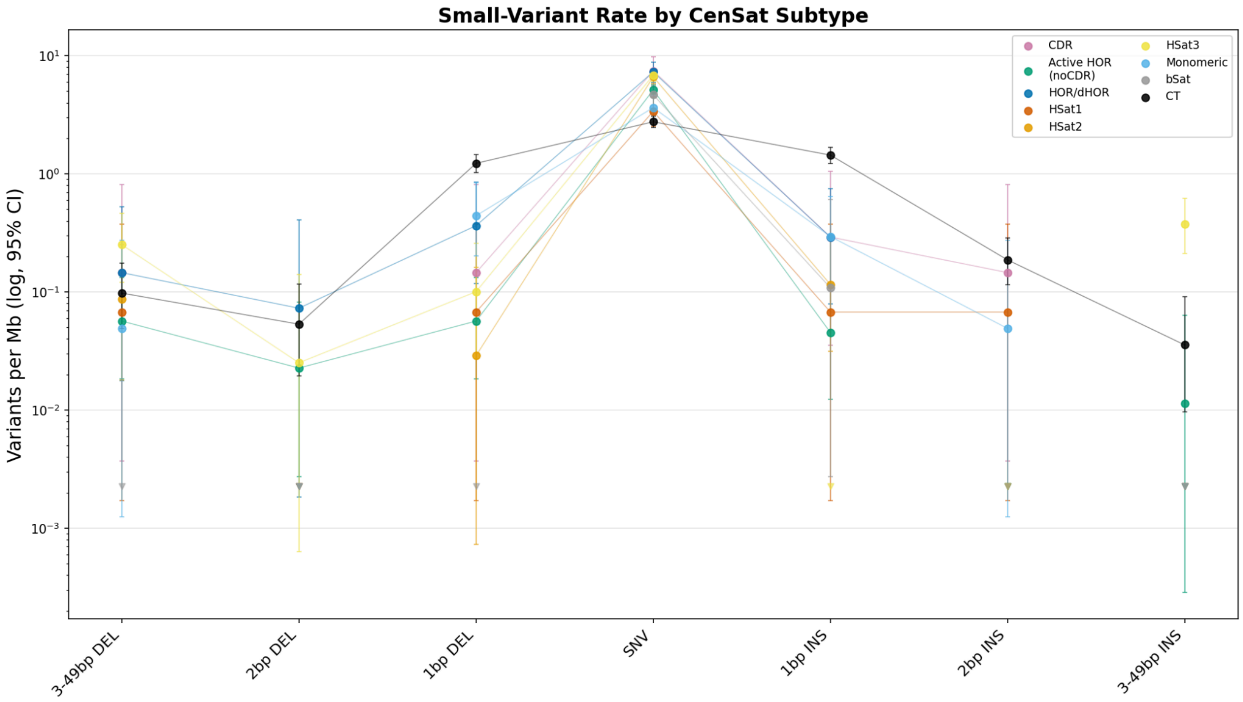


**Supplementary Fig. 27: Centromere satellite small variant rates vary by type and are higher for SNVs than in the centromere transition (CT) regions, which are similar to whole genome rates.**

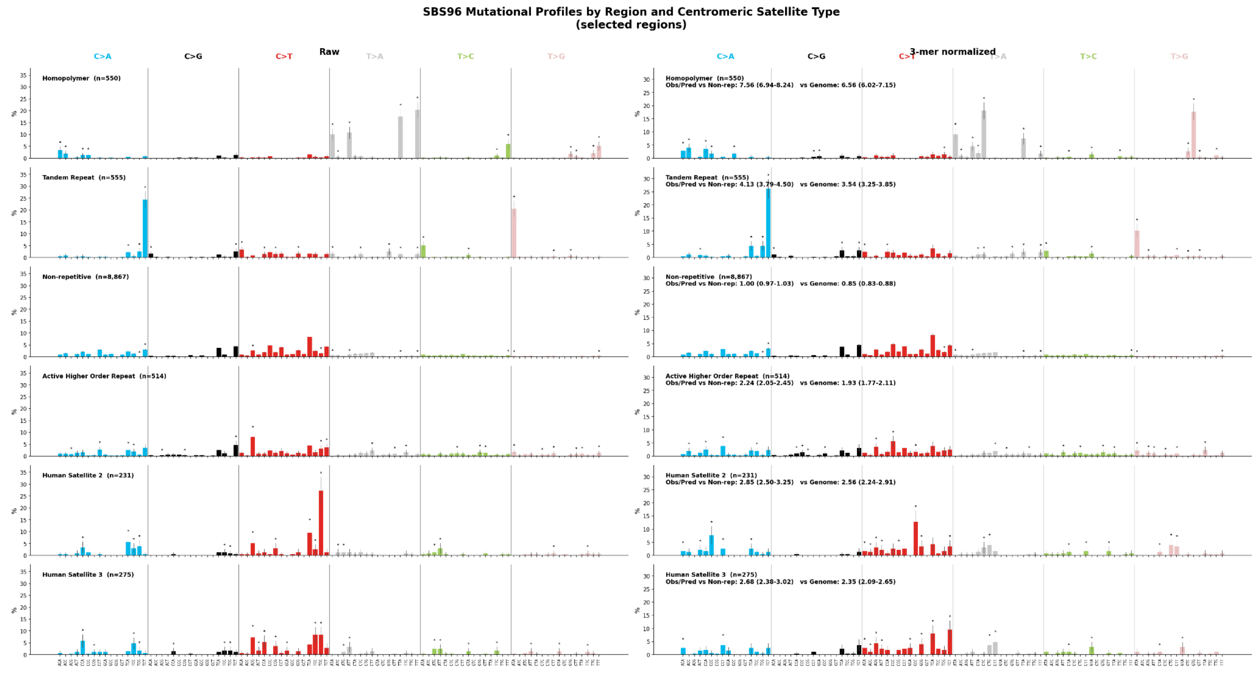


**Supplementary Fig. 28: 96-channel SBS spectrum across genomic regions, with and without trinucleotide frequency normalization. Mutational spectra also vary across repetitive regions, even after correcting for trinucleotide frequencies.** Homopolymer regions show the largest excess (Obs/Pred = 7.56 vs non-repetitive), driven primarily by T>A and T>G mutations in adjacent homopolymers and homopolymers next to dinucleotide TRs. Human Satellite 2 (Obs/Pred = 2.85) is dominated by C>T transitions at CpG dinucleotides, indicative of 5-methylcytosine deamination in hypermethylated heterochromatin. Active Higher Order Repeats (Obs/Pred = 2.24) and Human Satellite 3 (Obs/Pred = 2.68) display broader mutational spectra suggesting contributions from both CpG deamination and replication-associated errors, possibly due to less efficient repair in late-replicating heterochromatin. All repeat regions show mutation rates significantly exceeding predictions from trinucleotide composition. Each row shows the 96-channel SBS spectrum for a genomic region, with raw counts (left) and 3-mer normalized counts (right). Normalization corrects for differences in trinucleotide composition between regions by scaling each channel by the ratio of genome-wide to region-specific 3-mer frequency, preserving total mutation count. Error bars indicate 95% confidence intervals (binomial approximation). Asterisks denote channels where the per-base pair mutation rate is significantly different from the whole-genome rate (Poisson rate ratio z-test, 99% confidence). Panel annotations show the Obs/Pred ratio relative to non-repetitive and whole-genome baselines, where Observed is the measured SNV rate per bp and Predicted is the expected rate based on trinucleotide composition alone.

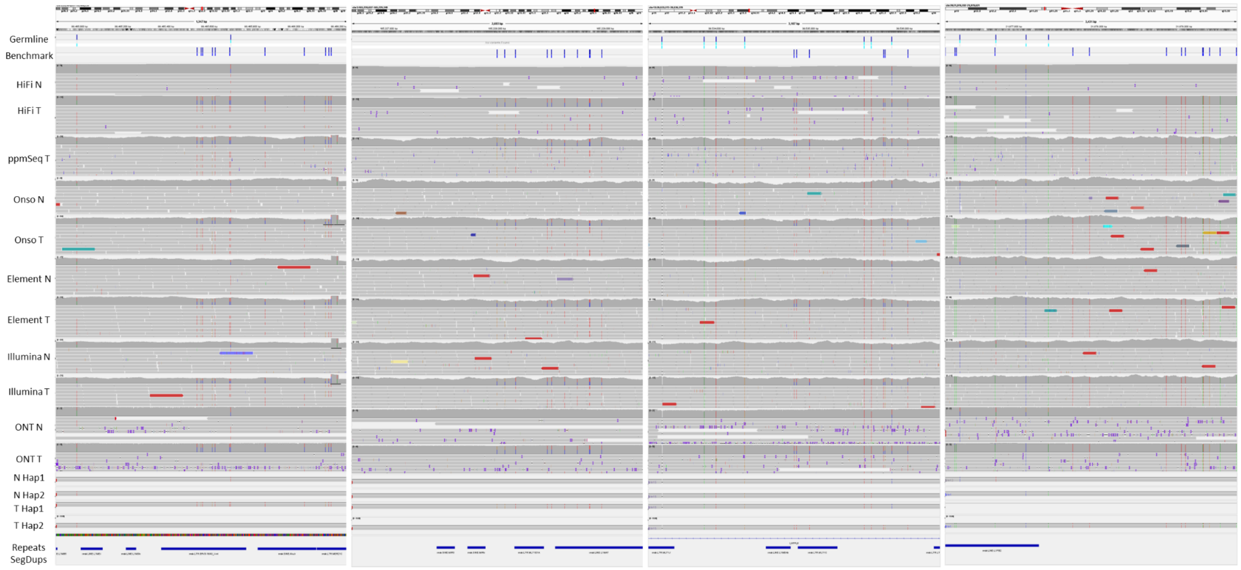


**Supplementary Fig. 29: Kataegis events.** There are some real clusters of somatic variants that appear to be typical kataegis events. From left to right, chr3:99465564 has ~10 C>T and 3 C>G within 500bp and similar clusters in chr3:165228003, both on hap1. There are also 6 SNVs around chr13:39534827 on hap2 and 14 SNVs in chr20:21976192-21979621.Examining clusters of SNVs, several appear to be in or near centromeres which were confirmed by centromere annotation including chr5_hap1:48340801-48343247 that seems to be due to suboptimal alignment of SV in centromere. A cluster of 6 SNVs in chr7_hap1:63886051-63886695 that is in the centromere track. Another cluster of 11 SNVs in chr20_hap2:30329346-30330733 between 2kb and 11kb deletions which is not an obvious suboptimal alignment since it has evidence in mapped reads too. Finally, a cluster of 5 SNVs in chr11_hap2:51767572-51767822 that is also within a centromere track.

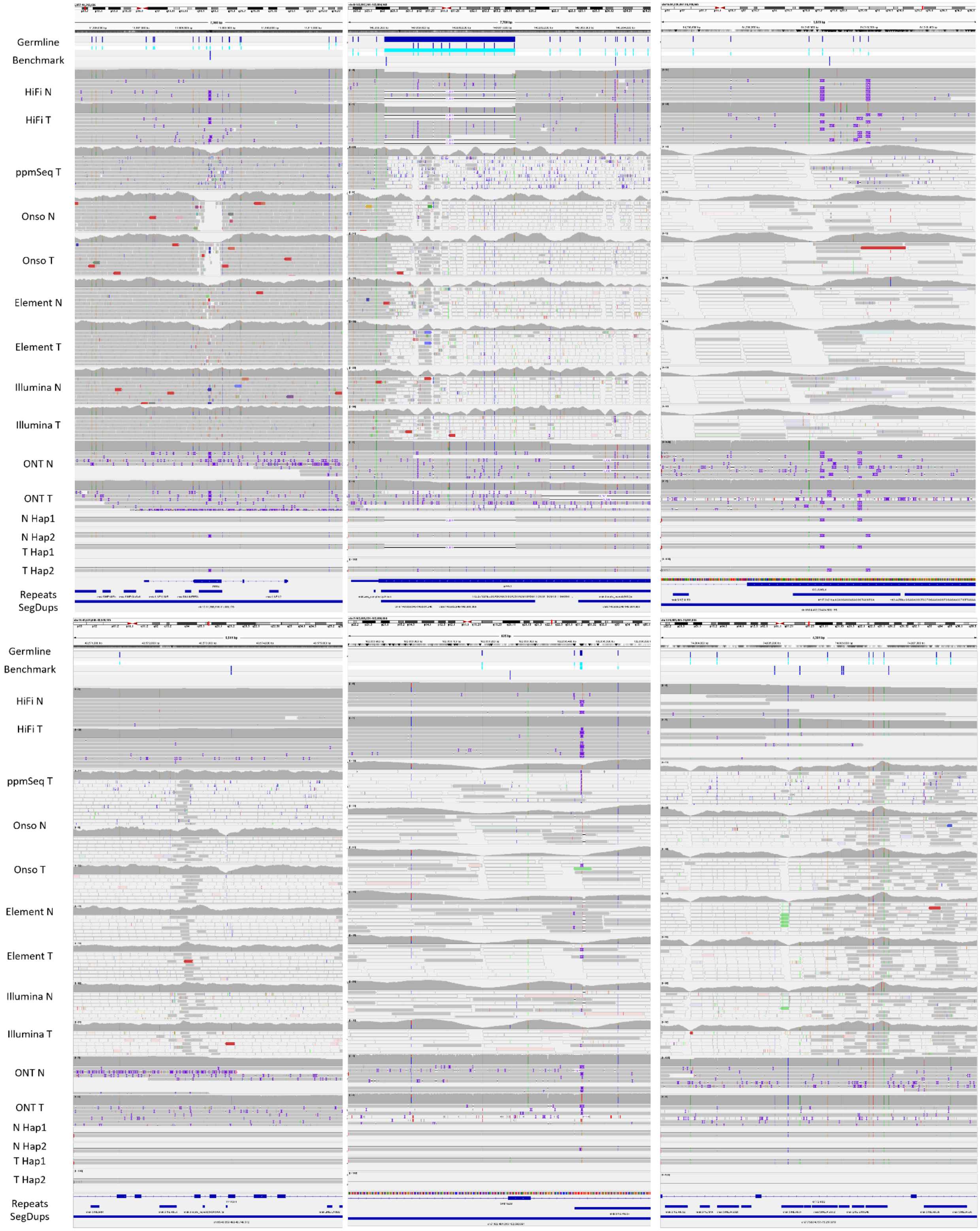
Germline
Benchmark
HiFi N
HiFi T
ppmSeq T
Onso N
Onso T
Element N
Element T
Illumina N
Illumina T
ONT N
ONT T
N Hap1
N Hap2
T Hap1
T Hap2
Repeats
SegDups
Germline
Benchmark
HiFi N
HiFi T
ppmSeq T
Onso N
Onso T
Element N
Element T
Illumina N
Illumina T
ONT N
ONT T
N Hap1
N Hap2
T Hap1
T Hap2
Repeats
SegDups

**Supplementary Fig. 30: Assemblies resolve true somatic variants in segmental duplications:** These regions typically have high false positive and false negative rates with mapping-based approaches. There are TR-associated variants in *PRB4*, *EPPK1*, *GOLGA6L4*, and as well as non-TR coding variants in PPIP5K1 (chr15:43573416), SPDYE2B exon 5, and a cluster of SNVs/indels in GTF2IRD2 (chr7:75074151-75217878). A particularly interesting TR example illuminated by the personalized reference is a stop-gain somatic SNV near the end of a 63 bp germline insertion in a VNTR in *PRB4* (BCSQ annotation of tumor insertion: stop_gained&inframe_insertion|PRB4|ENST00000279575|protein_coding|-|122R>122G*|11308619 T>TTTCTGGCTTTCCTGGATGAGGTGGGGGACCTTGGGACTGGTTGCCTCCTTGTGGGGGTCATCC) . The second codon from the end changes from CGA (arginine) in the normal to TGA (stop) in the reverse complement. While *PRB4* has only rarely been associated with cancer, this type of change would be very difficult to detect without long reads and a personalized reference. Relatedly, a coding VNTR in *EPPK1* has a synonymous SNV inside a 12 bp germline insertion (chr8:143863761 G>GGGCAGCGGCGGC), and a coding VNTR in *GOLGA6L4* has a synonymous SNV that causes a 21 bp germline insertion to shift alignment by 24 bp (chr15:84240066 G>GAACAGGAGGAGAGGCTGTGTA), both of which are difficult to detect with standard methods. *SPDYE2B* SNV in exon 5 that appears nonsynonymous but has no coverage in gnomAD due to low mappability.

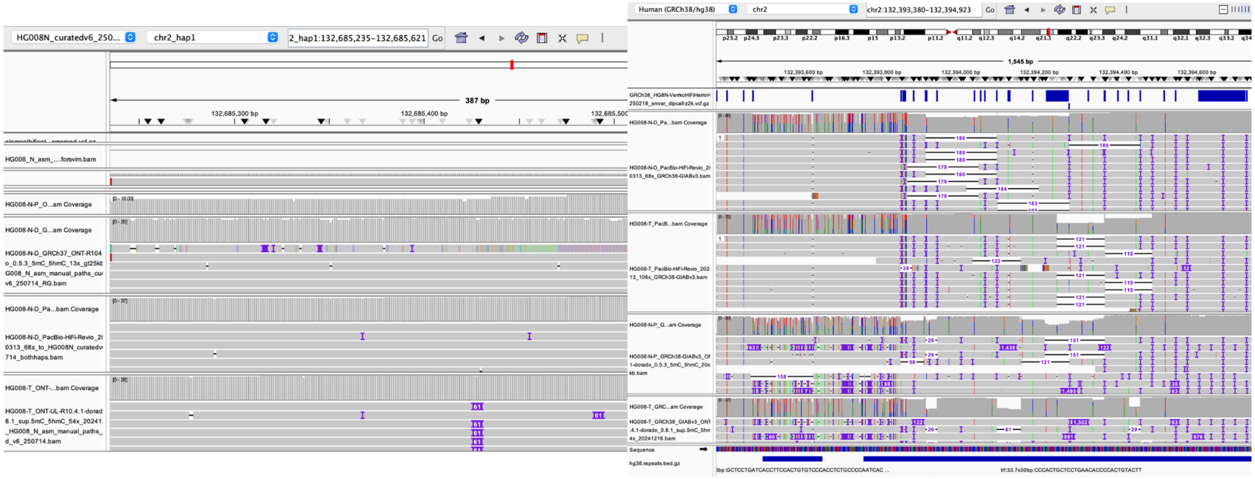


**Supplementary Fig. 31: Large STR and VNTR Expansions and Contractions:** These appear as simple insertions and deletions with respect to the normal assembly, but became complex with respect to GRCh38 due to overlap with germline variation. Most somatic variants in STRs and VNTRs were not identified by mapping-based approaches because all had germline SVs and/or large indels as well. Due to a 42 bp germline insertion in one VNTR, a 70 bp somatic deletion, which falls above the typical SV size cutoff, appears as a 28 bp deletion with respect to GRCh38, so that somatic SV callers would typically ignore this. There's also a 60 bp expansion of a VNTR around chr2:132394152 that contains very complex germline variation making it challenging to understand or represent the variant with respect to GRCh38, but it is very clear on the normal reference (https://github.com/jzook/HG008SVcuration/issues/323). In addition to VNTR expansions and contractions, breakpoints on one end of a complex 36 Mbp somatic inversion on chr7 fall inside a 611 bp VNTR that also has germline variation, which complicates detection and precise representation of the breakpoints on GRCh38, but breakpoints are clear when using the normal as the reference (https://github.com/jzook/HG008SVcuration/issues/199).

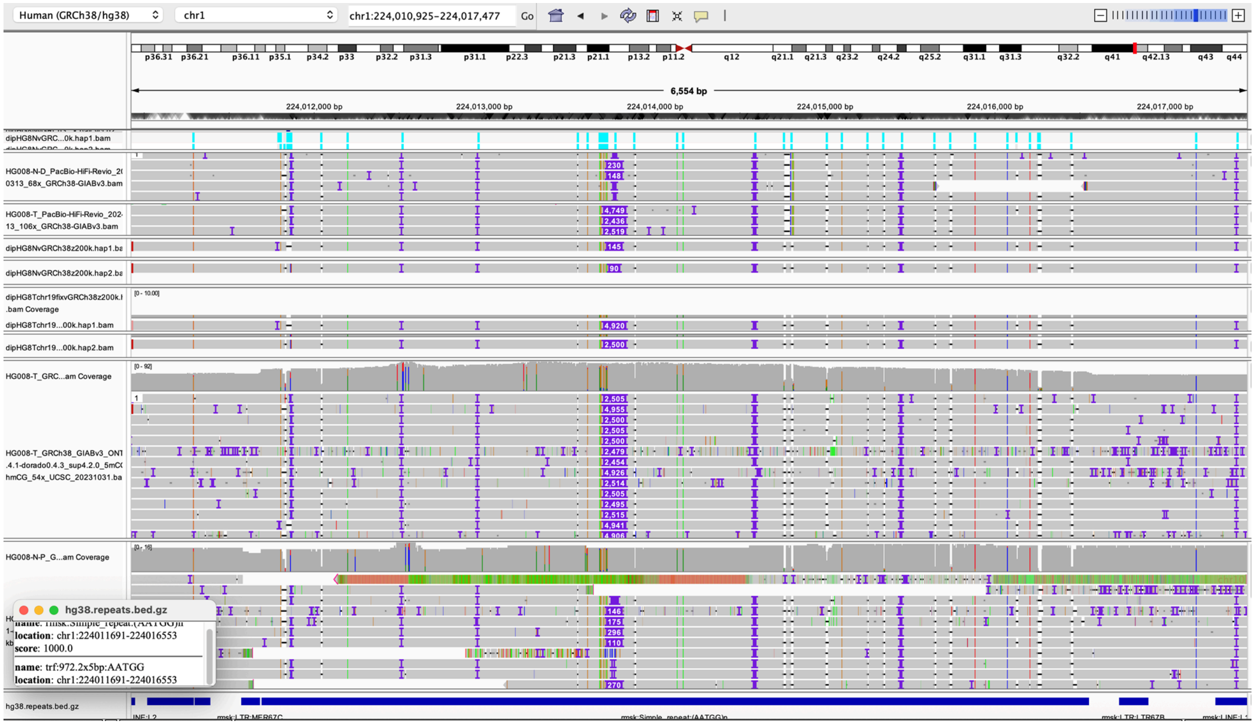


**Supplementary Fig. 32: Large intronic tandem repeats.** The assemblies resolve an intronic TR in the *SEPTIN7P13* segdup, which has ~4800 bp somatic expansion on hap1 and ~2400 bp somatic expansion on hap2 of the high-identity repeat in the middle of an imperfect repeat (https://github.com/jzook/HG008SVcuration/issues/326) we subsequently identified in a CenSat track as a small HSat3 array. It also appears to have subclonal variability in both normal tissues whereas the tumor cell line has much less mosaicism even though the repeat is longer, so it may be a particularly unstable TR in normal tissues (similar to a TR on chr4 https://github.com/jzook/HG008SVcuration/issues/386). This TR is also challenging because extra HiFi reads mismap here on GRCh38 that match a segdup at the beginning of chr1 on the normal assembly. The assemblies help both in minimizing mismapped reads as well as enabling resolution of the different expansions on each haplotype. There is an interesting CT-rich VNTR at chr6_hap2:101427602 that seems to have a somatic 60bp INS supported by almost all tumor reads, but the normal pancreas HiFi and ONT have a mix of lengths ranging from no INS to ~60 bp. Most insertions and many deletions 4 to 49 bp in size also fall in TRs, as described in the small variants section.

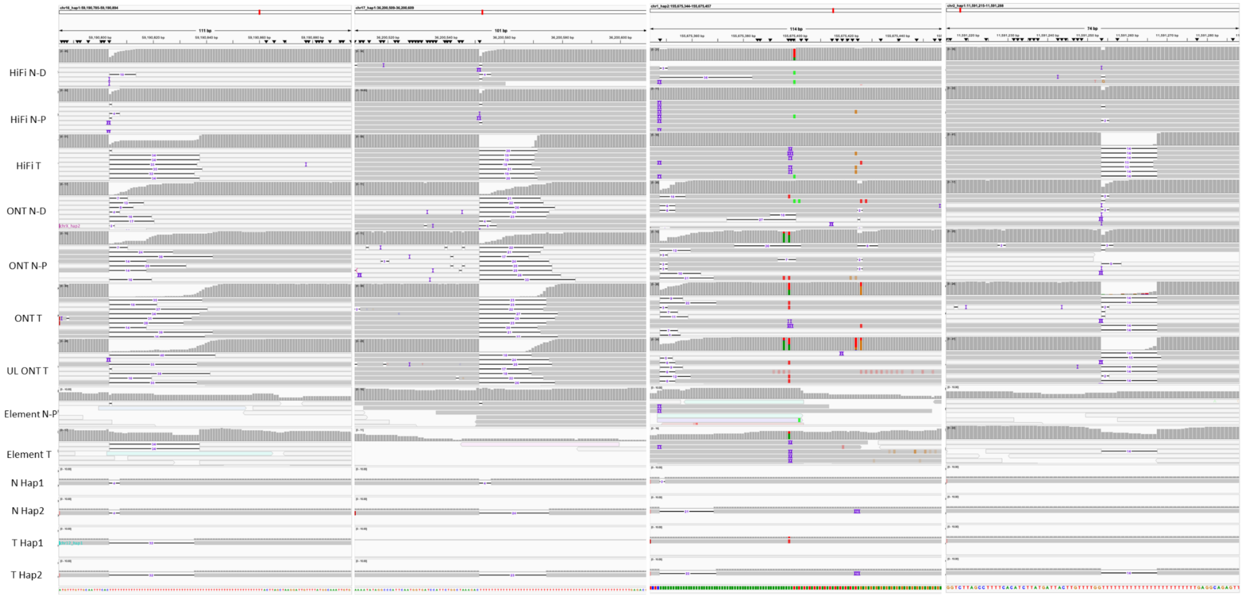


**Supplementary Fig. 33: Large Indels in Homopolymers and TRs**. When aligning to both haplotypes of HG008-N, several deletions and insertions are evident. Deletion examples include a somatic 34bp contraction of 58bp homopolymer at chr16_hap1:59190801 and a 19bp contraction of 52bp homopolymer at chr17_hap1:36200559. Also, 7 contractions of 10-30bp in homopolymers longer than 50bp are present in the benchmark. Further, there is a 7bp INS at a 52bp homopolymer 27bp diTR junction at chr1_hap2:155675392. For deletions, there are 14 deletions >=10bp in homopolymers <50bp, with the shortest being a 14bp contraction of a 24bp homopol at chr2_hap1:11591252. In contrast, all 5 insertions >10bp in homopolymers appeared likely to be assembly errors, mostly haplotype switches in the normal assembly. In addition, there are several large TR expansions in the benchmark: a 40bp INS at chr5_hap2:54605160, a 35bp INS at chrX_hap1:107227010, and a 34bp INS at chrX_hap1:23434627.

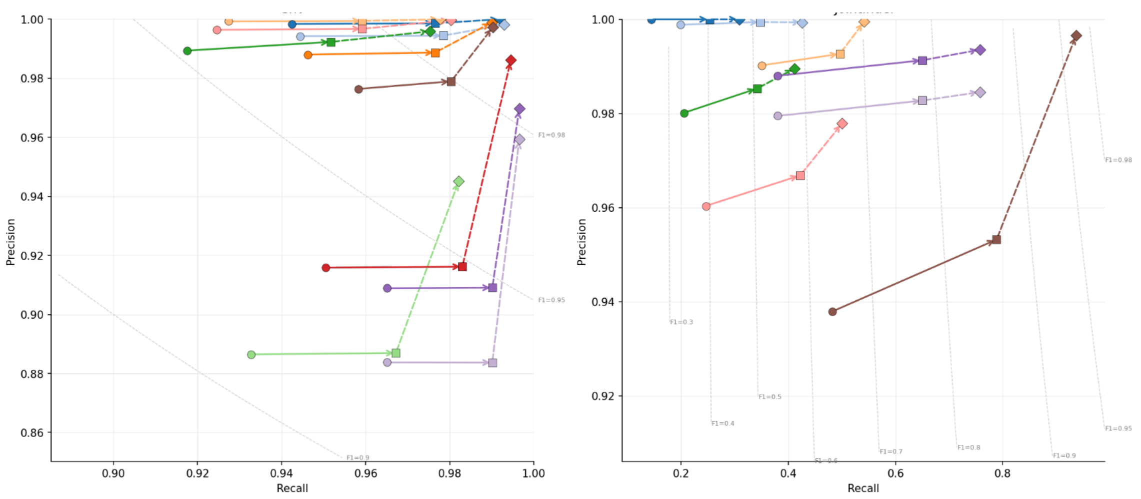


**Supplementary Fig. 34: Accuracy is lower for somatic variants near germline variants.** Performance across anonymized variant callers for SNVs (left) and indels (right) is better when somatic variants near germline variants are excluded (diamonds), or somatic variants whose representation is affected by germline variants are excluded (squares), vs. all GRCh38 benchmark somatic variants (circles).

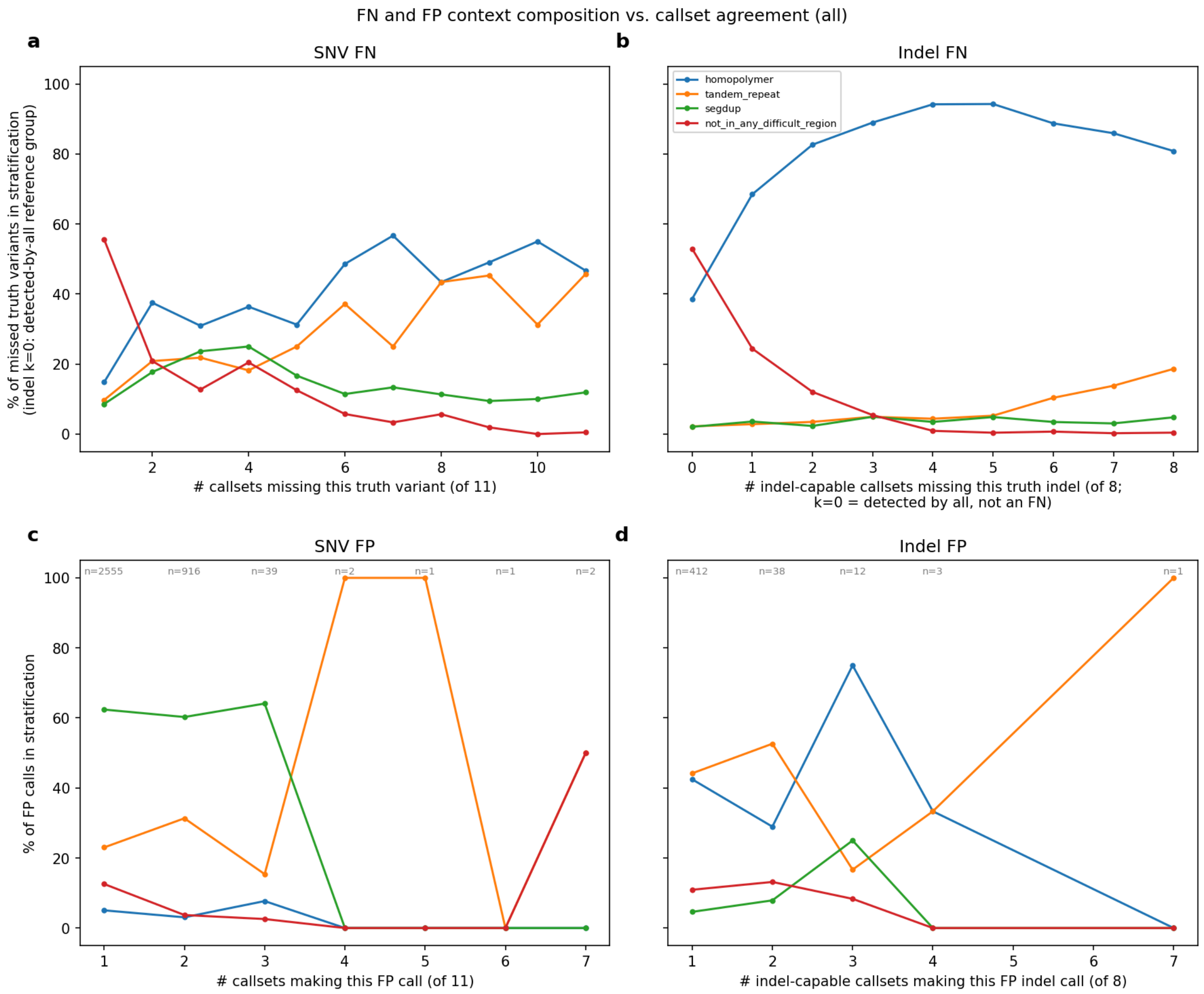


**Supplementary Fig. 35: Sources of FPs and FN.** Plotting the SNV and Indel FPs and FNs across different stratifications shows the impact of genomic context such as repetitive regions on variant calling performance.

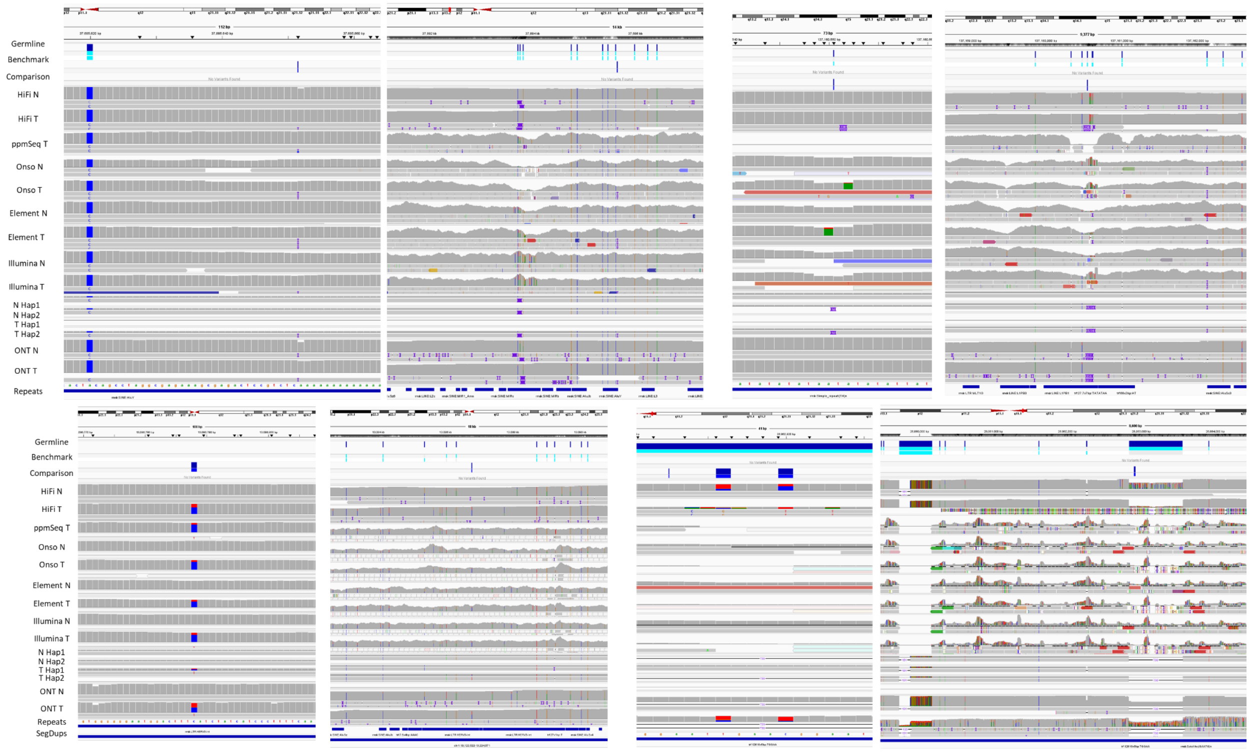


**Supplementary Fig. 36: False Positive and False Negatives Commonly Occur in Repetitive Regions** From visually inspecting over 180 discrepant small variants when comparing callsets to the draft benchmark, we identified that most of the discrepant calls are in repetitive regions such as homopolymers, SINEs, LINEs, tandem repeats, and segmental duplications. In this process, we considered a draft benchmark variant correct if the variant was supported by alignments of the tumor but not normal assemblies, and by tumor but not normal reads. We focused on reads from technologies with no evidence of bias or errors at the variant position, such as long reads in low mappability regions and segmental duplications or short reads that map across the repetitive unit in homopolymers or tandem repeats. Commonly, we identified that comparison callsets contained false positive somatic variants that matched or were near a germline variant in the normal sample. As a result of this process, we confirmed the benchmark reliably identified false positives and false negative SNVs and INDELs across comparison callsets. Examples include: (upper left) Homopolymers and SINEs - false negative 1bp insertion located in a SINE that is representative of many homopolymer FNs being in the polyA of Alus; (upper right) Tandem Repeats - false positive matches germline 14bp insertion in a tandem repeat and an LOH region; (lower left) Segmental Duplications - false negative due to difficult read mapping in Segmental Duplication; and (lower right) Small Variants within germline SVs – false positive SNV due to misaligned reads in germline 730bp deletion in a TR.

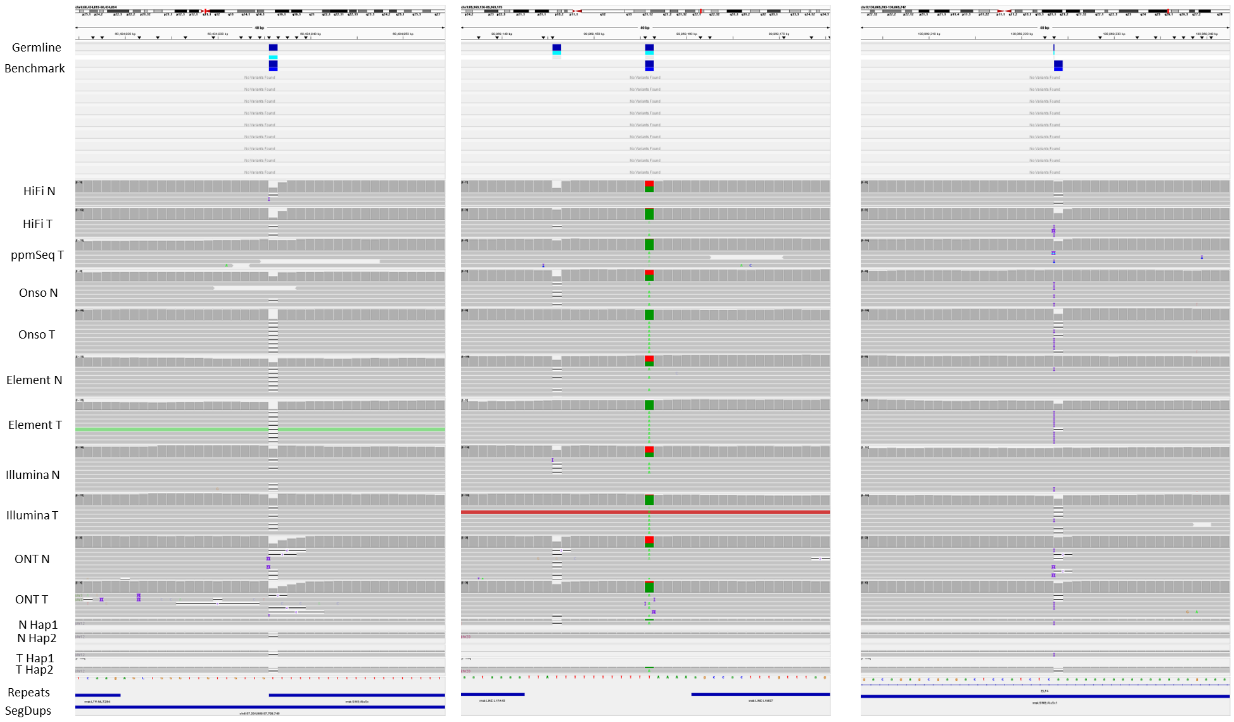


**Supplementary Fig. 37: Variants Missed All Mapping-Based Methods.** A class of somatic variants that are commonly missed in comparison between mapping-based methods and the somatic variant benchmark are those that modify germline variants, which usually occur in homopolymers and tandem repeats. These include the following types of modifications: those loci that appear to not be a variant in the tumor due to reversion of a germline variant to reference; those changing a germline variant to different variant; those that convert germline heterozygous to homozygous variant that looks similar to LOH (these appear in the current VCF file as heterozygous calls due to haplotype-specific calling); and other germline variation that makes it difficult to see true somatic change. There are also a small number of sites with somatic variants on both haplotypes.

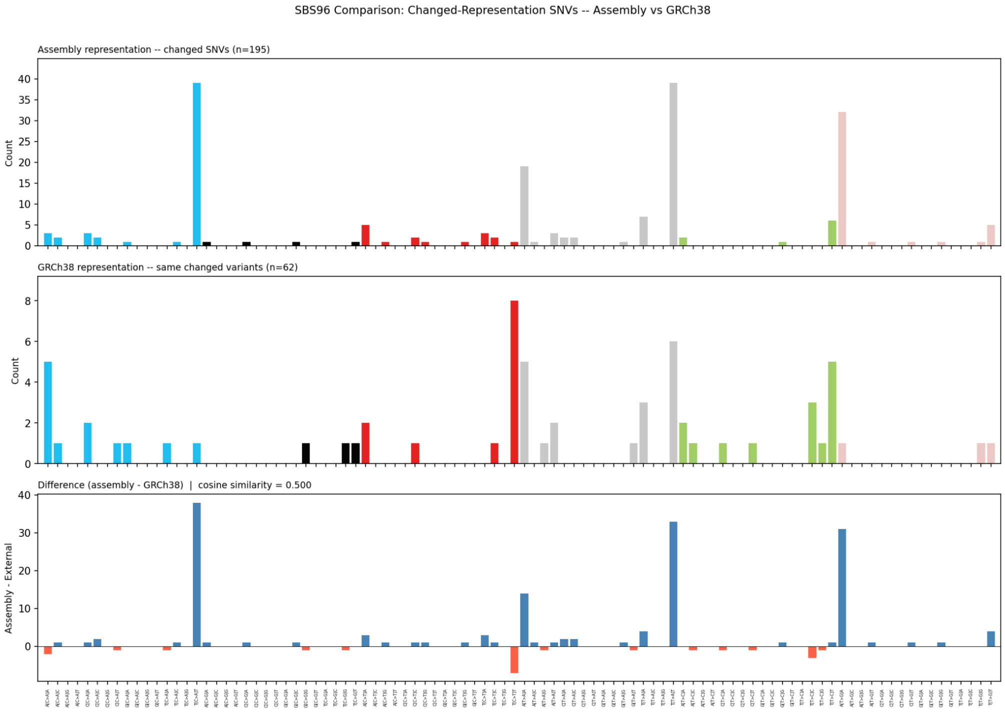
SBS96 Comparison: Changed-Representation SNVs -- Assembly vs GRCh38
Assembly representation -- changed SNVs (n=195)
Count
GRCh38 representation -- same changed variants (n=62)
Count
Difference (assembly - GRCh38) | cosine similarity = 0.500
Assembly - External

(a)

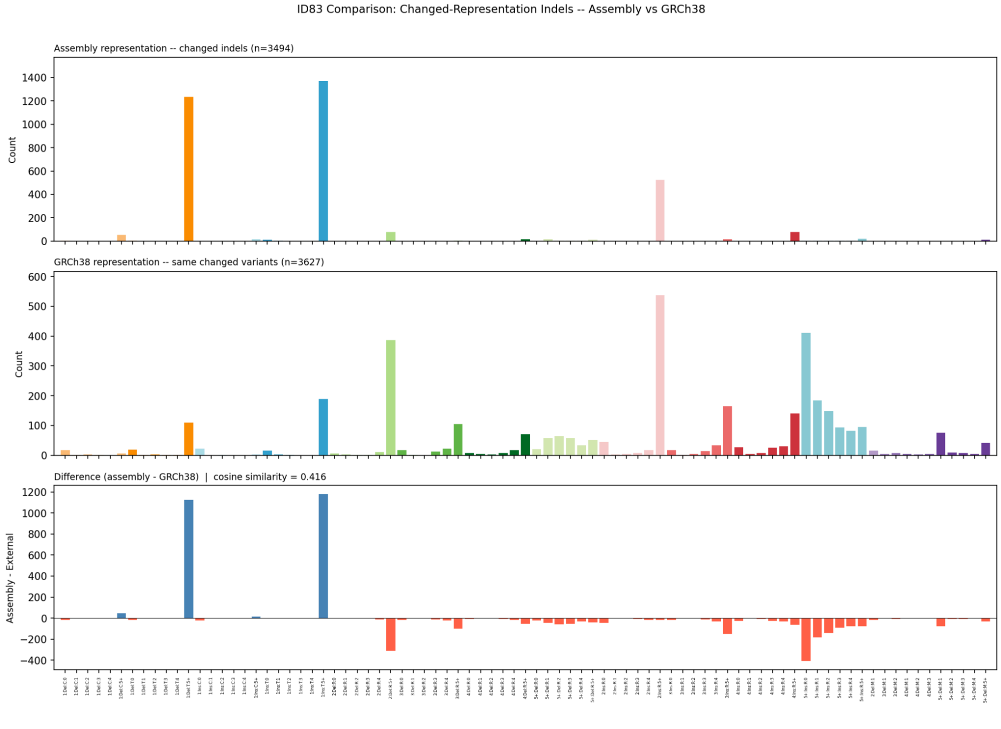
ID83 Comparison: Changed-Representation Indels -- Assembly vs GRCh38
Assembly representation -- changed indels (n=3494)
Count
GRCh38 representation -- same changed variants (n=3627)
Count
Difference (assembly - GRCh38) | cosine similarity = 0.416
Assembly - External

(b)

**Supplementary Fig. 38: Variants changing representation on GRCh38 affect mutational spectra.** The SBS 96-channel (a) and ID 83-channel (b) spectra for the correct normal assembly-based representation (top) differ when variants change representation on GRCh38 due to germline variation. SNVs on the normal assembly primarily fall in 4 channels because they fall at the boundaries of homopolymers and/or tandem repeats, but these change to many other channels on GRCh38. Similarly, indels are in 3 channels because they are in long homopolymers and tandem repeats, but change to many other channels on GRCh38 due to germline variation.

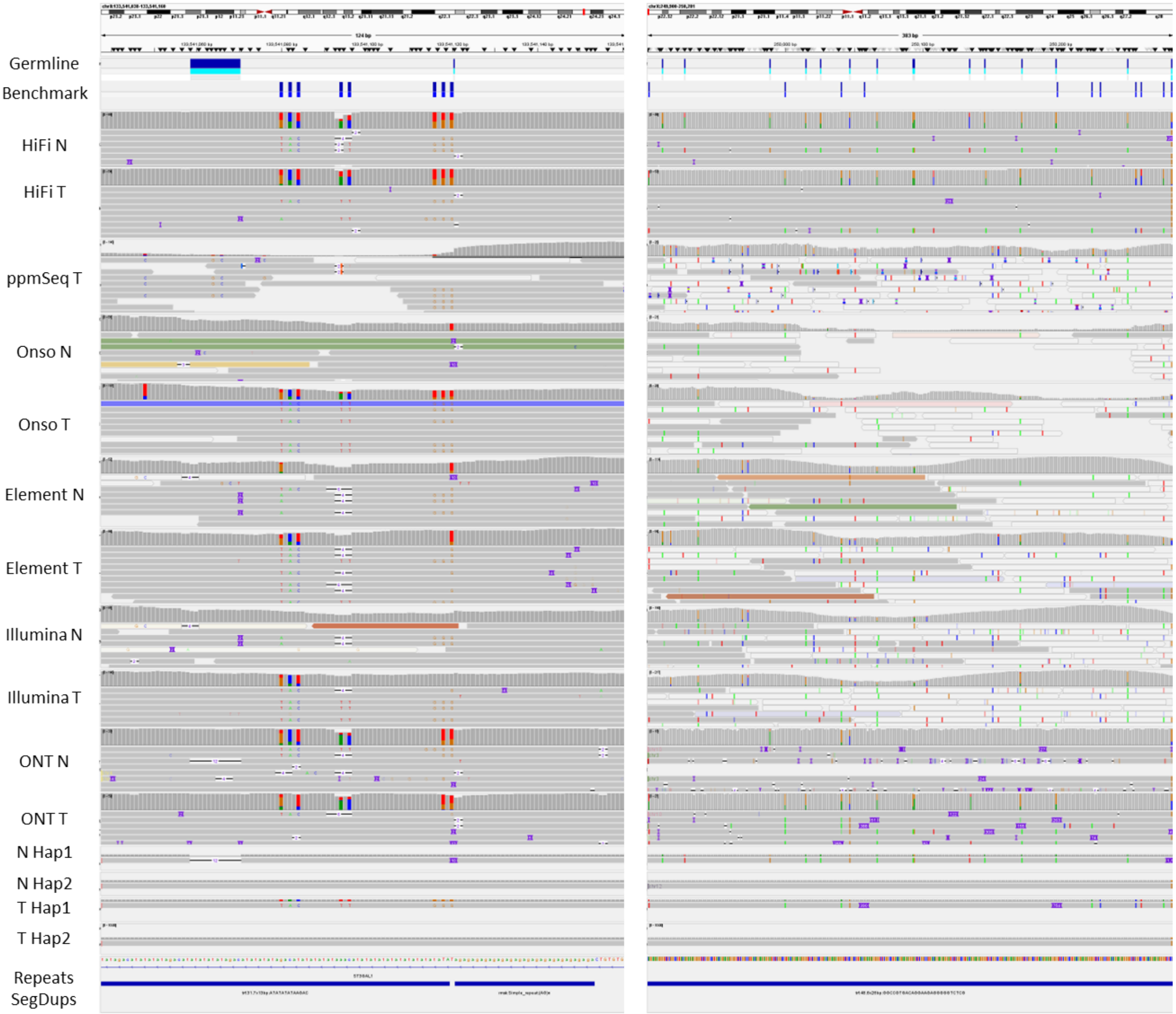


**Supplementary Fig. 39: Somatic variation frequently alters germline variants.** Looking at variants lifted to GRCh38 in homopolymers and tandem repeats, somatic indels can change in size or type or appear as SNVs in the tumor reads aligned to GRCh38. As an extreme example, in chr8:133541079-133541120, a 2 bp somatic insertion causes a 12bp deletion and 10bp insertion in the normal to change to 8 SNVs in adjacent diTRs. Another example is chrX:249900-250281, a 26 bp deletion in a VNTR causes 9 SNVs and 2 large insertion SVs. When clustering variants within 1 kbp on GRCh38, 47 of 104 clusters of >= 3 somatic variants are actually caused by a single somatic variant. These 47 somatic variants cause 204 apparent somatic variants on GRCh38, indicating that variant clusters on GRCh38 are often actually a simple single somatic change.

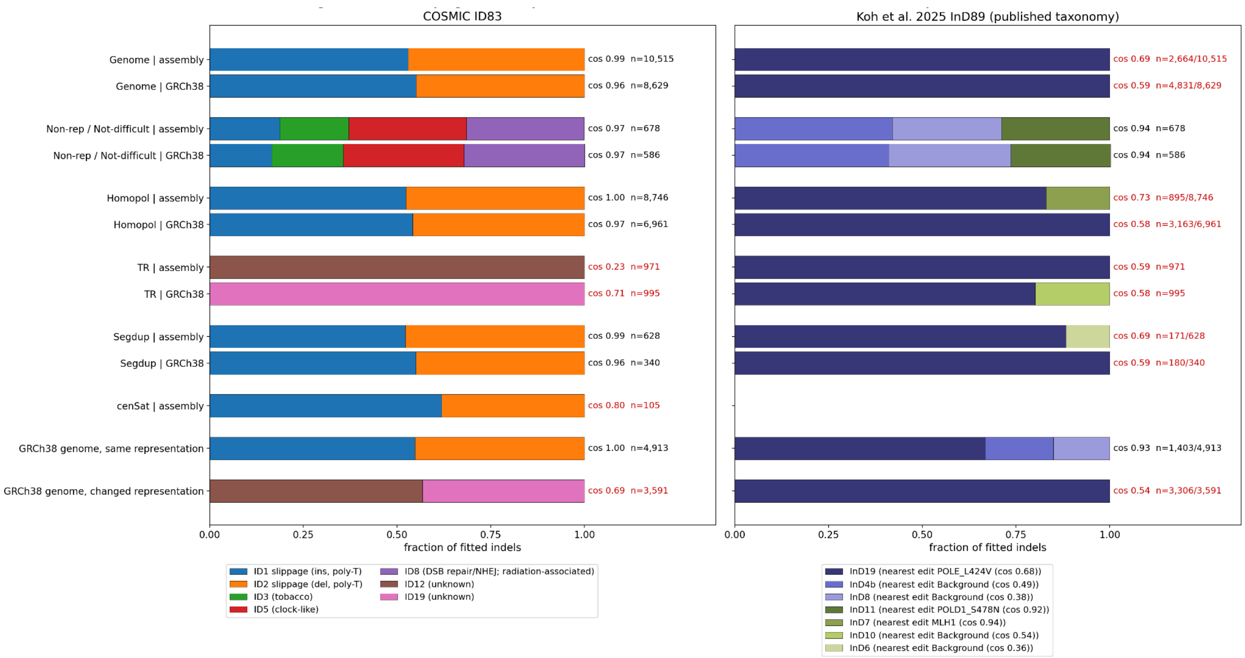


**Supplementary Fig. 40: Indel signatures vary across genomic regions.** V0.4 benchmark somatic indels were fitted to COSMIC v3.6 ID83 signatures (left) and to the 37 consensus indel signatures of Koh et al. (2025) under their published 89-channel classification (right). Homopolymers are highly enriched for 1 bp indels and STRs and VNTRs are highly enriched for >1 bp indels, particularly insertions. Because centromeric satellites have fewer homopolymers, STRs, and VNTRs, they are enriched for larger indels and microhomology-mediated deletions, similar to non-repetitive regions. The signatures can also change on a donor-specific reference vs GRCh38 because 1 bp somatic indels can change representation to larger indels or SNVs on GRCh38 due to germline variation. COSMIC used SigProfilerAssignment, and Koh et al. used non-negative least squares and indelsig.tools.lib. The assembly non-repetitive region excluded homopolymers, tandem repeats, centromeric satellites, and segmental duplications, similar to but not as comprehensive as the GIAB GRCh38 "not in all difficult regions" stratification. Centromeric satellites (cenSat) are shown for the assembly only, because the GRCh38 benchmark excludes satellite sequence; the Koh et al. fit is omitted for cenSat because fewer than 100 indels remained after classification. The final two rows split GRCh38 benchmark indels by whether the matching assembly call has the same or a changed representation. Signatures contributing at least 5% of indels in any bar are shown individually. Each Koh et al. signature is labelled with its most similar experimental gene-edit signature from that study and the corresponding cosine similarity. Text beside each bar gives the reconstruction cosine similarity, in red when below 0.9, and the number of indels fitted. Where n is given as a fraction, the second number is the total number of indels, because the published 89-channel classification excludes 1 bp indels in homopolymers of 10 bp or longer. In non-repetitive sequence, both signature sets fitted well and agreed between coordinate systems (cosine 0.94–0.97). COSMIC attributions there included ID8, which is associated with double-strand-break repair and ionizing radiation, and Koh et al. attributions included InD11, whose closest gene-edit signature is the POLD1 S478N polymerase mutant.

Homopolymer indels were explained almost entirely by the replication-slippage signatures ID1 and ID2 (cosine 0.97–1.00). However, the published Koh et al. classification fit repetitive regions poorly. Tandem-repeat indels were poorly explained by either signature set (cosine 0.23–0.71). In GRCh38 coordinates, indels with the same representation as the assembly call were explained by replication slippage (cosine 1.00). Indels whose representation changed relative to the assembly call were not well explained by any known signature (cosine 0.54–0.69).

.

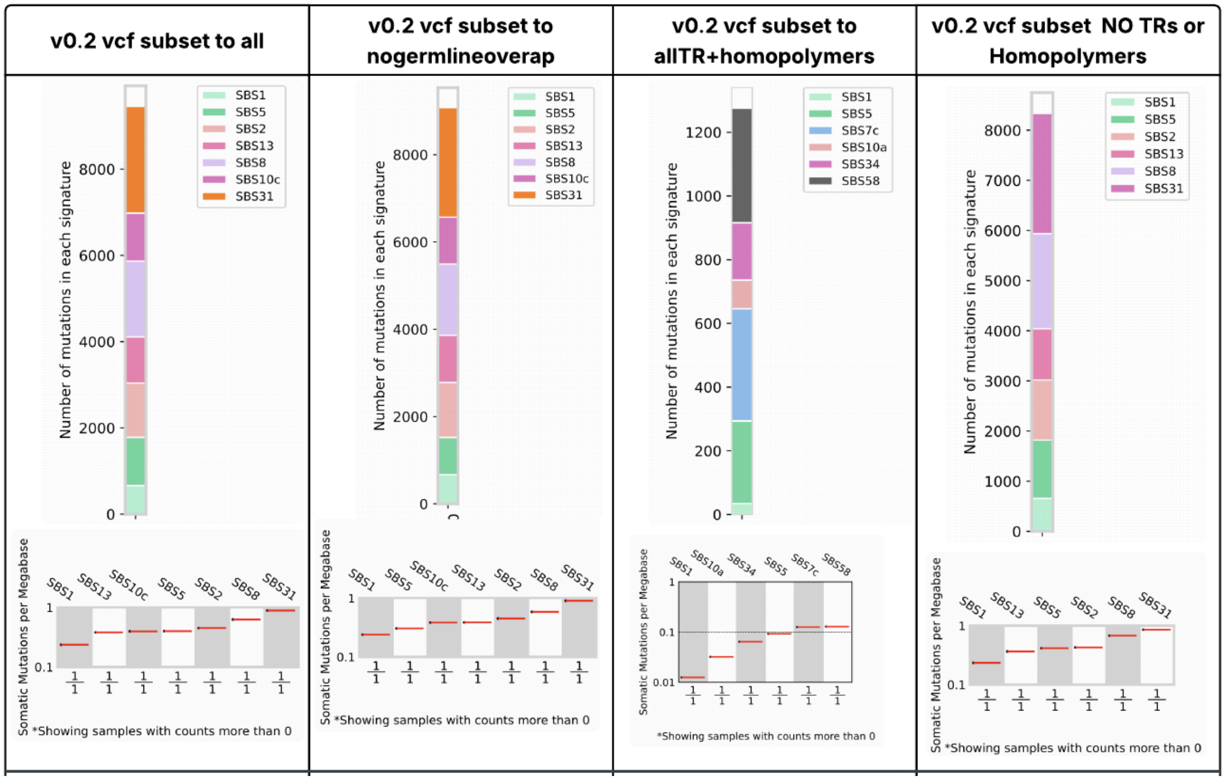


**Supplementary Fig. 41: Mutational Signatures in GRCh38 benchmark.** Using SigProfilerAssignment with the v0.2 small variant vcf subset to all benchmark regions (all), regions excluding somatic variants that alter germline variants (nogermlineoverlap), Tandem Repeats and Homopolymers (allTR+homopolymers), and outside Tandem Repeats and Homopolymers (No TRs or homopolymers). Single base substitutions (SBS) results for all.vcf seem consistent with PDAC (SBS1, 2, 5, 8, 13), and SBS31 is associated with platinum chemotherapy treatment which matches HG008-T intervention history. When subsetting to homopolymers and TRs, all of the signatures except SBS1 and SBS5 are enriched for motifs that occur at homopolymer/homopolymer, homopolymer/diTR, and diTR/diTR junctions where SNVs in these repeats often seem to occur.

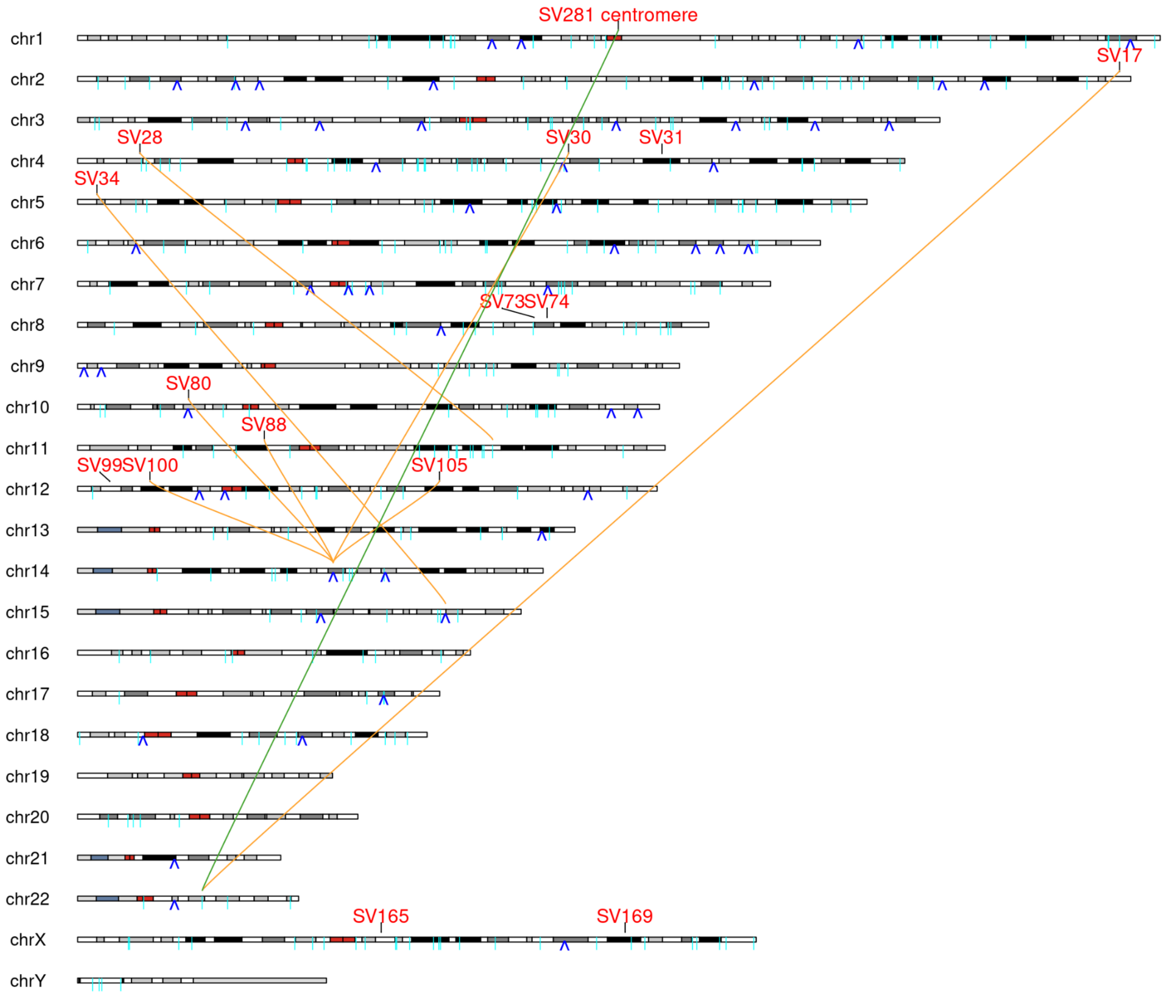


**Supplementary Fig. 42: Location of LINE retrotranspositions and their source (where determined) yellow and green lines**. Green line indicates insertion into chr1_hap2:124339632 (HG008Nv6.3). Cytoband data: http://hgdownload.cse.ucsc.edu/goldenpath/hg38/database/cytoBand.txt.gz. Cyan line markers: GRCh38 LINE-1 locations taken from https://www.repeatmasker.org/genomes/hg38/RepeatMasker-rm406-dfam2.0/hg38.fa.out.gz. Blue inverted V markers: LINE-1 locations from HG008-N-P HiFi pbsv Normal VCF.

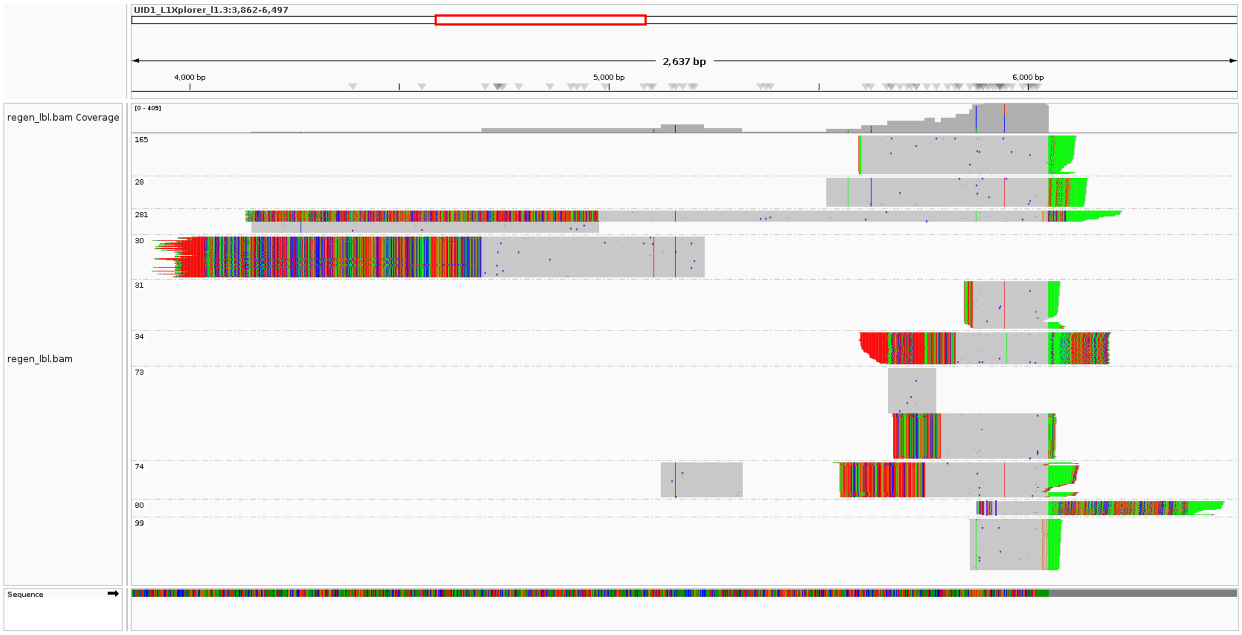


**Supplementary Fig. 43: Somatic L1 insertions mapped to a reference L1 sequence.** poly-A tails visible as green or red at end of insertions. Numbers are SV IDs from Supplementary Tables S10 and S12. Twin-priming events are also visible on 30, 34, 73, 74 and 281.

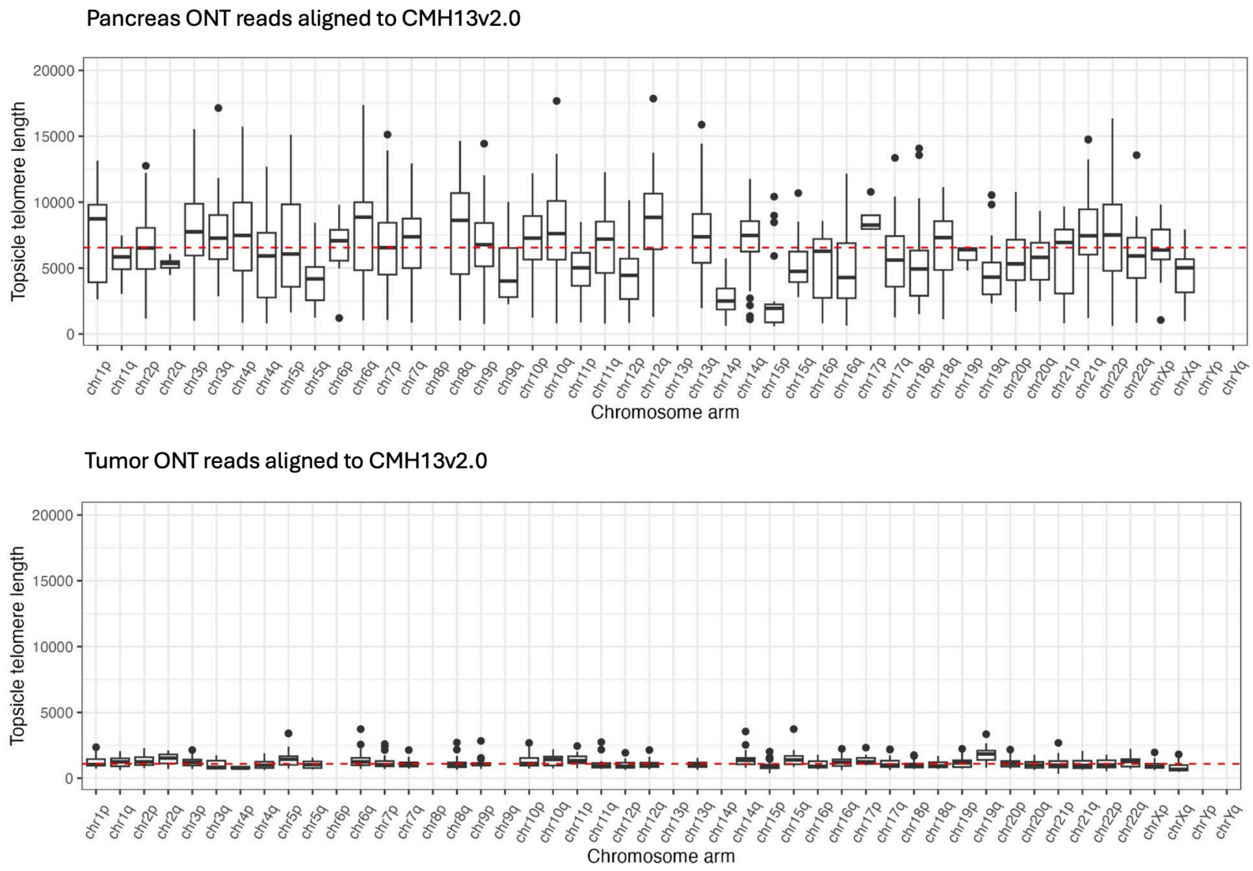


**Supplementary Fig. 44: Estimated lengths of telomeres in normal pancreatic tissue and tumor cell line using Topsicle with telomeres assigned based on alignments to GRCh38.** (a) Telomere lengths of all individual alleles (except unresolved 21p and 22p) in HG008-N-P, as measured from ONT long reads. (b) Telomere lengths of 66 alleles identified in HG008-T, as measured from ONT long reads. The absent alleles correspond to telomere ends that are not present in the tumor genome due to deletions or chromosome fusions.

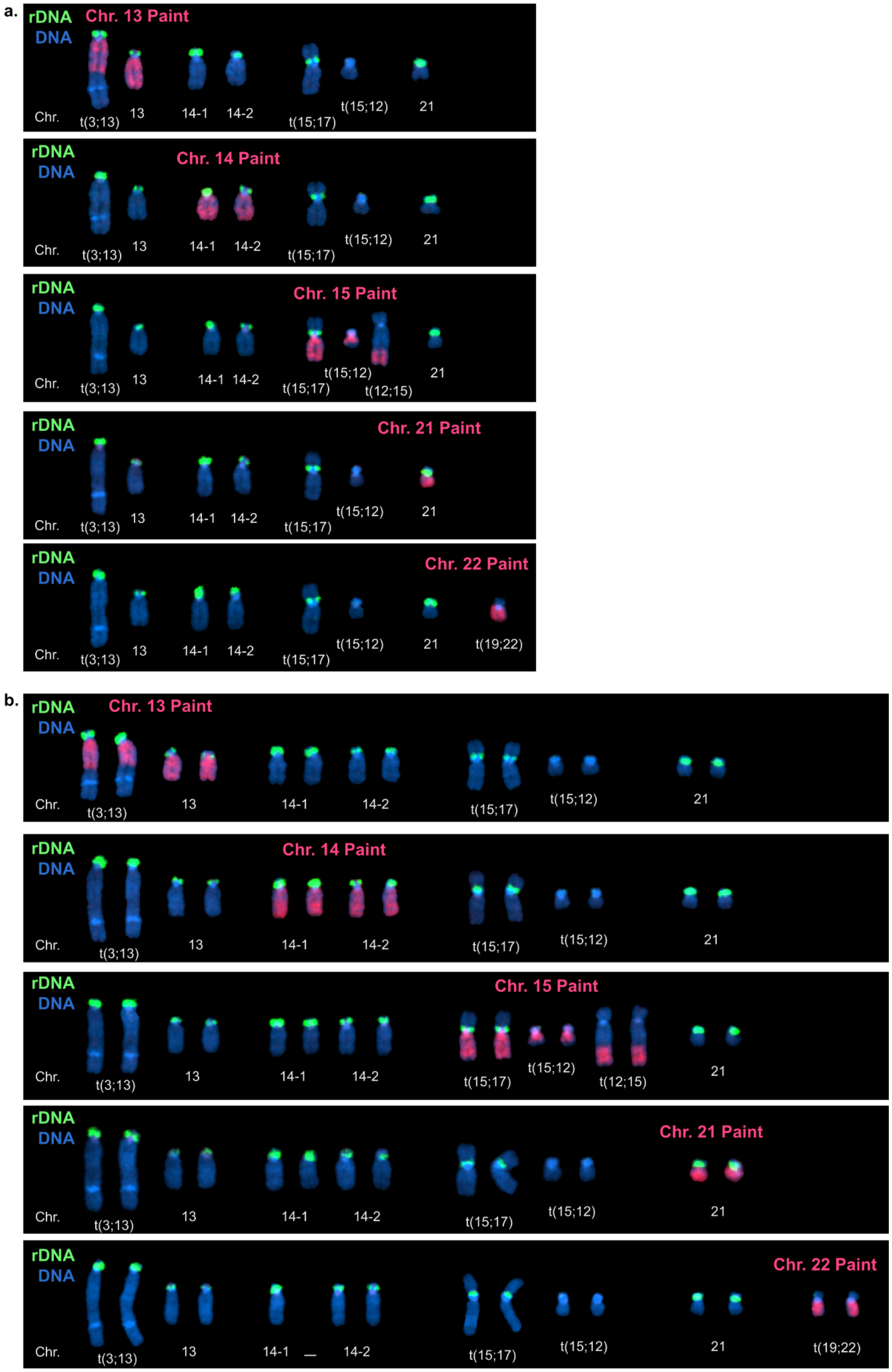
a.
rDNA
DNA
Chr. 13 Paint
Chr. 14 Paint
Chr. 15 Paint
Chr. 21 Paint
Chr. 22 Paint
Chr.
t(3;13)
13
14-1
14-2
t(15;17)
t(15;12)
t(12;15)
21
t(19;22)
b.
rDNA
DNA
Chr. 13 Paint
Chr. 14 Paint
Chr. 15 Paint
Chr. 21 Paint
Chr. 22 Paint
Chr.
t(3;13)
13
14-1
14-2
t(15;17)
t(15;12)
t(12;15)
21
t(19;22)

**Supplementary Fig. 45: Identities of 45S rDNA-containing chromosomes in HG008-T genome.** Representative karyograms of rDNA-containing chromosomes from hypodiploid (a) and genome-doubled (b) cells. Mitotic chromosome spreads from these cells were labeled by FISH with 45S rDNA probe (green) and indicated whole chromosome paints for five human acrocentric chromosomes – 13,14,15, 21 and 22 (red). DNA was counterstained with DAPI (blue). a. Only seven rDNA arrays were preserved in the hypodiploid tumor genome after loss of one copy of chromosome 21, one copy of chromosome 22, and deletion of acrocentric p-arm on another copy of chromosome 22 due to t(19;22) translocation. Rearranged chromosomes t(3;13) and t(15;17) retained rDNA arrays. b. This arrangement was preserved in cells after the genome doubling, with occasional loss of individual chromosomes.

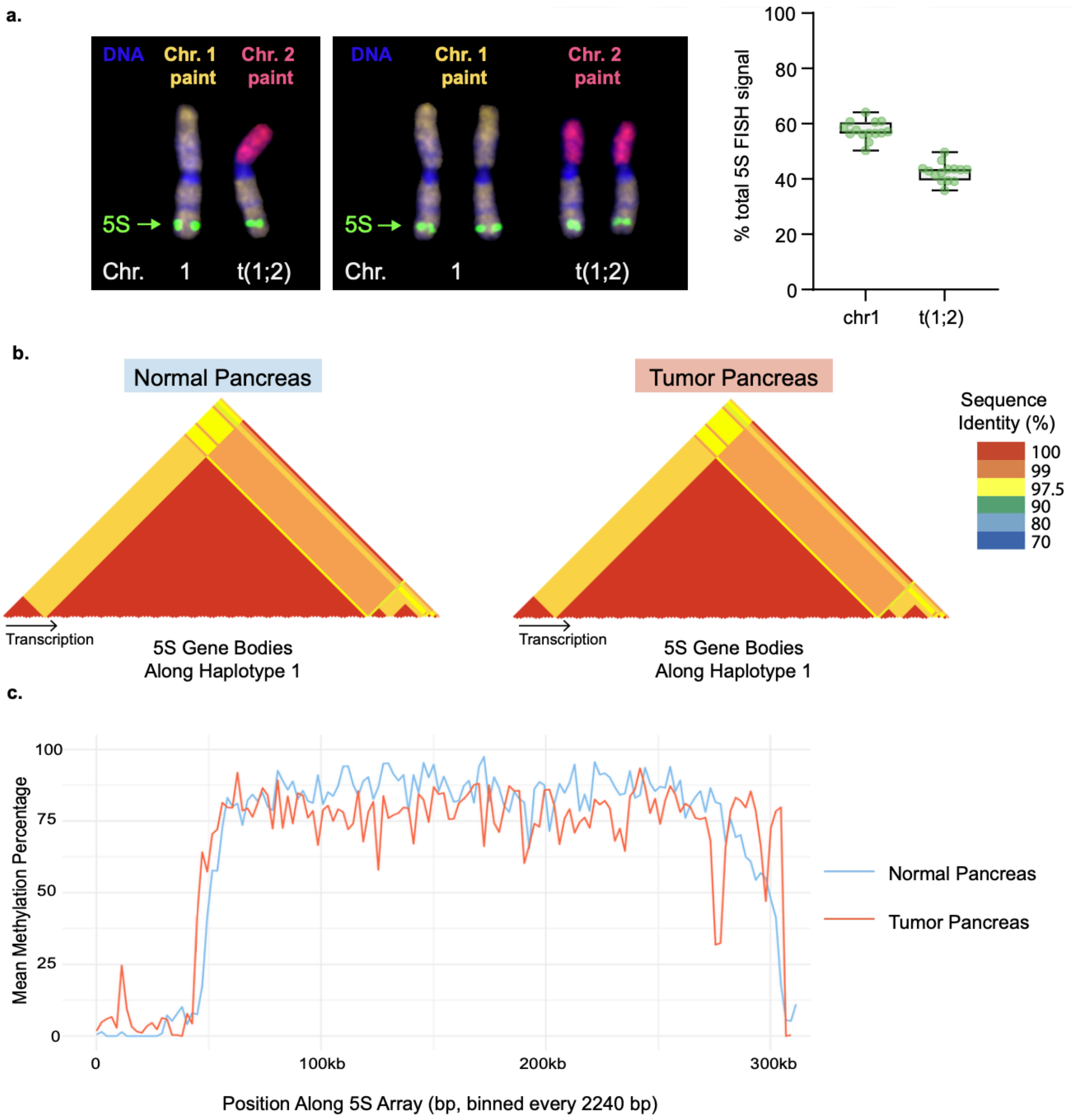
a.
DNA
Chr. 1 paint
Chr. 2 paint
5S
Chr. 1 t(1;2)
% total 5S FISH signal
100
80
60
40
20
0
chr1
t(1;2)
b.
Normal Pancreas
Tumor Pancreas
Sequence Identity (%)
100
99
97.5
90
80
70
Transcription
5S Gene Bodies Along Haplotype 1
c.
Mean Methylation Percentage
100
75
50
25
0
0
100kb
200kb
300kb
Position Along 5S Array (bp, binned every 2240 bp)
Normal Pancreas
Tumor Pancreas

**Supplementary Fig. 46: 5S Ribosomal DNA in Pancreatic Cancer Context.** a. Representative images of chromosomes containing 5S rRNA gene arrays from hypodiploid (left panel) and genome doubled (right panel) HG008-T cells. Chromosome spreads were labeled by FISH with 5S gene probe (green) and whole chromosome paints for chromosome 1 (yellow) and chromosome 2 (red). DNA was counterstained with DAPI (blue). 5S arrays were retained on both the intact chromosome 1 and the translocated chromosome t(1;2), and this arrangement was preserved after the genome doubling, with occasional loss of one copy of t(1;2) chromosome. The 5S array (haplotype 1) on the intact chromosome 1 is slightly larger than the one on the translocated copy (on average 57.5% vs 42.5% of the total 5S FISH signal). The boxplot on the right shows means with standard deviations from 14 hypodiploid spreads. b. Pairwise sequence identity of 5S genes within the array is high within normal and tumor haplotype 1 from HG008-T. Triangular heatmaps for normal (left) and tumor (right) show pairwise sequence identity, with values ranging from 100% to 97%, between every pair of 5S genes within a single array, ordered by array position in direction of transcription (position 1 = telomere proximal and first transcribed, established by BLAST strand orientation). c. Mean percent methylation was derived from ONT reads mapped to the respective assembly, binned in 2240 bp windows to represent the full 5S rRNA monomers, along the 5S rDNA array on haplotype 1 for HG008 pancreatic normal (blue) and tumor (red). Genomic position is rebased to the start of each array and oriented in the direction of transcription. One caveat is that copy number and DNA methylation analysis both require different computational approaches for the 45S and 5S arrays.

a.

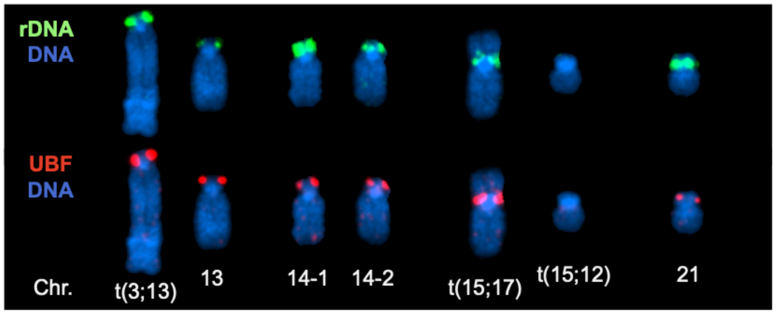


b.

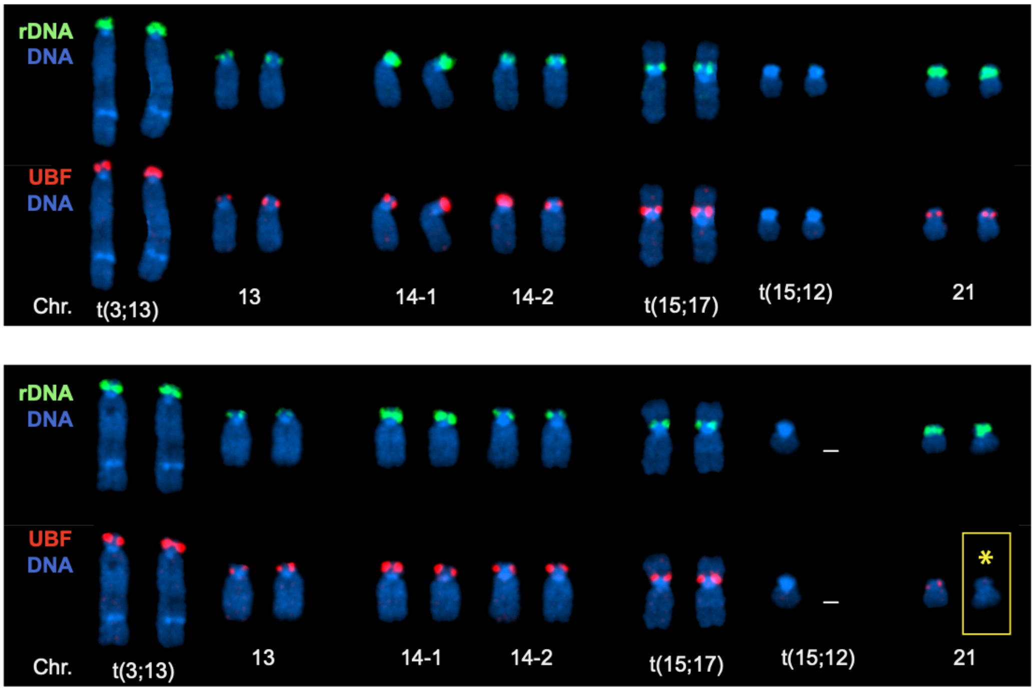


**Supplementary Fig. 47: 45S rDNA activity in HG008-T genome.** Representative karyograms of 45S rDNA-containing chromosomes labeled by immuno-FISH with rDNA probe (green, top panels) and an antibody against rDNA-specific transcription factor UBF (red, bottom panels). DNA was counterstained with DAPI (blue). a. In chromosomal spreads from hypodiploid cells all rDNA arrays were UBF-positive (active). b. In genome-doubled cells most rDNA arrays remained UBF-positive (upper panel), but in some cells one of the rDNA arrays on chromosome 21 became UBF-negative, indicative of transcriptional silencing (lower panel, with asterisk and outlined in gold).

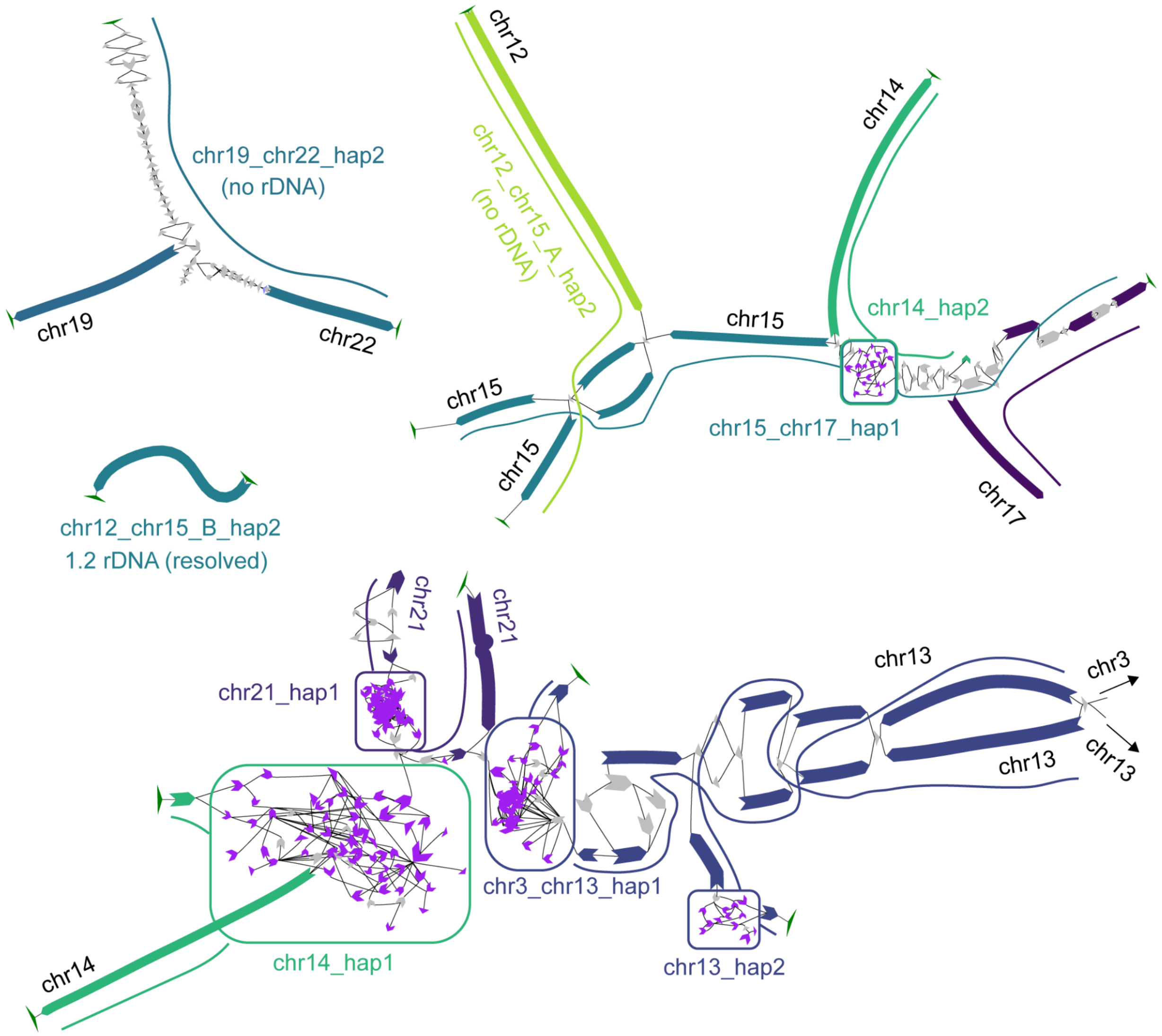


**Supplementary Fig. 48: Assembly graph of the tumor sample showing acrocentric chromosomes with rDNAs.** Nodes are colored and labeled based on alignments to CHM13. rDNA nodes are shown in purple, with tangles identified in rounded squares. One tangle (tangle 0 in Supplementary Table 11) is shared among chr14_hap2 and chr15_chr17_hap1. The rest of the tangles in the lower part were assignable to one chromosomal haplotype based on the hi-c based scaffolding.

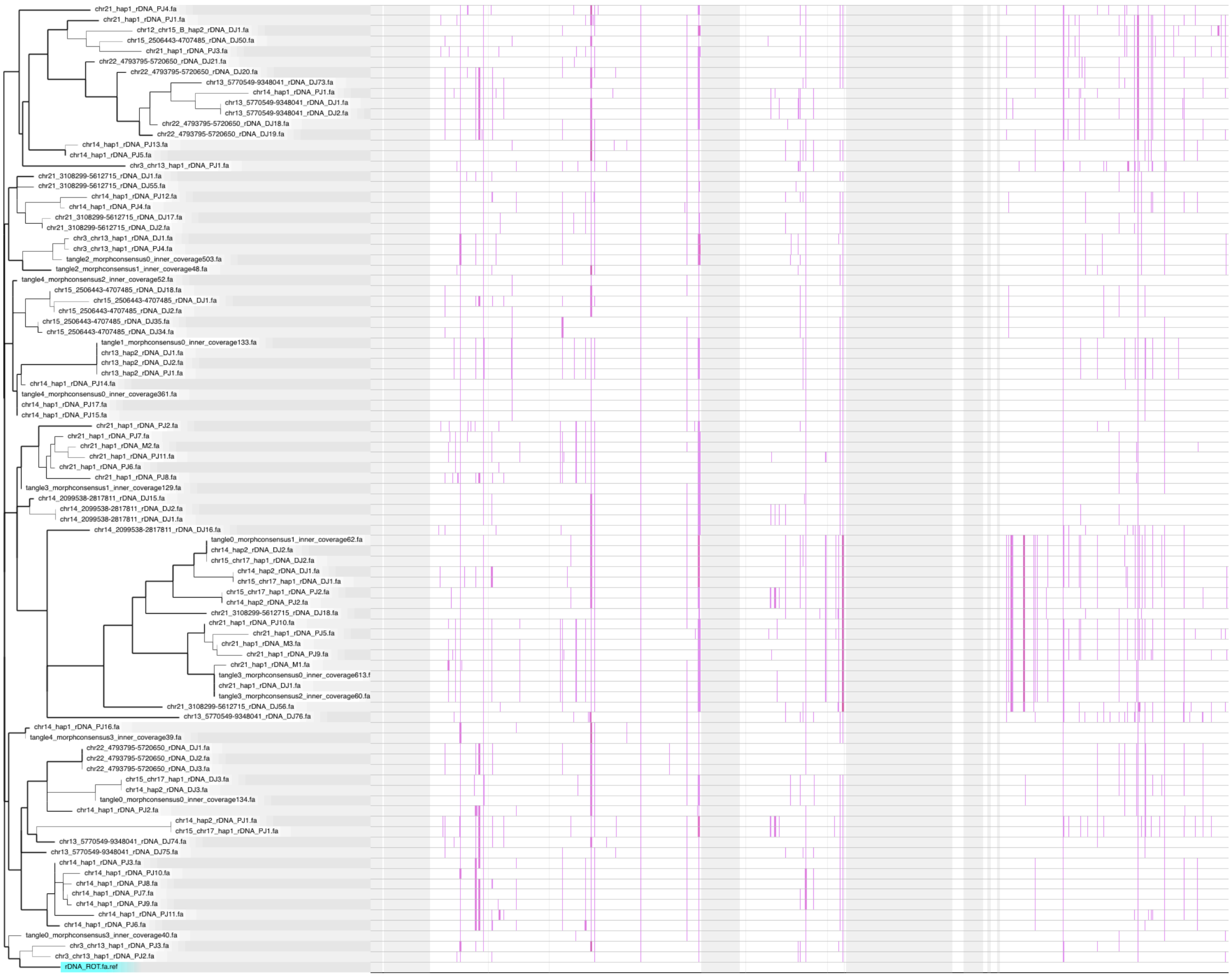
chr21_hap1_rDNA_PJ4.fa
chr21_hap1_rDNA_PJ1.fa
chr12_chr15_B_hap2_rDNA_DJ1.fa
chr15_2506443-4707485_rDNA_DJ50.fa
chr21_hap1_rDNA_PJ3.fa
chr22_4793795-5720650_rDNA_DJ21.fa
chr22_4793795-5720650_rDNA_DJ20.fa
chr13_5770549-9348041_rDNA_DJ73.fa
chr14_hap1_rDNA_PJ1.fa
chr13_5770549-9348041_rDNA_DJ1.fa
chr13_5770549-9348041_rDNA_DJ2.fa
chr22_4793795-5720650_rDNA_DJ18.fa
chr22_4793795-5720650_rDNA_DJ19.fa
chr14_hap1_rDNA_PJ13.fa
chr14_hap1_rDNA_PJ5.fa
chr3_chr13_hap1_rDNA_PJ1.fa
chr21_3108299-5612715_rDNA_DJ1.fa
chr21_3108299-5612715_rDNA_DJ55.fa
chr14_hap1_rDNA_PJ12.fa
chr14_hap1_rDNA_PJ4.fa
chr21_3108299-5612715_rDNA_DJ17.fa
chr21_3108299-5612715_rDNA_DJ2.fa
chr3_chr13_hap1_rDNA_DJ1.fa
chr3_chr13_hap1_rDNA_PJ4.fa
tangle2_morphconsensus0_inner_coverage503.fa
tangle2_morphconsensus1_inner_coverage48.fa
tangle4_morphconsensus2_inner_coverage52.fa
chr15_2506443-4707485_rDNA_DJ18.fa
chr15_2506443-4707485_rDNA_DJ1.fa
chr15_2506443-4707485_rDNA_DJ2.fa
chr15_2506443-4707485_rDNA_DJ35.fa
chr15_2506443-4707485_rDNA_DJ34.fa
tangle1_morphconsensus0_inner_coverage133.fa
chr13_hap2_rDNA_DJ1.fa
chr13_hap2_rDNA_DJ2.fa
chr13_hap2_rDNA_PJ1.fa
chr14_hap1_rDNA_PJ14.fa
tangle4_morphconsensus0_inner_coverage361.fa
chr14_hap1_rDNA_PJ17.fa
chr14_hap1_rDNA_PJ15.fa
chr21_hap1_rDNA_PJ2.fa
chr21_hap1_rDNA_PJ7.fa
chr21_hap1_rDNA_M2.fa
chr21_hap1_rDNA_PJ11.fa
chr21_hap1_rDNA_PJ6.fa
chr21_hap1_rDNA_PJ8.fa
tangle3_morphconsensus1_inner_coverage129.fa
chr14_2099538-2817811_rDNA_DJ15.fa
chr14_2099538-2817811_rDNA_DJ2.fa
chr14_2099538-2817811_rDNA_DJ1.fa
chr14_2099538-2817811_rDNA_DJ16.fa
tangle0_morphconsensus1_inner_coverage62.fa
chr14_hap2_rDNA_DJ2.fa
chr15_chr17_hap1_rDNA_DJ2.fa
chr14_hap2_rDNA_DJ1.fa
chr15_chr17_hap1_rDNA_DJ1.fa
chr15_chr17_hap1_rDNA_PJ2.fa
chr14_hap2_rDNA_PJ2.fa
chr21_3108299-5612715_rDNA_DJ18.fa
chr21_hap1_rDNA_PJ10.fa
chr21_hap1_rDNA_PJ5.fa
chr21_hap1_rDNA_M3.fa
chr21_hap1_rDNA_PJ9.fa
chr21_hap1_rDNA_M1.fa
tangle3_morphconsensus0_inner_coverage613.f
chr21_hap1_rDNA_DJ1.fa
tangle3_morphconsensus2_inner_coverage60.fa
chr21_3108299-5612715_rDNA_DJ56.fa
chr13_5770549-9348041_rDNA_DJ76.fa
chr14_hap1_rDNA_PJ16.fa
tangle4_morphconsensus3_inner_coverage39.fa
chr22_4793795-5720650_rDNA_DJ1.fa
chr22_4793795-5720650_rDNA_DJ2.fa
chr22_4793795-5720650_rDNA_DJ3.fa
chr15_chr17_hap1_rDNA_DJ3.fa
chr14_hap2_rDNA_DJ3.fa
tangle0_morphconsensus0_inner_coverage134.fa
chr14_hap1_rDNA_PJ2.fa
chr14_hap2_rDNA_PJ1.fa
chr15_chr17_hap1_rDNA_PJ1.fa
chr13_5770549-9348041_rDNA_DJ74.fa
chr13_5770549-9348041_rDNA_DJ75.fa
chr14_hap1_rDNA_PJ3.fa
chr14_hap1_rDNA_PJ10.fa
chr14_hap1_rDNA_PJ8.fa
chr14_hap1_rDNA_PJ7.fa
chr14_hap1_rDNA_PJ9.fa
chr14_hap1_rDNA_PJ11.fa
chr14_hap1_rDNA_PJ6.fa
tangle0_morphconsensus3_inner_coverage40.fa
chr3_chr13_hap1_rDNA_PJ3.fa
chr3_chr13_hap1_rDNA_PJ2.fa
rDNA_ROT.fa.ref

**Supplementary Fig. 49: Phylogenetic tree and clustering rDNA morphs.** Tree was built using parsnp and visualized with gingr. Morphs are labeled based on their origin. Assembled morphs in v3.3 assembly are named after the sequence name (chr_hap), with the Ns found after  the distal junction (DJ) or from the proximal junction (PJ). Predicted morphs from ribotin are named after the tangle (0-4), as shown in Supplementary Table 11, with morph consensus num. and coverage. Morphs from CHM13 are labeled after chr_start-end position, as found in chm13v2.0 counted from the one next to the DJ. Only full rDNA units (>90%) with sufficient coverage (>30) are included in the analysis. The first column is the 45S region. Gray areas indicate variable repeats. Purple bars indicate SNPs compared to the reference rDNA sequence, KY962518.1, rotated to begin at position 41,096 to match the start site after the DJ. The rDNA morphs formed a single cluster on each of the chromosomes 13-2 and t(15;12), suggesting localized propagation of the rDNAs in the tumor (Supplementary Table 11).  Assembled rDNAs on chr14-2 and t(15;17) clustered together with the predicted morph in three clusters, indicating strong homology between these two regions and possible gene conversion events among the rDNAs. Chromosomes t(3;13), 14-1 and 21 showed more diverse clusters (3, 4 and 7), yet localized at the leaf level. Some morphs were branched out closely from morphs found on the same homologous chromosome of CHM13, however at different degrees, suggesting some morphs underwent a more rapid turnover gaining more mutations (i.e. chromosome 14-1), while some preserved its form (i.e. one cluster in 21-1 and t(15;12)).

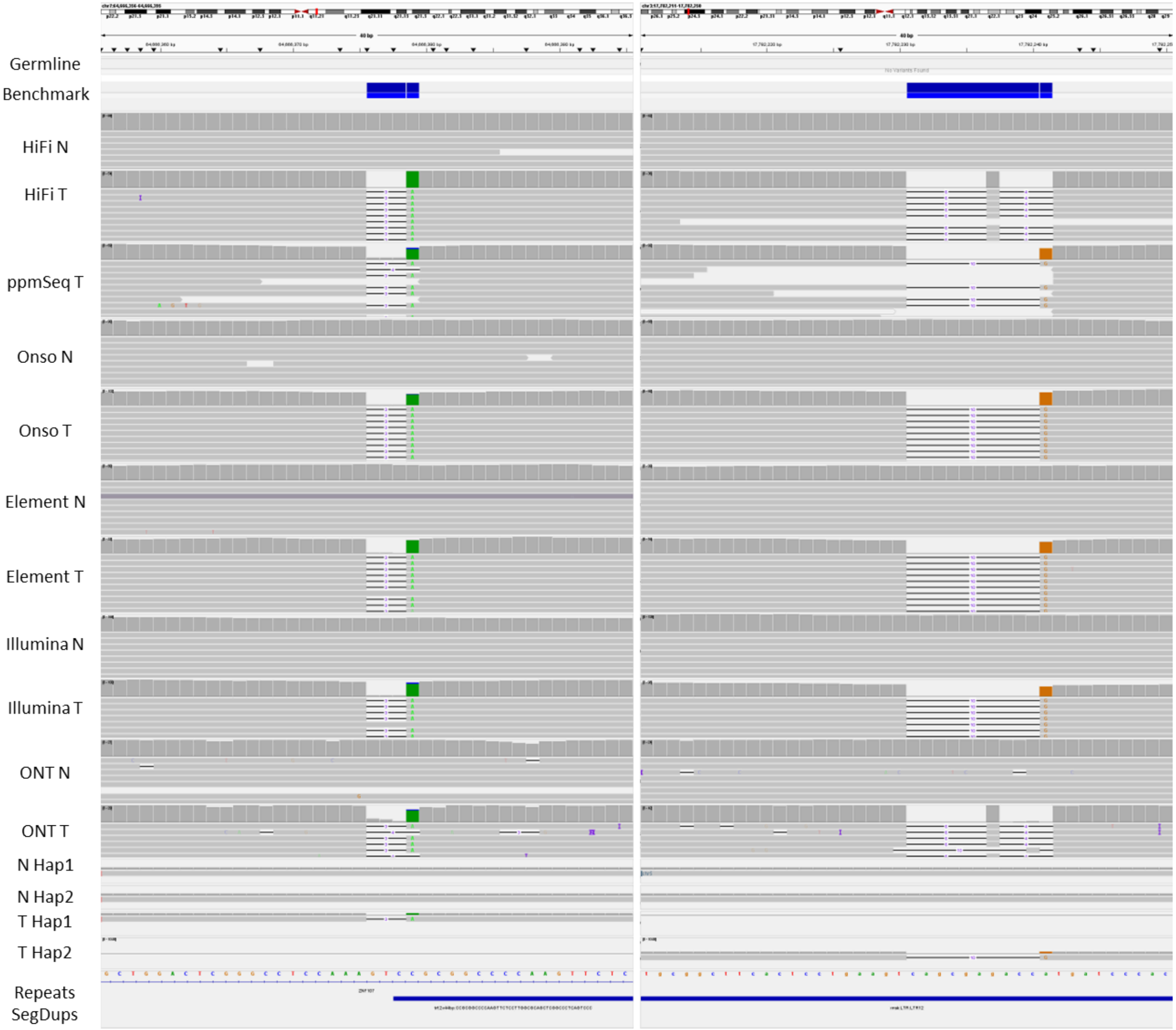


**Supplementary Fig. 50: Other types of complex variants.** These are two examples of complex variants though other events include MNVs. First, chr7:64666476 shows 4 bp replaced by 1 bp as seen in the alignments and assembly. Another example is at chr3:17782231 that has 11 bp replaced by 1 bp.

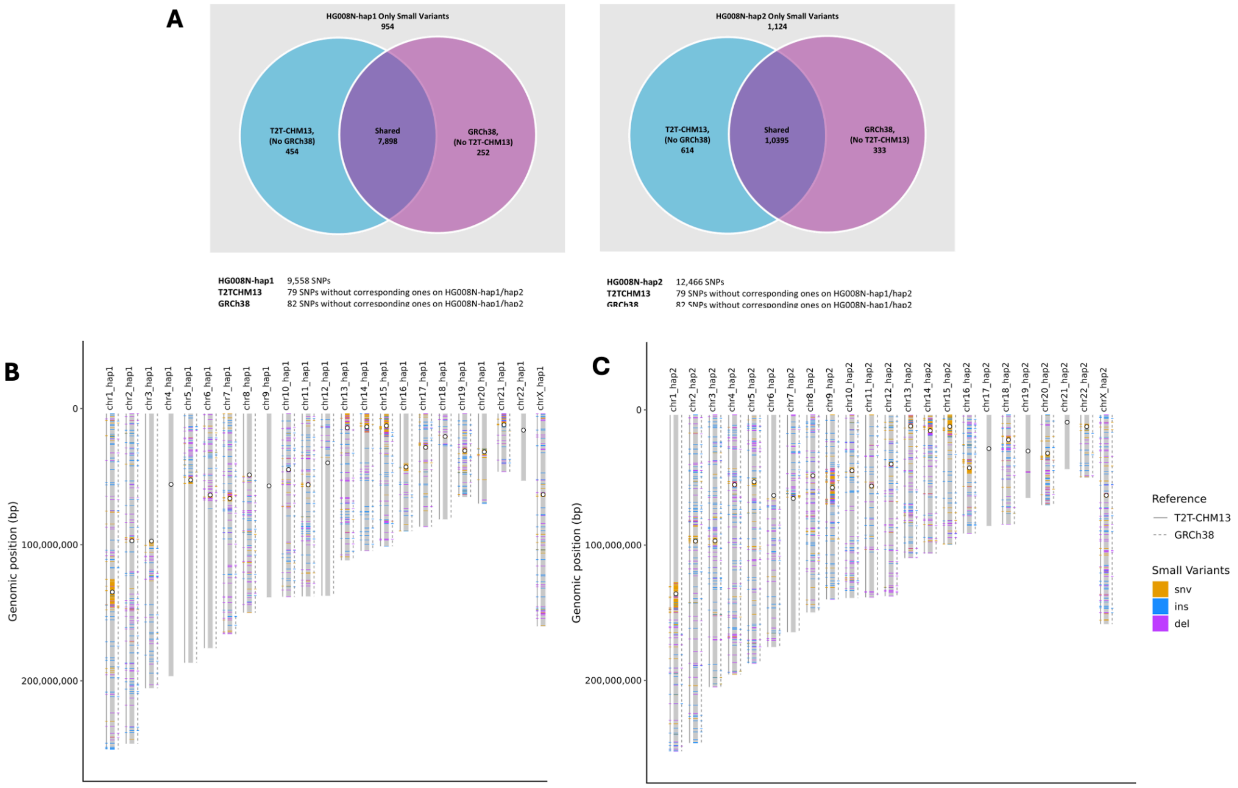


**Supplementary Fig 51: Benchmark small variants included in normal DSA, GRCh38, and T2T-CHM13v2.0.** (A) Venn diagram of small variants across references, with variants unique to the DSA, variants that could be lifted to both GRCh38 and T2T-CHM13v2, and variants that could only be lifted and represented on one. (B) and (C) Chromosomal distribution of small variants that are not shared across all three references. The center of the centromere is marked with an open circle.

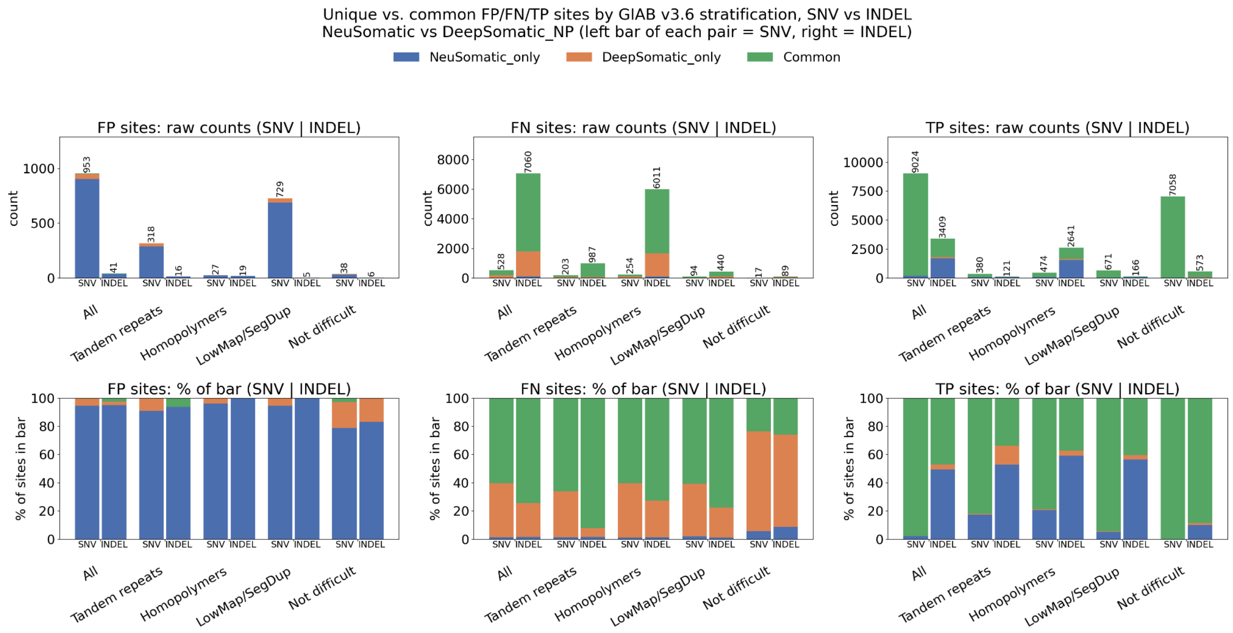
Unique vs. common FP/FN/TP sites by GIAB v3.6 stratification, SNV vs INDEL
NeuSomatic vs DeepSomatic_NP (left bar of each pair = SNV, right = INDEL)
NeuSomatic_only
DeepSomatic_only
Common
FP sites: raw counts (SNV | INDEL)
FN sites: raw counts (SNV | INDEL)
TP sites: raw counts (SNV | INDEL)
FP sites: % of bar (SNV | INDEL)
FN sites: % of bar (SNV | INDEL)
TP sites: % of bar (SNV | INDEL)
count
% of sites in bar
SNV
INDEL
All
Tandem repeats
Homopolymers
LowMap/SegDup
Not difficult

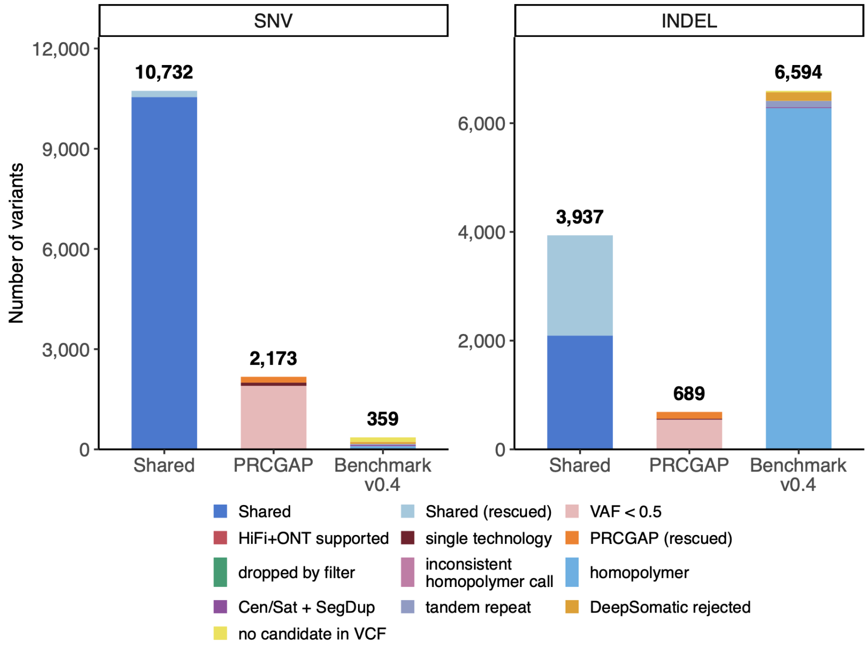
SNV
INDEL
Number of variants
10,732
2,173
359
3,937
689
6,594
Shared
PRCGAP
Benchmark v0.4
Shared
Shared (rescued)
VAF < 0.5
HiFi+ONT supported
single technology
PRCGAP (rescued)
dropped by filter
inconsistent homopolymer call
homopolymer
Cen/Sat + SegDup
tandem repeat
DeepSomatic rejected
no candidate in VCF

**Supplementary Fig. 52: Comparison of ensemble approaches to v0.4 small variant benchmark.** (a) v0.4 GRCh38 benchmark includes repetitive regions missing from previous consensus DeepSomatic and Neusomatic benchmarks. After removing VAF<10% to reduce FPs due to subclonal variation, very few FPs are in common, whereas many FNs are in common, primarily because the assembly-based benchmark includes harder variants in homopolymers, TRs, and segmental duplications. Also, the NeuSomatic vcf seems more tuned for sensitivity whereas the DeepSomatic vcf is tuned for specificity. This does not count the new benchmark variants that couldn't be lifted reliably to GRCh38, which would essentially be additional FNs. (b) v0.4 DSA-based benchmark includes many indels missing from consensus PRCGAP calls, particularly in homopolymers.

(a)

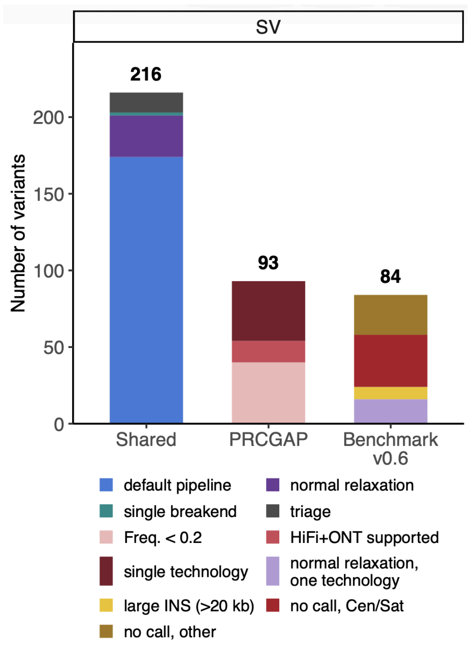


(b)

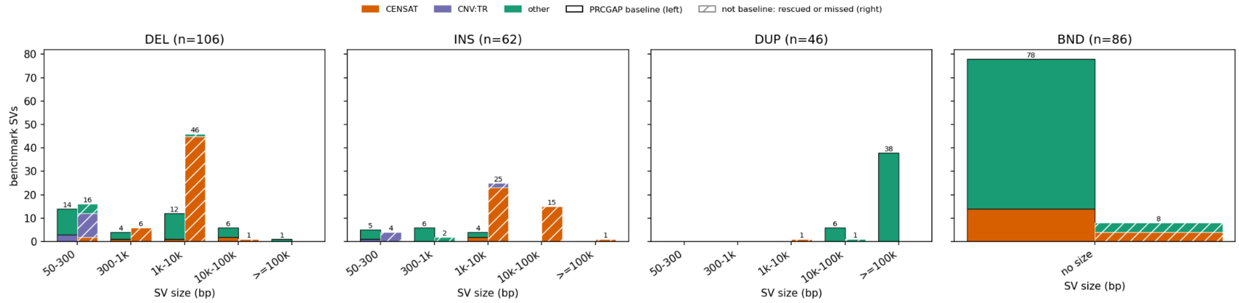


**Supplementary Fig. 53: Comparison of PRCGAP SV calls from long reads mapped to DSA vs. our curated v0.6 assembly-based benchmark.** (a) Putative TP, FP, and FN variants categorized. (b) Solid bars are PRCGAP calls with baseline parameters and filtering that match benchmark calls, and hashed bars are benchmark calls that are either filtered in or missing from PRCGAP calls.

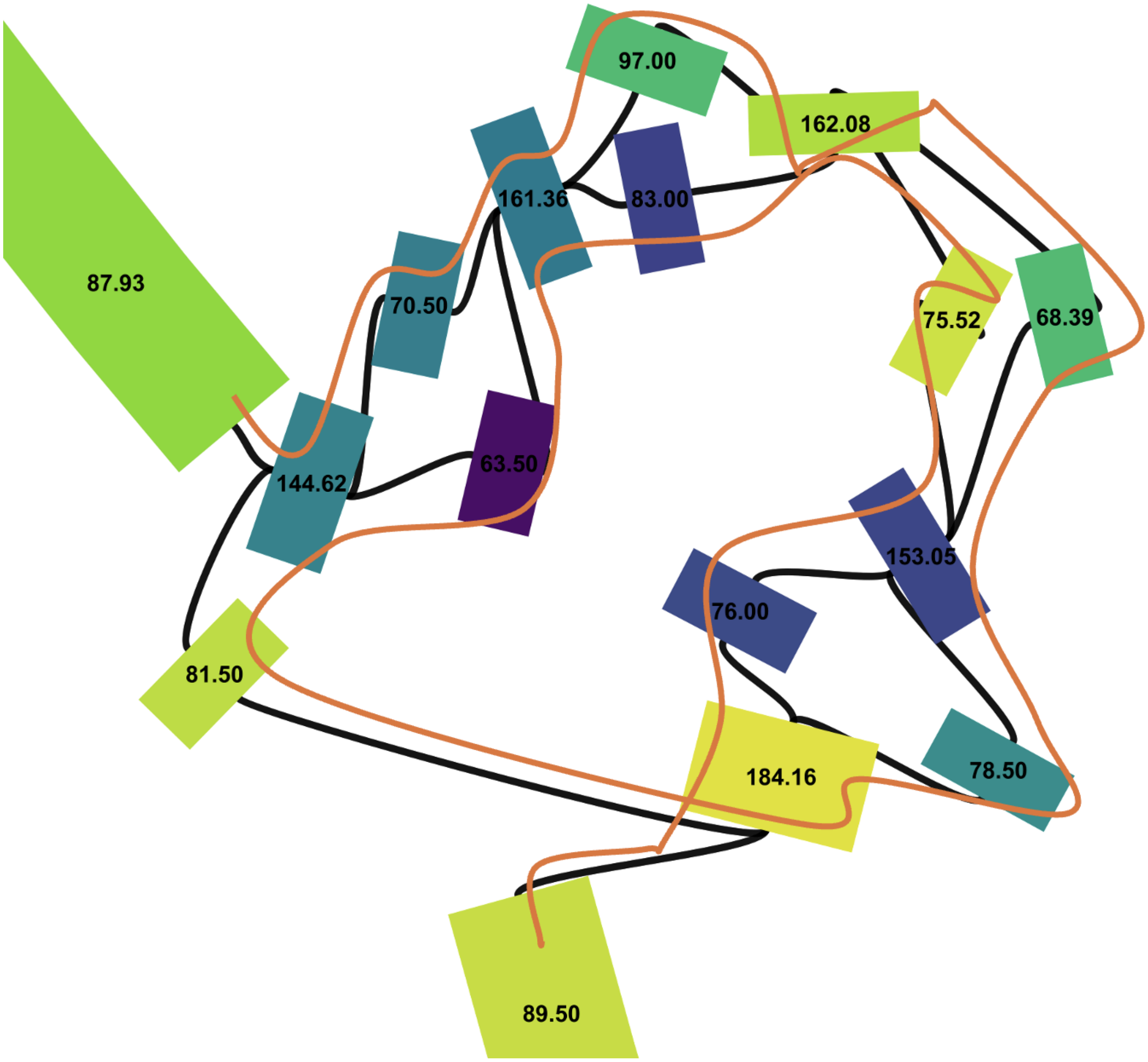


**Supplementary Fig. 54: Example Assembly Graph Manual Resolution.** Example of manual resolution of a 2.98 Mbp tandem duplication at chr7:4824648 (GRCh38), where numbers correspond to node coverage by HiFi and error-corrected ONT reads and orange is the manually chosen path. The nodes in each bubble were randomly assigned to each copy of the duplication so that each node in the bubble was used once and nodes with higher coverage between bubbles were used twice.

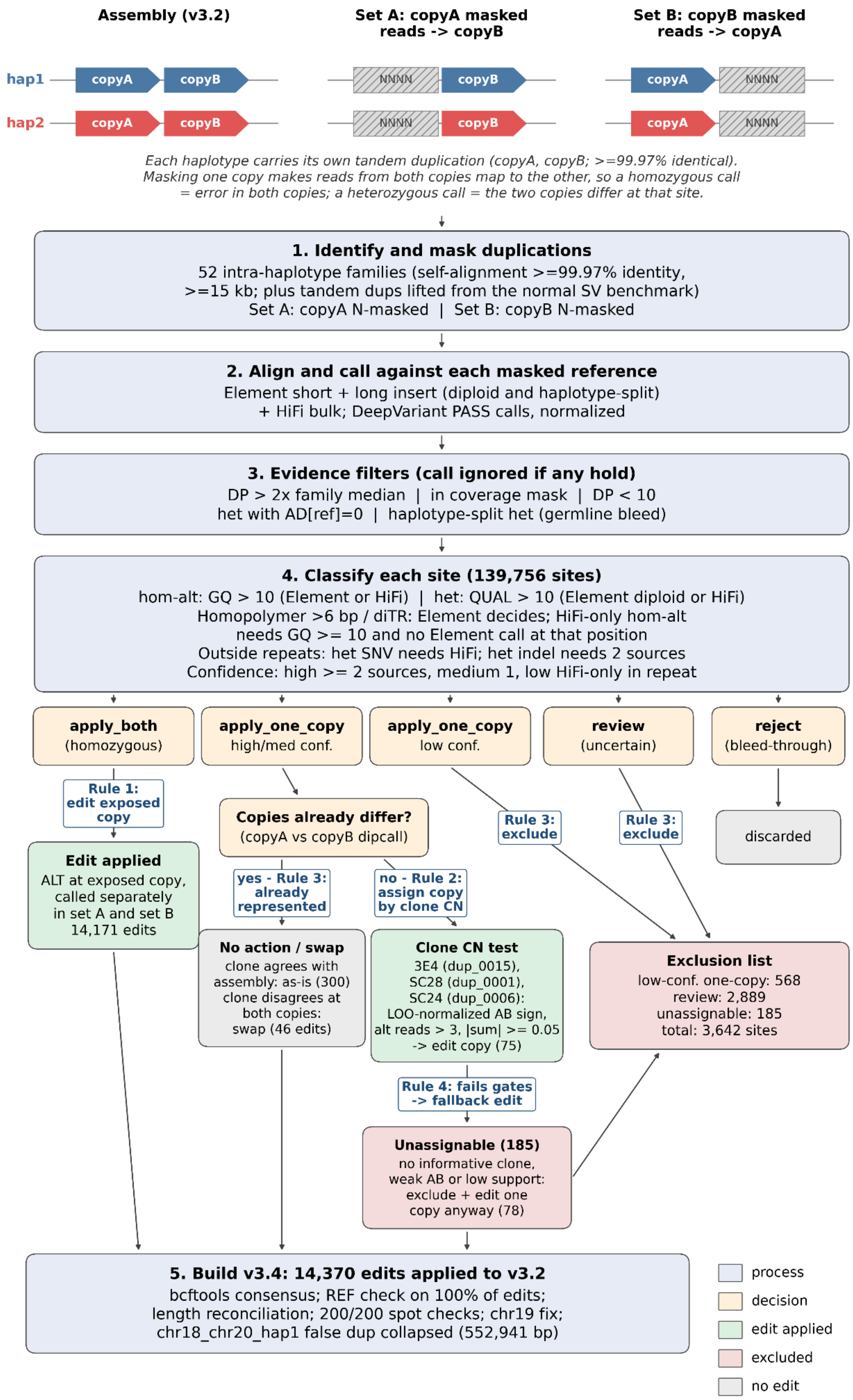
Assembly (v3.2)
Set A: copyA masked reads -> copyB
Set B: copyB masked reads -> copyA
hap1
hap2
copyA
copyB
NNNN
Each haplotype carries its own tandem duplication (copyA, copyB; >=99.97% identical).
Masking one copy makes reads from both copies map to the other, so a homozygous call = error in both copies; a heterozygous call = the two copies differ at that site.
1. Identify and mask duplications
52 intra-haplotype families (self-alignment >=99.97% identity, >=15 kb; plus tandem dups lifted from the normal SV benchmark)
Set A: copyA N-masked | Set B: copyB N-masked
2. Align and call against each masked reference
Element short + long insert (diploid and haplotype-split) + HiFi bulk; DeepVariant PASS calls, normalized
3. Evidence filters (call ignored if any hold)
DP > 2x family median | in coverage mask | DP < 10
het with AD[ref]=0 | haplotype-split het (germline bleed)
4. Classify each site (139,756 sites)
hom-alt: GQ > 10 (Element or HiFi) | het: QUAL > 10 (Element diploid or HiFi)
Homopolymer >6 bp / diTR: Element decides; HiFi-only hom-alt needs GQ >= 10 and no Element call at that position
Outside repeats: het SNV needs HiFi; het indel needs 2 sources
Confidence: high >= 2 sources, medium 1, low HiFi-only in repeat
apply_both (homozygous)
apply_one_copy high/med conf.
apply_one_copy low conf.
review (uncertain)
reject (bleed-through)
Rule 1: edit exposed copy
Copies already differ? (copyA vs copyB dipcall)
Rule 3: exclude
Rule 3: exclude
discarded
Edit applied
ALT at exposed copy, called separately in set A and set B
14,171 edits
yes - Rule 3: already represented
no - Rule 2: assign copy by clone CN
No action / swap
clone agrees with assembly: as-is (300)
clone disagrees at both copies: swap (46 edits)
Clone CN test
3E4 (dup_0015), SC28 (dup_0001), SC24 (dup_0006):
LOO-normalized AB sign, alt reads > 3, |sum| >= 0.05 -> edit copy (75)
Exclusion list
low-conf. one-copy: 568
review: 2,889
unassignable: 185
total: 3,642 sites
Rule 4: fails gates -> fallback edit
Unassignable (185)
no informative clone, weak AB or low support: exclude + edit one copy anyway (78)
5. Build v3.4: 14,370 edits applied to v3.2
bcftools consensus; REF check on 100% of edits; length reconciliation; 200/200 spot checks; chr19 fix; chr18_chr20_hap1 false dup collapsed (552,941 bp)
process
decision
edit applied
excluded
no edit

**Supplementary Fig 55 Custom process for polishing duplications by masking one copy of each duplication at a time.** After identifying duplications and classifying sites for edit into `apply_one_copy` or `apply_both_copies`, the following rules can be applied in the diagramed workflow. *Rule 1:* Every apply_both_copies site was edited to the alternate allele so set A calls edit copyB and set B calls edit copyA. Because a site is reported independently in each set, a base error shared by both copies is called and corrected in both, whereas an error confined to one copy produces a homozygous call only in the set exposing the erroneous copy so is corrected only there. We confirmed this behaviour after the build by lifting each *Rule 1* edit onto the other copy through a base-exact copyA/copyB alignment: of 13,816 such edits in the 13 self-alignment duplications, 74.6% had a matching edit on the other copy and 24.0% fell where the other copy already carried the alternate allele in v3.2; 0.7% left the other copy uncorrected because the corresponding call in the other set failed a gate, and these were largely already on the exclusion list. *Rule 2 - copy assignment by clone copy number:* A high- or medium-confidence apply_one_copy site indicates that one copy differs from the other but not which. For three duplications we had a single-cell-derived clone in which one duplicate copy had changed copy number (dup*0015/clone 3E4, dup*0001/clone SC28, dup_0006/clone SC24), which shifts the allele balance of a variant residing on the gained or lost copy. We force-genotyped every candidate site in HiFi and Illumina alignments of all clones with freebayes (--only-use-input-alleles, minimum MAPQ 0), computed allele balance from AD, and normalized each clone's allele balance by subtracting the median of the other clones at the same site. A site was assigned if the informative clone had > 3 alternate-supporting reads and the sum of its normalized Illumina and HiFi allele balances had magnitude ≥ 0.05; the sign, calibrated against 329 sites at which the alternate-carrying copy was independently known from assembly-to-assembly comparison, determined which copy received the edit. Sites also present in the copyA-versus-copyB dipcall comparison, in which the assembly already represents a difference between the copies, were treated as follows: if the clone evidence agreed with the assembly's assignment, no action was taken; if the clone evidence pointed at the opposite copy at both the site and its lifted counterpart in the other set, both edits were applied, which exchanges the alleles between copies without erasing the difference (46 such swaps); otherwise no action was taken. Sites failing the support, magnitude, or assignment tests went to the exclusion list. *Rule 3 - fallback for confident but unassignable sites:* For apply_one_copy sites that failed only the copy-assignment step, the alternate allele was applied to one copy so that the assembly carries the well-supported allele somewhere, while the region remained excluded from the benchmark bed. The edited copy was whichever set's coordinate the site was reported in; when a site was reported in both sets as matching through copyA/copyB alignment within 3 bp for SNVs or 100 bp for indels, only one was edited, and no edit was applied if the counterpart had already been edited by *Rule 1* or *2*.

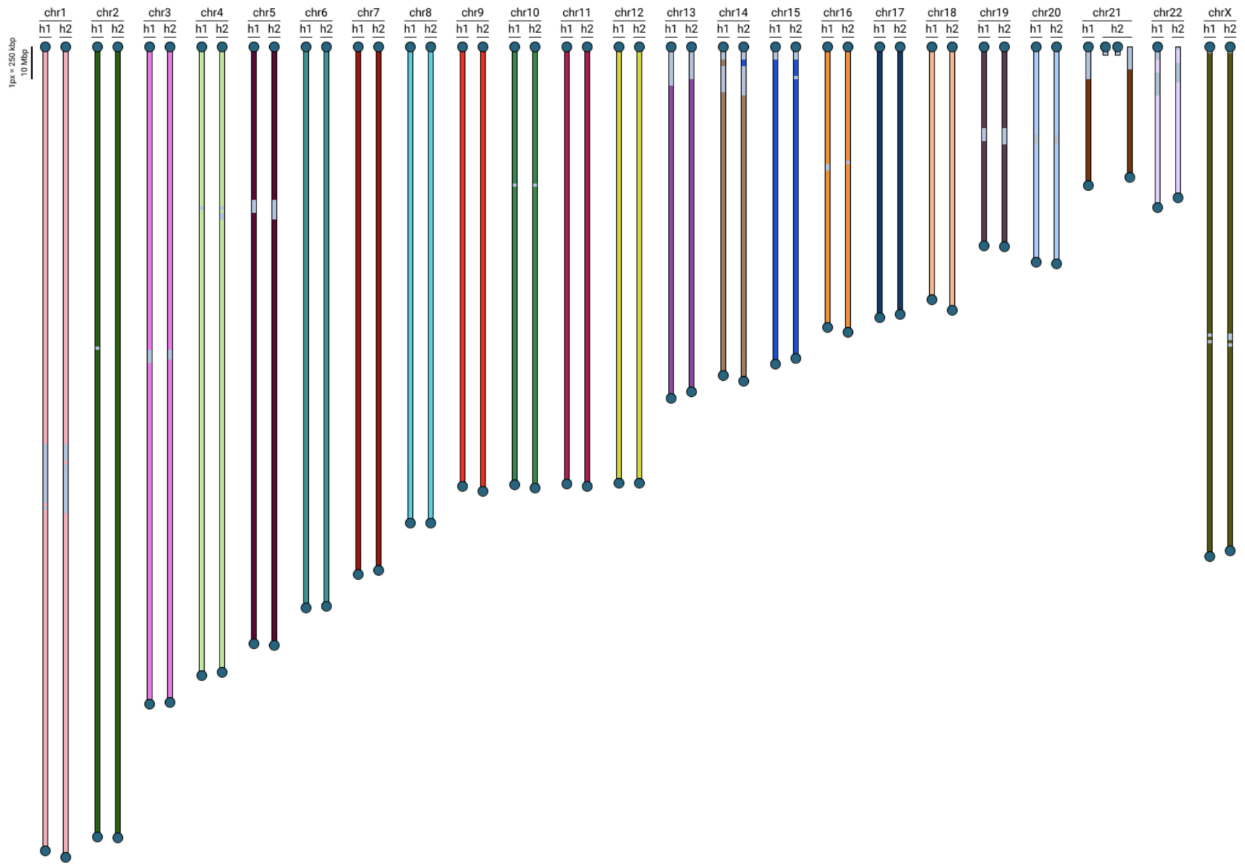


**Supplementary Fig. 56: Karyoscope orientation of HG008Nv6.3 large contigs based on CHM13 k-mers.**
